# Engineered display of ganglioside-sugars on protein elicits a clonally and structurally constrained B cell response

**DOI:** 10.1101/2023.06.03.543556

**Authors:** Lachlan P. Deimel, Xiaochao Xue, Aziz Khan, Lucile Moynie, Charles J. Buchanan, Guoxuan Sun, Ryan McBride, Heiko Schuster, Charles Gauthier, Regis Saliba, Karolis Leonavicus, Leanne Minall, Guillaume Bort, Rebecca A. Russell, Erdinc Sezgin, James C. Paulson, Daniel C. Anthony, Andrew J. Baldwin, James Naismith, Torben Schiffner, Benjamin G. Davis, Quentin J. Sattentau

**Author notes:** Correspondence (QJS), (BGD). S1, Sugar Synthesis and Log Generation; S2, Further Discussion of Structural Analyses of Bar1 Fab; S3, Supplementary Glycan Microarray Document.

## Abstract

Ganglioside sugars, as Tumour-Associated Carbohydrate Antigens (TACAs), are long-proposed targets for vaccination and therapeutic antibody production, but their self-like character imparts immunorecessive characteristics that classical vaccination approaches have to date failed to overcome. One prominent TACA, the glycan component of ganglioside GM3 (GM3g), is over-expressed on diverse tumours. To probe the limits of glycan tolerance, we used protein editing methods to display GM3g in systematically varied non-native presentation modes by attachment to carrier protein lysine sidechains using diverse chemical linkers. We report here that such presentation creates glycoconjugates that are strongly immunogenic in mice and elicit robust antigen-specific IgG responses specific to GM3g. Characterisation of this response by antigen-specific B cell cloning and phylogenetic and functional analyses suggests that such display enables the engagement of a highly restricted naïve B cell class with a defined germline configuration dominated by members of the *IGHV2* subgroup. Strikingly, structural analysis reveals that glycan features appear to be recognised primarily by antibody CDRH1/2, and despite the presence of an antigen-specific Th response and B cell somatic hypermutation, we found no evidence of affinity maturation towards the antigen. Together these findings suggest a ‘reach-through’ model in which glycans, when displayed in non-self formats of sufficient distance from a conjugate backbone, may engage ‘glycan ready’ V-region motifs encoded in the germline. Structural constraints define why, despite engaging the trisaccharide, antibodies do not bind natively-presented glycans, such as when linked to lipid GM3. Our findings provide an explanation for the long-standing difficulties in raising antibodies reactive with native TACAs, and provide a possible template for rational vaccine design against this and other TACA antigens.

**Highlights:**

- GM3g synthetically coupled via a longer, orthogonal (from backbone) glycoconjugate (LOG) presentation format (thioethyl-lysyl-amidine) display elicits high-titre IgG responses in mice.
- The germinal centre experience of LOG glycoconjugate-specific B cell responses is directly influenced by the protein backbone.
- Structural characterisation of the antibody response to LOGs reveals highly restricted germline-encoded glycan-engaging motifs that mediate GM3g recognition.
- Failure of antibodies to bind the native trisaccharide highlights barriers to be overcome for the rational design of anti-TACA antibodies.

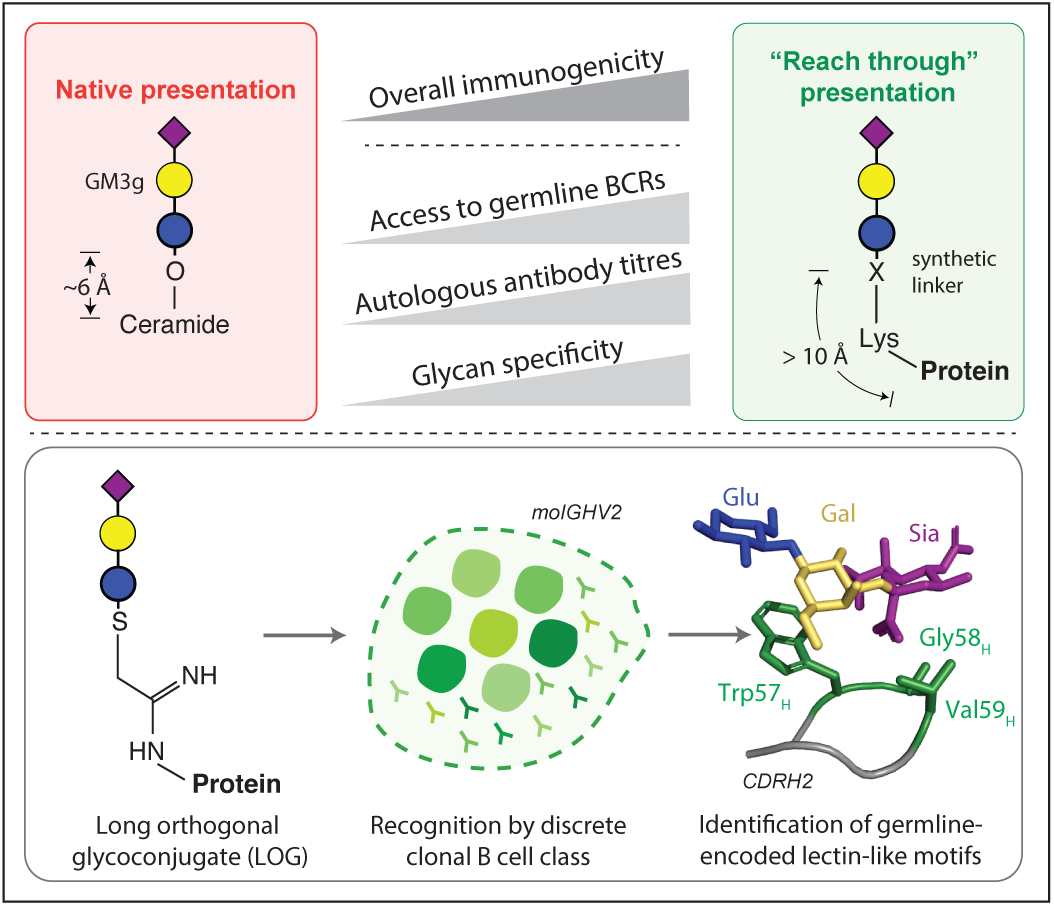

## Introduction

Glycosylation is a widespread enzymatic process with critical roles in modulating and controlling protein and lipid structure, function, and stability. Since mammalian glycans are endogenously added and processed, they and their native glycoconjugates have restricted immunogenicity to avoid autoreactive responses. Immunogenicity may be limited by different mechanisms, including central and peripheral immunological tolerance and intrinsic lack of antigen immunogenicity due to biophysical constraints such as antigen size, charge and accessibility. Many adaptive tolerance mechanisms are well-understood and can be partitioned into several main themes: i) B cell negative selection that eliminates self-glycan-reactive precursors during their development in the bone marrow (*1*); ii) antigen interactions with immunoinhibitory lectins such as CD22 and Siglec-G (*2, 3*), and iii) lack of T cell help to class-switch and affinity-mature B cells (*4, 5*). However, in contrast to self-glycans, antibodies can be readily elicited against foreign glycans, such as those from bacteria, where the same tolerance constraints do not apply. Correspondingly, classical glycoconjugate vaccines artificially display bacteria-derived polysaccharides in a format where the sugar polymer is typically presented in a non-specific, ‘parallel’-mode that the immune system readily responds to (*6, 7*). By contrast, the immunological ‘blind-spots’ present for self-like glycans and glycoconjugates may render the host vulnerable to pathogen attack and other pathology such as cancers that routinely exploit glycosylation processes as an immune evasion tactic. This limits the effective targeting of certain pathology-associated glycans via vaccination.

The selective breaking of B cell tolerance to glycans may therefore have profound utility in certain settings, a highly relevant example being approaches to developing neutralising antibody-based vaccines against HIV-1 (*8*). The HIV-1 envelope glycoprotein (Env), the only target of neutralising antibody elicitation and attack, exploits glycosylation to shield underlying sensitive peptidic epitopes (*9, 10*). However, a small subset of HIV-1-infected individuals develop rare B cell clones that produce potent broadly neutralising antibodies (bNAbs) which interact with glycans or glycan-protein composite epitopes on Env, and mediate broad and potent neutralisation (*11– 14*). In general, these bNAbs have undergone significant somatic hypermutation (SHM) resulting in the progressive accommodation of Env glycans into their respective epitopes via affinity maturation (*15, 16*). Thus, most inferred germline revertants (iGL) of bNAbs fail to recognise natively-glycosylated Env. This creates a formidable ‘moving target’ for vaccine design as the extent of SHM generated during natural infection is difficult to recapitulate by current vaccination approaches. Other approaches may therefore prove necessary for success.

In principle, the odds of achieving a functional anti-self-like glycan response by vaccination may be improved by reducing the need for SHM. In this scenario, the target glycan would ideally be recognised directly by the germline repertoire to initiate a B cell response and so avoid the requirement for extensive SHM for initial target epitope recognition. To our knowledge, this has not yet been observed but would prove widely valuable.

Tumour-associated carbohydrate antigens (TACAs), which are also self-like, may be over-expressed or modified on tumour cells compared to their normal counterparts (*17*). TACAs present even greater vaccine design challenges compared with viral glyco-antigens because i) several major classes of TACAs are presented on glycosphingolipids (including gangliosides) rather than proteins, removing the T helper (Th) component of the adaptive response, and ii) proximity of the carbohydrate to the membrane likely limits B cell receptor accessibility and downstream antibody engagement (*18*). The glycan component of the ganglioside GM3 (GM3g, 3’-*O*-sialyllactosyl) is of particular interest for its elevated expression in melanoma and neuroectodermal tumours (*19, 20*). GM3 has only very weak immunogenicity, with experimental studies showing limited success in eliciting anti-GM3 antibody responses to GM3 or GM3g on various carrier proteins (*21–25*). Early mouse immunisation studies with purified GM3 reported an apparent borderline IgM response (*21, 26*). Notably, conjugation of GM3g to carrier proteins such as Keyhole Limpet Haemocyanin (KLH) or bacterial cells (*21*) provided T cell help to the emerging B cell responses but in these early studies it remained unclear from the serological analyses whether the resulting antibody responses were truly SiaLac/GM3-specific or instead driven in part by artefacts of linker immunogenicity and cross-reactivity in associated assays (*27*).

To further probe our understanding of antibody responses against potentially useful small self-glycans in a manner that might enable vaccination approaches, we combined synthetic glycan-protein engineering with detailed B cell immunological analyses to probe germline-targeted responses leading to the elicitation of glycan-reactive antibodies. We have exploited bespoke chemical linkages of precisely modulated format and length, not found in nature, using an orthogonal/‘side-on’ mode from protein carrier side-chains to probe glycan tolerance mechanisms. The resulting ‘reach through’ presentation by longer, orthogonal glycoconjugates (LOGs) was designed to allow glycans to engage a subset of otherwise inaccessible naïve B cells.

We demonstrate here proof-of-principle of this concept by presenting GM3g on different carrier protein backbones. GM3g-specific IgG titres were readily elicited via a highly restricted clonotypic B cell response using distinct B cell receptor (BCR) heavy chain-mediated glycan recognition. Antibodies binding GM3g do not react with GM3 itself confirming the key role that the presentation of the glycan plays. These data not only provide a rational basis for the key role of glycan presentation in the specificity of corresponding B cell clones elicited, but also represent the first evidence of a TACA-directed germline-targeting immunogen with implications for the future design of glycan reactive antibody-based vaccine approaches.

## Results

### Differing presentation of GM3g modulates B cell immunogenicity

GM3 presents its glycan (GM3g) (**Fig 1a**) natively at short distance (estimated at 6 Å based on native *O*-glycoside, three-bond *O*-hydroxymethyl spaced display from the head group) from its native macromolecular (lipid membrane) assembly surface.

**Figure 1:**
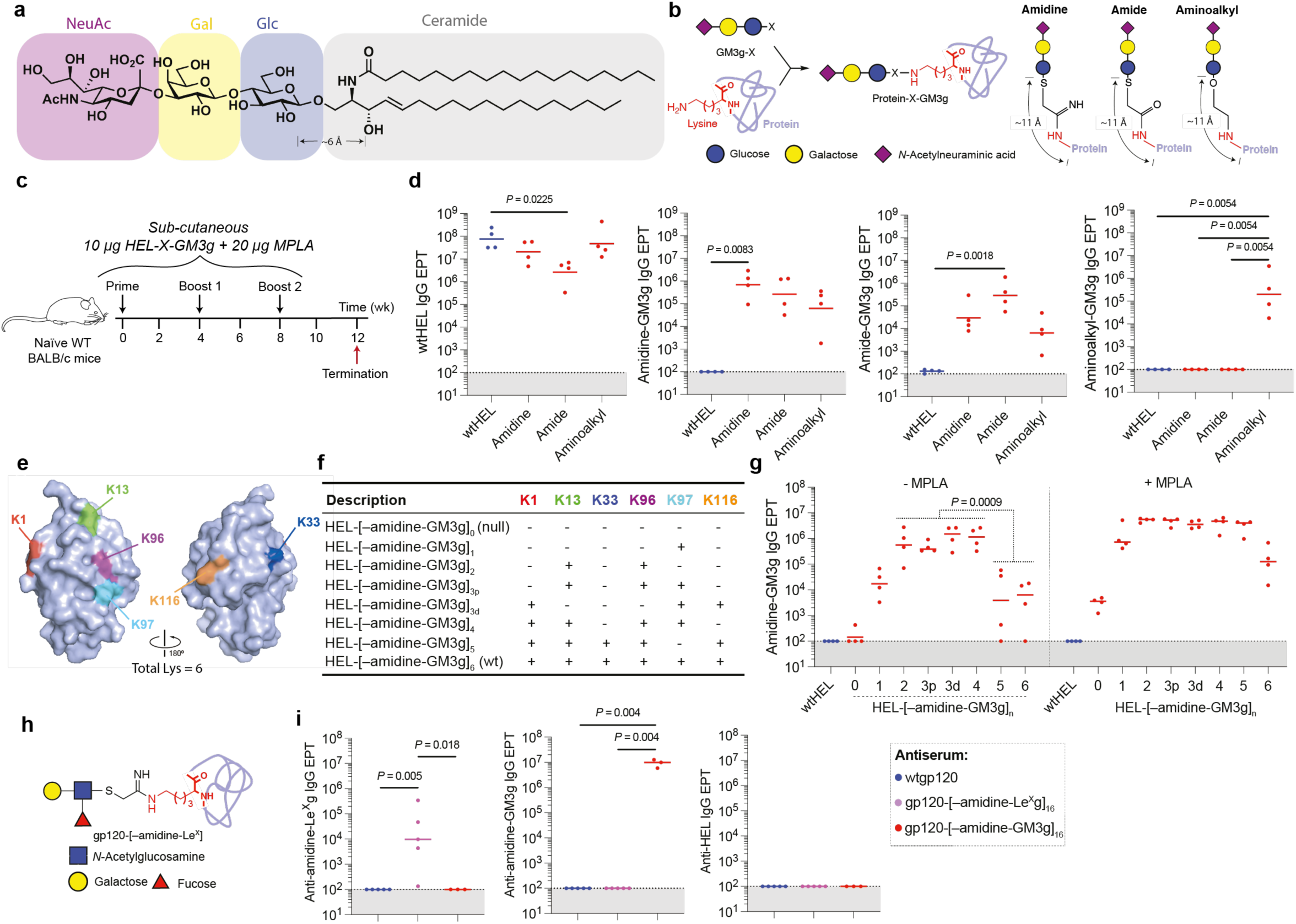
Immunogenicity of semi-synthetic, non-native GM3g-based LOGs in mice. **(a)** Native mammalian ganglioside and TACA, GM3. **(b)** ‘Tag-and-modify’ approaches to chemically coupling GM3g to lysine sidechains of diverse protein carriers. **(c)** Immunisation schedule. **(d)** Terminal LOG-specific IgG endpoint titers of immunized mice. **(e,f)** Position of lysine residues on HEL (PDB: 193L) and their select substitution to arginine (‘-’) in mutant set. **(g)** Evaluation of autologous LOG-specific IgG post-immunisation. **(h)** gp120-[–amidine-Le^X^] LOG design. **(i)** Terminal endpoint titers. Data were compared via Dunn’s test. HEL mutant Ig titres were clustered into low (0–1), medium (2–4) and high (5–6) glycan occupancy and pairwise compared.

We first chose to interrogate the inherent immunogenicity in mice of natively-presented GM3g in the context of intact GM3 lipid. Assembly of GM3-bearing liposomes (PG:PC:Chol:GM3 = 39:39:19:3) created an appropriate macromolecular assembly bearing multi-copy GM3g (**Fig S1a**). Following immunisation of WT BALB/c mice formulated with the TLR-4-agonist-based adjuvant Monophosphoryl-Lipid A (MPLA) (Mata-Haro et al., 2007), antisera against both the GM3-containing and GM3-free control liposome displayed modest IgM reactivity with ceramide (median EPT across groups of 2,156), and similarly low anti-GM3 IgM titres (EPT = 492) (**Fig S1b–d**). These findings are consistent with non-specific IgM binding and the absence of specific GM3g-binding antibodies, reflecting the low-affinity, high-avidity nature of IgM in ELISA formats (*29, 30*). Unsurprisingly, in the absence of T cell help (classically provided by protein in the antigenic complex), antigen-specific IgG was not detected against either ceramide or GM3 (**Fig S1e,f**). These data confirm the profoundly limited immunogenicity of GM3g in this macromolecular format.

We next explored an alternative non-native macromolecular assembly upon which to display GM3g. Orthogonal display on macromolecular protein scaffolds has the potential to mimic membrane-like multi-copy GM3g display, yet allowing control of copy-number density, site-specific conjugation and, critically, distance from the surface in terms of longer orthogonal display. The use of precise protein-editing methods via lysine (Lys)-selective (*31*) ‘tag-and-modify’ methods (*32*) allows GM3g presentation in diverse protein scaffolds (**Fig 1b**). Synthetic, protein-compatible methods were developed that accessed three presentation modes that were systematically varied for both O- *vs* S- glycoside display via different amidine [– C(NH)NH–], amide [–C(O)NH–], or aminoalkyl [–(CH_2_)_2_NH–] linkers all at a similar, extended nine/ten-bond length, corresponding to ∼11 Å from the peptide backbone (for synthesis, refer **Document S1**). These chemistries probed diverse non-native linkage motifs with features that modulate charge/pKa, hydrogen-bonding ability and hydrophobicity that are absent from the mammalian glycome, to create longer, orthogonal glycoconjugates (LOGs).

Initial application to the model protein antigen wild-type Hen Egg Lysozyme (wtHEL) generated corresponding biochemically homogeneous LOG products HEL-[amidine-GM3g]_n_, HEL-[amide-GM3g]_n_ and HEL-[aminoalkyl-GM3g]_n_ with full (n = 6) glycan occupancy in an efficient manner (**Fig S2a–c**) and LOG products were screened for endotoxin (**Fig S2d,e**; **Document S1**).

Mice were immunised with the three different HEL-[–X-GM3g]_6_ LOG antigens in MPLA adjuvant. Serum IgG titres against the glycoconjugate was assayed by ELISA against a corresponding LOG constructed from an unrelated protein carrier, gp120-[–amidine-GM3g]_16_ (**Fig 1c,d**). Antibody responses against the autologous LOG were considered a proxy for overall immunogenicity, whereas responses against the heterologous LOG indicated glycan cross-reactivity. High IgG titres were detected against autologous LOGs in antisera from all LOG-immunised mice. Antisera from HEL-[amidine-GM3g]_6_ and HEL-[amide-GM3g]_6_ were mutually cross-reactive with each other, reflective of only a small atomic variation (O versus NH) in display (**Fig 1d**). However, strikingly, whilst antisera from the aminoalkyl LOG was cross-reactive with both amidine- and amide-LOGs, the converse was not the case: amidine- and amide-LOG antisera were not reactive with the aminoalkyl LOG. Antibody titres against the protein carrier HEL revealed high-titre IgG in all immunisation groups, indicating that all methods used to achieve LOG glycoconjugation largely preserved native HEL epitopes. Since the amidine-based LOG provided the highest homologous anti-GM3g titres, we chose to prioritise its investigation.

### Conjugation to protein is necessary for LOGs to elicit glycoconjugate-specific IgG

To evaluate whether covalent linkage to the carrier protein was required for LOG immunogenicity, we compared immunisation with either LOG HEL-[amidine-GM3g]_6_ or instead with wtHEL that had been non-covalently mixed with stoichiometrically equivalent (n = 6) amounts of a corresponding, non-conjugated (‘free’) side-chain-only amidine-GM3g [–C(NH)NH-GM3g] (**Fig S2f**). To test GM3g-specific effects in particular, cross-reactive IgG titres were measured via ELISA against gp120-[amidine-GM3g]_16_. The HEL-based LOG antiserum had substantial titres of amidine-GM3g-reactive IgG even in the absence of exogenous adjuvant (EPT = 1,430), which increased with the addition of adjuvant. However, no LOG-specific responses were detected in the groups immunised with mixed, unconjugated HEL-plus-amidine-GM3g (*P* < 0.0001, Tukey’s post-hoc) (**Fig S2g,h**). These data imply that LOG immunogenicity is contingent on conjugation of the glycan to a protein carrier to facilitate B cell activation and isotype switching via T cell help, anticipated from classical hapten-carrier biology (*33*).

### Precise LOG-editing maps the role of glycan site and stoichiometry in modulating immunogenicity

Given the robust immunogenicity of LOGs, we next set out to better understand the molecular basis for glycan moiety immunogenicity by mapping the functional roles of both glycan site and copy number in precise structure-activity relationships. Importantly, our ‘tag-and-modify’ LOG construction methods (*32*) allowed ready LOG ‘editing’ simply via corresponding control of ‘tag’ site and copy number. In this way, site-directed mutagenesis of Lys to Arg allowed codon assignment whilst leaving global protein physicochemical properties including charge essentially unchanged. We designed a set of mutant HEL constructs to control the number of Lys and subsequently GM3g copy number and spacing (**Fig 1e,f; Fig S3**). This set of mutants permitted the dissection of features including moiety spacing, such as proximal versus distal GM3g glycoconjugates in HEL-[–amidine-GM3g]_3p_ and HEL-[–amidine-GM3g]_3d_). Notably, predicted pI values were essentially unaltered: wtHEL was 9.32, whereas HEL-null (in which all Lys were mutated to Arg) was 9.48. In this way, full control of Lys sites and copy numbers (n = 0–6) allowed editing of the GM3g in corresponding LOGs to generate a comprehensive panel of HEL LOGs (HEL-[– amidine-GM3g]_0-6_). These allowed dissection of the individual contributions of HEL-[– amidine-GM3g]_6_ in what represents, to our knowledge, an unprecedented parsing of the site-specific roles of glycan moieties in probing glycoconjugate immunogenicity. Strikingly, these revealed that not only is copy number a determining factor, but that contrary to prior avidity-centric perceptions, maximal loading does not deliver maximum titres. Indeed, optimal sugar loading with respect to anti-glycoconjugate antibody production was not proportional to the number of modifications but was found to be 2–4 (for HEL-[–amidine-GM3g]_2-4_) in the absence of adjuvant, with significant reductions in IgG titres for HEL-[–amidine-GM3g]_5_ and HEL-[–amidine-GM3g]_6_ (*P* < 0.0001) (**Fig 1g**). Interestingly, the glycoconjugate spacing in the case of HEL-[– amidine-GM3g]_3p_ and HEL-[–amidine-GM3g]_3d_ had no obvious bearing on the final GM3g-specific IgG titres.

To understand the origins of this counterintuitive outcome, we evaluated possible mechanisms. First, we tested whether the increased GM3g-specific titres arising from HEL-[–amidine-GM3g]_2-4_ immunisation were a consequence of Lys-to-Arg mutations changing the T cell immunogenicity of the protein backbone, possibly introducing artificial T cell epitopes that enhanced the response rather than a genuine GM3g loading effect. To assess this, we immunised mice with incompletely amidine-GM3g-modified wtHEL derived from chemical modification conditions adjusted to instead yield a product where the mean glycan occupancy was lowered to ∼3.7 per HEL. Mice immunised with this alternative lower copy product again showed greater GM3g-specific IgG titres compared to the high copy number LOG, HEL-[–amidine-GM3g]_6_ (**Fig S4a,b**) (*P* = 0.029), implying that the differential GM3g titres were unlikely to result from protein carrier amino acid substitutions impacting T cell help. Notably, HEL is a weak T cell antigen in BALB/c mice(*34*), and though the high IgG titres imply that sufficient T help is generated to facilitate reliable antigen-specific isotype switching, we were unable to detect Th recall responses, including in mice that had received HEL in MPLA (**Fig S4c–g**).

To further probe the relationship between glycoconjugate occupancy and the downstream humoral response, we evaluated the anti-GM3g IgM response two weeks post-prime (**Fig S4h**). These titres reflect the early humoral response which may not necessitate Th support. Although IgM titres were lower and data more dispersed compared to IgG, the trends with respect to glycan occupancy were the same, again implying that this is likely to be a Th cell-independent effect. This GM3g occupancy phenomenon was distinct from that observed against the HEL backbone, which was found to largely be adjuvant-(*P* < 0.0001) rather than sugar loading-dependent (*P* = 0.3496, two-way ANOVA) effect (**Fig S4i**). Collectively, these data therefore highlight that glycan occupancy may have a substantial effect on antibody outcomes, suggesting that the titration of optimal loading can be leveraged to deliver higher titres.

Interestingly, HEL-[–amidine-GM3g]_0_ in which all lysines were mutated to arginine elicited a low titre anti-GM3g response (EPT = 2,940) in formulation with MPLA (**Fig 1g**). These data, along with mass spectrometric analysis (**Fig S3b**) suggest that even partial incorporation of GM3g onto the *N*-terminal primary amine is sufficient to initiate a response against the glycoconjugate.

### GM3g-specific antibodies raised with multiple protein carriers

Having demonstrated that HEL LOGs elicit substantial IgG titres even with relatively low glycan copy numbers, we next tested the immunogenicity of the amidine-GM3g LOG on a different protein carrier, truncated gp120. This provided an excellent additional test of the LOG method, with more potential Lys ‘tag’ sites and a backbone that supplies multiple Th epitopes. Notably, while the total number of lysines on the gp120 construct used was 25, after application of the same benign editing methods for LOG generation, we estimated via electrophoretic analysis and densitometry data that amidine-GM3g loading delivered a mean of approximately 16 modifications (gp120-[–amidine-GM3g]_16_, **Fig S5a,b**). This partial lysine occupancy may be a consequence of the heavy endogenous *N*-linked glycosylation on gp120 reducing the accessibility of some lysine sidechains.

To assay longitudinal outcomes, animals were immunised with gp120 or gp120-[– amidine-GM3g]_16_ and bled periodically (**Fig S5c**). gp120-[–amidine-GM3g]_16_ rapidly induced GM3g-reactive IgG even after a single immunisation in the absence of adjuvant (IgG EPTs ∼10^3^), which further increased after boosting (∼10^5^–10^6^), unlike the unmodified gp120-only counterpart (*P* < 0.0001) (**Fig S5d–f**). These titres further increased with adjuvantation, with titres approximately an order of magnitude greater at the terminal timepoint (*P* = 0.006). Interestingly, gp120-[–amidine-GM3g]_16_ antisera displayed dramatically less antibody reactivity against the unmodified gp120 protein backbone compared with the unmodified gp120 antiserum against the unmodified gp120 protein backbone, implying that the GM3g modifications disrupted or masked immunodominant native gp120 epitopes (**Fig S5g,h**). This is consistent with GM3g ‘plugging gaps’ between the extensive native *N*-linked glycosylation sites. Similar antibody outcomes were also observed after immunisation with a corresponding LOG based on influenza A virus H1N1-NC99-HA-trimer (*P* = 0.016) (**Fig S5i–k**), H1N1-HA-[–amidine-GM3g]_26_. These data collectively demonstrate that GM3g-reactive antibody responses may be elicited regardless of the carrier protein. These responses were also irrespective of mouse sex and genetic background (**Fig S6**).

### Antigen-specific T helper responses are unaltered in LOGs

Any protein alteration, including the methods we used here to generate LOGs, may also affect downstream peptide processing and antigen presentation. We therefore tested the specific impact of LOGs on T cell antigen-specific recall responses. Whole spleen suspensions from gp120-[–amidine-GM3g]_16_-immunised mice were stimulated *in vitro* with unmodified gp120, gp120-[–amidine-GM3g]_16_ and HEL-[–amidine-GM3g]_6_ for 16 h (adding Brefeldin A for the final 6 h). IFN-ᵧ^+^ CD4 T cells were quantified and contrasted between the vaccination and re-stimulatory conditions (**Fig S7a–d**). Detectable antigen-specific responses were found only in the adjuvanted groups irrespective of the GM3g-presentation status of the immunogen. Moreover, the recall response was of equal magnitude whether gp120 or gp120-[–amidine-GM3g]_16_ were used. HEL-[–amidine-GM3g]_6_ did not induce any recall responses, confirming the important role of the conjugated carrier in providing T cell help. Together these suggested that the presentation of GM3g with LOGs did not inhibit the capacity for corresponding antigen to be processed nor for corresponding T cells to recognise anchored peptide (*P* > 0.9999). We further evaluated secretion of a broader panel of cytokines in supernatant after 72 h and observed similar trends in both IL-2 and IL-4 (**Fig S7e–g**). As is classical in the Th2-biased BALB/c background, IgG1 was the predominant isotype, with the TLR-4/Th1-biasing MPLA adjuvant bolstering IgG2a production (**Fig S7h,i**).

### Variation of the glycan in LOGs elicits orthogonal antibody outcomes

Having demonstrated that GM3 LOGs may be created in forms that are strongly immunogenic for B cell responses, we tested the extension of this phenomenon to other self-glycans. We chose the Lewis group trisaccharide Lewis-X (Le^X^) as another representative glycan for its similar size (trisaccharidic) and yet differing sugar content and arrangement (branched, non-linear) and charge state (neutral) (**Fig 1h**). Corresponding gp120-[–amidine-Le^X^]_n_ LOG was constructed in an essentially identical manner and used in formulation with MPLA in identical immunisation protocols. Antibodies were similarly raised against the Le^X^ LOG, with significantly greater titres compared with animals immunised with unmodified gp120 (*P* = 0.005) (**Fig 1i**). Notably, antiserum raised against either corresponding Le^X^g or GM3g LOGs were orthogonal, strictly binding autologous glycan, implying tight glycan specificity.

### B cell clonality against GM3g LOG is narrow

To dissect the molecular mechanisms underpinning the surprisingly robust B cell response against the LOGs, we conducted comprehensive clonotyping using animals primed with the HEL-[–amidine-GM3g]_6_ LOG. Antigen-specific B cells were sorted from mice, sorting on pre-gated IgD^-^ B cells according to molecular probes specific either to the glycoconjugate or the protein backbone (**Fig 2a,b**; **Fig S8a**). Heavy chain variable regions (V_H_) were recovered from one mouse and sequenced from 87 events, for which the majority (80/87) were GM3g-specific (**Fig 2c**). Clonality was defined according to the inferred heavy chain VDJ gene origins (**Fig 2d**; **Fig S8b**). Antigen-specific events were found in the spleen and bone marrow rather than inguinal lymph nodes, suggesting that draining follicular responses had ceased by four weeks post-administration (**Fig 2e**).

**Figure 2:**
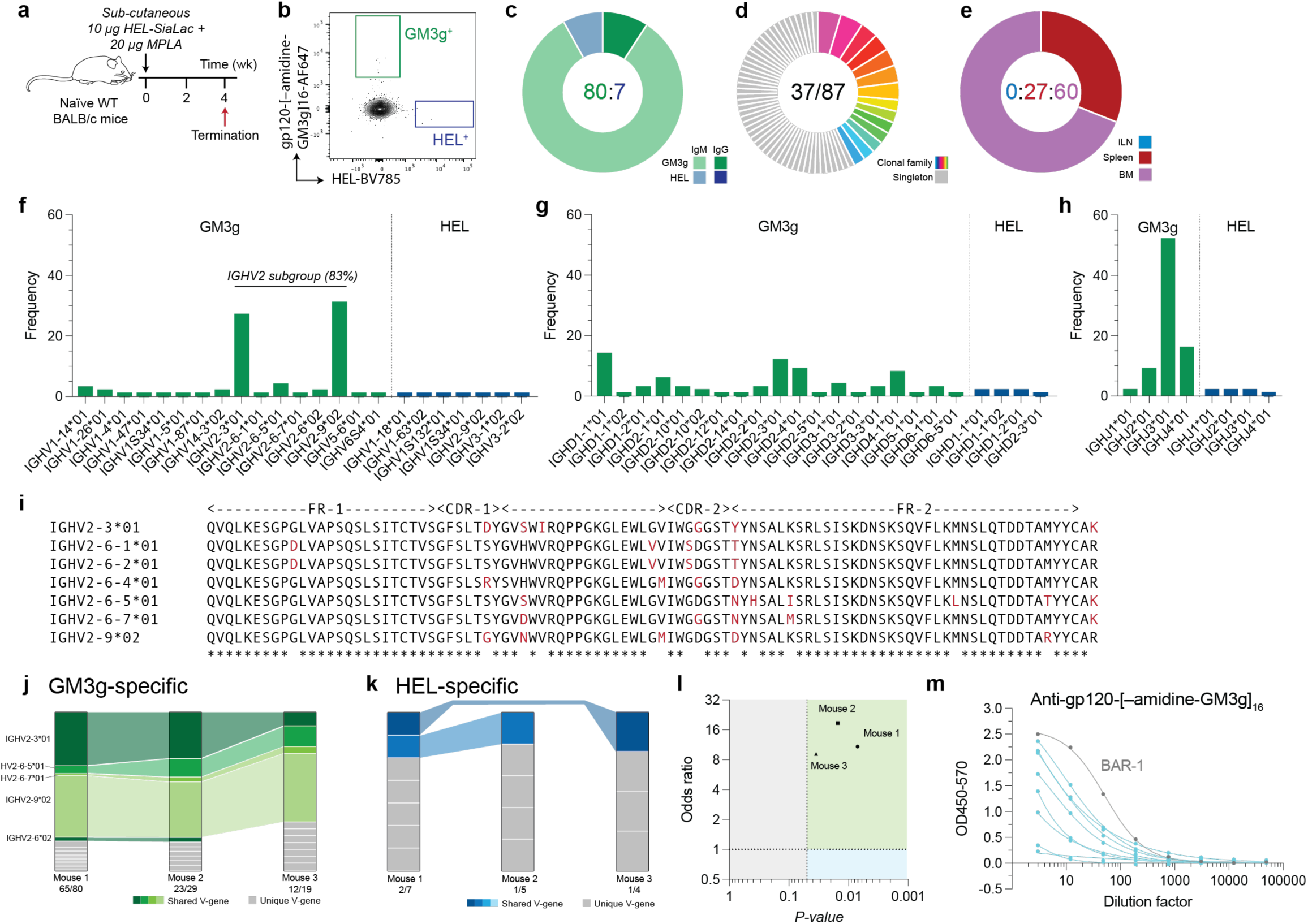
Evaluation of the B cell clonality raised against the semi-synthetic amidine-GM3g LOG. **(a)** Immunisation schedule. **(b, c)** Antigen-specific sorting strategy on pre-gated IgD^-^ B cells. **(d)** B cell clonality as inferred from the VH sequences. **(e)** Location the B cells were recovered. **(f–h)** VH gene segment utilisation. **(i)** Alignment of enriched *IGHV2* gene segments. **(j,k)** Distribution of common V-genes from B cells that bound either the GM3g- or HEL-specific probes. **(l)** Odds ratios were calculated for the proportion of share V_H_-genes as a proxy for germline restrictiveness. Statistical analysis was conducted parametrically on the log of the odds ratio. **(m)** Supernatant from representative *IGHV2*-origin B cells were screened against gp120-[–amidine-GM3g]_16_ via ELISA. **(c–h)** Representative data from Mouse 1.

The specific gene segments present in the isolated clones (**Fig 2f–h**; **Fig S8c**) reveal striking homology in their IGHV utilisation. In particular, the *IGHV2* subgroup was the predominant V_H_-gene class used in the GM3g-specific events and was expressed in > 80% of sorted B cells. The phylogenetically-related *IGHV2-3*01*, *IGHV2-6-5*01* and *IGHV2-9*02* members were the most well-represented in the GM3g-binders (**Fig 2i**). By contrast, the proportionality of V-genes utilised among HEL-binding B cells was significantly more diverse. Furthermore, D- and J-gene usage was highly diverse among these clones, implying that they tolerate broad CDRH3s and joining orientations.

V_H_-gene utilisation was also highly related between animals, implying a striking consistency in the use of this V_H_-gene-dependent clonal class in facilitating LOG binding (**Fig 2j**). This was unlike the HEL-binding clones; for these a broader, more diverse set of clonotypes was isolated, fully consistent with the larger antigenic protein surface compared with the more restricted but seemingly immunodominant glycan surface in corresponding LOGs (**Fig 2k**). The corresponding odds ratio that a given V-gene would be shared with respect to the antibody binding target revealed that for all animals, there is significantly narrower V-gene utilisation against LOG than the protein backbone alone (**Fig 2l**).

Given the strikingly restricted clonotypology of the anti-[–amidine-GM3g] response in the context of the broad tolerance to diverse D_H_ and J_H_ genes, the LOG was hypothesised to access a high frequency of naïve B cells. To interrogate this, LOG-binding naïve B cells from murine splenocytes were detected at a strikingly high frequency of 0.025% of IgD^+^IgM^mid-hi^ B cells (**Fig S8d,e)**. These events were sequenced from one mouse, revealing similar enrichment of the *IGHV2* subgroup (88%) compared with the immunised mice (**Fig S8f,g**).

A representative subset of several GM3g-binding IgGs from the IGHV2-subgroup origin were recombinantly synthesised and supernatant screened against gp120-[– amidine-GM3g]_16_ (**Fig 2m**) – all bound specifically, confirming functionality. The best binder amongst these antibodies, termed BAR-1 with inferred germline V_H_-gene IGHV2-9*02 (**Fig. 2m**), was purified for further analysis.

### The influence of the protein backbone on B cell outcomes does not perturb narrow anti-glycan clonal responses

Having isolated and identified the role of the *IGHV2* subgroup in the binding of HEL LOGs, we aimed to determine the effects of the protein backbone on B cell clonal outcomes. We similarly sorted B cells from gp120-[–amidine-GM3g]_16_-immunised mice (4-weeks post-prime). B cells that bound the gp120 backbone were not identified (**Fig 3a,b**), consistent with undetectable gp120 serum antibody binding in these animals (**Fig 3c**) and other animals primed with this LOG as an immunogen (**Fig S5**). Strikingly, the V-gene usage of antibodies raised against gp120-[–amidine-GM3g]_16_ again revealed that the *IGHV2* subgroup dominates, representing > 90% of clones (**Fig 3d,e**), of the same clonotype as that observed in the HEL-[–amidine-GM3g]_6_-immunised mice.

**Figure 3:**
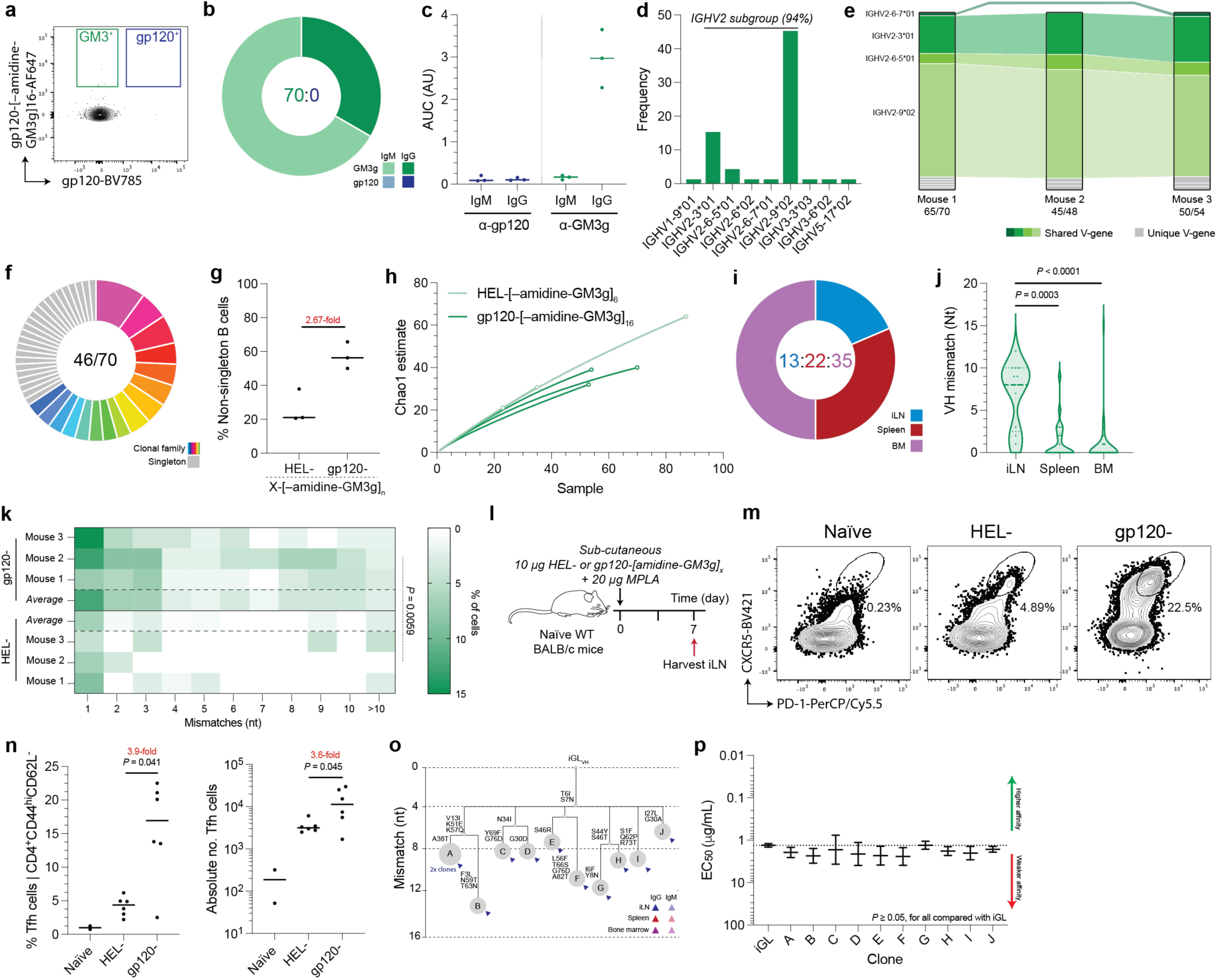
Protein backbone determines germinal center experience and clonal maturation of GM3g LOG-specific B cell. WT 6-week-old BALB/c mice were primed with 10 µg gp120-[–amidine-GM3g]_16_ + 20 µg MPLA and culled 4-weeks post-prime. **(a)** Antigen probe sorting strategy on pre-gated IgD^-^ B cells and **(b)** probe binding specificities of sorted events. **(c)** Terminal serum antibody reactivity was determined via ELISA. **(d,e)** IGHV gene utilisation. **(f)** B cell clonality was inferred according to the iGL VH sequences. **(g)** Percentage of clonal family members of isolated B cells with respect to protein backbone. **(h)** Chao1 estimates for mice immunized with either HEL- or gp120-based GM3g LOGs. **(i)** Location of isolated antigen-specific B cells. **(j)** VH mismatches compared to the iGL with respect to the compartment the B cells were isolated from. **(k)** Distributions of SHM rates with respect to protein backbone. **(l–n)** Evaluation of the Tfh population in draining iLNs 7 days post-prime. **(o)** Example clonal family phylogeny. **(p)** Relative EC50 values of recombinant GM3g-specific clonal family titrated against HEL-[– amidine-GM3g]_6_. **(b,f,i,j)** Representative data from mouse 1. Data were compared via non-parametric Kruskal-Wallis and post-hoc Dunn’s test.

We observed in gp120-[–amidine-GM3g]_16_-immunised mice that a higher proportion of B cells were members of clonal families compared with HEL-[–amidine-GM3g]_6_-immunised mice, with an average of 2.67-fold increase in the proportion of non-singleton B cells (**Fig 3f,g**). This may imply that the gp120 protein backbone offers greater clonal expansion, probably as a function of its improved T cell immunogenicity compared with HEL (**Fig S4, Fig S7)**. We further assessed the impact of the protein backbone on clonal diversity by performing a Chao1 estimate test (*35, 36*). While there was a trend for lower class sampling values in gp120-[–amidine-GM3g]_16_-immunised mice (which implies narrow clonal diversity), this was not statistically significant (**Fig 3h**). We also observed that at four-weeks post-prime, there were some antigen-specific B cells found in the iLN (**Fig 3i**) – this was not seen in the HEL-[–amidine-GM3g]_6_-immunised mice and may suggest that the different protein backbone maintains activated B cells within the secondary or tertiary lymphoid organ (S/TLO) structures, where much of the antigen persists, driving increased maintenance of the follicular response. We observed in the sequences isolated from gp120-[–amidine-GM3g]_16_-immunised mice that the degree of SHM undergone was compartment-specific (**Fig 3j**): the mean nucleotide mismatch of VH sequences derived from the lymph node was 6.8, spleen was 1.4 and bone marrow 0.8. Moreover, the extent of SHM undergone by the clones raised against gp120-[–amidine-GM3g]_16_ were significantly greater than that against HEL-[–amidine-GM3g]_6_ (*P* = 0.0059, Kolmogorov–Smirnov test) (**Fig 3k**). These data implicate the protein backbone in determining the maintenance of the primary germinal centre (GC) reaction conditions.

To understand the cellular underpinnings of the improved GC experience of gp120-[– amidine-GM3g]_16_-raised clones, we measured the induction of follicular helper T (Tfh) cells with respect to protein carrier. We demonstrated that the gp120 carrier elicits a larger Tfh population (**Fig 3l–n; Fig S9**), which is coordinate with the concept that the extent of SHM experienced by the glycoconjugate-specific B cells can be toggled by changing the T cell immunogenicity of the carrier protein.

### LOGs induce minimal affinity maturation despite SHM

Having shown differential SHM rates with respect to the protein carrier, we next evaluated the functional effect of SHM on antibody affinity. First, we analysed the mutation frequencies across the V_H_ gene in an unbiased manner to identify whether there were codons that were commonly mutated across the gp120-[–amidine-GM3g]_16_-immunised mice (**Fig S10a**) and identified that positions in CDRH1––namely T6I and S7N––were frequently mutated across multiple animals (**Fig S10b**). To evaluate the effect of these mutations, we introduced these changes into BAR-1 and screened their binding via ELISA; the data revealed no significant differences in binding compared to the wild-type mAb (**Fig S10c**), implying a lack of affinity maturation associated with these mutations. Second, we selected the largest clonal family, which had undergone significant expansion and diversification and was of an inferred *IGHV2-9*02* origin (**Fig 3o**). These antibodies were expressed recombinantly and screened via ELISA against HEL-[–amidine-GM3g]_6_ and the EC_50_ values were compared against that of the iGL (**Fig 3p**). Our data showed no evidence of increased affinity against the glycoconjugate, despite substantial SHM, collectively suggesting a strongly limited capacity for B cells to further improve binding against the carbohydrate.

### LOGs raise a specific anti-sugar polyclonal antibody specificity

To dissect anti-glycan specificity, we screened antisera derived from gp120 and from gp120-[–amidine-GM3g]_16_ LOG against a panel of 137 mammalian glycans (**Fig 4a; Table S3, Table S4**)(*37, 38*). This broad assessment revealed strikingly focused and specific binding against only nine glycans of >220. Indeed, cross-reactivity was seen only to very subtly altered features: ⍺-2,3 → ⍺-2,6 → monohydroxylated *N*-acetyl-Neu → *N*-glycolyl-Neu or OH-2-Glc → NHAc-2-GalNAc. To further interrogate the specificity of the polyclonal antibody response, we designed a soluble ligand competition assay for the binding of the antiserum to arrayed HEL-[amidine-GM3g]_6_. Consistent with the glycan panel analysis, GM3g antiserum bound essentially equivalently to its OH-2-Glc- and NHAc-2-GalNAc variants (IC_50_ of 7.76 mM and 9.90 mM, respectively) (**Fig 4b,c**). Two truncated variants further mapped GM3g specificity and saccharidic moiety dependency: the disaccharide variant SiaGal competed relatively weakly (47.8 mM), implying some role for the ‘inner’ reducing-end interactions, whereas GM3g disaccharide lacking ‘tip’ non-reducing-end Sia showed no detectable competition, implying the presence of more critical contacts made with the terminal sialic acid. These findings were rationalised by our subsequent structural analysis.

**Figure 4:**
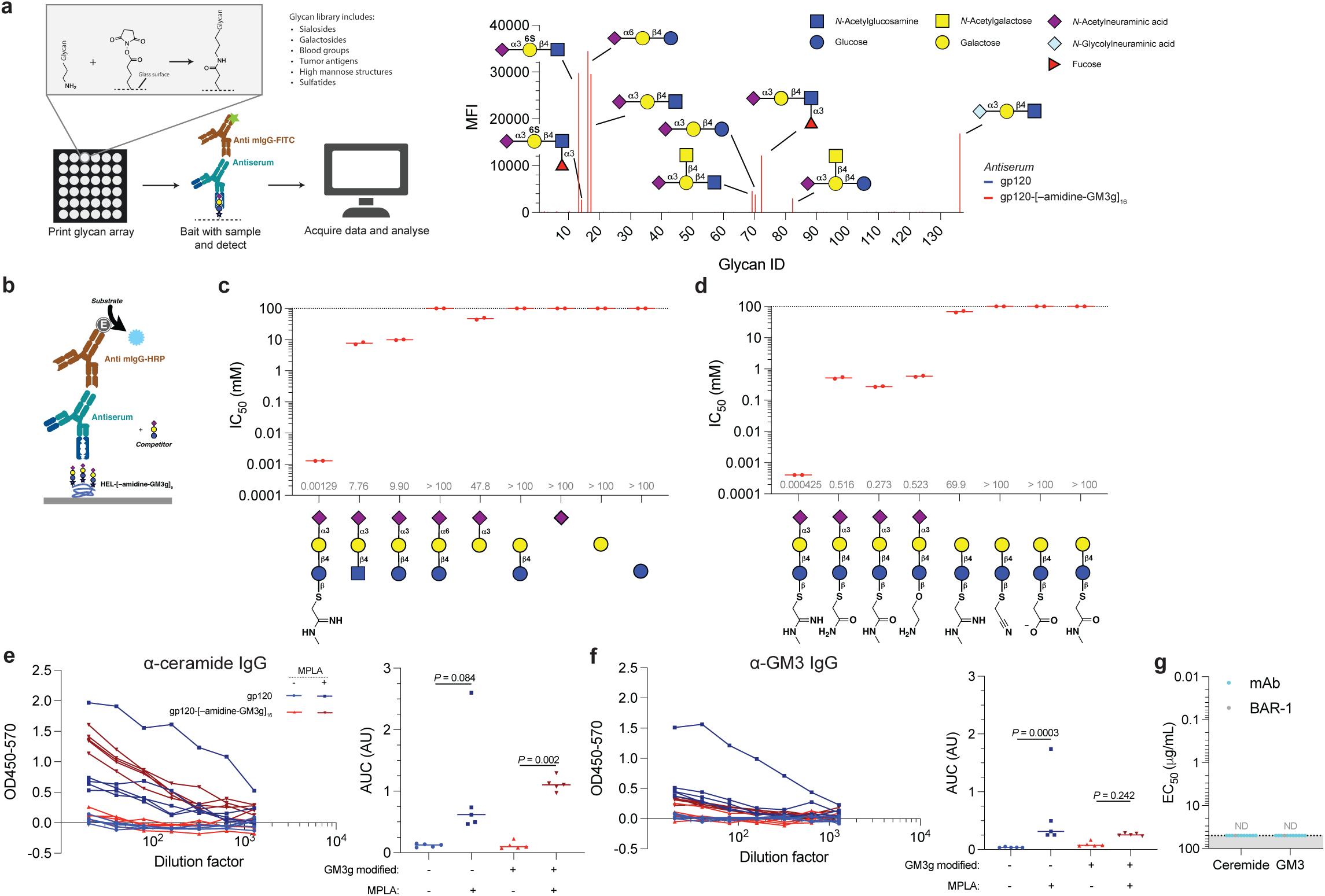
Glycan specificity, linker engagement and context dependency of amidine-GM3g LOG-raised antibodies. **(a)** Antiserum from gp120-[–amidine-GM3g]_16_-immunized mice was screened for reactivity against a mammalian-derived glycan library. **(b–d)** A competition ELISA was conducted to evaluate the polyclonal antibody dependency on component and adjacent ligand segments, as well as the antibody tolerance of alternative linker formats. Polyclonal reactivity against **(e)** a negative ceramide control and **(f)** native presentations of SiaLac on GM3 was evaluated via direct ELISA using serum raised in gp120-[–amidine-GM3g]_16_ antiserum. **(g)** These analyses were additionally screeded using a set of purified GM3g-specific recombinant monoclonal antibodies later characterised. Data were compared using a post-hoc Dunn’s test.

Although the data imply that the complete GM3g glycan structure is a required component of antibody binding, we also observed broad, substantial contributions from differing non-reducing aglycones (**Figure 4d, left**): enhanced binding for amide and aminoalkyl aglycones was potentiated further by the presence of an amidine. Any such potentiation was notably lost in the absence of incorrect glycan (**Fig 4d, right**), further highlighting the role of tight glycan recognition in driving affinity, despite apparent engagement both of glycan and aliphatic constituents. Thus, although these data suggest that the antibody response targets the linker-glycan motif, it is nonetheless specific to the GM3g glycan.

We next interrogated the binding of GM3g LOG-raised antibodies against native GM3g display through ELISA screening gp120-[–amidine-GM3g]_16_ antiserum against GM3 and a ceramide control. Data revealed no indication of GM3-specific binding, but rather elevated non-specific reactivity with both ceramide and GM3 in an MPLA-dependent manner, potentially a function of the adjuvant mounting non-specific antibody responses with a substantial hydrophobic element (**Fig 4e,f**). To eliminate any serological background and control for the non-specific binding observed in the MPLA-adjuvanted gp120-[–amidine-GM3g]_16_ antiserum, GM3g LOG-reactive monoclonal antibodies of an *IGHV2* origin were purified and again screened via ELISA. No binding was detected against either ceramide or GM3 in any of the 11 clones tested (**Fig 4g**). These data imply that the antibodies raised against GM3g presented synthetically in this manner fail to elicit reactivity against native glycan presentation.

### Biophysical, biochemical, and structural properties of the dominant GM3g-engaging clonal class

We generated and purified the Fab of the GM3g-binding mAb clone, BAR-1 and quantified binding using surface plasmon resonance (SPR) against an amidine (C(NH)NH)-GM3g-coated surface, bearing the same extended side-chain motif as used in LOGs, generating a K_D_ = 17 ± 1 μM (**Fig 5a**). Next, we synthesised an equivalent soluble ligand, Lys–amidine-GM3g, as a representative minimal LOG motif, and a truncated variant Me–amidine-GM3g and conducted solution-phase isothermal titration calorimetry (ITC), generating respective similar K_D_ = 5.4 ± 1.2 μM (Lys– amidine-GM3g, **Fig 5b, Fig S11a)** and K_D_ = 2.1 ± 0.7 μM (Me–amidine-GM3g, **Fig S11b,c**). Notably, consistent with LOG design, rather than display entropic cost, both displayed balanced binding thermodynamics (TΔS = -1 kcal/mol and - 5.7 kcal/mol, respectively). Competition ELISAs using these soluble ligands were consistent with that observed using polyclonal sera, namely, that binding could be competed out using soluble GM3g, but that Me–amidine-GM3g was more competitive (**Fig 5c**).

**Figure 5:**
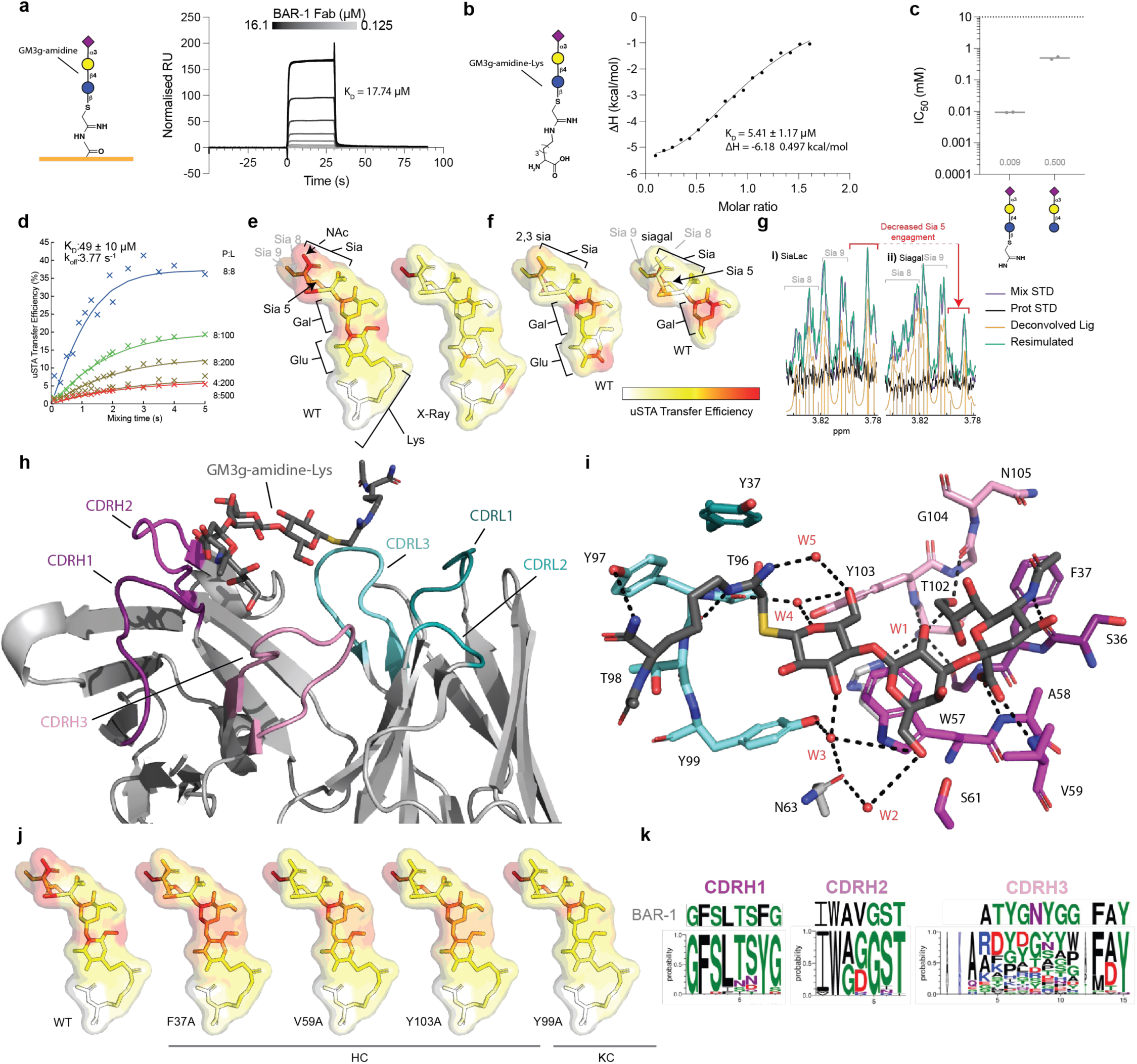
Structural, biophysical and biochemical characterization of BAR-1 reveals germline-encoded lectin-like GM3g-engaging motifs. **(a)** The binding kinetics of the BAR-1 Fab was evaluated via SPR using an amidine-GM3g-coated chip. **(b)** ITC was conducted with a soluble reductionist LOG mimic, Lys-amidine-GM3g. **(c)** Competition ELISA using BAR-1 soluble ligands. (d) *K*_D_ and k_off_ for the amidine/Bar-1 complex were determined using uSTA NMR measurements repeated over titrated proton/ligand concentrations and analysing data from the NAc proton signal. **(e)** NMR uSTA analysis of the BAR-1:Lys–amidine-GM3g binding in solution. **(f)** Truncation of Lys–amidine-GM3g to GM3g and its tip disaccharide (Neu5Ac-Gal; Siagal) notably leads to a readjustment of GM3g focused even more upon the ‘foothold’ interaction of Gal. **(g)** Example raw uSTA data illustrating residue level precision that reveals the subtle readjustment of Lys–amidine-GM3g to GM3g effect, here analysed via the H-5’’, H-8’’ and H-9’’ positions. **(h)** Side view of the 1.9-Å x-ray structure of the BAR-1 Fab bound to the Lys-amidine-GM3g. Both heavy and light chain CDRs are marked. Binding side of GM3g-amidine-Lys. **(i)** Residues within 4.0 Å of the polysaccharide are displayed and hydrogen bonds are shown as black broken lines. Water is marked in red. Lys–amidine-GM3g is shown as sticks. **(k)** Logo plots of the CDRH residues from isolated GM3g-specific *IGHV2* subgroup-encoded B cells.

Dynamic structural interrogation of the BAR1•Lys–amidine-GM3g complex using universal standard transfer analysis (uSTA) protein NMR (*39*) (**Fig 5d,e**; **Fig S11c–l**) gave a *K*_D_ = 49 ± 10 μM and *k_off_* = 3.77 ± 3 s^-1^, consistent with values obtained by complementary methods (**Fig 5a,b**). Atomic-level ‘heat maps’ of magnetization transfer in uSTA revealed a ligand pose with primary engagement of BAR1 with the glycan motif of GM3g over the Lys–amidine-linker moiety. Interestingly, analyses of the interaction of two truncated ligand variants – GM3g itself and just the tip disaccharide Neu5Ac-Gal (**Fig 5e–g**) – further identified relaxation of GM3g alone into a pose that creates even greater contact of the Gal upon removal of the LOG longer-linker moiety in Lys–amidine-GM3g. This suggested topological frustration in the complex with Lys–amidine-GM3g (and by extension the LOG) that, when removed, allows a relaxation further into the binding motif.

Next, the atomic level basis of these interactions was probed through complementary methods, allowing structural analysis of BAR-1 in complex with Lys–amidine-GM3g. Crystallization of the BAR1•Lys–amidine-GM3g complex revealed a striking, seemingly LOG-specific arrangement in the 3D-structure of the *holo* complex (**Fig 5h,I** and **Fig S12a**). Notably, consistent with design, the longer length of the LOG moiety allowed the GM3g to ‘reach through’ a seemingly flexibly-engaged CDRL3 region to engage key residues in CDRH2, and, also, to some extent CDRH1, leaving the part of the groove formed by CDRH3, CDRL1 and CDRL2 unoccupied. The antibody binding pocket is largely hydrophobic in character.

The crystal contained two complete copies of the complex which are largely identical (rmsd of light chain 0.5 Å). In both the electron density is well ordered for all three sugar rings and the amidine of Lys–amidine-GM3g, but less well ordered for the ‘reach through’ lysine side-chain. As a seemingly key ‘foothold’ the indole of Trp57_H_ stacks against the alpha-face of the Gal sugar of GM3g to create a classical pi-CH interaction (**Fig 5i** and **Fig S12c**) found in diverse so-called carbohydrate modules (CBMs) (*40, 41*). This is supported by binding of the tip Neu5Ac sugar of GM3g, which makes five hydrogen bonds to BAR1 backbone, including a striking bidentate interaction of its C-1 carboxylate with amide nitrogen atoms of Ala58_H_ and Val59_H_ but notably there is no charge-driven interaction. Several highly coordinated water molecules (W) also contribute to binding, as well as a hydrophobic pi-CH interaction with Phe37_H_. The reducing-end Glc of GM3g also makes hydrogen bonds to three water molecules, two of which bridge to the protein (including W3 which bridges to Tyr99_L_, Asn63_H_ and galactose) but only three direct van der Waal contacts with the protein. The amidine linkage of Lys–amidine-GM3g makes hydrogen bonds to the protein (Tyr97_L_) and to a water molecule (W5) that bridges to the glucose and, intriguingly, a cation-pi interaction with Tyr37_L_ confirming a contribution from the longer amidine linker to binding. The aliphatic side chain of the lysine makes van der Waal contacts with Tyr97_L_.

To probe specific contributions to binding, including the ‘foothold’ Trp57_H_, we probed the residues lining the binding site of BAR-1 through Ala-scanning mutagenesis. uSTA protein NMR allowed us to look at the modulation of the binding pose adopted by Lys– amidine-GM3g. Strikingly, whilst alterations of lining residues Phe37_H_, Val59_H_, Tyr103_H_ and Tyr99_K_ retained residual binding in an ELISA format (**Fig S14**), their interaction surfaces were all essentially similar (**Fig 5j**). By contrast, no interactions at all were observed between Lys–amidine-GM3g and ‘foothold’ mutant Trp57Ala, further emphasizing its key role (**Fig S14c**). Specifically, when exciting the ‘ligand only’ sample at 8 ppm, small residual signal is seen in the STD spectrum. This was found to be of identical magnitude to the spectrum of the Trp57_H_ BAR-1 mutant, revealing that there was no detectable binding between ligand and protein.

Together, these structural and biophysical analyses highlight the key residues important in driving binding. These residues were notably conserved amongst the *IGHV2* subgroup-containing clones that we had validated for GM3g binding, with particularly high sequence similarity in their V_H_-encoded CDRH1 and CDRH2 loops (**Fig 5k**), and are also consistent with our mutagenic and structural analyses. These data also showed that involvement of CDRH3 in ligand binding was limited, which aligns not only with our structural analyses but also with the observation from our broad set of B cells isolated against the ligand and their tolerance of exceptionally diverse CDRH3.

## Discussion

For self-glycans, there is a heavy incentive to skew the naïve B cell repertoire to avoid the presence of self or self-like glycan-reactive B cells to prevent generation of autoreactive antibodies (*1, 8*), as supported by evidence of anti-glycan responses associated with various autoimmune conditions (*42, 43*). Notably, previous studies have failed to reliably raise high-titre antibodies responses against GM3 using conventional autologous formulations (*21, 26, 44*).

The LOG modular format has potential advantages compared with immunisation with autologous GM3, namely: i) the docking of the sugar to a peptidic carrier allows for associated T cell help, and ii) non-native presentation of otherwise immunorecessive TACAs via a bespoke chemical linker may bypass the tolerogenic constraints that prevent antibodies being raised against native glycan presentations in endogenous glycoconjugates. Our discovery that GM3g-specific IgG responses were readily mounted in a mouse (predominantly by the *IGHV2* subgroup) reveals that the LOG modular format of self-glycans can access a subset of naïve B cells that native presentations of the same glycan do not.

We have rationalised the lack of native glycoconjugate cross-reactivity by combining immunogenetic, structural, biochemical and biophysical-based analyses. The structure of the BAR-1 Fab with Lys–amidine-GM3g reveals that that the sugar portion is recognised by the CDR1 and CDR2 loops in the V_H_ domain. Intriguingly, the recognition of the galactose and sialic acid sugars closely resembles (**Fig S13**) the arrangement seen for a hexasaccharide binding antibody (*45*). Although showing a different ‘reach through’ orientation, there is striking conservation of several features of the final tip glycan interactions: primarily a classical pi-CH interaction between the CH beta-face of Gal and the indole ring of Trp (Asensio et al., 2013), the recruitment of bridging water molecules, a bidentate hydrogen bond between the carboxylate of the terminal sialic acid with the protein backbone and the engagement of the methyl of the *N*-acetyl by an aromatic residue. We suggest this serves as generic ‘foothold’ (focused on the Trp) in the antibody, and by implication BCR, for these two sugars. With LOGs, the linker of Lys–amidine-GM3g reaches through to the sugar-binding pocket by spanning the V_H_V_L_ interface to be recognised by the CDR3 loop of the V_L_ domain. In native GM3 the trisaccharide has a glycosidic link in close proximity to the branched arrangement of the ganglioside (displaying stearic acid and sphingosine). A ‘reach through’ mode observed for Lys–amidine-GM3g would therefore be unavailable to ‘self’ GM3. Close embedding of GM3 into the membrane creates prohibitive van der Waal clashes for antibody engagement.

The surprising immunogenicity of the LOG amidine-GM3g is, we hypothesise, a function of the set of V(D)J configurations tolerated by the BAR-1 clonal class. The binding of a naïve B cell receptor to an epitope is dependent on the combinatorial effect of the distinct V(D)J configuration of the cell. However, the structural dimensionality imparted by the germline configuration — a function of the unique combinatorial gene segment composition –– is, we show here, drastically reduced in this LOG-raised clonal class. These clones tolerate highly diverse sets of D_H_ and J_H_ segments, and our structural characterisation demonstrates that the corresponding CDRH3 has no substantial contribution to glycan recognition.

This suggests a clone-by-clone basis under which these anti-TACA antibody elicitation approaches ought to be considered. Taken together, our data now suggest that future approaches necessitate analyses of how a given glycoconjugate might engage TACA:linker specific B cell clonotype, using comprehensive structural and biochemical analysis of clonal reactivity, consideration of increased scope for SHM-directed affinity maturation, and determination of native-recognition in relation to context-dependent TACA-specific recognition.

Use of different protein platforms (e.g. HEL-[–amidine-GM3g]_6_ and gp120-[–amidine-GM3g]_16_ led to the same germline configurational response against the GM3g self-glycan moiety. This was also associated with a clear modulation of the germinal centre experience the clones had undergone with respect to the amount of follicular help offered by the different protein backbones. There was a direct relationship between the amount of T cell help detected and the ensuing antigen-specific B cell IgG response. This in turn suggests future application using heterologous boosters: yet more T-immunodominant backbones conducive to greater somatic hypermutation rates may then lead to clonal affinity maturational outcomes. Moreover, heterologous immunisation strategies based on TACAs presented in the context of systematically varied LOGs conjugated to different chemical linkers might exploit affinity maturation processes to ‘walk’ clones towards native glycan reactivity. This is unlike classical germline-targeted approaches, which use isolated and highly mutated antibodies of known functional effect as a template germline clonal class (*15, 47*). However, in the instance described here, it is not known whether ‘up-mutation’ of the BAR-1 class can move towards a functional effect to yield native GM3g recognition.

Finally, the explicit demonstration also of the presence of germline-encoded lectin-like motifs (*48–50*) present in the murine BCR germline is striking. This not only challenges the dogma associated with the perceived poor immunogenicity of glycans (*51*) but may also provide an explanation for the greatly divergent views and results that have in the past been obtained from immunisations with glycoconjugates. Not only may this be a consequence of conjugate presentation format (e.g. ‘parallel’ *versus* ‘orthogonal’ or shorter *versus* longer linkage), as we argue here, but may also be a consequence of the restricted clonotypic response that we have discovered here. It may be that only upon engagement of the correct glycoconjugate or glyco-epitope would a large proportion of naïve B cells be activated by using appropriate ‘predisposed’ germline BCRs, thus improving the frequency of B cell activation events *in vivo* and explaining the relatively high titres of anti-GM3g antibodies elicited after a limited immunisation regimen. We therefore propose that the logical design of the entire conjugate and not just, for example, the glycan as has been typical, is important to properly exploit these rare, correct engagement events in the effective design of future immunogens.

## Methods

### Generation of synthetic GM3 liposomes

All lipids were from Avanti Polar lipids. Lipid mixtures (as indicated in the results) in chloroform at 1 mg/mL were dried under the flow of nitrogen, rehydrated with buffer (150 mM NaCl, 10 mM HEPES, 2 mM CaCl_2_) and vortexed to form multilamellar vesicles. Then, suspension of the multilamellar vesicles were sonicated at power 3, duty cycle 40% for 10 mins using Branson Sonifier 250. Liposomes were stored under nitrogen atmosphere.

### Recombinant protein expression

Proteins were expressed recombinantly in-house using the HEK 293Freestyle expression system (Life Technologies). Cells were transfected using PEI Max (Polysciences) and relevant expression vector. Proteins were purified from supernatant using either Protein A agarose beads (Life Technologies), or immunoaffinity chromatography (D1.3 for HEL and 2G12 for gp120) where columns were prepared with AminoLink Plus resin (Life Technologies), both used according to the manufacturer’s instruction. Purified protein was tested for endotoxin contamination prior to immunisation. These analyses were conducted using either a HEK293T TLR4-CD14-MD2 IL-18 reporter line (Invivogen) or the RAW-Blue Cell assay (Invivogen). These readouts were acquired according to the manufacturer’s protocol. Protein preparations where the reporter endotoxin readouts were less than the < 0.125 ng/mL LPS control was considered clean.

### Synthesis of LOGs

Details of chemical synthesis and characterisation are outlined in **Document S1**. ***HEL-[amidine-GM3g]_6_:*** GM3g-SCN **1** (404 mg, 587 μmol) was activated in sodium methoxide solution (20 mM in 29 ml, <u>1.0 eq. of CH_3_ONa</u>) by following the standard protocol (as shown in the preparation of GM3g-imidate **7**). After stirring for 4 days at room temperature, THF (87 mL) or ether (29 mL) was added to precipitate the sugar. The white solid was separated by centrifugation, the supernatant was discharged and the white residue was then dried under vacuum before being used immediately for protein modification.

The precipitated sugar was dissolved in PBS buffer (2.8 mL, pH = 7.4). A fresh solution of protein in PBS buffer (0.7 mL, 5 mg/mL, pH = 7.4) was added (final concentration of HEL was 1 mg/mL) and the mixture was incubated at 25 °C for 12 h (checked by SDS-PAGE and LC_MS if necessary). The reaction was desalted by PD-10 column twice (note: desalting is not sufficient to completely remove sugar. Excess sugar was in the post fraction; GM3g-CN **1** could be recovered by purification). Dialysis was subsequently performed in PBS buffer at 4 °C (4 h × 2 and 12 h × 1). The solution was concentrated, sterilized and stored at 4 °C. Concentration of HEL-[–amidine-GM3g]_6_ was analyzed by BCA assay (7.13 mg/mL, 0.6 mL, endotoxin free-PBS buffer, pH = 7.4). Notably, modifications using [–amidine-Le^x^] were prepared in an essentially identical manner.

#### HEL-[–aminoalkyl-GM3g]_6_

A mixture of HEL (11ul, 220ug, 20mg/ml in PBS, pH = 7.4), GM3gOCH_2_CHO (41.56 ul, 30 mg/ml in H_2_O, 20 eq./lysine, 6*lysine), freshly prepared NaBH_3_CN solution (23.19 ul, 5 mg/ml in H_2_O, 20 eq./lysine, 6*lysine), topped to 220ul with H_2_O (144.25 ul, final protein concentration was 1mg/ml) was incubated in 37 °C for 24 h without shaking. Then solution was immediately dialyzed to PBS at 4 °C.

#### HEL-[–amide-GM3g]_6_

15.83 ul of **S-Short-NHS** (8 eq./lysine, 60mg/ml in DMSO, freshly prepared) was added into HEL (50ul, 1mg, 20mg/ml in PBS, pH = 7.4) solution, mixed and incubated for 3 h at room temperature. Following immediate desalting into water, protein was concentrated to 2 mg/ml and checked with LC-MS. To a solution of dimeric GM3g **28** (1 mg) in water (75.6 ul), 1.54 ul of TCEP solution (1.0 eq, 0.5 M in water, freshly prepared from TCEP (free acid) solid, neutralized by 3 eq. of NaOH) was added and, after incubation at room temperature for 30 mins, transferred it into 287 ul of iodo-HEL solution (574 ug, 2mg/ml, 6.4 eq. GM3g-SH/Lysine) above. 172 ul of sodium borate (100 mM, pH = 8.5) was added, topped to 1mg/ml (protein solution) with 37 ul of water. This mixture was incubated at RT for 3 hours (LC-MS checking) before being dialysed against PBS (pH = 7.4), concentrated, sterilized, and analyzed by BCA assay.

### SDS-PAGE

Proteins were evaluated via SDS-PAGE. Samples were ran on precast NuPAGE Bis-Tris gels (Life Technologies) with 1X MOPS buffer (Life Technologies) according to the manufacturer’s instructions. Proteins were stained using InstantBlue Coomassie protein stain (Abcam). Molecular weights were manually estimated according to the Novex Sharp protein marker. To obtain sharper bands on native N-linked glycosylated proteins, such as gp120, sample was pre-treated with PNGase F (New England Biolabs) according to the manufacturer’s instructions.

### Primary amine ELISA

Following the conjugation of the synthetic glycoconjugates to a carrier protein, relative free amines were contrasted to estimate the degree of lysine modification. This protocol was adapted (*52*). Briefly, protein samples (5 μg) in 10 μL PBS were mixed in 40 μL 0.1M sodium bicarbonate buffer. 5% solution of 2,4,6-trinitrobenzenesulfonic acid (TNBSA) was diluted 1:500 in the bicarbonate buffer, and 25 μL of this mixture was added to the protein sample. After 2 h incubation at 37°C. 25 μL 10% SDS and 12.5 μL of 1M HCl was added. The absorbance at 335 nm was measured using a Spectramax M5 spectrophotometer.

### Animal experimentation

Wild-type pathogen-free 6-week-old BALB/c mice were purchased from Charles River. Animals were monitored daily and provided standard chow and water *ad libitum*. Immunisation schedules are outlined in the results, and mice were bled periodically from the tail vein. Animals were sacrificed via a rising CO_2_ gradient and subsequent cervical dislocation schedule 1 procedure.

### ELISA assays

Samples were serially diluted and incubated on antigen-coated and blocked SpecraPlate-96 HB (PerkinElmber) plates. Antibodies were detected with either anti-mouse IgG-HRP (STAR120P, Bio-Rad), anti-mouse IgG1-HRP (STAR132P, Bio-Rad), anti-mouse IgG2a-HRP (STAR133P, Bio-Rad), anti-mouse IgM-HRP (II/41, BD Bioscience), or anti-human IgG-HRP (Jackson ImmunoResearch). ELISAs were developed using 1-Step Ultra TMB ELISA substrate (Life Technologies), with the reaction being terminated with 0.5M H_2_SO_4_. For competition ELISAs, sample was pre-incubated with the coating antigen for 1 h, before adding the competitor for an additional hour for a new equilibrium to be reached. Detection and development were subsequently conducted, as per the direct ELISA protocol. Cytokine ELISAs were performed using commercially available kits. IL-2, IL-4 and IFN-ᵧ in supernatant was measured according to the manufacturer’s protocol (all Life Technologies).

Optical densities were measured at 450 and 570 nm on the Spectramax M5 plate reader (Molecular Devices). After background subtraction, logistic dose-response curves were fitted in GraphPad Prism. Endpoint titres were determined as the point at which the best-fit curve reached an OD_450-570_ value of 0.01, a value which was always > 2 standard deviations above background.

### Intracellular cytokine analysis

Whole splenocyte suspensions (5 × 10^5^ cells in 200 μL in a flat-bottom 96-well plate) were stimulated *in vitro* with 10 μg/mL antigen in cRPMI for 16 h. For the final 6 h, 5 μg/mL brefeldin A (Biolegend) was added to all wells to suspend ER–Golgi trafficking and block cytokine secretion. Cells were subsequently washed with PBS with ice cold 2mM EDTA and stained with TruStain FcX Plus (Biolegend) and LIVE/DEAD Fixable Blue on ice for 30 minutes. Surface markers were subsequently stained on ice for 20 minutes: anti-mouse CD3-PE (1:200, 17A2, Biolegend), anti-mouse CD4-APC (1:200, RM4-5, Biolegend) and anti-mouse CD8-AF700 (1:200, RPA-T8, Biolegend). Following treatment with fixation and permeabilization buffers (Biolgend), cytokine was stained for using anti-mouse IFN-ᵧ-PE/DAZZLE 954 (1:100, XMG1.2, Biolegend)—this was conducted on ice for 40 minutes. Cells were washed twice with FACS buffer (PBS with 2% FCS and 0.05% sodium azide; FB) before acquiring data on the BD Fortessa X-20 (BD Bioscience).

### Vaccine-specific B cell isolation

Antigen probes were synthesised via modification using NHS-esterified biotin, AF647 or AF488 protein modification kits, as per the manufacturer’s instructions (Life Technologies). Successful modification was confirmed by both SDS-PAGE and subsequent fluorescent gel scanning on a ChemiDoc (Bio-Rad), as well as via mass spectrometry.

Immunised BALB/c mice were immunised 4 weeks prior to B cell isolation. Single cell suspensions were generated from the spleen, inguinal lymph nodes and bone marrow (femur and tibia). Fc receptors were blocked and stained with LIVE/DEAD Fixable Blue, as outlined earlier. The following surface stain cocktail was prepared: anti-mouse F4/80-PE (1:200, BM8, Biolegend), anti-mouse Gr-1 (1:200, RB6-8C5, Biolegend), anti-mouse CD3-PE (1:200, 17A2, Biolegend), anti-mouse CD4-PE (1:200, RM4-5, Biolegend), anti-mouse CD8-PE (1:200, RPA-T8, Biolegend), anti-mouse B220-eFluor450 (1:100, RA3-6B2, BD Biosciences), anti-mouse IgD-AF700 (1:200, 11-26c.2a, Biolegend), anti-mouse IgM-PE/Cy7 (1:200, R6-60.2, BD Biosciences), anti-mouse IgG1-FITC (1:200, A85-1, BD Biosciences), anti-mouse IgG2a/2b-FITC (1:200, R2-40, BD Bioscience), antigen probes as indicated in the results (10 μg/mL). Cells were stained on ice for 1 h. Cells were washed with FB and sorted immediately on a BD FACSAriaFusion (BD Biosciences). LIVE/DEAD^-^DUMP^-^B220^mid–hi^IgD^-^ (IgM^+^/IgG^+^)Ag^+^ B cells were singly sorted into MicroAmp Optical 96-Well PCR Plates (Life Technologies) containing 5 μL 1X TCL buffer supplemented with 1% 2-ME. Immediately following sorting, plates were centrifuged at 1,500 g for 1 minute. Plates were stored at -80°C until use.

### B cell receptor variable region recovery

Recovery of the antigen-specific B cell receptor variable regions was carried out, adapted from previous publications (*53, 54*). We are happy to share a detailed step-by-step protocol upon request. Briefly, single cell lysates were thawed on ice and RNA was captured on RNAClean XP beads (Beckman Coulter), subsequently washing with 70% ethanol. RNA was eluted and cDNA libraries were synthesised using SuperScript III (Life Technologies) with random primers (Life Technologies). VH and VK regions were recovered using the first PCR primer sets (**Table S1**) and Q5 polymerase. VH amplicons were purified and sequenced using 5’ MsVHE. These sequences were used to determine B clonality.

To confirm the recovered sequences were truly antigen-specific, antibodies were synthetised recombinantly. To incorporate the variable regions into an expression vector, vector-overlapping adapters were incorporated via PCR (**Table S1**), and the V regions were inserted into cut backbone vectors (heavy chain: FJ475055; kappa chain: FJ75056) via Gibson reaction (New England Biolabs). Successful clones were prepared, and vector products were transiently transfected into HEK 293Freestyle cells, as previously outlined.

### Immunogenetic analyses

Immunogenic analyses were performed on the VH regions of successfully recovered clones (**Table S2**). Sequences were screened against the mouse germline gene segment repertoire using the Immunogenetics Information System (IMGT; https://www.imgt.org/IMGT_vquest/input). These outputs were used to determine both clonality (i.e. their inferred V(D)J configuration) and the SHM rates. Sequences that returned either no result (indicative of poor sequence read quality) or was unproductive (e.g. premature stop codon) was excluded from our analysis. Clonal lineage trees were determined using GCTree (*55*) and rendered on Adobe Illustrator.

### Glycan array

Glycan arrays were custom printed on a MicroGridII (Digilab) using a contact microarray robot equipped with StealthSMP4B microarray pins (Telechem) as previously described (*38*). Briefly, samples of each glycan were diluted to 100 μM in 150 mM Na_3_PO_4_buffer, pH 8.4. Aliquots of 10 μl were loaded in 384-well plates and imprinted on NHS-activated glass slides (SlideH, Schott/Nexterion), each containing 6 replicates of each glycan. Remaining NHS-ester residues were quenched by immersing slides in 50 mM ethanolamine in 50 mM borate buffer, pH 9.2, for 1 hr. Blocked slides were washed with water, centrifuged dry, and stored at -20 °C until use.

Serum samples were diluted 1:200 in PBS + 0.05% Tween-20 and applied directly to the array for 1h incubation. Following 1h, samples were rinsed from the array surface by dipping 4 × each in PBS-Tween, PBS and deionized water, respectively. Washed arrays were reprobed with anti-mouse-IgG-AlexaFluor488 (10ug/mL) for 1h incubation. Following secondary incubation, arrays were washed again by dipping 4 × each in PBS-Tween, PBS and deionized water, respectively, and dried by centrifugation. Dried slides were scanned for 488 signal on an Innoscan 1100AL scanner (Innopsys) and signal intensities were calculated using Mapix (Innopsys) and graphed using Excel (Microsoft).

### Surface plasmon resonance

SPR was performed using a Biacore T200 instrument. GM3g-IME was immobillised onto an S-CM5 sensor chip as previously described (*39*). For analysis of Fab binding, serial delusions were sequentially injected at a flow rate of 10 μL/minute. An anti-c-Myc Fab (clone: 9E10) was used as a negative control.

### Isothermal titration calorimetry

Affinities of Fab BAR1 for the Amidine-GM3g and Lys–amidine-GM3g were measured by isothermal titration calorimetry using an automated PEAQ-ITC instrument (MicroCal) at 25 °C. Titrations were performed using 18 × 2 μL injections of 200-300 μM of the polysaccharide into 20-30 μM of the protein in PBS buffer. The heats of dilution measured from injection of the ligands into the buffer were subtracted, and titration curves were fitted with one-site binding model.

### Universal standard transfer analysis

All NMR experiments were recorded at 298K on a 950-MHz spectrometer with Bruker Avance III HD console and 5-mm TCI CryoProbe, running TopSpin 3.6.1 and using a SampleJet. All ligands in this work were first assigned using selective 1D Hartmann-Hahn TOCSY and HSQC experiments.

The uSTA experiments were either recorded with the same stddiffesgp.2 as previously described (*39*), or a pseudo 3D version that used an inputted file vdlist to increment the saturation times. The number of points were set to 32768 or 65536 and sweep width to 16.05ppm for an acquisition time of 2.150s and 4.300s. All spectra were processed using nmrPipe within the uSTA workflow as previously described (*39*), resulting in ‘transfer efficiencies’ that quantify the strength of the saturation transfer and inform on the binding pose in the complex.

*K*_D_ and k_off_ rates for amidine-lysine/WT BAR-1 complex were obtained by repeating the uSTA analysis over a range of protein/ligand concentrations and globally analysing the build-up curves for the NAc proton (**Fig S11**) as described previously (*39*). The interaction surface for the X-ray data was calculated from the structure using a <1/r^6^> expectation value between each proton in the ligand, and all protons in the protein, as described previously.

In Lys–amidine-GM3g, we would anticipate a range of R_1_ relaxation times which could affect the transfer efficiencies. To address this, we measured the R_1_ and R_2_ relaxation rates of each proton of the ligand and developed a correction that allowed us to rescale the transfer efficiencies to account to variations in the relaxation rate. The adjustments to the interaction surfaces by performing this operation were modest (**Fig S11**).

R_1_ and R_2_ relaxation rates were recorded using bespoke pulse sequences derived from the Bruker t1ir and cpmg sequences, with water suppression achieved by using excitation sculpting from the zgesgp sequence. The R_1_ experiment employed the zgesgp phase cycle had no cycle on the inversion pulse. The R_2_ experiment employed the zgesgp phase cycle with a y, -y pulse sequence on the CPMG refocusing pulse, which was performed before the water suppression sequence to avoid interference. In the final measurements, the interscan delay was set to 5s to allow significant relaxation of protons. Both relaxation spectra were recorded with 8 transients and 4 dummy scans per increment, 65536 acquisition points and a sweep width of 15.96 ppm (950Mhz) for an acquisition time of 2.163 s. Spectra to obtain R_1_ were acquired with 13 delays: 5, 0.001, 0.05, 0.1, 0.25, 0.5, 0.8, 1, 1.5, 2, 3, 4 and 5s. Spectra to obtain R_2_ were recorded with 12 delays using a spin-echo time of 800us (2×400us) per cycle and 0, 400, 40, 80, 120, 160, 200, 240, 280, 320, 360 and 400 cycles. The 90-degree 1H times were calibrated manually.

To perform the correction of the transfer efficiencies, we first took the fitting parameters obtained from the full *K*_D_ analysis of Lys–amidine-GM3g / BAR-1 complex interaction. The R_1_ rate for the NAc proton in the ligand (0.37 s^-1^) was essentially identical to that measured using the R_1_ experiment (0.4 s^-1^), providing independent support for our analysis. We then simulated the transfer efficiencies expected as we systematically vary R_1_ and R_2_. The expected transfer efficiency was largely invariant of R_2_, but could vary by a factor of 2 as R_1_ varies by one order of magnitude. We used these curves to interpolate the expected transfer efficiency as both R_1_ and R_2_ tend to zero, thus to a reasonable first approximation, removing variation in ligand relaxation between atoms from the uSTA measurement. The R_1_ correction was more significant than the R_2_ correction. The largest variation of R_1_ in the dataset was a factor of 3.5, ranging from 0.4 s^-1^ (NAc) to 1.4 s^-1^, leading to only modest adjustments of the transfer efficiency.

### X-ray crystallography

BAR-1 Fab was loaded onto a gel filtration Superdex 200 column (GE Healthcare) in 10 mM Tris-HCl, pH 7.5, 150 mM NaCl. Co-crystals appeared at 20 °C after a few weeks from a hanging drop of 0.1μL of protein solution (19 mg/mL with 1.2 mM Lys– amidine-GM3g) with 0.1 μL of reservoir solution containing 30% (w/v) PEG MME 2000, 0.1 M potassium thiocyanate in vapor diffusion with reservoir. Crystals were frozen with the same solution containing 16% glycerol and 4 mM Lys–amidine-GM3g . Data were collected at the Diamond light source oxfordshire (beamlines I24). Data were processed with XIA2 (*56–60*). Structure has been solved by molecular replacement using PHASER and pdb file 6ug7 (*45*), and the structure was builded with Autobuild program, refined with REFINE of PHENIX with NCS restraints (*61*) and adjusted with COOT (*62*). Coordinates and topologies of ligands were generated by PRODRG (*63*) Structures were refined at 1.9 Å. Two Fab molecules are present in the asymmetric unit (H/L and A/B). The two molecules are very similar (rmsd of 0.4852 Angstroems for 213 residues). Final refinement statistics are given in Table 1. Atomic coordinates and structure factors have been deposited in the Protein Data Bank (8BJZ). The quality of all structures was checked with MOLPROBITY (*64*). The Ramachandran statistics are as follows: 98% favoured and 2% allowed.

### Data processing and statistical evaluation

Flow cytometry data was evaluated in FlowJo V.10.8. Graphs were generated in GraphPad Prism V9.4 and using GCTree (*55*), and later edited in Adobe Illustrator for aesthetics. Statistical analysis was conducted either in GraphPad Prism V.9.4 or in RStudio V.4.1. Chao1 estimates were performed using the RStudio iNEXT package (*65*). Statistical test details are provided in the results, figures and associated figure legends.

## Acknowledgments

This work was supported by the Rosetrees Trust Interdisciplinary Award ID2020/100023. Upgrades of 600-MHz and 950-MHz NMR spectrometers were funded by the Wellcome Trust (grant ref: 095872/Z/10/Z) and the Engineering and Physical Sciences Research Council (grant ref: EP/R029849/1), respectively, and by the University of Oxford Institutional Strategic Support Fund, the John Fell Fund, and the Edward Penley Abraham Cephalosporin Fund. This project has received funding from the European Research Council (ERC) under the European Union’s Horizon 2020 research and innovation programme (grant agreement 101002859). E.S. is supported by Swedish Research Council Starting Grant (2020–02682). QJS is a Jenner Vaccine Institute Investigator and a James Martin School Senior fellow. The Chemistry theme at the Rosalind Franklin Institute is supported by the EPSRC (V011359/1 (P)).

## Author contributions

Conceptualization of project: LPD, TS, BGD, QJS

Methodology: LPD, XX, AK, LM, CLB, HS, CG, RS, KL, ES, JP, AJB, JN, TS, BGD, QJS

Investigation: LPD, XX, AK, LM, CJB, RM, HS, CG, RS, KL, RAR, ES

Funding acquisition: BD, QJS

Project administration: BD, QJS

Supervision: ES, JP, AJB, JN, BGD, QJS

Writing – original draft: LPD

Writing – review & editing: LPD, BGD, QJS

---

**Figure S1:**
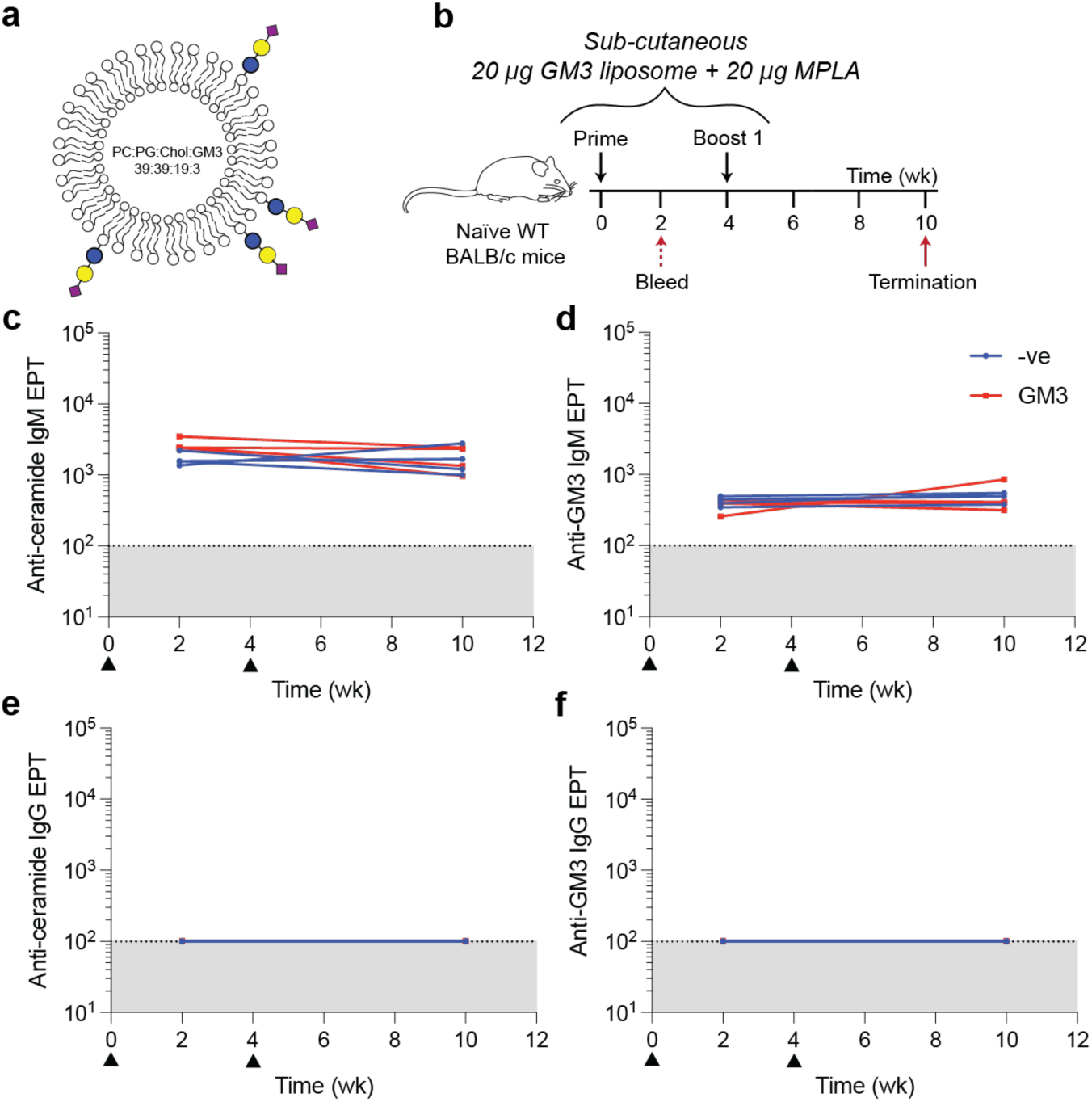
Immunorecessiveness of GM3 liposomes in mice. **(a)** Liposomes were synthesized both with and without GM3. **(b)** Immunization schedule. **(c-f)** Serum IgM and IgG reactivity was screened via direct ELISA against both ceramide and GM3 over the immunisation period.

**Figure S2:**
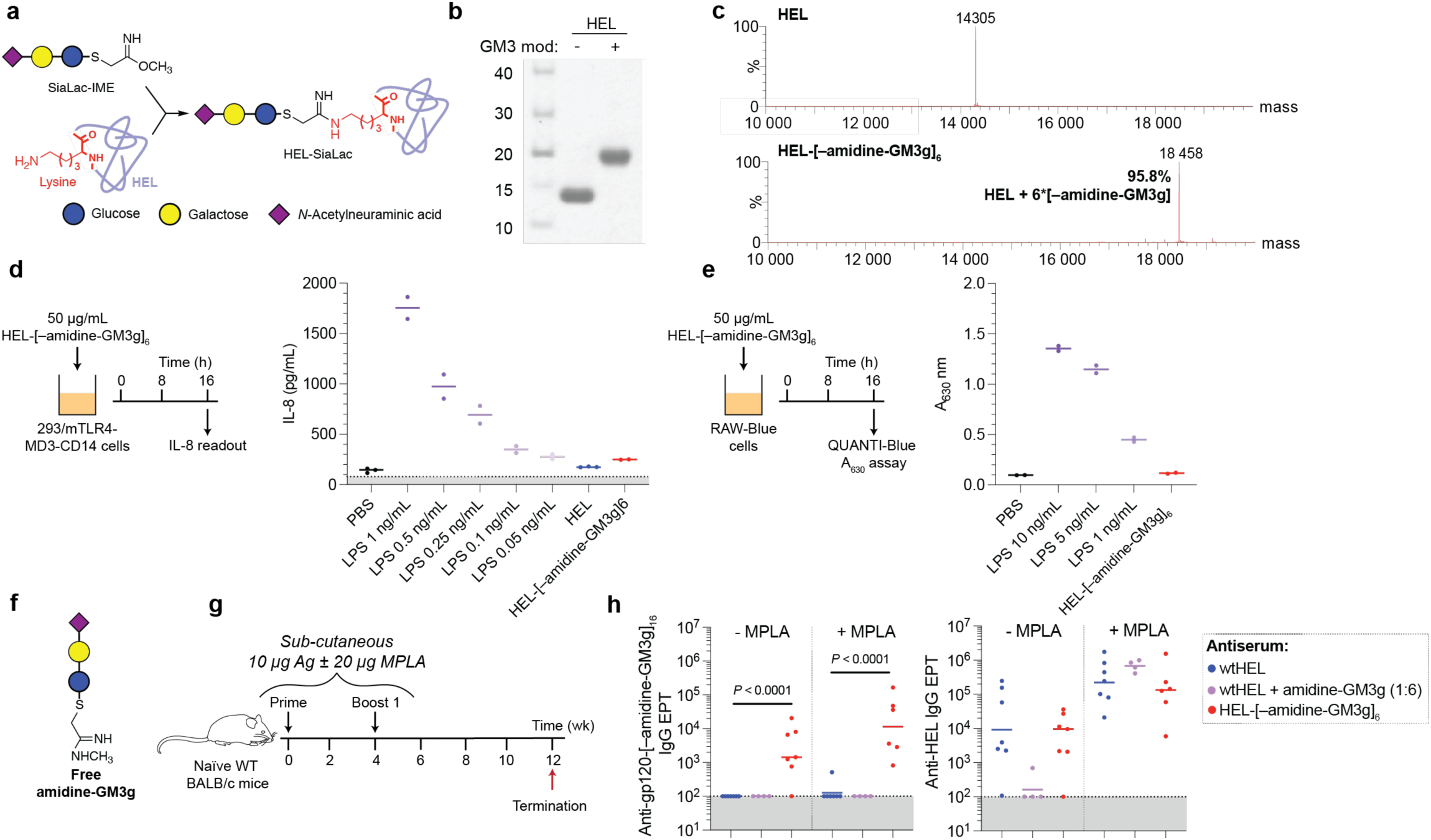
Chemical and immunological characterisation of the amidine-GM3g LOG. **(a)** Overview of HEL-[–amidine-GM3g]6 synthesis. **(b)** SDS-PAGE of HEL following [–amidine-GM3g] conjugation. **(c)** Mass spectra of HEL-[– amidine-GM3g]_6_ sample. **(d)** LPS contamination was tested via incubating 293/mTLR4-MD3-CD14 cells with HEL-[–amidine-GM3g]6. IL-8 production was evaluated via ELISA. **(e)** Broader endotoxin contamination was screened using RAW-Blue cells. **(f)** Free amidine-GM3g design. **(g)** Immunisation schedule. **(h)** Terminal IgG endpoint titres. Data were compared via Tukey’s post-hoc multiple comparison test.

**Figure S3:**
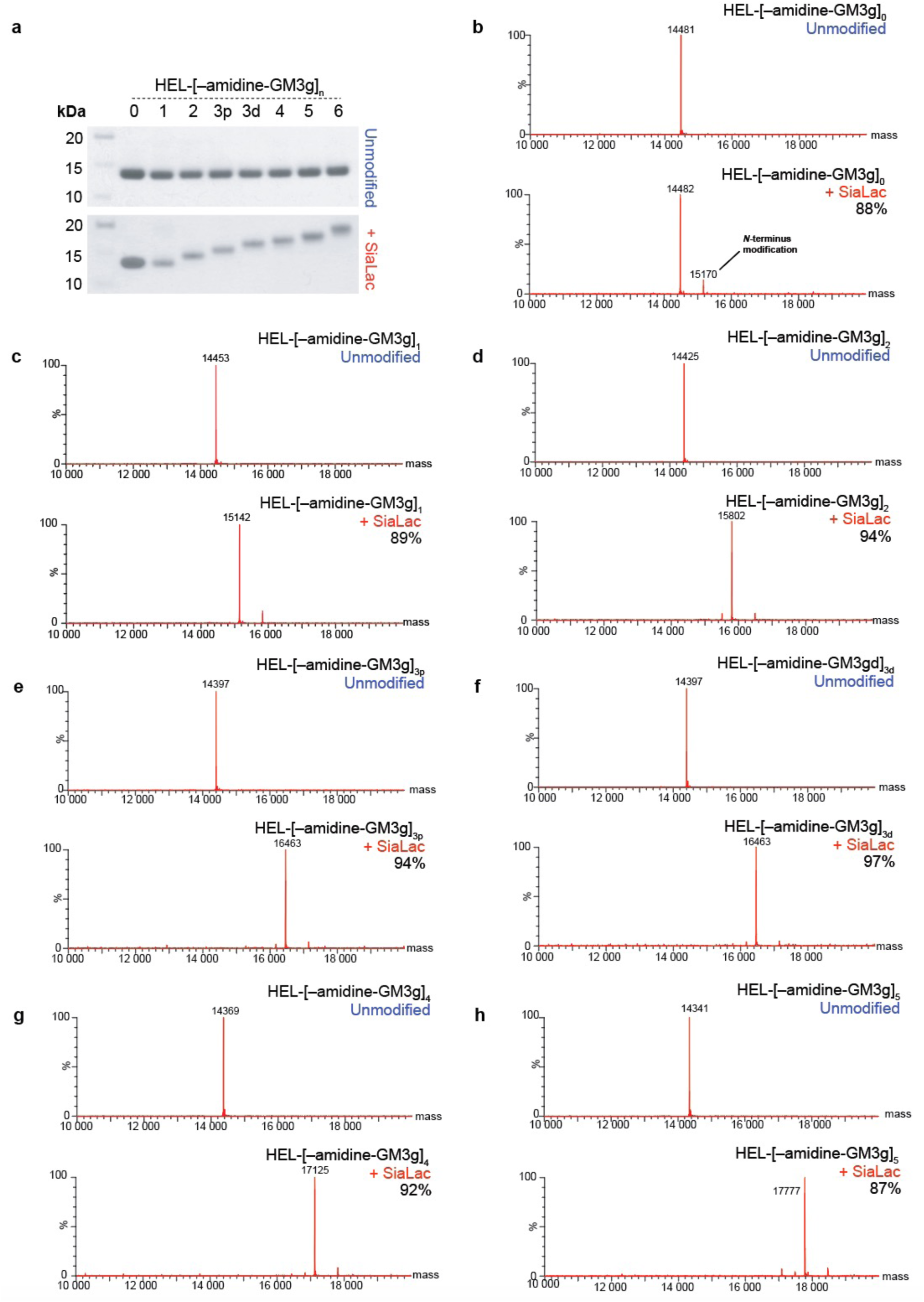
Chemical characterization of amidine-GM3g-modified HEL mutants. **(a)** SDS-PAGE of the purified HEL mutants and their GM3g-modified counterparts. **(b–h)** Mass spectra of the modified HEL mutant products. Refer to Figure 1f for K->R mutation code.

**Figure S4:**
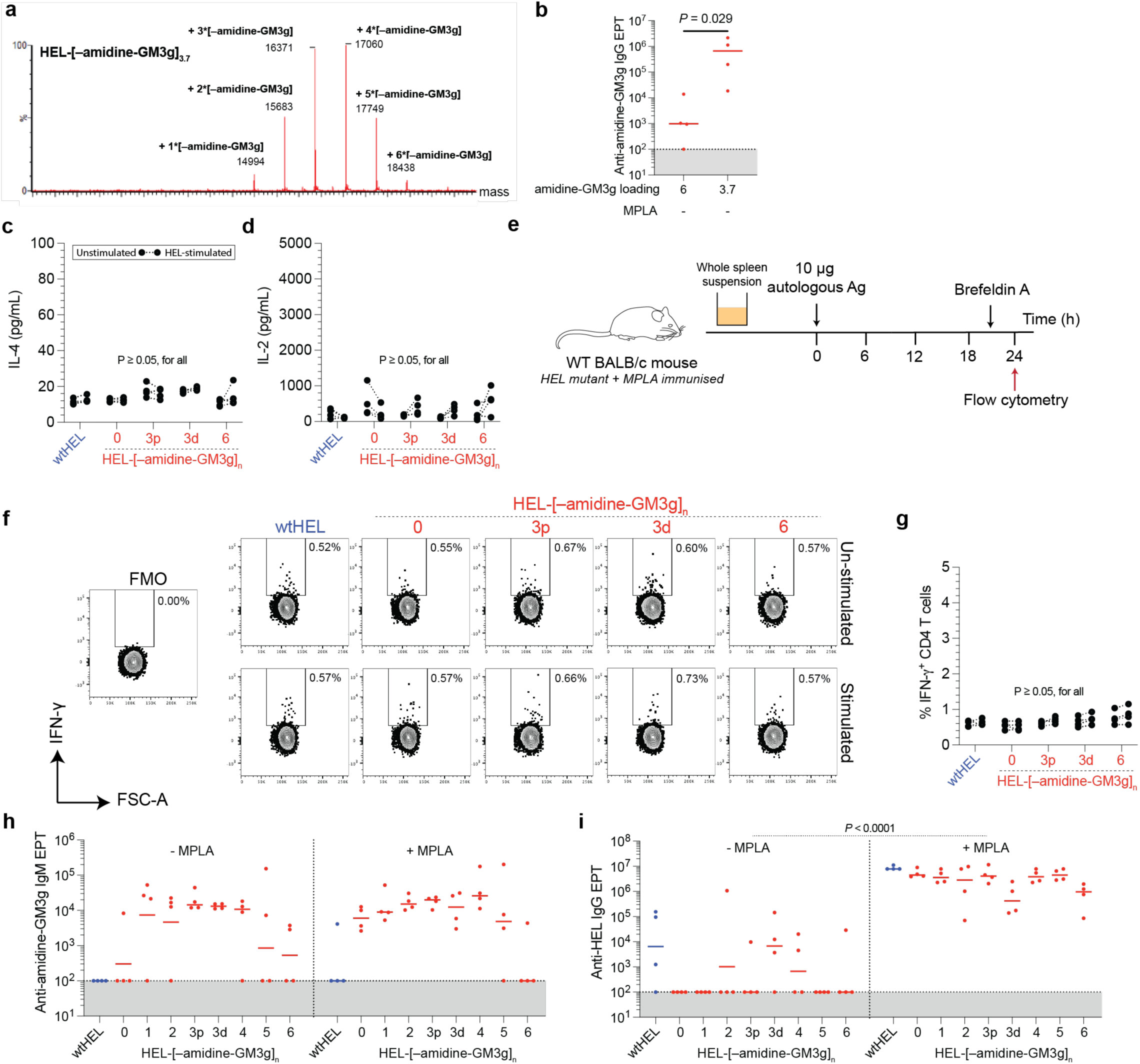
Dissecting whether modifications to the HEL protein backbone implicate Th responses. **(a)** Mass spectra of wtHEL partially modified with amidine-GM3g, producing HEL-[–amidine-GM3g]3.7. **(b)** Terminal gp120-[amidine-GM3g]16-reactive IgG endpoint titers of mice primed and boosted with HEL-[–amidine-GM3g]3.7. **(c,d)** Whole splenocytes of animals immunized with amidine-GM3g-modified HEL mutants were stimulated *in vitro* for 72 h and cytokine release in supernatant was screened. **(e–g)** Intracellular IFN-ᵧ was detected via flow cytometry on pre-gated CD4^+^ cells. **(h)** IgM endpoint titres were screened against gp120-[–amidine-GM3g]16 two-weeks post-prime. **(i)** Terminal IgG-specific IgG. Data were compared using Dunn’s tests, except **(i)** where two-way ANOVA contrasted adjuvant and sugar loading effects.

**Figure S5:**
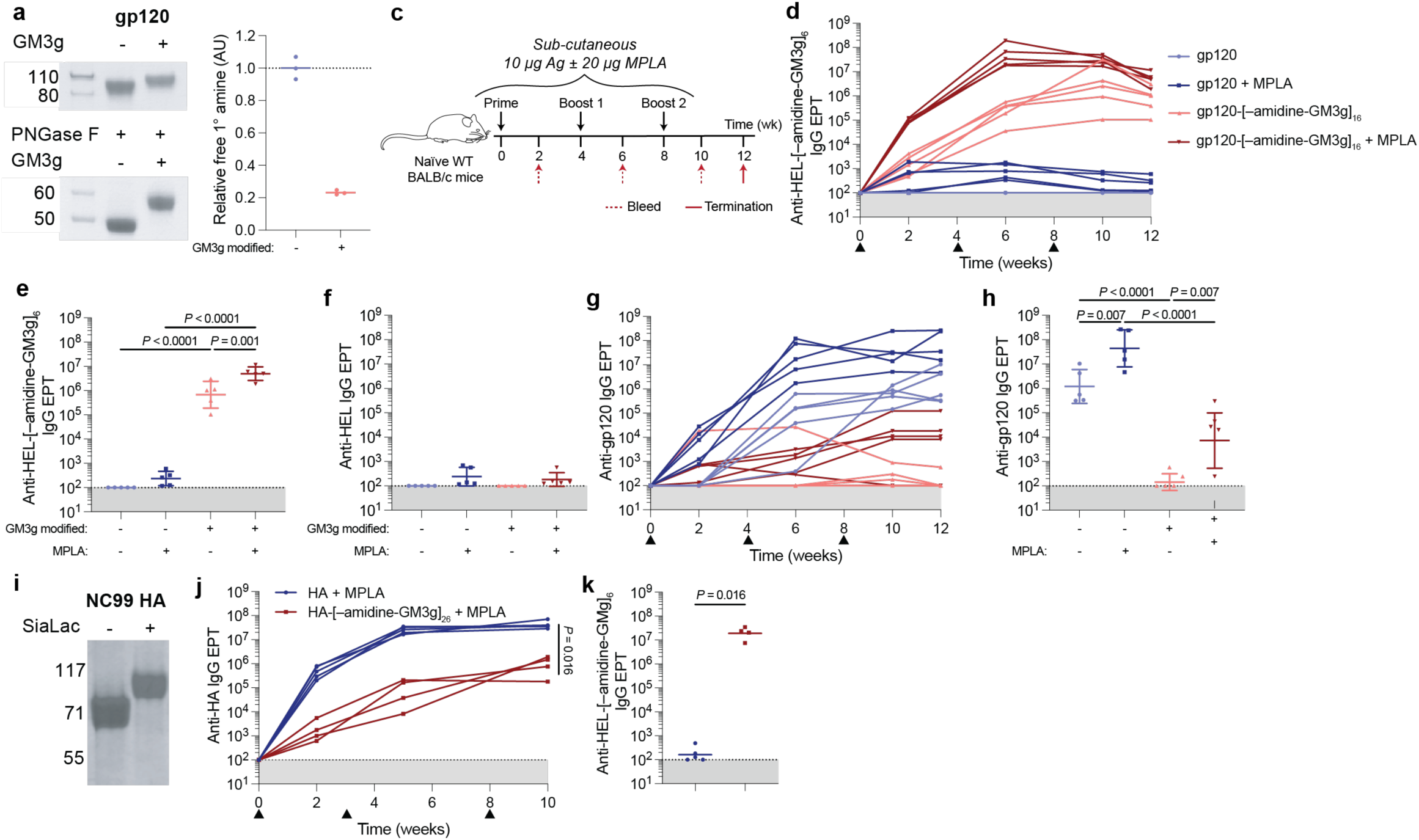
LOG-specific antibody responses occur against amidine-GM3g across multiple protein carrier proteins. **(a)** SDS-PAGE of amidine-GM3g modified and unmodified proteins pre- and post-PNGase F treatment. **(b)** Free amine ELISA post-LOG modification. **(c)** Immunisation schedule. **(d–h)** Longitudinal or terminal serum IgG endpoint titres against LOG-specific and protein carrier constructs in animals immunised with gp120-[–amidine-GM3g]16. Data were evaluated using a post-hoc Tukey’s test. **(i)** SDS-PAGE of IAV-derived H1N1 (NC99) HA post-GM3g modification. **(j,k)** Longitudinal and terminal IgG endpoint titres. Data were evaluated using Mann-Witney tests.

**Figure S6:**
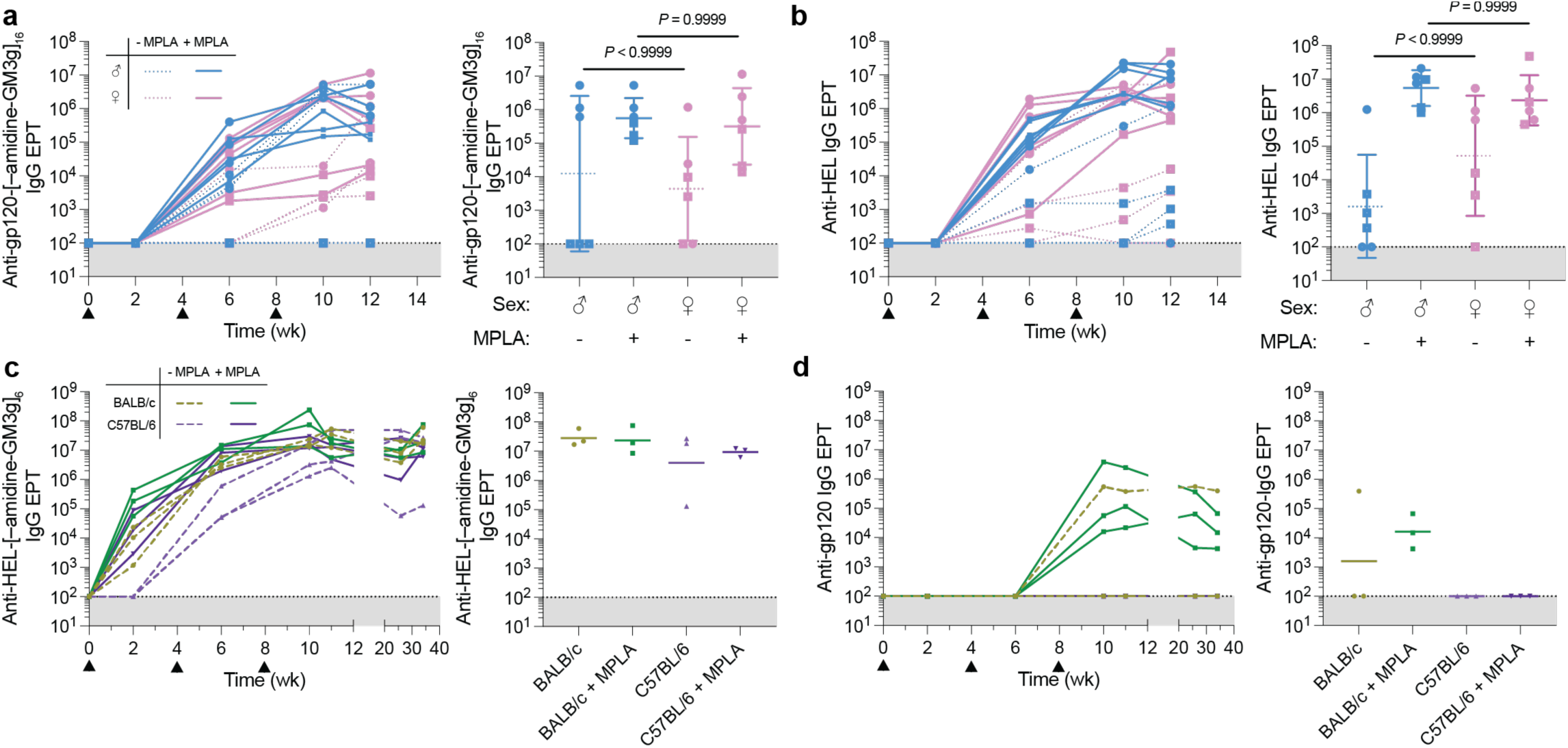
Evaluating the sex and murine background effects on the anti-amidine-GM3g LOG response. **(a,b)** Male and female WT BALB/c mice were immunised three times (▲) with 10 µg HEL-[–amidine-GM3g]6 ± 20 µg MPLA. Both LOG and protein backbone-specific serum IgG endpoint titres were determined both longitudinally and at the terminal timepoint. Data were compared via Dunn’s multiple comparison test. **(c,d)** BALB/c and C57BL/6 mice were immunised with 10 µg gp120-[–amidine-GM3g]16 ± 20 µg MPLA. Serum IgG endpoint titres against antigen components, LOG and protein backbone, were measured.

**Figure S7:**
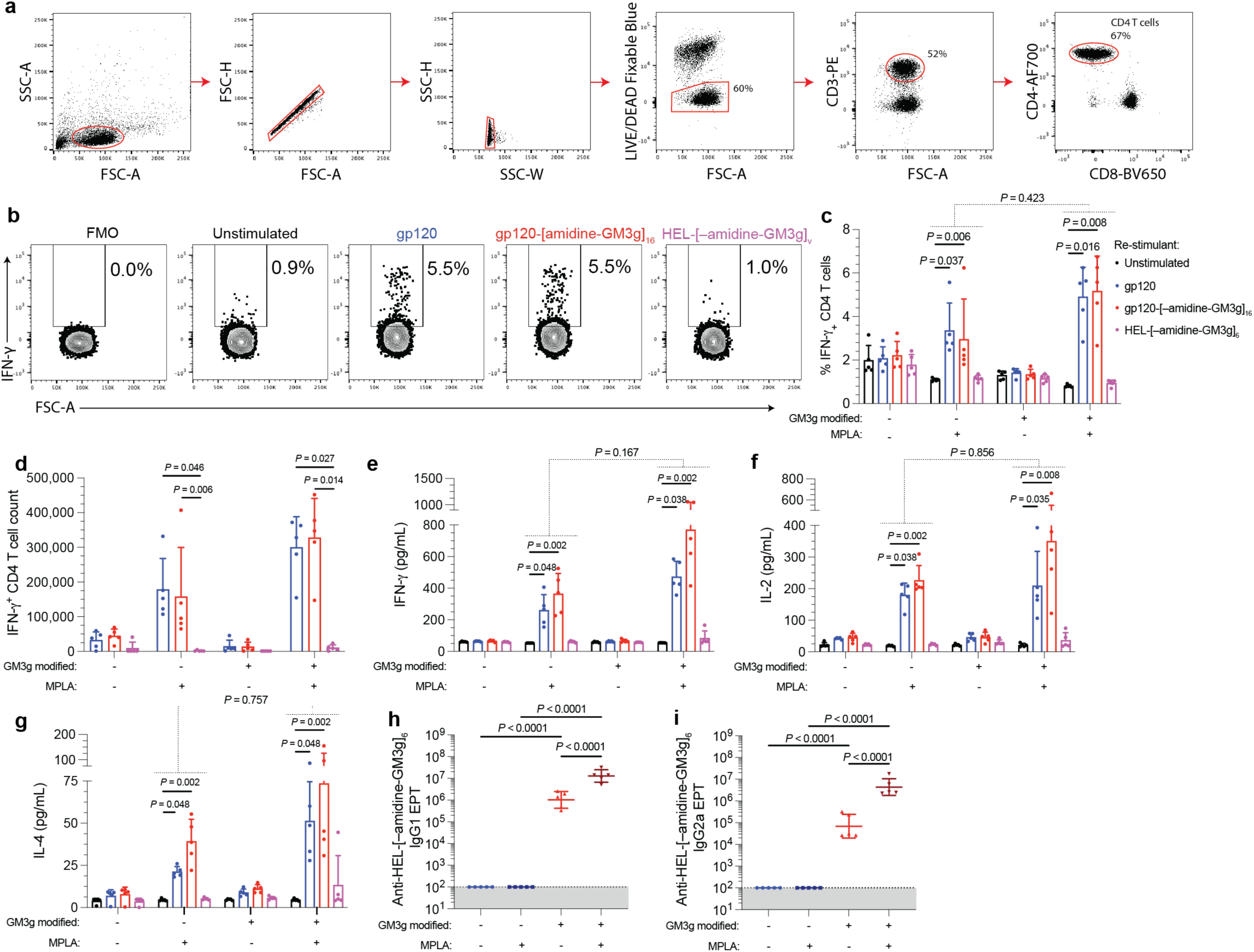
Th cell recall responses in mice immunized with gp120-[–amidine-GM3g]16. **(a–d)** Intracellular cytokine staining was performed on splenocytes of immunised mice, restimulated *in vitro* with different protein antigens, as indicated. IFN-y production among CD4^+^ T cells was compared between vaccination and restimulatory conditions. **(e–g)** Cytokine release was similarly compared in splenocytes restimulated for 72 h via ELISA. **(h,i)** Serum LOG-specific IgG subclass endpoint titres were measured via ELISA. Data were compared pairwise via Tukey’s post-hoc test. Establishment of an interaction effect between vaccination and restimulatory conditions were determined via two-way ANOVA.

**Figure S8:**
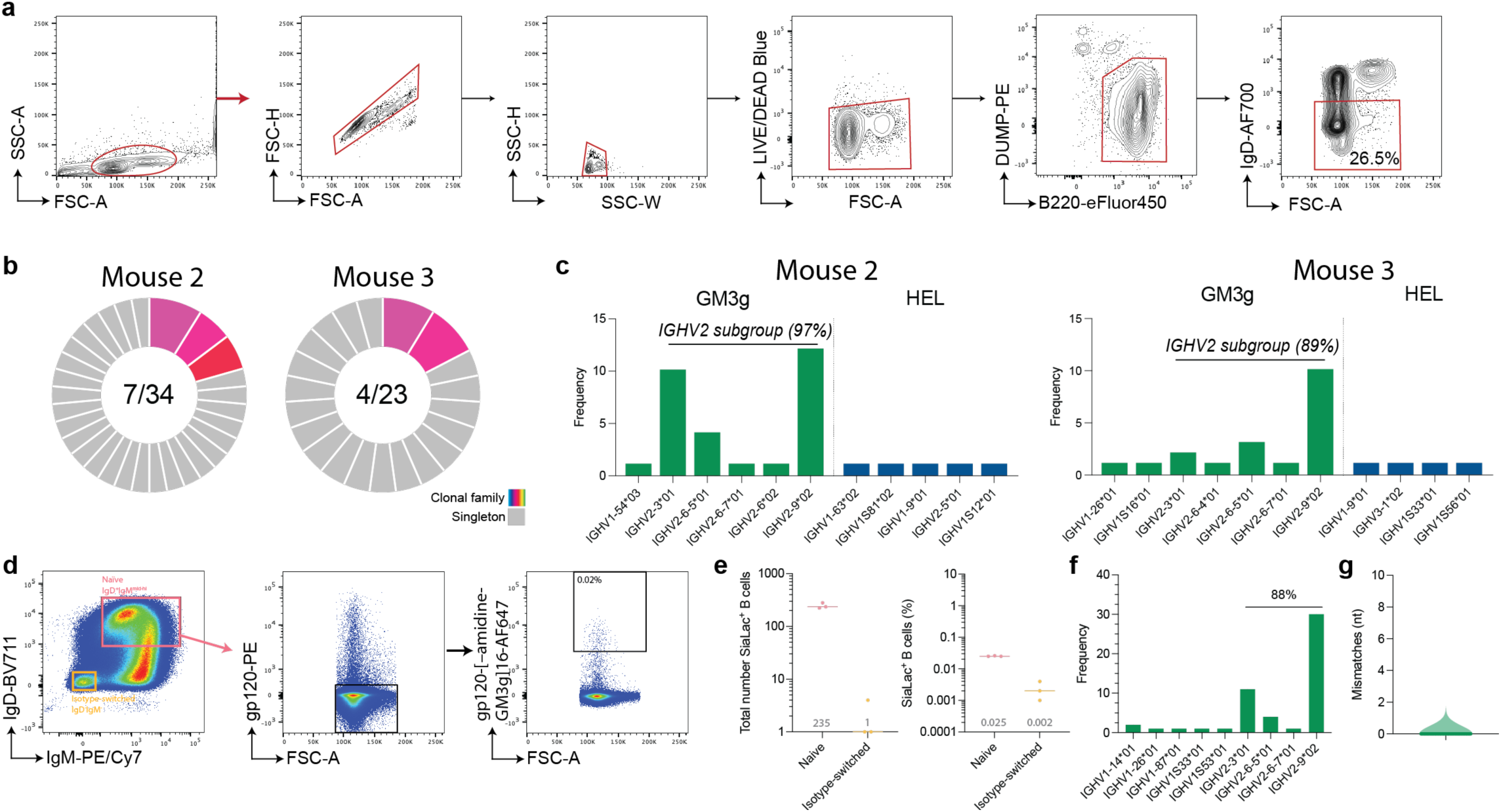
Clonotyping of HEL-[–amidine-GM3g]6-immunized mice. **(a)** Gating strategy for IgD^-^ B cells. **(b)** Clonal family clustering and **(c)** *IGHV* gene-segment utilisation in mice primed with HEL-[–amidine-GM3g]6. **(d)** Gating strategy to identify the antigen-specific naïve B cell population from splenocytes. **(e)** The absolute number and percentage of [–amidine-GM3g]^+^ B cells. **(f,g)** Heavy chain V-regions were recovered and sequence-validated from one mouse, confirming their clonotypic origins and GC inexperience.

**Figure S9:**
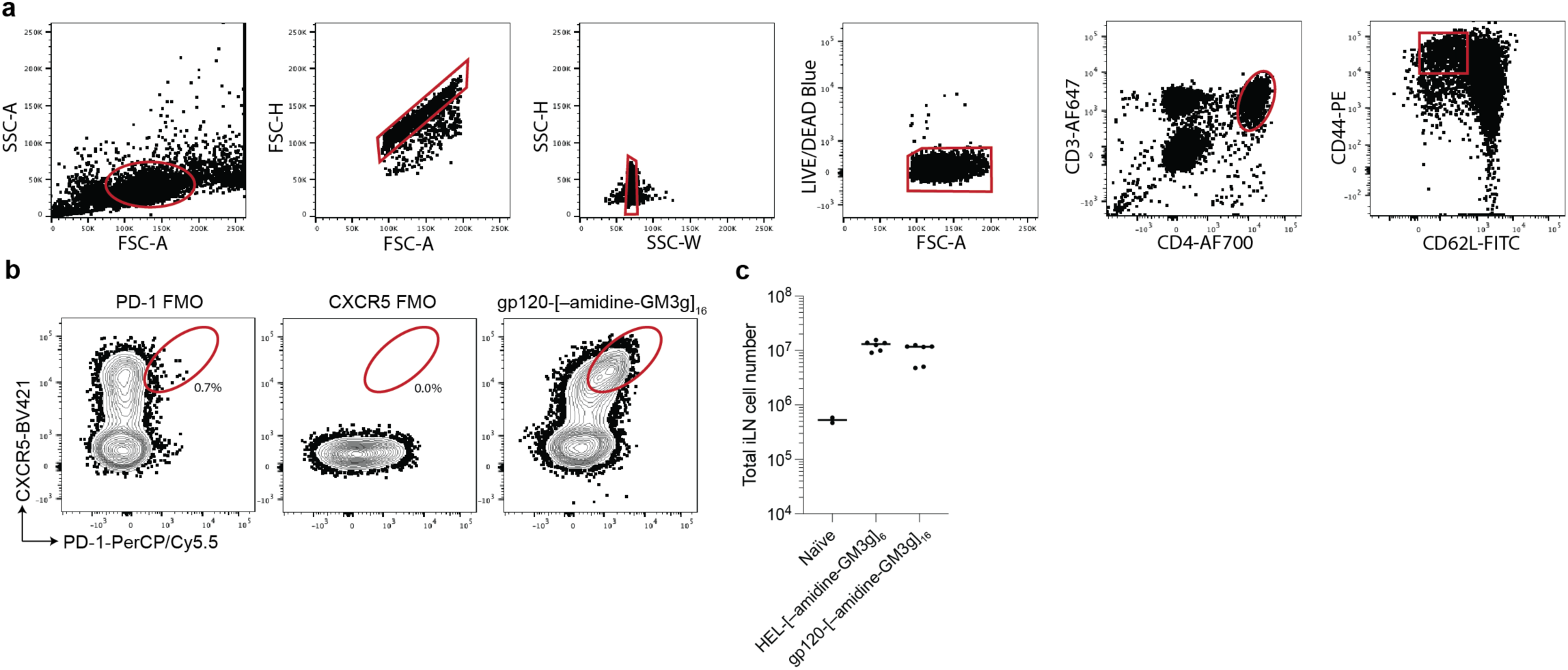
Determining the backbone-dependent Tfh population size in LOG-immunized animals post-prime. **(a)** Gating strategy for CD4^+^CD62L^-^CD44^hi^. **(b)** Representative FACS plots for determining the Tfh population. **(c)** Absolute total cell count in iLNs.

**Figure S10:**
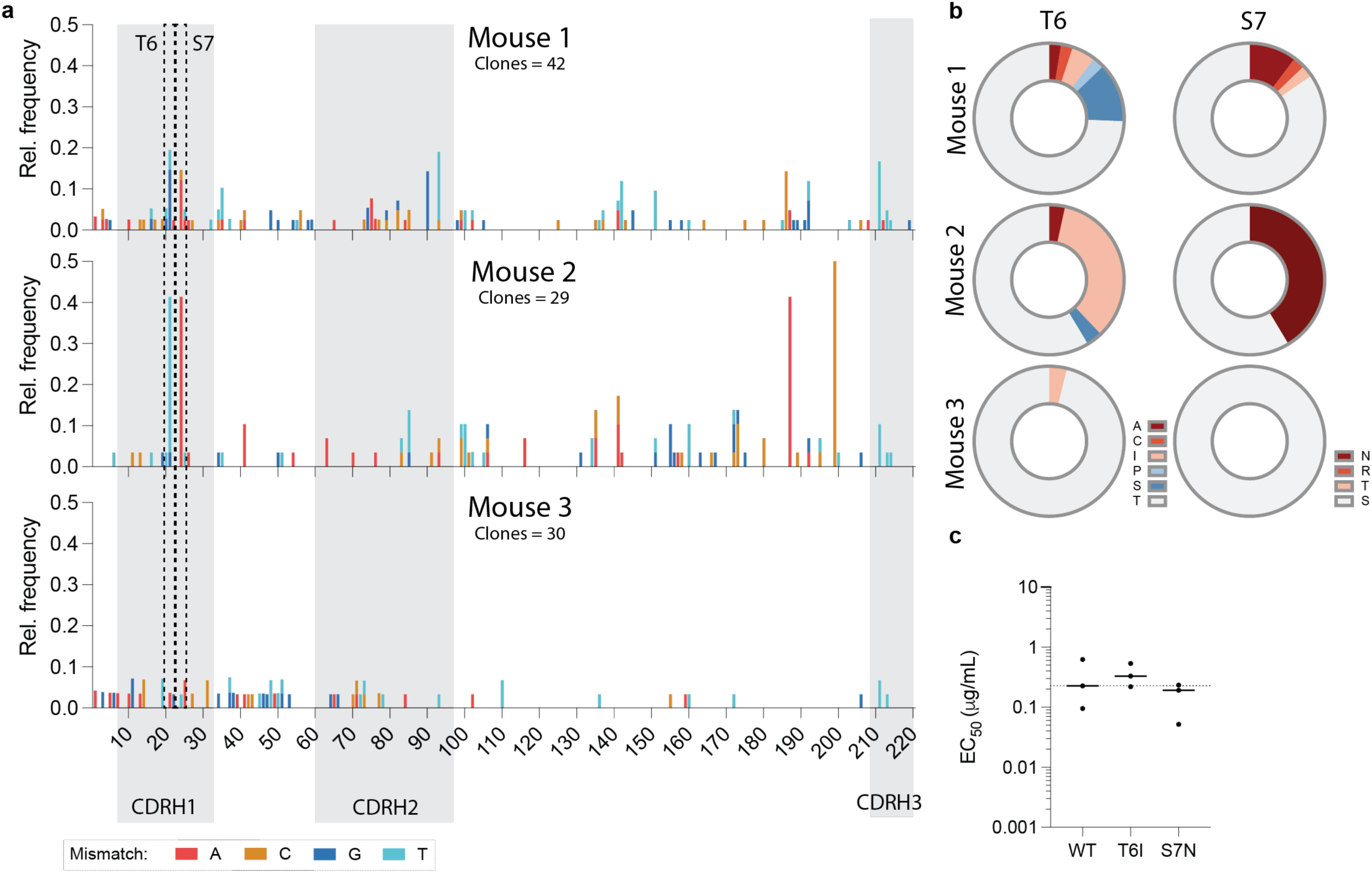
Mutation frequencies observed across the gp120-[–amidine-GM3g]16-raised *IGHV2* subgroup population. **(a)** Manhattan plot of the nucleotide mismatches from all isolated IGHV2-origin GM3g-binding B cell raised against the gp120-[–amidine-GM3g]_16_ LOG. **(b)** Substitutional implications at mutation hotspot codons, where the wild-type encodes T6 and S7. **(c)** The most common substitions were mutated into the WT BAR-1 sequence and their relative binding against gp120-[–amidine-GM3g]_16_ was compared via ELISA.

**Figure S11:**
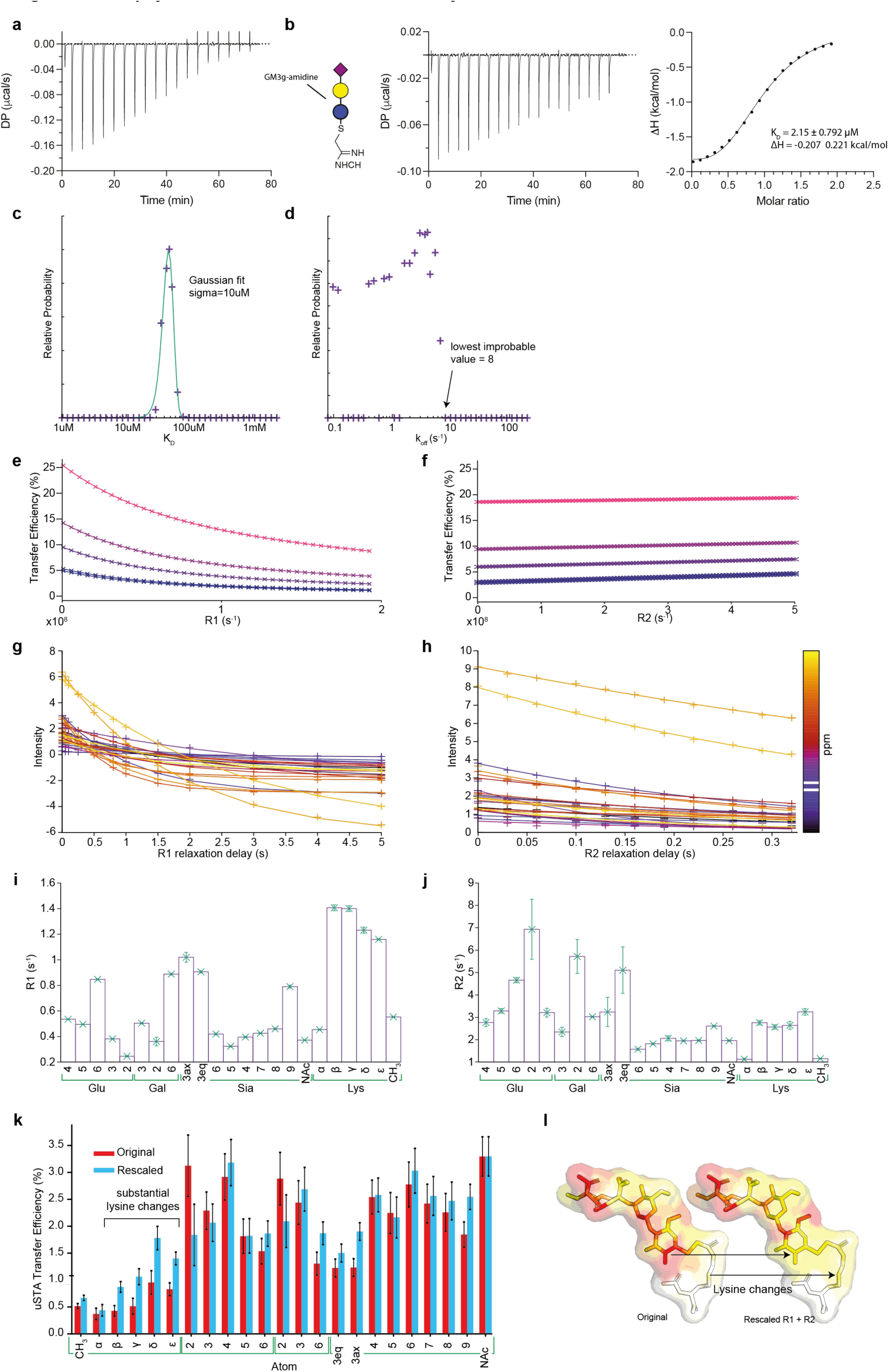
Details of Lys–C(NH)NH-GM3g•BAR1 complex by uSTA NMR. **(a)** Raw titration data of BAR-1 Fab against Lys–amidine-GM3g. **(b)** ICT performed against amidine-GM3g. **(c)** Results of the Bloch-McConnell fitting of BAR-1 with Lys–C(NH)NH-GM3g. **(d)** These reveal good quality fits of the data. Iteratively changing and fixing the *K*D value, refitting the data and following the variation in the probability of the model being correct (exp(-chi^2^/2)) allows construction of an error surface. To an excellent approximation, the variation in the fitted *K*D follows a gaussian distribution **(e)**. Performing the same analysis on the koff parameter resulted in a non-central distribution, indicating that in this case, while *K*D is well determined, *koff* is not. The distribution is reasonably interpreted by a log-normal distribution, resulting in the most probable value being 3.77 s^-1^ but with asymmetric error bars, +4 s^-1^, -2 s^-1^. The distribution can be interpreted as placing a limit on koff, such that koff <8 s^-1^. **(f,g)** R1 and R2 relaxation rates were obtained for each proton in amidine-lysine. The variation in relaxation rates approximately by a factor of 3, prompted us to consider the effects of this on the transfer efficiency. Notably, the R1 determined from the *K*D analysis for the NAc proton (0.37 s^-1^) was consistent with the value measured directly and independently (0.4 s^-1^) supporting the quantitative uSTA analysis. **(h,i)** The simulated parameters from the *K*D analysis in **c** were used to simulate the variation in transfer efficiency as a function of R_1_ and R_2_, revealing almost no variation with R_2_, but a modest variation with R_1_. **(j)** These curves were interpolated using a biexponential function for R1 and a linear function for R2, and were used to provide a rescaling factor to adjust the transfer efficiencies of each atom to the value expected if relaxation was identical to the NAc proton. The largest correction was for the lysine delta proton (R_1_ 1.4 s^-1^) which was furthest from the NAc R1 (0.4 s^-1^). In this extreme case, the correction to the transfer efficiency was a factor of 2. **(k)** The original and rescaled interaction surfaces for Lys–C(NH)NH-GM3g. The overall pattern observed is largely invariant of the rescaling, with some positions varying more than others. The main conclusions drawn from inspection of the surface, that the NAc methyl group and the sialic acid moiety dominate the interaction, that protons in all GM3g sugars are important, and that the lysine does not contribute substantially to the interaction are independent of the relaxation correction. In the manuscript, all interaction surfaces shown have had the transfer efficiencies ‘corrected’ using this method.

**Figure S12:**
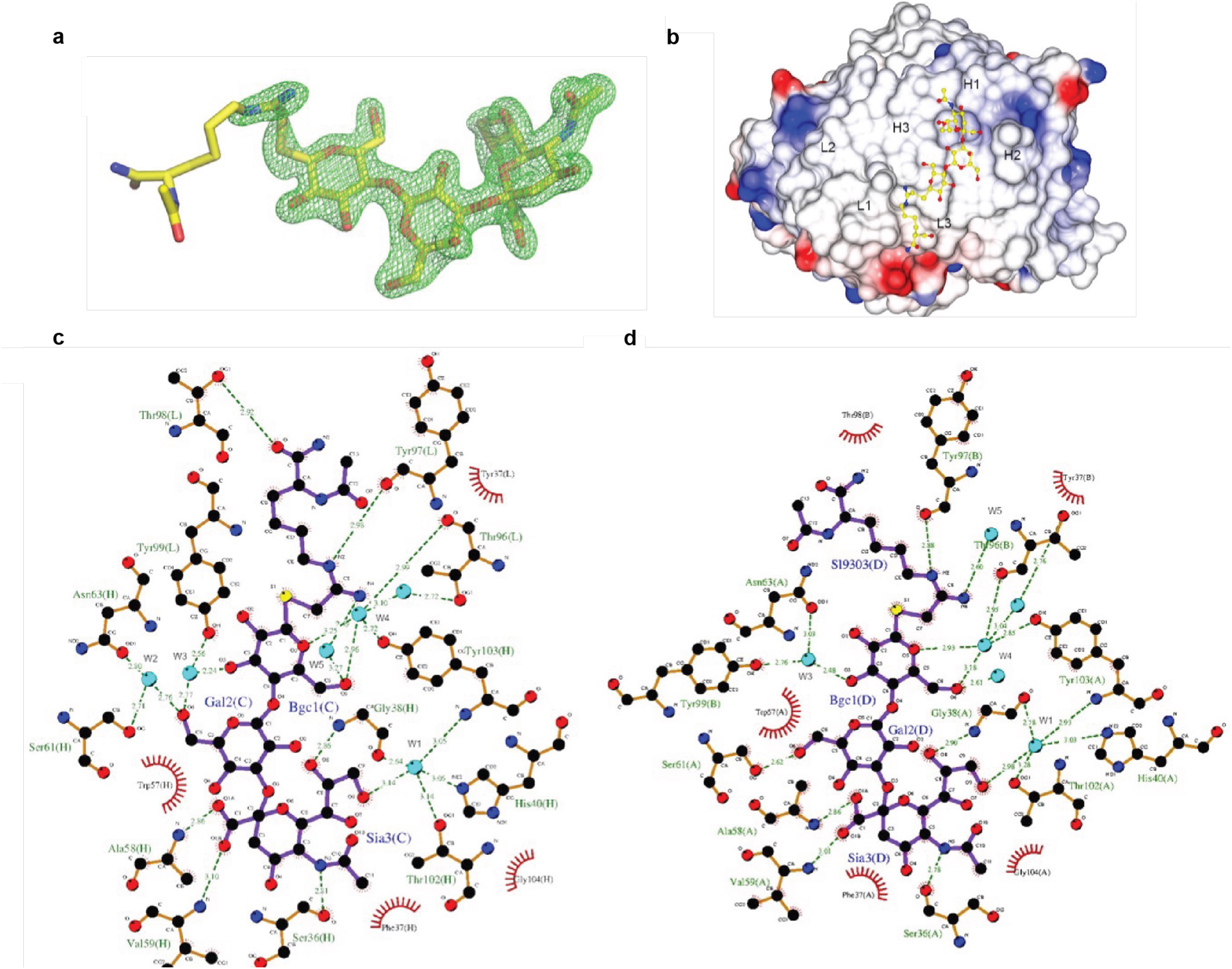
Details of the X-ray structure of Lys–C(NH)NH-GM3g•BAR1. **(a)** FO–FC electron density omit map at 3 σ around SiaLac-amidine-Lys molecule. SiaLac-amidine-Lys is shown as sticks with carbon atoms coloured in yellow, nitrogen in dark blue and oxygen in red. **(b)** Surface of the binding side of BAR-1/SiaLac-amidine-Lys complex structure. The surface of Bar-1 is colored by electrostatic charges calculated in CCP4MG (red for negative potential, white for neutral and blue for positive). SiaLac-amidine-Lys is shown as sticks with the carbon in yellow. CDR loops have been labelled. **(c,d)** Ligplot diagrams illustrating BAR-1/siaLac-amidine-Lys interactions for chain H/L and A/B. Covalent bonds of the polysaccharide and the protein residues are in purple and brown sticks, respectively. Hydrogen bonds are represented by green dashed lines and hydrophobic contacts are shown as red semi-circles with radiating spokes

**Figure S13:**
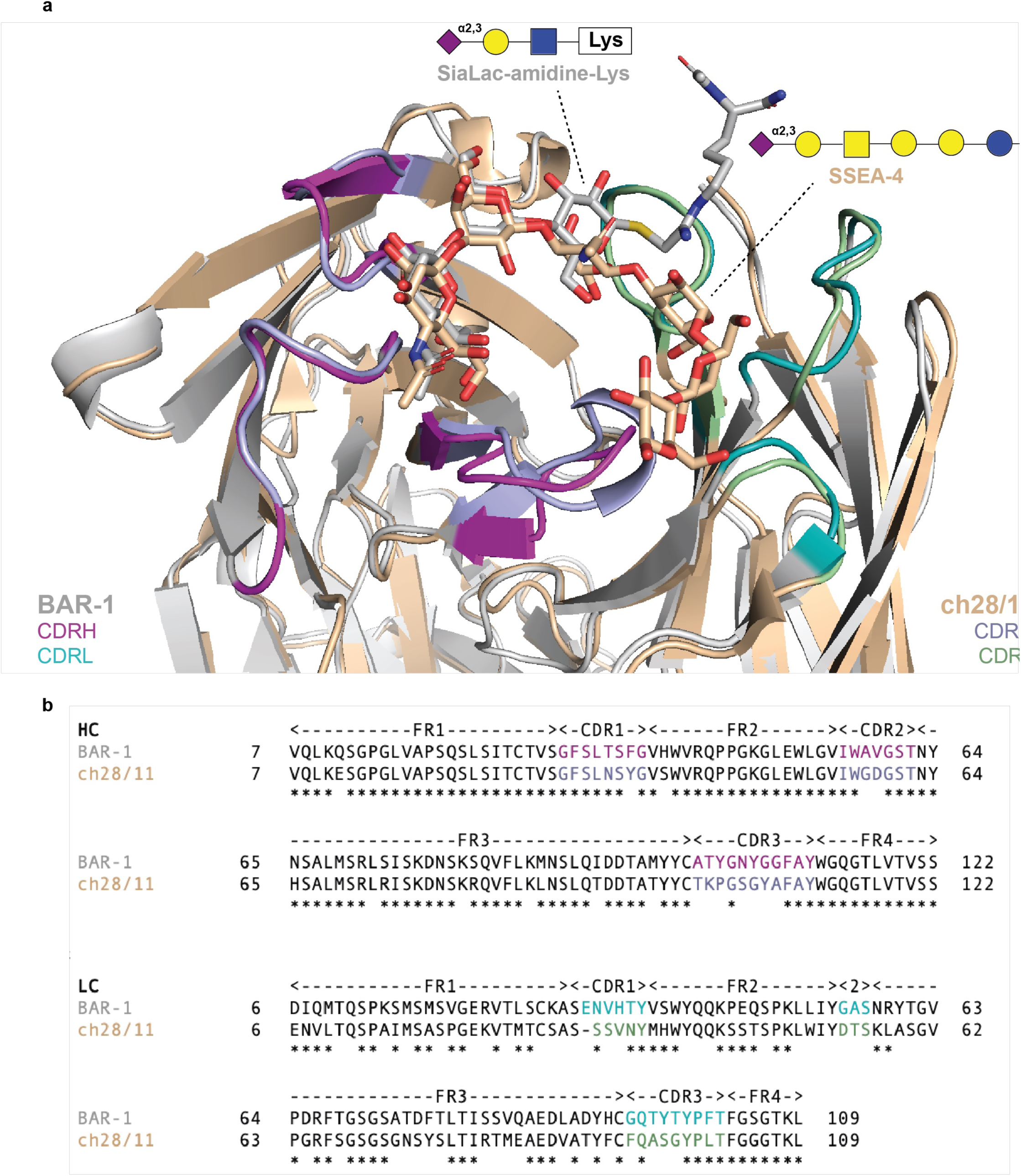
Comparison between BAR-1 and ch28/11. **(a)** Structural alignment of the BAR-1 x-ray structure bound to Lys–C(NH)NH-GM3g and that of ch28/11 to SSEA-4. (b) Sequence alignment of the two structures. Highlight similar SiaGal-recognition pattern as indicated by similar CDRH1/2 motifs.

**Figure S14:**
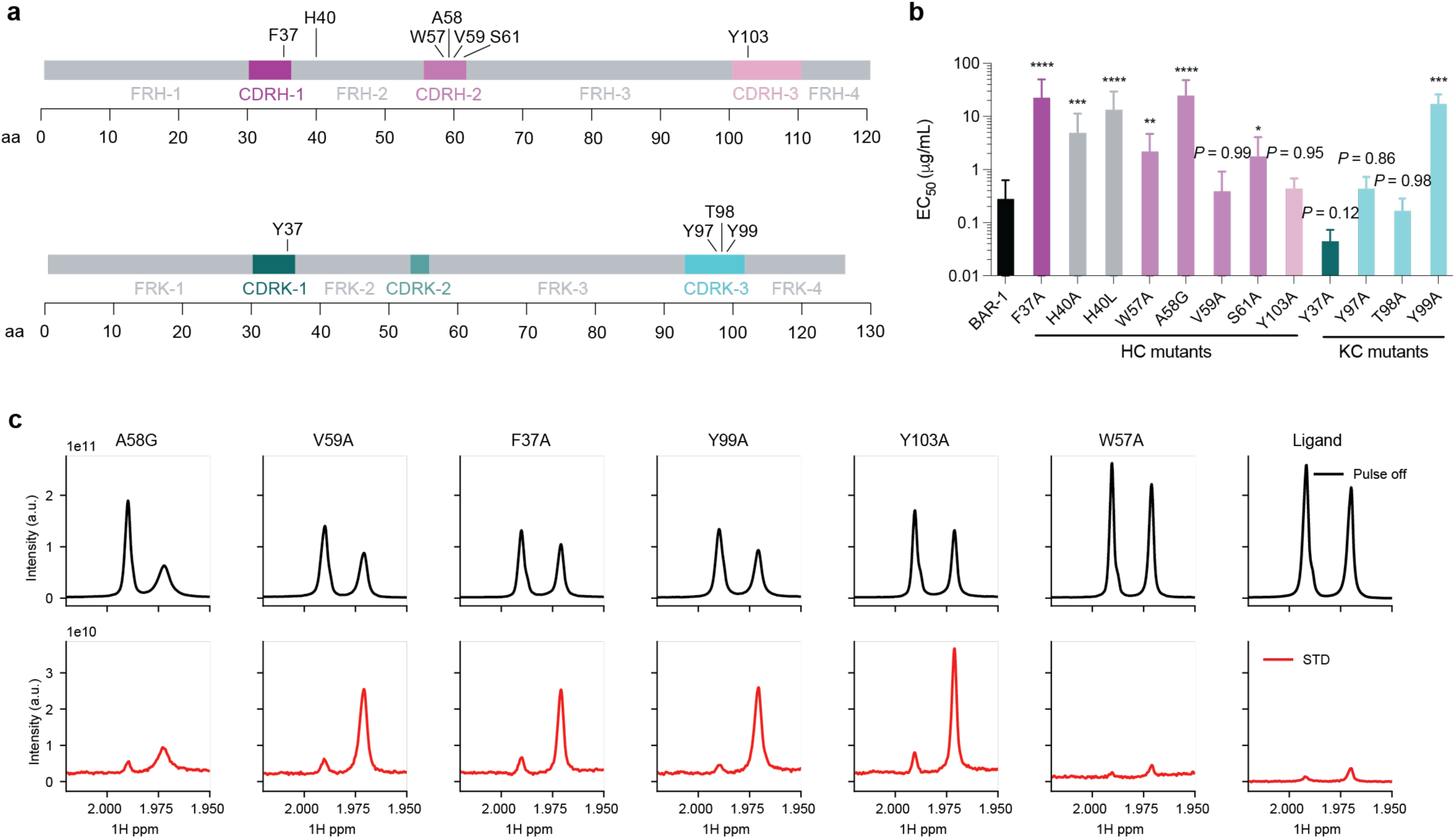
Alanine scanning of BAR-1. **(a)** Sequence schematic of BAR-1 and select residues targeted for mutagenesis. **(b)** ELISA EC50 binding was compared against gp120-[–amidine-GM3g]16 binding (*n* = 4). Data were compared via Tukey’s post-hoc multiple comparison test. *P*-value denotations: ’****’ *P* < 0.0001, ’***’ *P* < 0.001, ’***’ *P* < 0.01 and ’*’ *P* < 0.05. **(c)** ‘Pulse off’ 1D NMR (black) and saturation transfer difference (STD) spectra for the various BAR-1 mutants considered, showing specifically the distinctive NAc methyl groups that terminate the Lysine moiety (Left hand peak) and the Sialic acid (Right hand peak).

