## Supplementary Information for "Engineered display of ganglioside-sugars on protein elicits a clonally and structurally constrained B cell response"

#### Synthetic notes

#### I. HEL-[–amidine-GM3g]

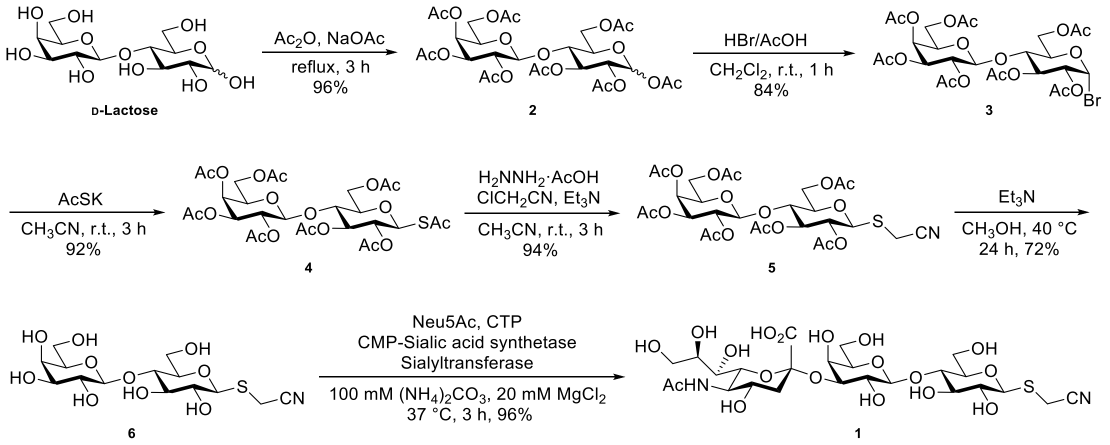


**Scheme I-1.** Chemoenzymatically synthetic route of cyano-functionalized 3′-SiaLac **1**.

As shown in **Scheme I-1**, starting from commercially available D-lactose, acetylation in a combination of $Ac_2O$ and NaOAc in reflux gave lactose octaacetate **2** with a yield of 96%, and treated acetate **2** with acetic HBr solution at room temperature for 1 h, yielding the bromosugar **3** in 84% after recrystallization from chilled ether.

The conversion from **3** to thioacetate **4** was carried out in DMF. Later, $CH_3CN$ was employed as the media to access a readily workup and purification, the corresponding yield was 92%, which was comparable with product synthesised when DMF was used.

Subsequently, selective hydrolysis of the anomeric thioacetate and $S_N2$ reaction were set up in one-pot, giving the cyano-functionalized lactose **5** in 94% yield; following hydrolysis of acetates in hot alkaline methanol generated **6** in 72% yield, which served the substrate for the enzymatic reaction.

*Expression and Purification of NmCSS/PmST3*

Enzymes associated must be expressed prior to enzymatic sialylation. Plasmids pET23a-*NmCSS* (sialic acid-CMP synthetase from *Neisseria meningitidis*) and pET23a-*PmST3* (2,3-sialyl-CMP transferase from *Pasteurella multocida*) were transferred into XL10-gold competent cells. After culture and extraction, plasmids were sequence-validated.

Enzymes were expressed in BL21(DE3) competent *E.* coli. Expression was confirmed by SDS-PAGE and Western Blotting, as shown in **Figure I-2**.

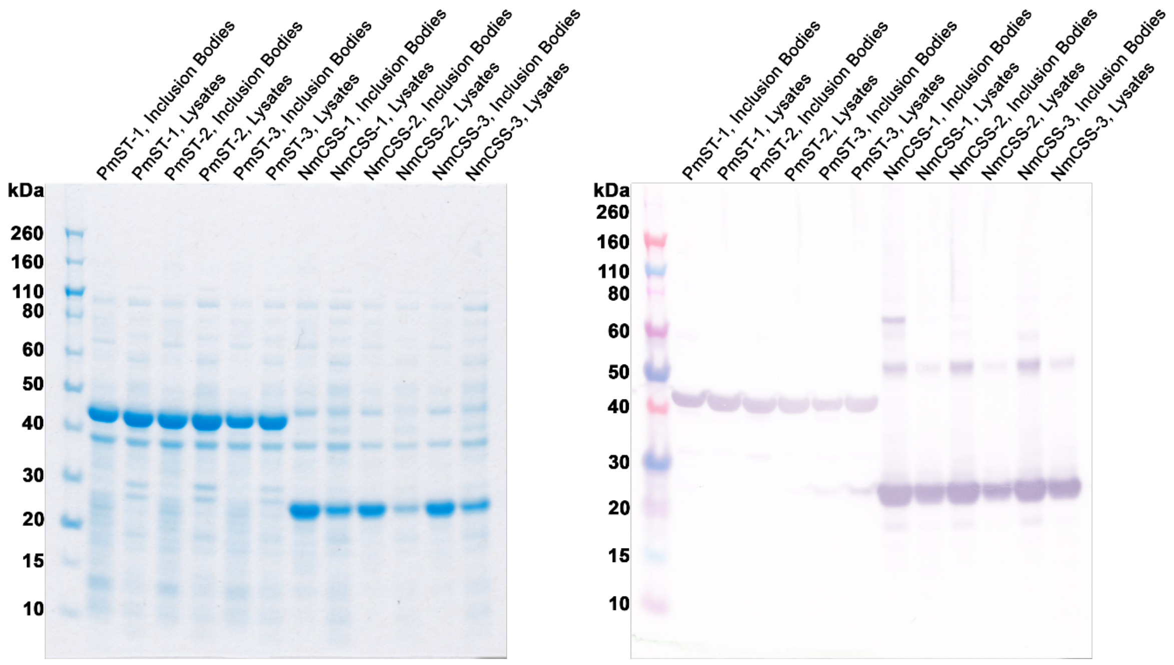


**Figure I-2.** SDS-PAGE and western blotting analysis of expressed *NmCSS* and *PmST3*.

Purification was then carried out manually by using Ni-resin column followed by desalting. Concentrations were analysed by BCA assay, giving enzymes *NmCSS* (11.81 mg/mL, 3.5 mL) and *PmST3* (14.47 mg/mL, 3.5 mL) from 1 L of culture, separately. Enzymes were then characterized by mass spectrometry (**Figure I-3**), aliquoted, and stored at – 78 °C for use

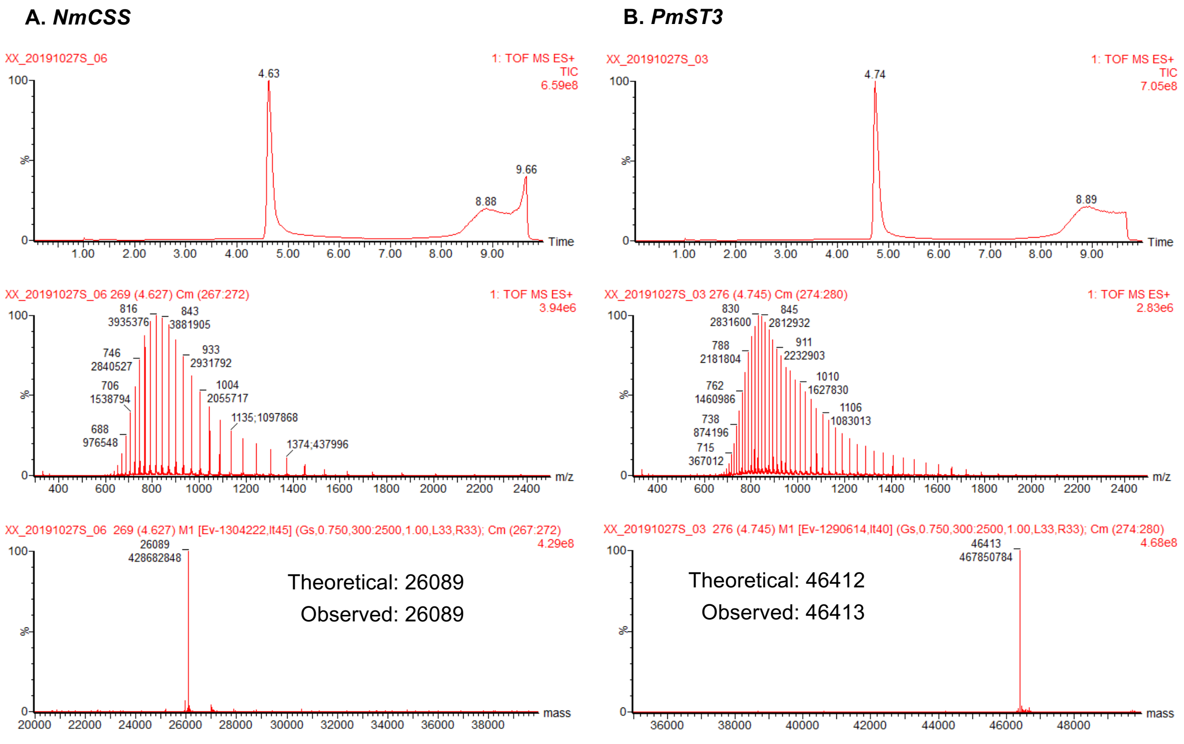


**Figure I-3.** LC_MS of purified *NmCSS* and *PmST3*.

*Enzymatic assembly of cyanomethyl 3'-SiaLac* ***1***

Assembly of 2,3-linked sialic glycoside has been investigated (*1–5*). Efficient 2,3-sialylation was achieved under varying conditions, but the yield was highly substrate-dependent. In the analysis shown in **Figure I-4**, the reaction condition was optimized, monitoring the outcome by TLC.

First, sialylation was tested in different buffers which have been reported previously (*1–5*). As shown in **Figure I-4A**, compared with the Tris·HCl buffer, ammonium carbonate (pH = 8.5) delivered a more efficient conversion. Also, we found that this conversion was related to the pH value of the reaction since the activities of enzymes were pH-dependent (*6, 7*). Furthermore, increasing the reaction time and the addition of more CTP did not improve yield.

As reported in the literature (*8*), *PmST3* may hydrolyse sialic acid-CMP (or CMP-Neu5Ac), the donor of sialylation, when too much enzyme was incubated (the mechanism of sialylation was shown in **Scheme 1-2**). The reaction was titrated with different amounts of *NmCSS* and *PmST3*. Notably, where the enzyme concentrations were lowered, there was no improvement other than a slower transformation (**Figure I-4B**).

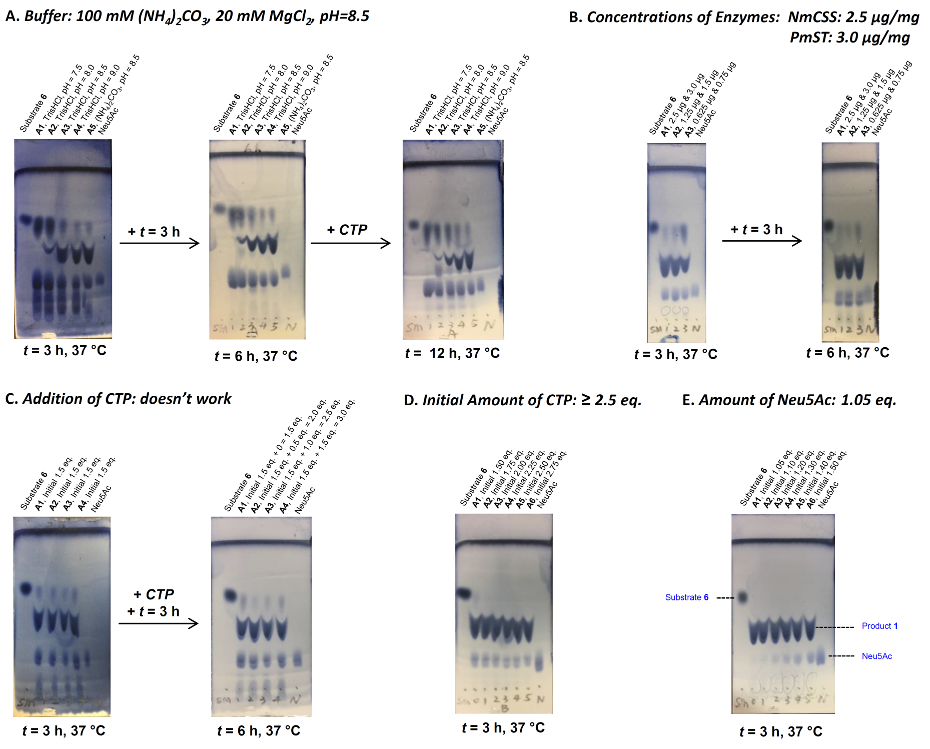


**Figure I-4.** Optimization of enzymatic condition (only monitored by TLC). TLCs were developed in $H_2O$–*i*PrOH–EtOAc (1:2:4) system followed by visualization in cerium ammonium molybdate (CAM) stain; description of spots was marked in **E**.

Sialic acid-CMP could be hydrolysed to sialic acid and CMP in alkaline reaction buffer, which was essential for maintaining activities of enzymes. This resulted a consumption of CTP. Therefore, additional CTP could contribute to a better yield. As we expected, a higher conversion was observed when more CTP was appended. While the substrate **6** couldn't be consumed even 3.0 eq. (1.5 eq. plus 1.5 eq.) of CTP was entirely used (**Figure I-4C**).

Later, reaction was carried out by using vary amounts of initial CTP. Complete sialylation was obtained when the initial amount of CTP was over 2.5 eq. (**Figure I-4D**), this was different from when CTP was added sequentially (**Figure I-4C**).

Although sialic acid-CMP was hydrolysed in reaction buffer, the resulting free sialic acid could be continuously recycled when CTP content was sufficient (over 2.5 eq.). Less sialic acid could make subsequent purification much easier. As shown in **Figure I-4E**, 1.05 eq. of sialic acid gave complete conversion, based on the TLC.

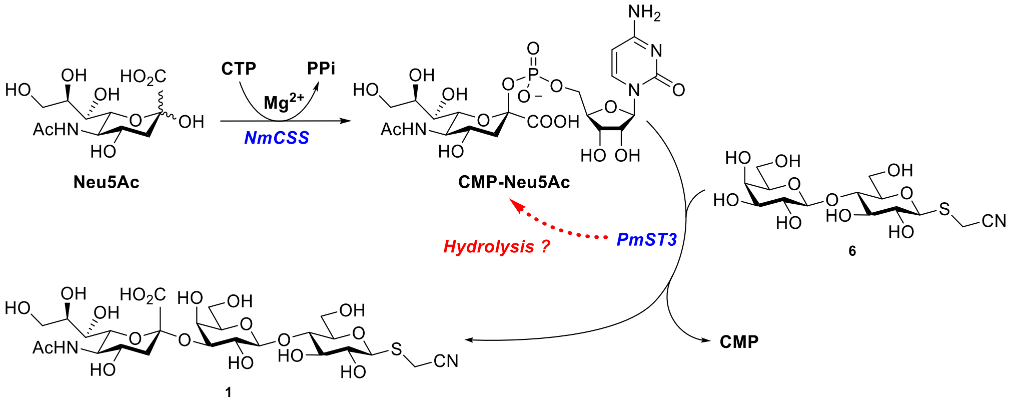


**Scheme I-2.** Mechanism of 2,3-silylation in a one-pot two-enzyme system.

Having optimised the reaction conditions, 2,3-sialylation was scaled up to 200 mg. A flash column chromatography (silica gel) followed by a size exclusive chromatography (LH20) revealed the desired trisaccharide **1** with a 96% yield.

The formed sialosidic linkage was characterized by NMR. A significant downfield-movement of H-3′ in proton spectrum (3.69 *ppm* to 4.14 *ppm*) and a strong correlation between H-3′ and C-2″ in HMBC spectrum verified the 2,3-linkage. Also, the coupling constant between C-1″ and H-3″$_{ax}$ ($^3J$ = 4.7 Hz) indicated the expected α-sialoside (*9–11*).

*Glycan reagent activation*

Treatment of a catalytic amount of $CH_3ONa$ in methanol, cyano-functionalized trisaccharide **1** can be converted into the imidate-linked 3′-SiaLac **7**, as shown in **Scheme I-3**. This product can react with free primary amines of lysine residues on peptides and proteins.

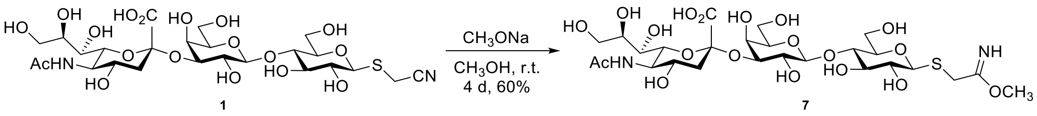


**Scheme I-3.** Equilibrium between cyano-functionalized 3′-SiaLac **1** and imidate-linked 3′-SiaLac **7**.

Considering previously reported conditions in the literature (*12–15*), activation of trisaccharide **1** was performed in dry methanol-$d_4$ using 1.0 eq. of $CD_3ONa$. The real-time conversion was monitored by $^1H$ NMR spectrum. As shown in **Figure I-5A**, in the anhydrous conditions ($CD_3OD$ was dried over 4Å MS), the cyano group was mildly converted into the corresponding imidate. The maximum conversion, 60%, was reached after 3–4 days at room temperature (**Figure I-6**). Continuous observations illustrated that the formed imidate was quite stable (up to 10 days) in the reaction solution.

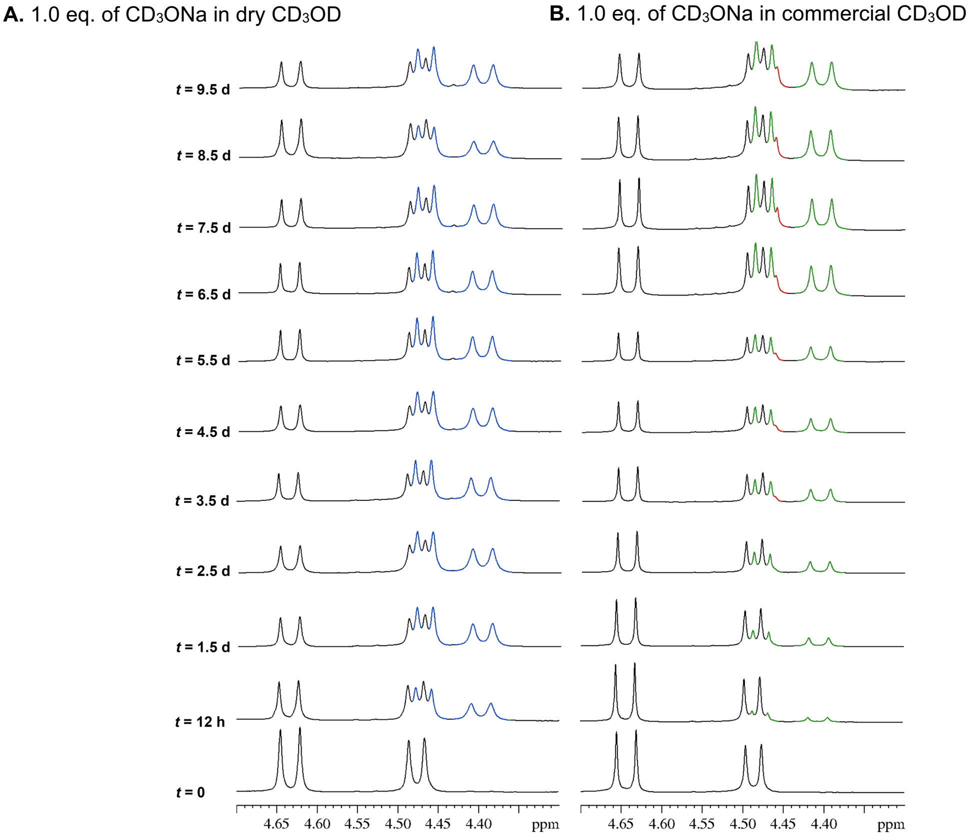


**Figure I-5.** Real-time $^1H$ NMR spectra of activation of **1**. Black peaks showed anomeric protons (H-1 and H-1′) of **1**; blue ones (**A**) or green ones (**B**) represented anomeric protons (H-1 and H-1′) of **7**; red signals in **B** indicated byproduct formation which probably was from hydrolysis of imidate **7**.

When the activation was directly tested in the commercially available $CD_3OD$, a much slower conversion was obtained. It took around one week to get 50% yield as the maximum conversion, as shown in **Figure I-6**. Moreover, according to $^{1}H$ NMR spectra illustrated in **Figure I-5B**, additional peaks, highlighted as red, were around which indicated the formation of a byproduct.

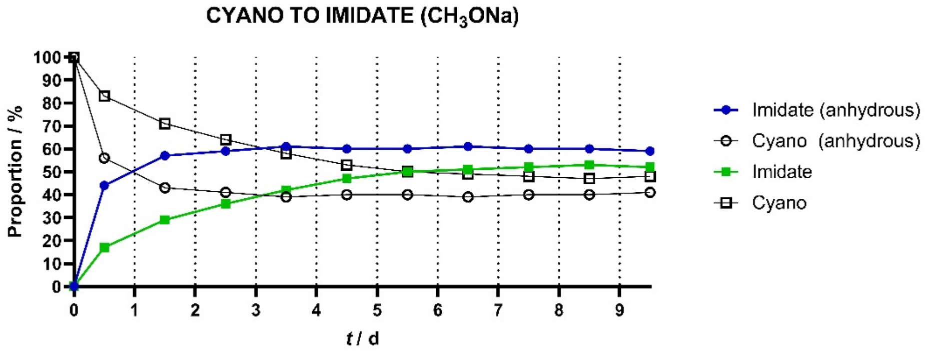


**Figure I-6.** Conversion from cyano **1** to imidate **7** in $CD_3ONa$ solution.

Activation of glycan **1** was tested under various "forcing" conditions (*14*), including reflux with $CH_3ONa$ or *t*BuOK in methanol, however, for example, the conversion was much worse, with the final yield of imidate **7** at around 20%.

*Detailed parameters affecting glycan reagent activation*

Although these methods allow ready activation prior to LOG formation, to gain additional insight on the influence of experimental parameters on the conversion of cyano-functionalized $Le^x$ into imidate-linked $Le^x$ (**$Le^x$-IME**), we also employed a response surface methodology (RSM). RSM is a statistical technique based on design of experiments (DoE). For our study, we used the Doehlert experimental design, which necessitates conducting $N = k^2 + k + N_0$ experiments, where $k$ represents the number of the parameters and $N_0$ represents the number of center runs. Our investigation comprised 25 experiments with 5 conducted at the center of the domain.

*O*-Acetylated (alcohol function protection) cyano-functionalized $Le^x$ (50 mg, 54 µmol) was suspended in dry $CH_3OH$ (final concentration *X4*) to which was added $CH_3ONa$ (*X2* equiv.). The reaction was left stirring at *X3* °C for *X1* h. The reaction mixture was analysed by $^{1}H$ NMR in $CH_3OH$ solvent (annihilation of solvent signal) to avoid any kinetic isotope effect resulting from deuterated methanol. The final ratio of the activated (unprotected) **$Le^x$-IME** on the cyano-functionalized (unprotected) $Le^x$ was used as output of the system.

**A.**

| Exp. | X1 | X2 | X3 | X4 |
|---|---|---|---|---|
| 1 | 1 | 0 | 0 | 0 |
| 2 | -1 | 0 | 0 | 0 |
| 3 | 0,5 | 0,866 | 0 | 0 |
| 4 | -0,5 | -0,866 | 0 | 0 |
| 5 | 0,5 | -0,866 | 0 | 0 |
| 6 | -0,5 | 0,866 | 0 | 0 |
| 7 | 0,5 | 0,2887 | 0,8165 | 0 |
| 8 | -0,5 | -0,2887 | -0,8165 | 0 |
| 9 | 0,5 | -0,2887 | -0,8165 | 0 |
| 10 | 0 | 0,5774 | -0,8165 | 0 |
| 11 | -0,5 | 0,2887 | 0,8165 | 0 |
| 12 | 0 | -0,5774 | 0,8165 | 0 |
| 13 | 0,5 | 0,2887 | 0,2041 | 0,7906 |
| 14 | -0,5 | -0,2887 | -0,2041 | -0,7906 |
| 15 | 0,5 | -0,2887 | -0,2041 | -0,7906 |
| 16 | 0 | 0,5774 | -0,2041 | -0,7906 |
| 17 | 0 | 0 | 0,6124 | -0,7906 |
| 18 | -0,5 | 0,2887 | 0,2041 | 0,7906 |
| 19 | 0 | -0,5774 | 0,2041 | 0,7906 |
| 20 | 0 | 0 | -0,6124 | 0,7906 |
| 21 | 0 | 0 | 0 | 0 |
| 22 | 0 | 0 | 0 | 0 |
| 23 | 0 | 0 | 0 | 0 |
| 24 | 0 | 0 | 0 | 0 |
| 25 | 0 | 0 | 0 | 0 |
| Levels | 5 | 7 | 7 | 3 |

**B.**

| Parameter | Code | Level (-1) | Level (0) | Level (1) |
|---|---|---|---|---|
| Reaction time (h) | X1 | 4 | 62 | 120 |
| Equiv $CH_3ONa$ | X2 | 0,1 | 3,05 | 6 |
| Reaction Temperature (°C) | X3 | 4 | 27 | 50 |
| $Le^x$ concentration (mM) | X4 | 30 | 80 | 130 |

**C.**

| Experiment | 1 | 2 | 3 | 4 | 5 | 6 | 7 | 8 |
|---|---|---|---|---|---|---|---|---|
| X1 (h) | 120 | 4 | 91 | 33 | 91 | 33 | 91 | 33 |
| X2 (equiv.) | 3.05 | 3.05 | 5.61 | 0.50 | 0.50 | 5.61 | 3.90 | 2.20 |
| X3 (°C) | 27 | 27 | 27 | 27 | 27 | 27 | 46 | 8 |
| X4 (mM) | 80.0 | 80.0 | 80.0 | 80.0 | 80.0 | 80.0 | 80.0 | 80.0 |
| IME/cyano ratio | 0.95 | 1.07 | 1.07 | 0.93 | 0,97 | 1.03 | 0.57 | 1.58 |
| **Experiment** | **9** | **10** | **11** | **12** | **13** | **14** | **15** | **16** |
| X1 (h) | 91 | 62 | 33 | 62 | 91 | 33 | 91 | 62 |
| X2 (equiv.) | 2.20 | 4.75 | 3.90 | 1.35 | 3.90 | 2.20 | 2.20 | 4.75 |
| X3 (°C) | 8 | 8 | 46 | 46 | 32 | 22 | 22 | 22 |
| X4 (mM) | 80.0 | 80.0 | 80.0 | 80.0 | 119.5 | 40.5 | 40.5 | 40.5 |
| IME/cyano ratio | 1.93 | 1.6 | 0.67 | 0.57 | 0.88 | 1.23 | 1.25 | 1.28 |
| **Experiment** | **17** | **18** | **19** | **20** | | | | **21-22-23-24-25** |
| X1 (h) | 62 | 33 | 62 | 62 | | | | 62 |
| X2 (equiv.) | 3.05 | 3.90 | 1.35 | 3.05 | | | | 3.05 |
| X3 (°C) | 41 | 32 | 32 | 13 | | | | 27 |
| X4 (mM) | 40.5 | 119.5 | 119.5 | 119.5 | | | | 80.0 |
| IME/cyano ratio | 0.71 | 0.83 | 0.74 | 1.77 | | | | 1.01 |

The outcomes of this study resulted in the development of a specific guiding model for **$Le^x$-IME** activation, as an example:

$$Y = 1.004 + 0.016\, b_1 + 0.040\, b_2 - 0.722\, b_3 - 0.040\, b_4 + 0.224\, b_{33} + 0.087\, b_{44} - 0.275\, b_{13} + 0.141\, b_{23} + 0.091\, b_{14} - 0.202\, b_{34}$$

, a second-order polynomial equation obtained through the Doehlert experimental design where Y represents the imidate/cyano molar ratio ($^1$H NMR). The coefficients $b_x$ and $b_{xx}$ are first and second-order terms respectively (related to each parameter *X1*, *X2*, *X3* and *X4*), and the coefficients $b_{xy}$ are the first-order terms related to the interactions between two parameters. $b_1$: reaction time, $b_2$: equivalence of $CH_3ONa$, $b_3$: reaction temperature, $b_4$: $Le^x$ concentration. Statistical parameters of the model: $R^2$: 0.971, $R^2$A: 0.930 (NemrodW, version 9901). The model gives IME/cyano ratio based on selected parameters. [To utilize this model, the coefficients $b_1$, $b_2$, $b_3$ and $b_4$ need to be replaced with the coded values (ranging from -1 to 1) corresponding to the selected parameters (*X1*, *X2*, *X3* and *X4*). For example, $b_1$=1 if the reaction time (*X1*) is set to 120 h. The coefficients $b_{33}$ and $b_{44}$ represent the second-order terms of the coded values (for instance $b_{11}=1^2$ if the reaction time *X1* is set to 120 h). The interference coefficients $b_{13}$, $b_{23}$, $b_{14}$, and $b_{34}$, need to be replaced with the product of the corresponding coded values (for instance $b_{13}=b_1*b_3=1*(-1)$ if the reaction time *X1* is set to 120 h and the temperature *X3* is set to 4 °C. This model also allows us to determine the optimal conditions for the reaction, which are *X1*=120 h, *X2*=0.1 equiv., *X3*=4 °C and *X4*=130 mM. Under these conditions, the imidate/cyano ratio is 2.68, corresponding to 72.8 %mol of activated imidate.

Whilst all effects are small, reaction temperature has the most significant influence on the outcome, and several pairs of parameters strongly interact with each other, such as temperature/time, temperature/Le$^x$ concentration, temperature/equivalence of $CH_3ONa$ and reaction time/Le$^x$ concentration, in descending order of influence. It is important to note that this model equation is validated within the studied domain but highlights the general principle of how activation may be yet further optimized if needed.

*LOG formation*

3′-SiaLac modified HELs (HEL-SiaLac$_x$ ≣ HEL-[–amidine-GM3g]$_x$) can be achieved when carried out in PBS buffer (pH = 7.4). In a recent report using similar chemistries, sodium borate (SB) buffer (pH = 8.5) was introduced as well (*15*). Therefore, generation of HEL-[–amidine-GM3g]$_x$ was tested in PBS buffer or SB buffer by using the alkaline (crude) or neutral **7** (which was neutralized by resin (hydrogen form) after activation).

As illustrated in **Figure I-7**, the modifications with crude sugar **7** in PBS (**Figure I-7B & C**) was faster and more efficient than that in SB buffer; coupling was more processed in SB buffer, giving the over glycosylated conjugate HEL-SiaLac$_7$ for 5% yield. When the reaction mixture was neutralized by DOWEX 50WX8 (100~200 mesh, hydrogen form) after the formation of imidate **7**, it failed in PBS buffer (**Figure I-7C**), while coupling in SB buffer (**Figure I-7D**) worked but was much slower in comparison to that when crude **7** was used (**Figure I-7B**).

**A.**

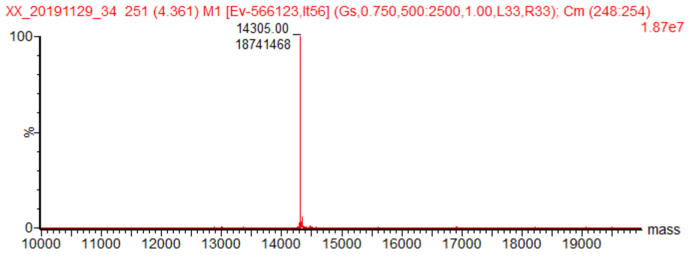


| | | |
|---|---|---|
| HEL | MW: 14305 | |
| HEL-SiaLac$_1$ | MW: 14993 | (P1) |
| HEL-SiaLac$_2$ | MW: 15681 | (P2) |
| HEL-SiaLac$_3$ | MW: 16369 | (P3) |
| HEL-SiaLac$_4$ | MW: 17058 | (P4) |
| HEL-SiaLac$_5$ | MW: 17746 | (P5) |
| HEL-SiaLac$_6$ | MW: 18434 | (P6) |
| HEL-SiaLac$_7$ | MW: 19122 | (P7) |

**B.**

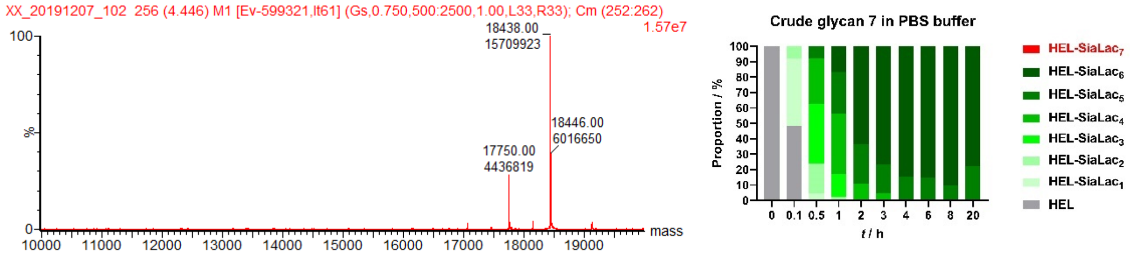


**C.**

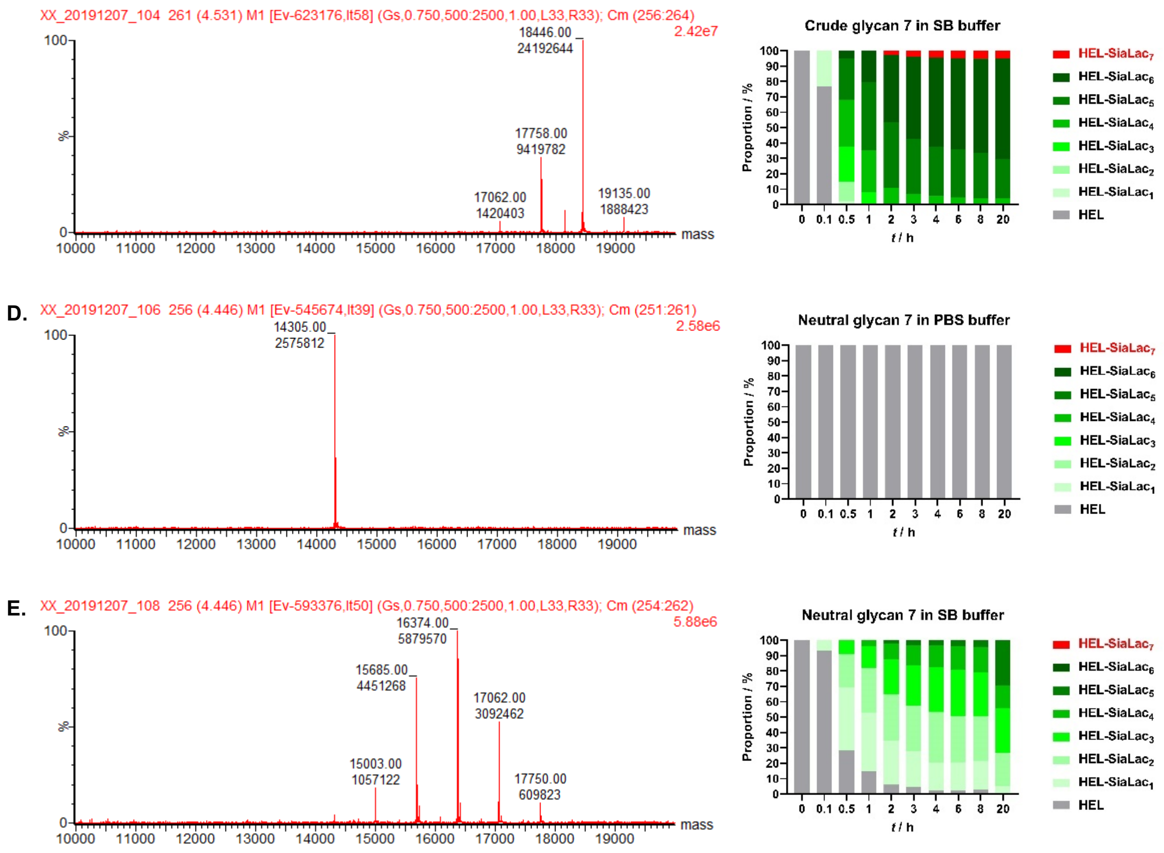


**Figure I-7.** Kinetic analysis and MS spectra at the final point. 60 Eq./Lys of imidate **7** were added and six lysine residues were considered for all cases. A full list of MS data was attached in section 7.2.

According to these preliminary data, several key points were identified: 1) PBS buffer was favourable for modification; 2) resin-neutralized glycan **7** didn't work as expected or, at least, acidic resin was not suitable for neutralization since it could easily lead to over-neutralization which subsequently altered the pH of the reaction; 3) reaction rate and efficiency were pH-dependent; 4) the stability of formed glycoconjugates (HEL-SiaLac$_x$) should be evaluated. As shown in **Figure 1-7B**, HEL-SiaLac$_6$ appeared to be hydrolysed back to HEL-SiaLac$_5$ when the reaction time was extended from 8–20 h, and 5) 60 eq. of glycan **7** for each lysine was not enough to get a complete conversion, much more sugar was required to furnish the hexavalent conjugate HEL-SiaLac$_6$ as the sole product.

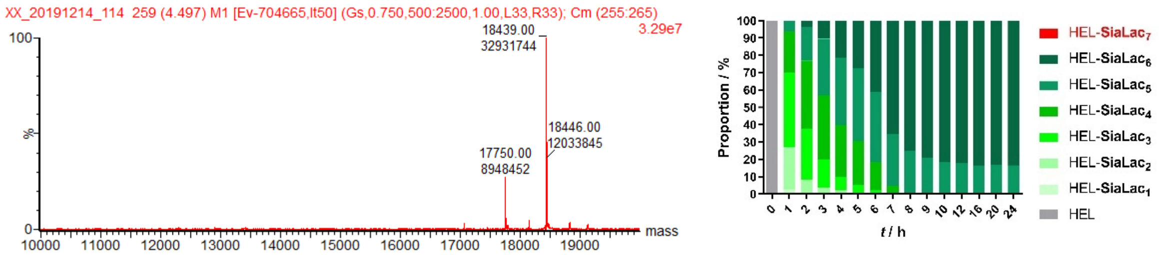


**Figure I-8.** Stability Investigation of HEL_SiaLac$_x$. 60 eq. of glycan **7** was used for one lysine and six lysine residues were considered.

To figure out the stability of conjugates HEL-SiaLac$_6$, modification was tried again in PBS buffer by using 60 eq. of crude glycan **7** for each lysine residue. As illustrated in **Figure I-8**, coupling was done in 12 h, yielding conjugates HEL-SiaLac$_5$ (17%) and HEL-SiaLac$_6$ (83%) as a mixture. Also, the formed conjugates were stable in this reaction solution for at least 24 h at room temperature.

Since the crude imidate **7** is more effective for HEL modification, it will be employed for exploring the reaction condition tentatively. Crude sugar **7** derived from direct concentration was submitted into PBS buffer (pH = 7.4). The resulting sugar solution was then mixed with HEL. The EP tube was incubated at 25 °C and the modification progress was monitored by LC_MS after desalting. As shown in **Figure I-9**, the coupling rate and progress increased as more sugar was used, the actual pH value of the reaction had a certain correlation with the sugar amount: the amount of sugar and associated pH value determine the reaction rate and the final conversion.

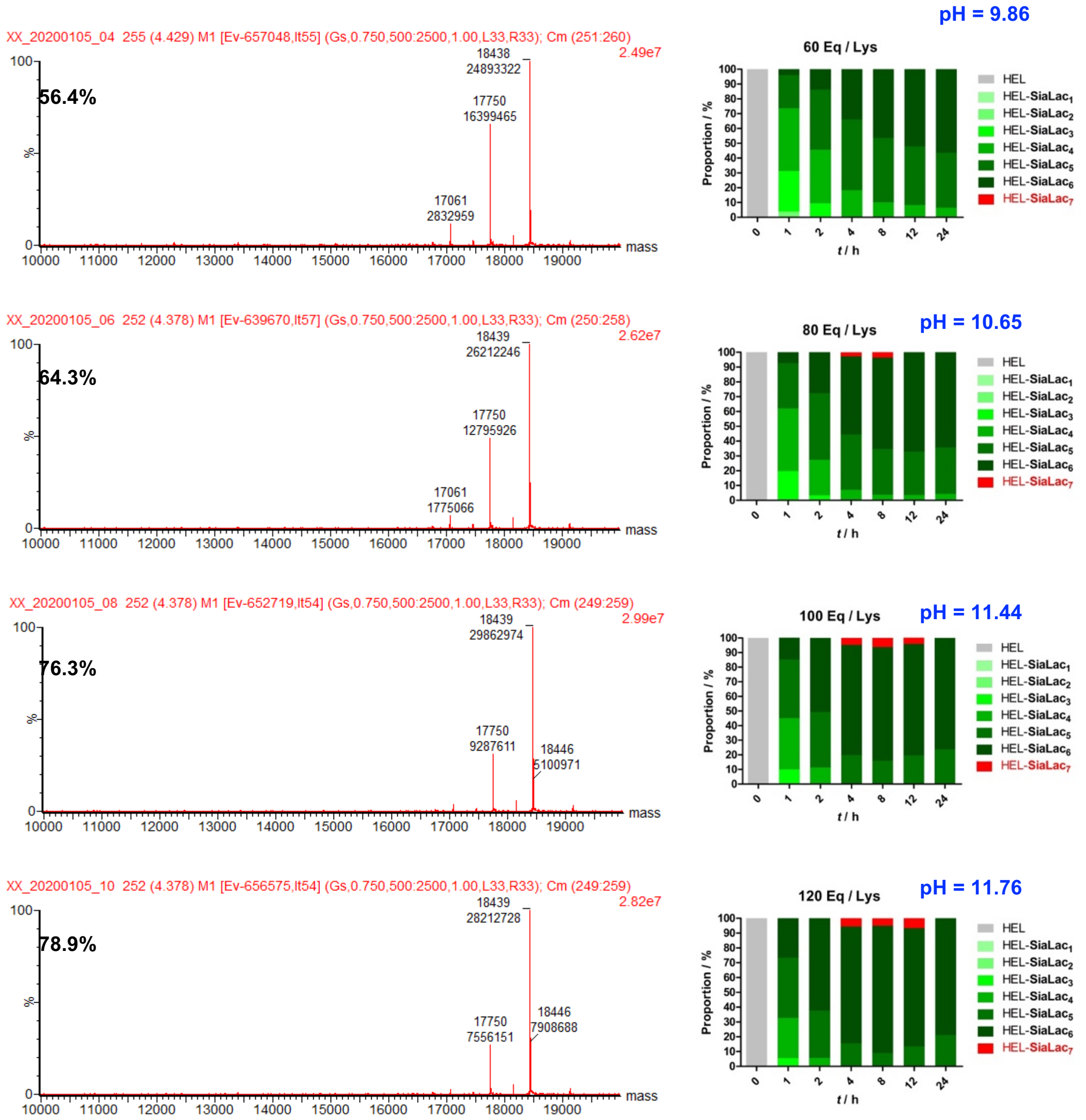




pH = 11.89

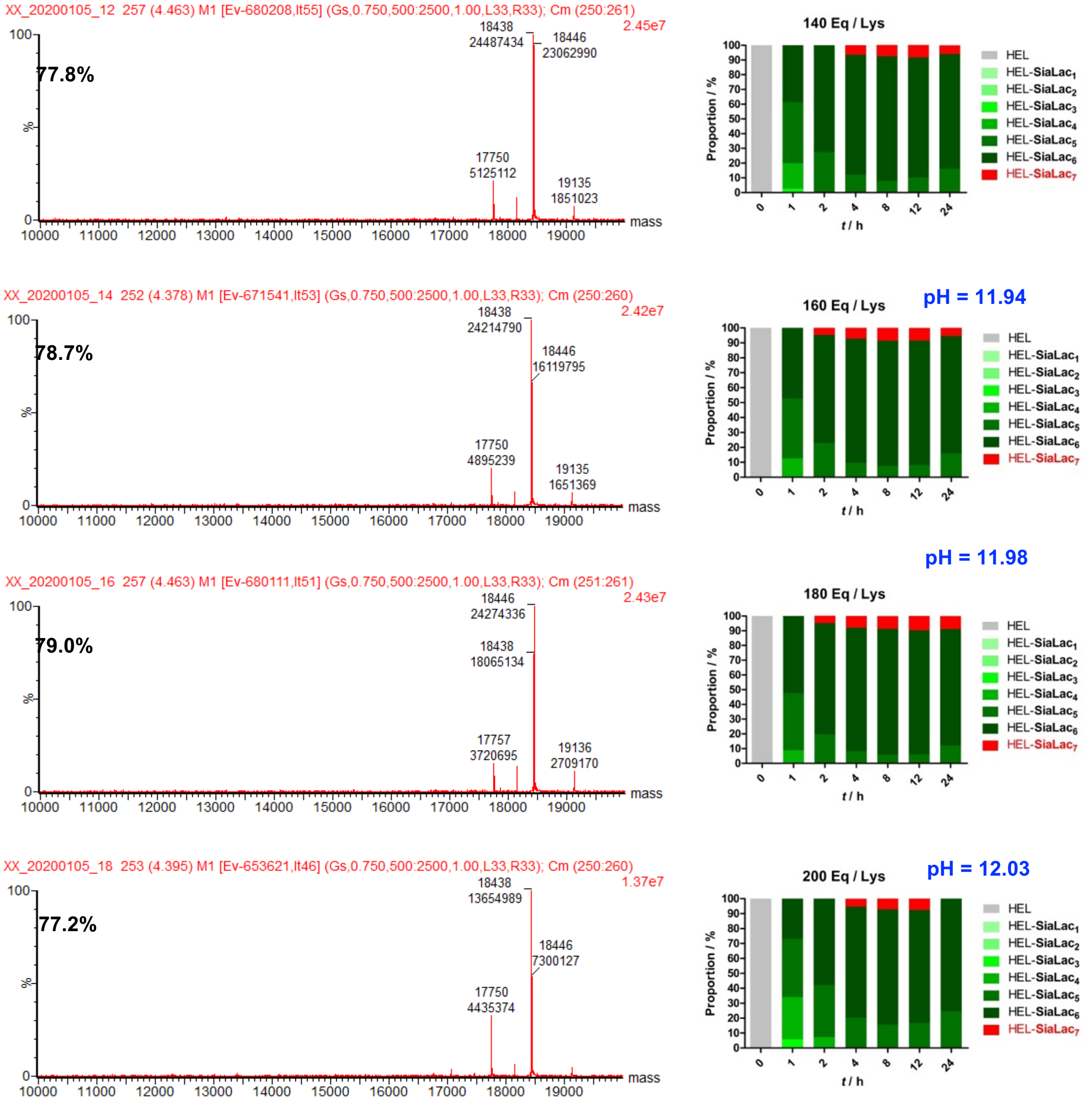


**Figure I-9.** HEL modification by using the stale, crude sugar **7**. The actual pH value of mixture and the corresponding yield of HEL_SiaLac$_6$ were noted. Illustrated LC_MS was the one when t = 24 h.

Here, the yield of HEL-SiaLac$_6$ was poor. A potential reason was that the sugar was not freshly prepared. Imidate **7** used in the reaction was stored at -78 °C for two weeks after concentration. This suggests that decomposition may have occurred during storage.

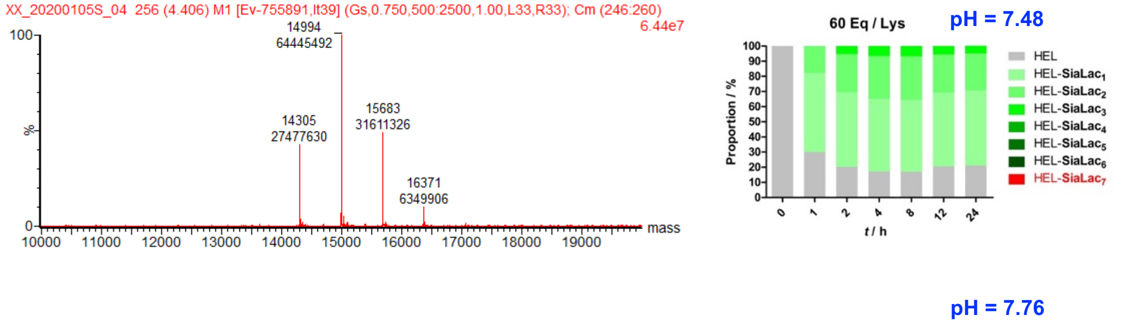

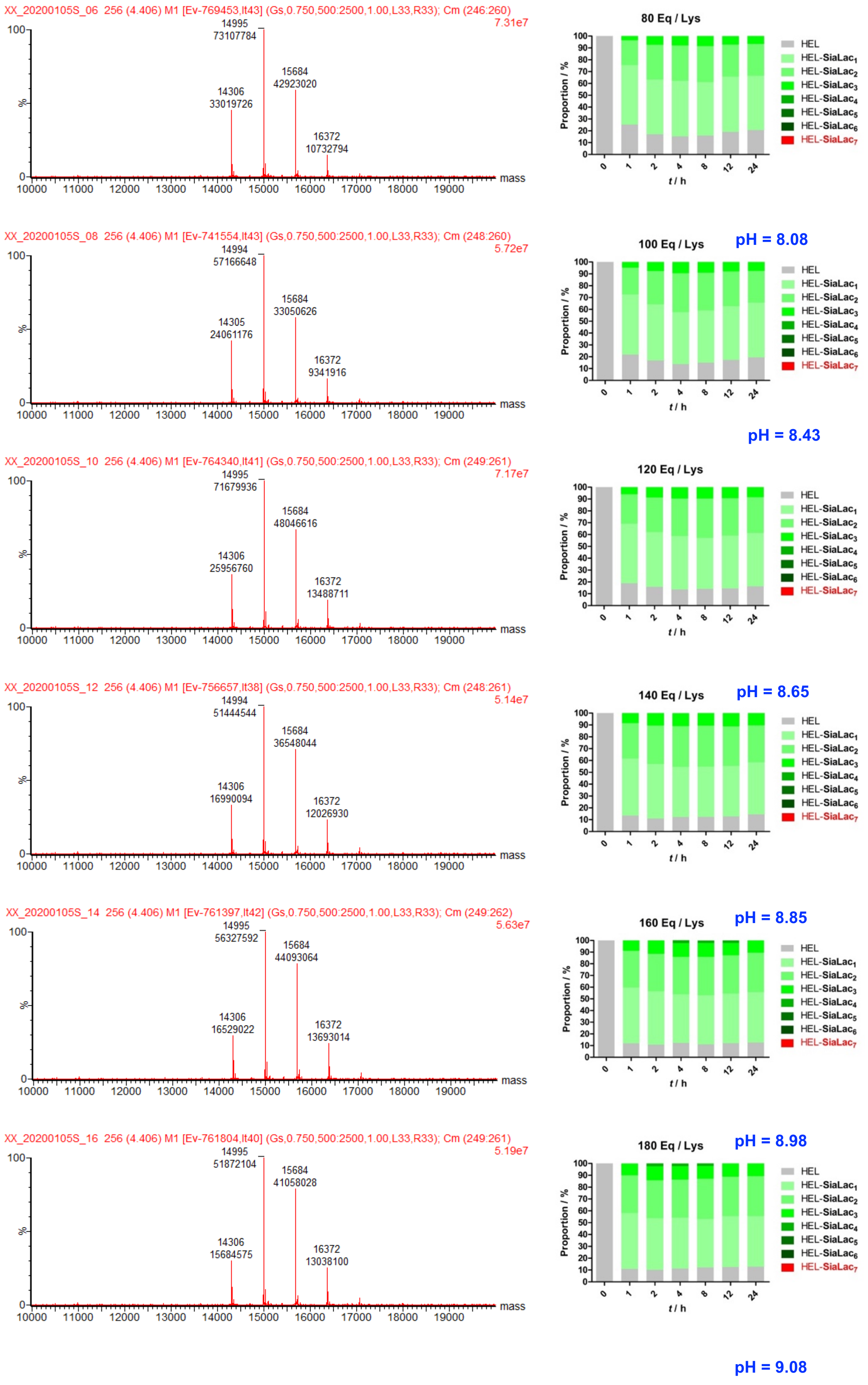

XX_20200105S_06 256 (4.406) M1 [Ev-769453,It43] (Gs,0.750,500:2500,1.00,L33,R33); Cm (246:260)
7.31e7
14995
73107784
15684
42923020
14306
33019726
16372
10732794
80 Eq / Lys
XX_20200105S_08 256 (4.406) M1 [Ev-741554,It43] (Gs,0.750,500:2500,1.00,L33,R33); Cm (248:260)
5.72e7
14994
57166648
15684
33050626
14305
24061176
16372
9341916
100 Eq / Lys
pH = 8.08
pH = 8.43
XX_20200105S_10 256 (4.406) M1 [Ev-764340,It41] (Gs,0.750,500:2500,1.00,L33,R33); Cm (249:261)
7.17e7
14995
71679936
15684
48046616
14306
25956760
16372
13488711
120 Eq / Lys
XX_20200105S_12 256 (4.406) M1 [Ev-756657,It38] (Gs,0.750,500:2500,1.00,L33,R33); Cm (248:261)
5.14e7
14994
51444544
15684
36548044
14306
16990094
16372
12026930
140 Eq / Lys
pH = 8.65
XX_20200105S_14 256 (4.406) M1 [Ev-761397,It42] (Gs,0.750,500:2500,1.00,L33,R33); Cm (249:262)
5.63e7
14995
56327592
15684
44093064
14306
16529022
16372
13693014
160 Eq / Lys
pH = 8.85
XX_20200105S_16 256 (4.406) M1 [Ev-761804,It40] (Gs,0.750,500:2500,1.00,L33,R33); Cm (249:261)
5.19e7
14995
51872104
15684
41058028
14306
15684575
16372
13038100
180 Eq / Lys
pH = 8.98
pH = 9.08
100
%
0
mass
10000 11000 12000 13000 14000 15000 16000 17000 18000 19000
Proportion / %
t / h
0 1 2 4 8 12 24
HEL
HEL-SiaLac1
HEL-SiaLac2
HEL-SiaLac3
HEL-SiaLac4
HEL-SiaLac5
HEL-SiaLac6
HEL-SiaLac7

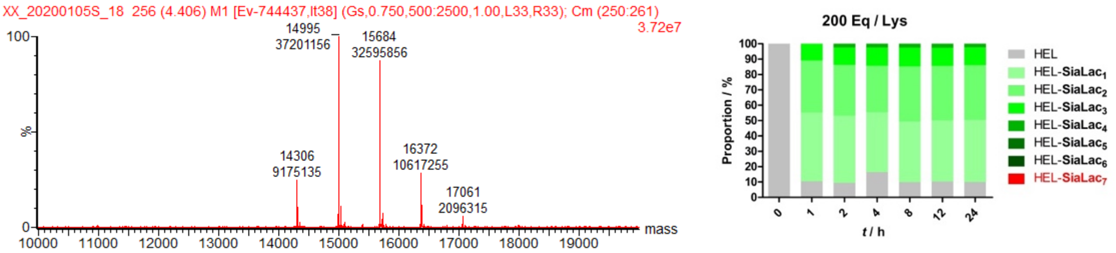


**Figure I-10.** HEL modification by using the stale, neutralized sugar **7**. The actual pH value of mixture and the corresponding yield of HEL_SiaLac6 were noted. Illustrated LC_MS was the one when t = 24 h.

When the old, neutralized sugar was used for HEL modification, the reaction became slow. the pH value of the mixture was slightly altered by the neutral sugar ($CH_3Ona$ was neutralized with an equivalent amount of acetic acid) (**Figure I-10**).

For comparison, fresh sugar **7** was neutralized by acetic acid again. The resulting sugar was used for modification immediately. In this case, the pH value of the reaction was the same as the one when the old, neutral sugar was introduced (**Figure I-10**). As illustrated in **Figure I-11**, the process was very slow, only giving a mixture of glycoconjugates with slight modification. If the pH value of reaction was forced to back to the initial pH of PBS buffer (pH = 7.4), we observed the effect of sugar amount on modification. As shown in **Figure I-12**, progress was limited for all batches; sugar amount and reaction time was largely irrelevant, which verified that the pH value is the primary determinant of the modification outcome.

pH = 7.48

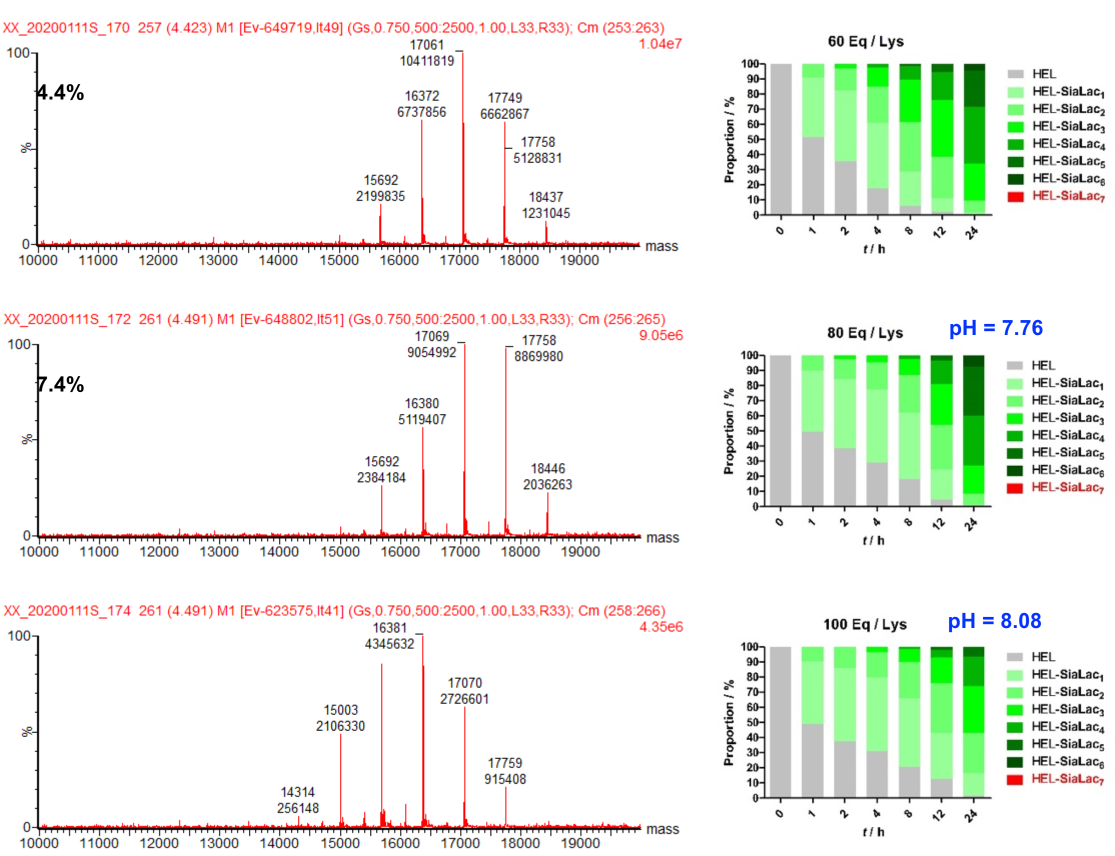




pH = 8.43

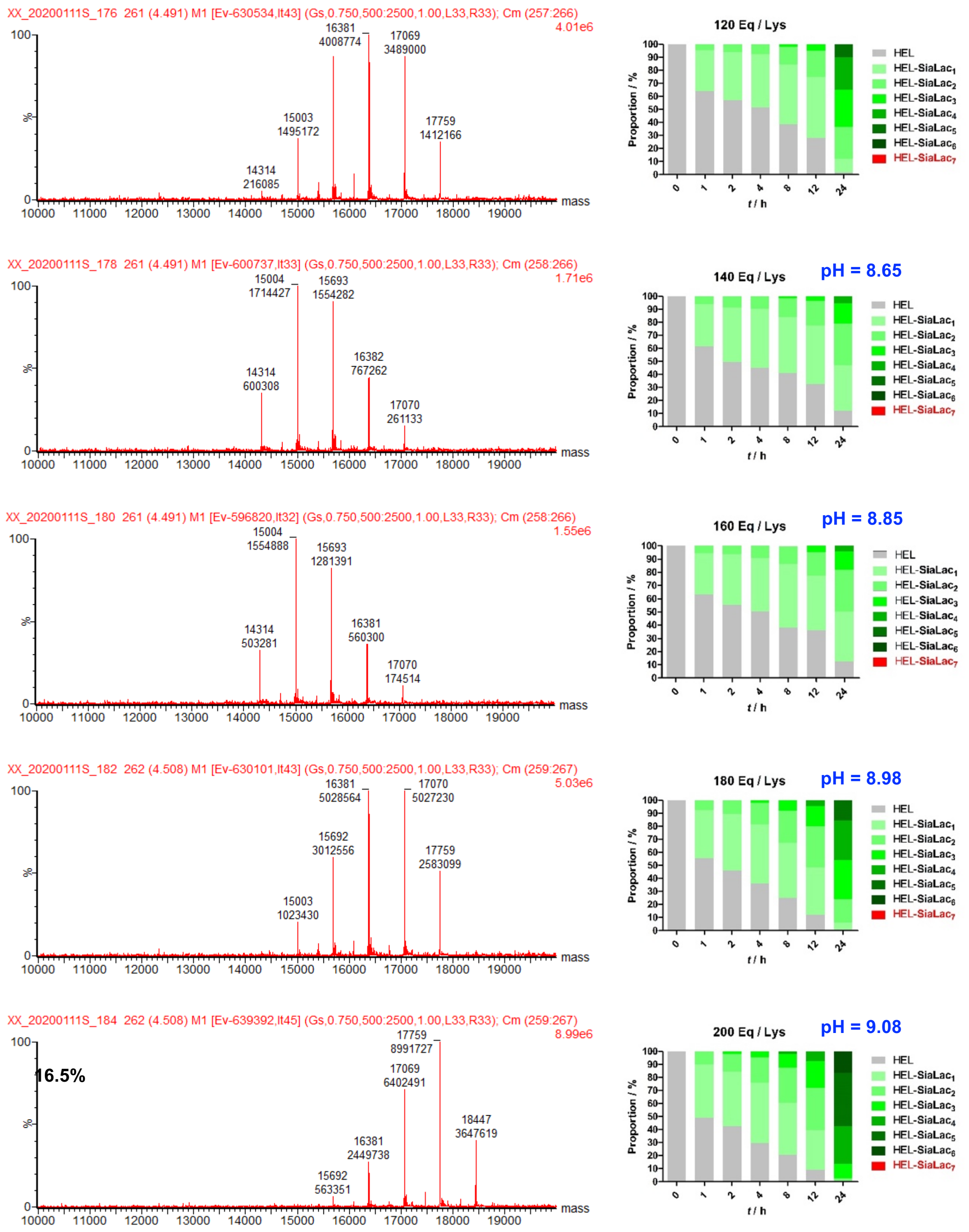


**Figure I-11.** HEL modification by using the freshly neutralized sugar **7**. The actual pH value of mixture and the corresponding yield of HEL_SiaLac6 were noted. Illustrated LC_MS was the one when t = 24 h.

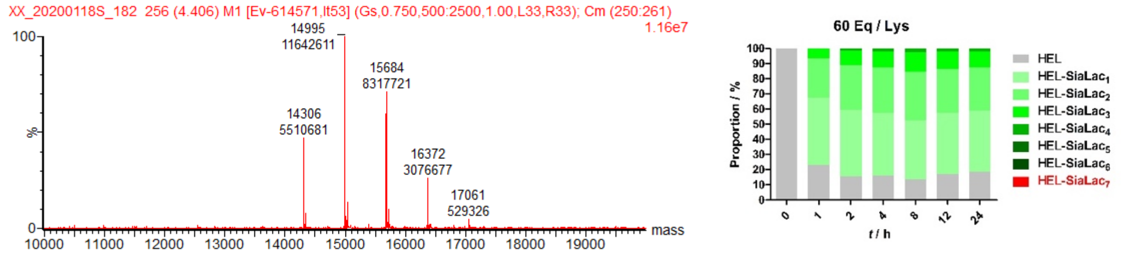

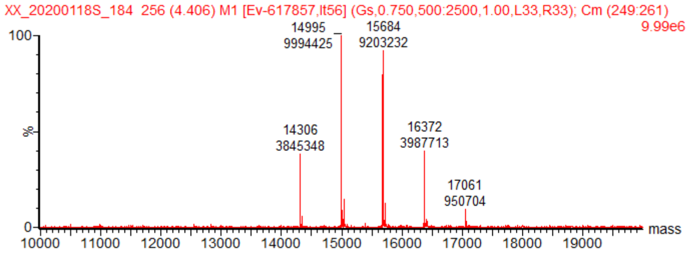
XX_20200118S_184 256 (4.406) M1 [Ev-617857,It56] (Gs,0.750,500:2500,1.00,L33,R33); Cm (249:261)
9.99e6
14995
9994425
15684
9203232
14306
3845348
16372
3987713
17061
950704
mass

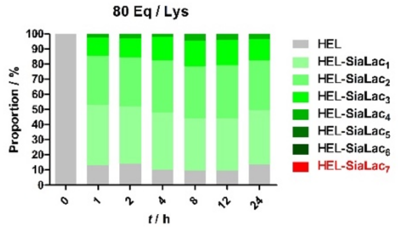
80 Eq / Lys
Proportion / %
t / h
HEL
HEL-SiaLac$_1$
HEL-SiaLac$_2$
HEL-SiaLac$_3$
HEL-SiaLac$_4$
HEL-SiaLac$_5$
HEL-SiaLac$_6$
HEL-SiaLac$_7$

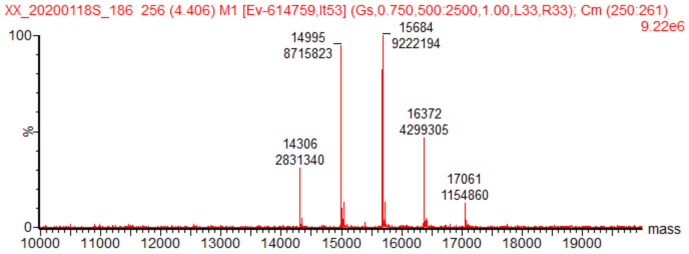
XX_20200118S_186 256 (4.406) M1 [Ev-614759,It53] (Gs,0.750,500:2500,1.00,L33,R33); Cm (250:261)
9.22e6
14995
8715823
15684
9222194
14306
2831340
16372
4299305
17061
1154860
mass

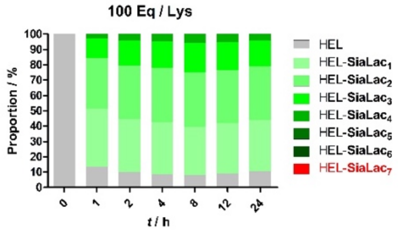
100 Eq / Lys
Proportion / %
t / h
HEL
HEL-SiaLac$_1$
HEL-SiaLac$_2$
HEL-SiaLac$_3$
HEL-SiaLac$_4$
HEL-SiaLac$_5$
HEL-SiaLac$_6$
HEL-SiaLac$_7$

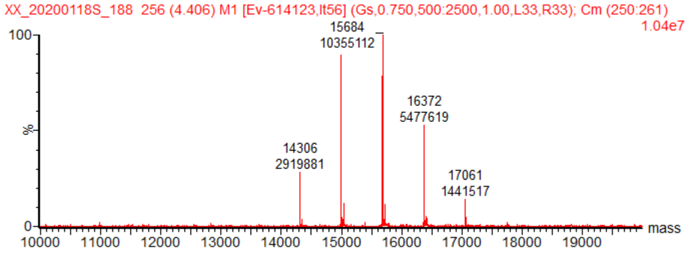
XX_20200118S_188 256 (4.406) M1 [Ev-614123,It56] (Gs,0.750,500:2500,1.00,L33,R33); Cm (250:261)
1.04e7
15684
10355112
14306
2919881
16372
5477619
17061
1441517
mass

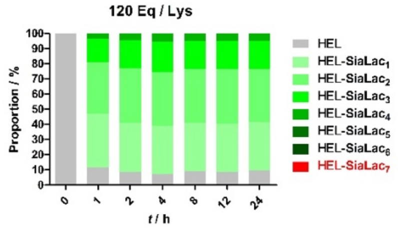
120 Eq / Lys
Proportion / %
t / h
HEL
HEL-SiaLac$_1$
HEL-SiaLac$_2$
HEL-SiaLac$_3$
HEL-SiaLac$_4$
HEL-SiaLac$_5$
HEL-SiaLac$_6$
HEL-SiaLac$_7$

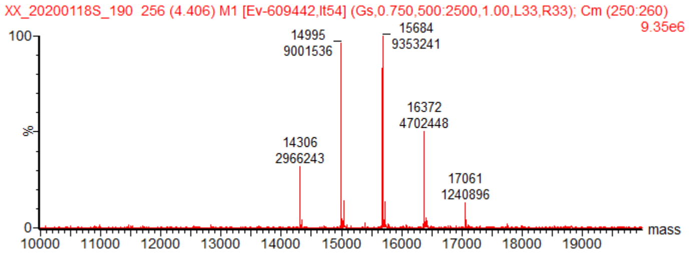
XX_20200118S_190 256 (4.406) M1 [Ev-609442,It54] (Gs,0.750,500:2500,1.00,L33,R33); Cm (250:260)
9.35e6
14995
9001536
15684
9353241
14306
2966243
16372
4702448
17061
1240896
mass

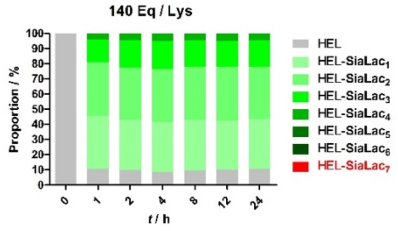
140 Eq / Lys
Proportion / %
t / h
HEL
HEL-SiaLac$_1$
HEL-SiaLac$_2$
HEL-SiaLac$_3$
HEL-SiaLac$_4$
HEL-SiaLac$_5$
HEL-SiaLac$_6$
HEL-SiaLac$_7$

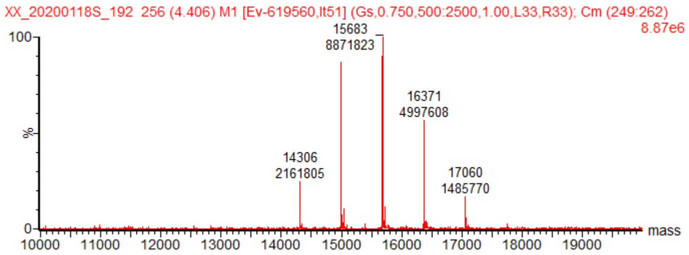
XX_20200118S_192 256 (4.406) M1 [Ev-619560,It51] (Gs,0.750,500:2500,1.00,L33,R33); Cm (249:262)
8.87e6
15683
8871823
14306
2161805
16371
4997608
17060
1485770
mass

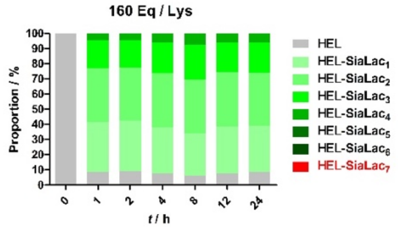
160 Eq / Lys
Proportion / %
t / h
HEL
HEL-SiaLac$_1$
HEL-SiaLac$_2$
HEL-SiaLac$_3$
HEL-SiaLac$_4$
HEL-SiaLac$_5$
HEL-SiaLac$_6$
HEL-SiaLac$_7$

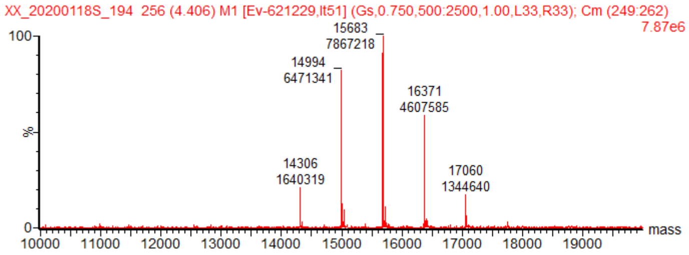
XX_20200118S_194 256 (4.406) M1 [Ev-621229,It51] (Gs,0.750,500:2500,1.00,L33,R33); Cm (249:262)
7.87e6
15683
7867218
14994
6471341
14306
1640319
16371
4607585
17060
1344640
mass

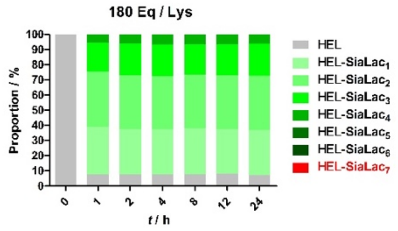
180 Eq / Lys
Proportion / %
t / h
HEL
HEL-SiaLac$_1$
HEL-SiaLac$_2$
HEL-SiaLac$_3$
HEL-SiaLac$_4$
HEL-SiaLac$_5$
HEL-SiaLac$_6$
HEL-SiaLac$_7$

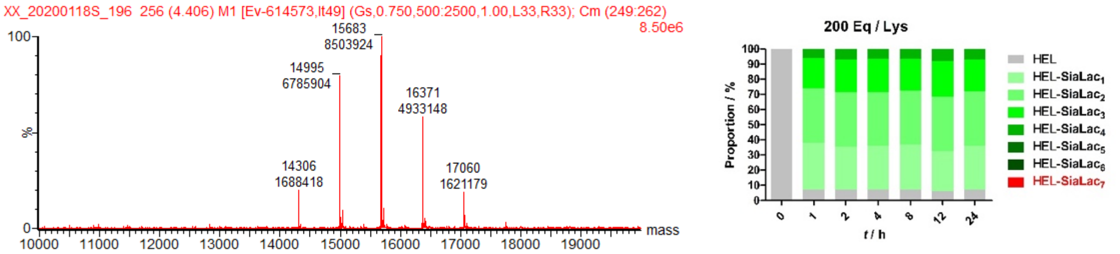


**Figure I-12.** HEL modification by using the freshly neutralized sugar **7** in PBS buffer (pH = 7.4). The pH was adjusted by HCl solution (1 M in MQ water). Illustrated LC_MS was the one when t = 24 h.

Subsequently, we tried to purify the sugar. Imidate **7** was slightly more polar than cyano **1**, but it is unstable in a silica column if an aqueous eluent was used. Another candidate was precipitation. Both cyano **1** and imidate **7** were very polar. The activation solution in methanol was slightly cloudy. Therefore, we speculated that sugar **7** could be precipitated by adding solvent with low polarity, while $CH_3ONa$ would be left in supernatant.

Once activation was done, an equal volume of THF was added. The white precipitate was collected by centrifugation, the supernatant was discharged and the solid was dried in vacuum for HEL modification.

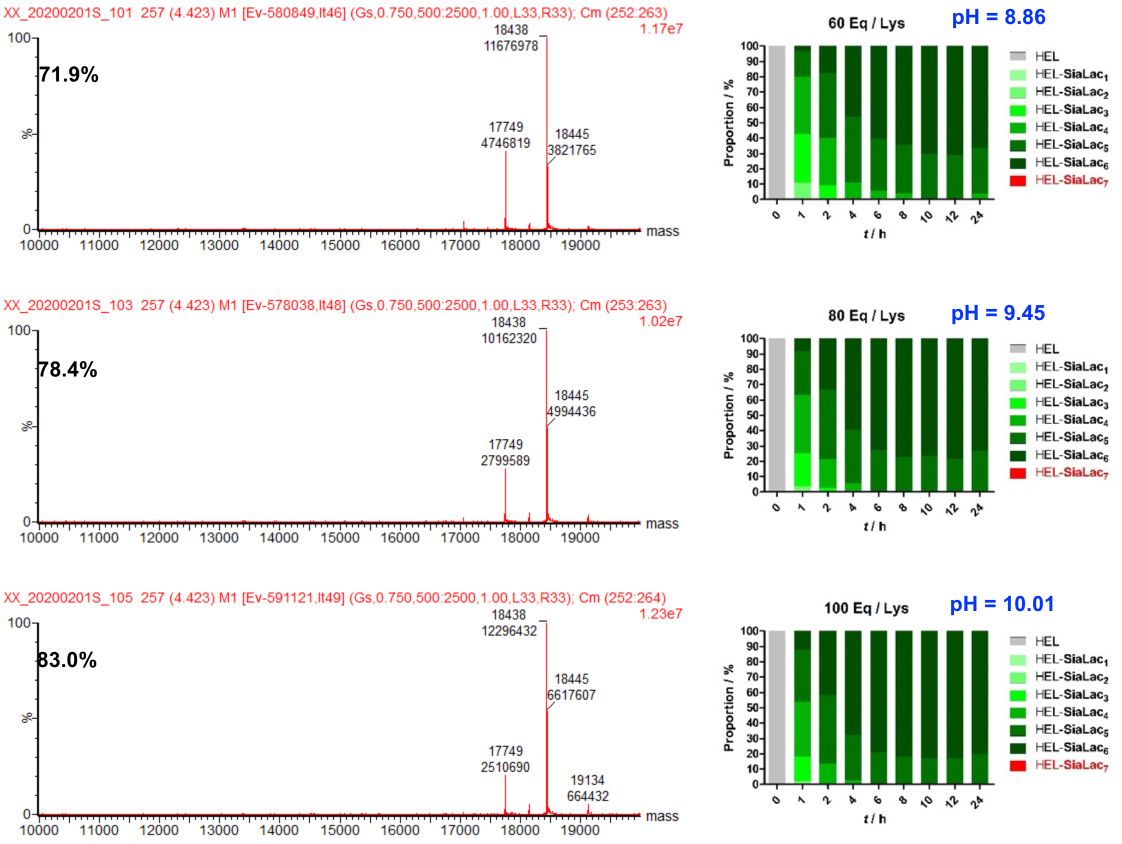


pH = 10.31

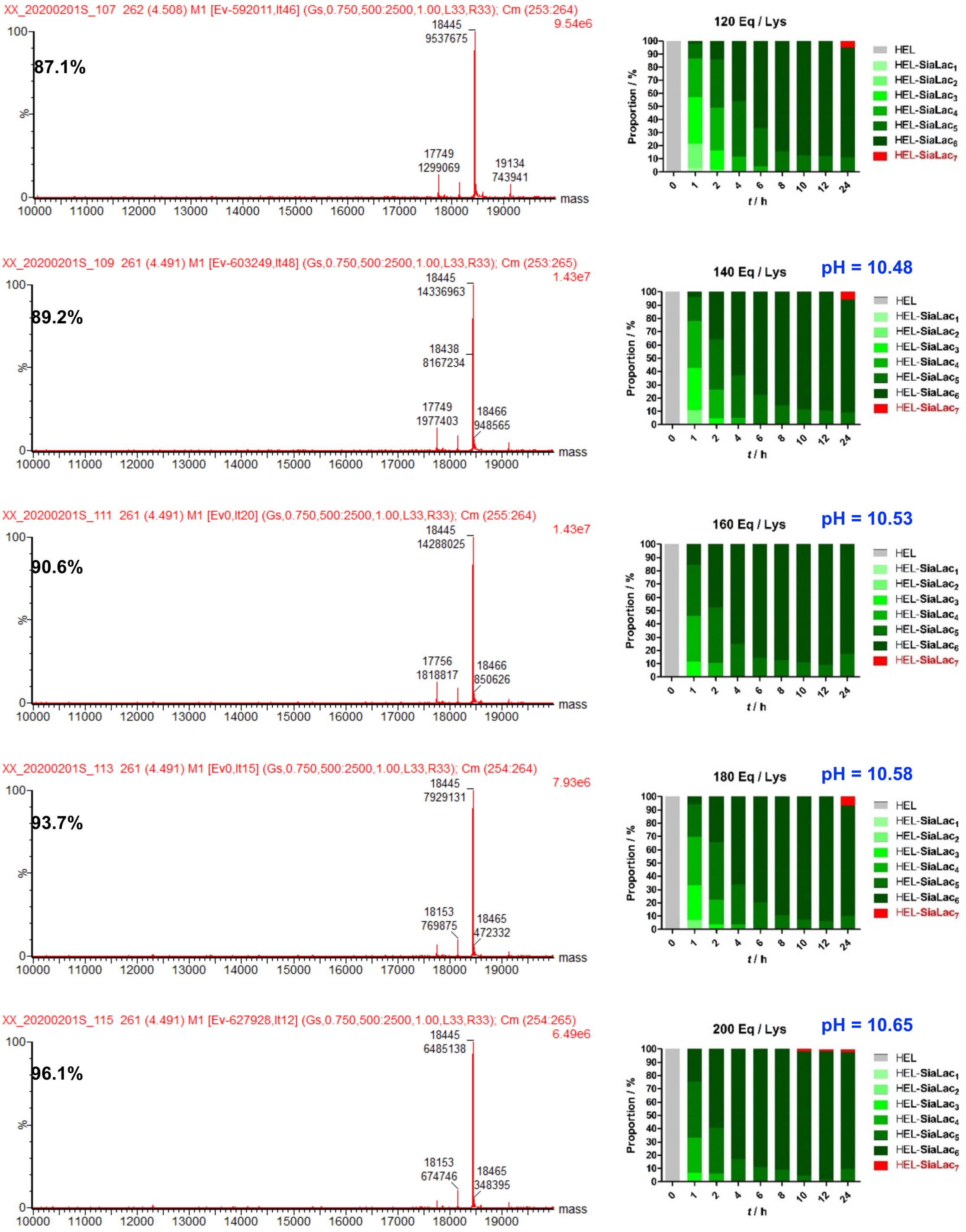


**Figure I-13.** HEL modification by using the freshly precipitated sugar **7** in PBS buffer. The actual pH value of mixture and the corresponding yield of HEL_SiaLac$_6$ were noted. Illustrated LC_MS was the one when t = 24 h.

As shown in **Figure I-13**, the conversion was highly efficient, especially where 200 equivalents of imidate **7** (per lysine) was added. The optimal yield of HEL-SiaLac$_6$ (96%) occurred at 12 h, where excessive reaction time caused a slight loss of yield. One possible reason was that one of SiaLac attached on lysine residues was unstable in the alkaline solution. As shown in **Figure I-13**, HEL-SiaLac$_6$ was slightly hydrolyzed to HEL-SiaLac$_5$.

Interestingly, LC_MS of HEL_SiaLac$_6$ gave a pair of peaks in all batches. The smaller one was what we expected, and the larger one appeared to be added seven Dalton. The ratio was quite dependent on the reaction condition (**Figure I-13**) and time (attached in Section 7.2). One speculation was that they are two states of one molecule.

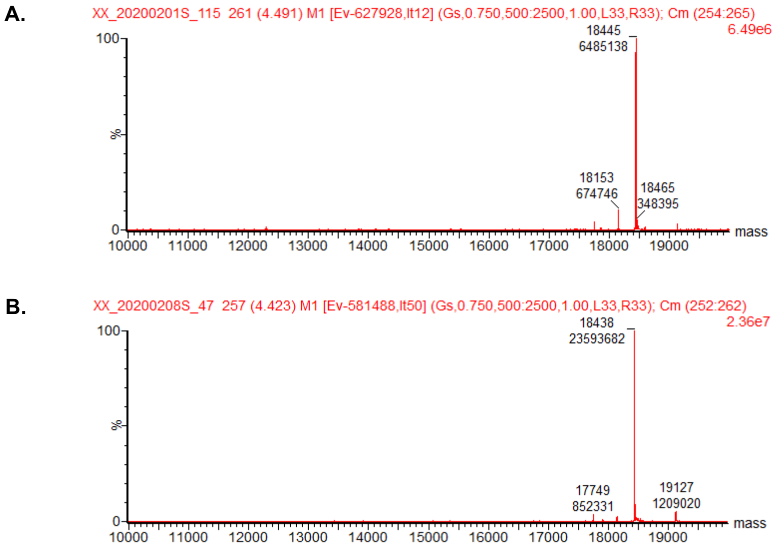


**Figure I-14.** LC_MS of SiaLac$_6$. A: in ammonium acetate (100 mM, pH = 8.0) buffer; B: in formic acid (1% in MQ water) solution.

Generally, LC_MS was collected in ammonium acetate (100 mM, pH = 8.0) buffer. However, when the LC_MS was checked again in formic acid (1%) solution, only the expected peak was observed; the extra one disappeared. (**Figure I-14**). These clarified that the mass spectrum of HEL_SiaLac$_6$ was pH-dependent.

With the optimal condition in hand, a large scale of HEL modification (3.5 mg of HEL) was performed. The reaction mixture was submitted for desalting on PD-10 column twice followed by dialysis in PBS buffer, after concentration and sterilization, the concentration was characterized with BCA assay, yielding HEL-SiaLac$_6$ in 95% (7.13 mg/ml in 0.6 ml) (**Figure I-15**).

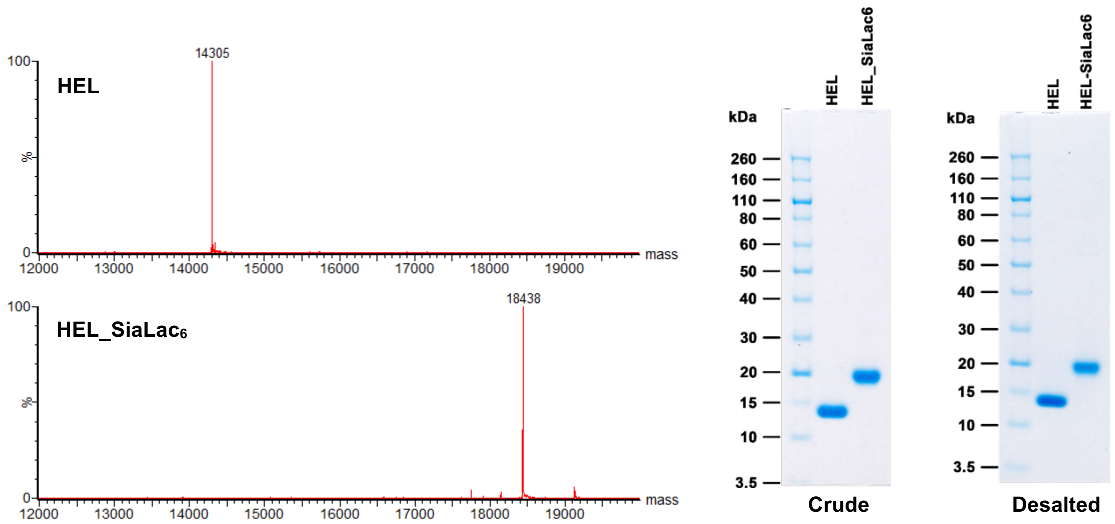


**Figure I-15.** Characterization of HEL_SiaLac$_6$.

#### II. gp120-[–amidine-GM3g]

In addition to HEL and its mutants, gp120 is another carrier protein we plan to do. A pilot of gp120_SiaLac preparation was done by using the standard condition developed previously. The modification process was monitored with SDS-PAGE, a clear migration proved the successful conjugation (**Figure II-1A**). A following big batch modification failed because of a massy sugar activation (**Figure II-1B**). The protein was then recovered from the reaction mixture for re-modification, but it's much worse when the recovered protein was used for some reason, unfortunately, as shown in **Figure II-1C**. One more batch worked nicely in which a fresh gp120 was employed (**Figure II-1D**), yielding 2.74 mg of gp120_SiaLac.

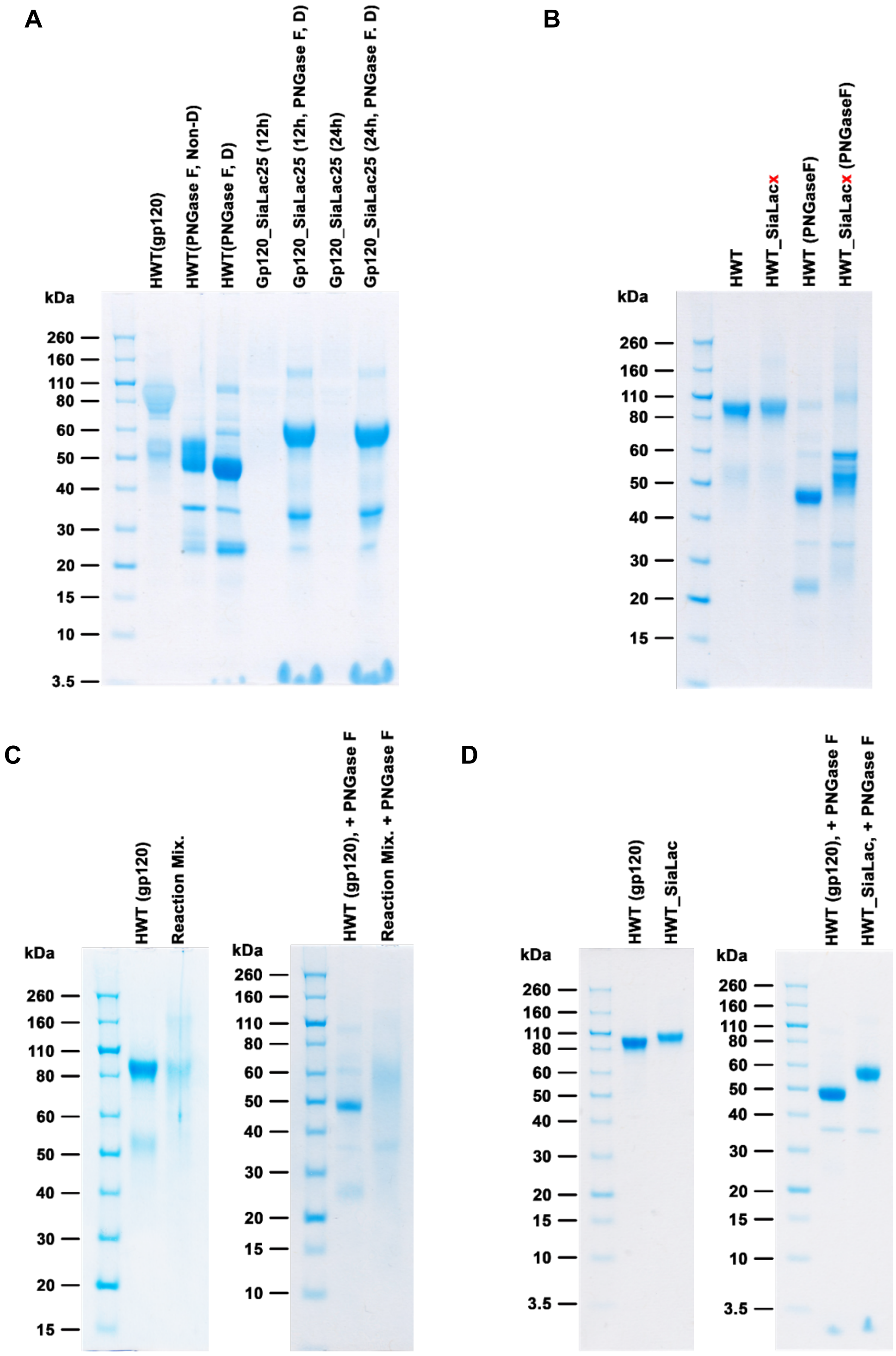


**Figure II-1.** SDS-PAGE of gp120_SiaLac.

#### III. HEL-[–alkylamine-GM3g]

To investigate the immunogenicity mechanism of the amidine linker in SiaLac modified wtHEL and HEL mutants, in this section, we aimed to conjugate SiaLac to HEL with an alternative linker: an aminoalkyl or *O*-link (SiaLacOCH$_2$CH$_2$NH-HEL).

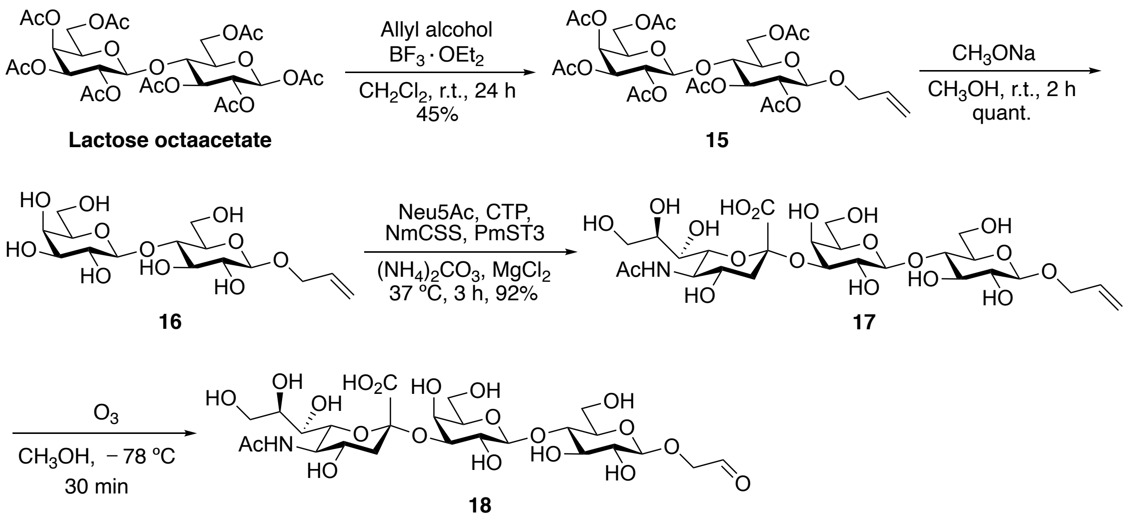


**Scheme III-1.** Synthesis of SiaLac-CHO (**18**)

As illustrated in **Scheme III-1**, Starting from lactose octaacetate, allyl glycoside **15** was synthesised using BF$_3$·OEt$_2$ as the promotor. After removal acetyl groups in sodium methoxide solution, allyl glycoside **16** was sialylated in the standard enzymatic condition, yielding the desired sialyllactose trisaccharide **17** with the desired a-2,3 linkage. Ozonolysis of the terminal alkene of **17** generated the aldehyde **18**, which was directly used for protein modification without further purification.

Protein modification was carried out in a reductive amination condition: a mixture of HEL and fresh aldehyde **18** was dissolved in distilled water followed the addition of sodium cyanoborohydride solution. The resulting reaction was incubated overnight at 37 °C and the crude solution was checked by SDS-PAGE (**Figure III-1**) and LC-MS (**Figure III-2**).

| *No.* | *Sugar* (eq/lys) | NaCNBH$_3$ (eq/lys) | *No.* | *Sugar* (eq/lys) | NaCNBH$_3$ (eq/lys) |
|---|---|---|---|---|---|
| **A1** | 5 | 10 | **C1** | 5 | 30 |
| **A2** | 10 | 10 | **C2** | 10 | 30 |
| **A3** | 20 | 10 | **C3** | 20 | 30 |

| | | | | | |
|---|---|---|---|---|---|
| **A4** | 50 | 10 | **C4** | 50 | 30 |
| **B1** | 5 | 20 | **D1** | 5 | 40 |
| **B2** | 10 | 20 | **D2** | 10 | 40 |
| **B3** | 20 | 20 | **D3** | 20 | 40 |
| **B4** | 50 | 20 | **D4** | 50 | 40 |

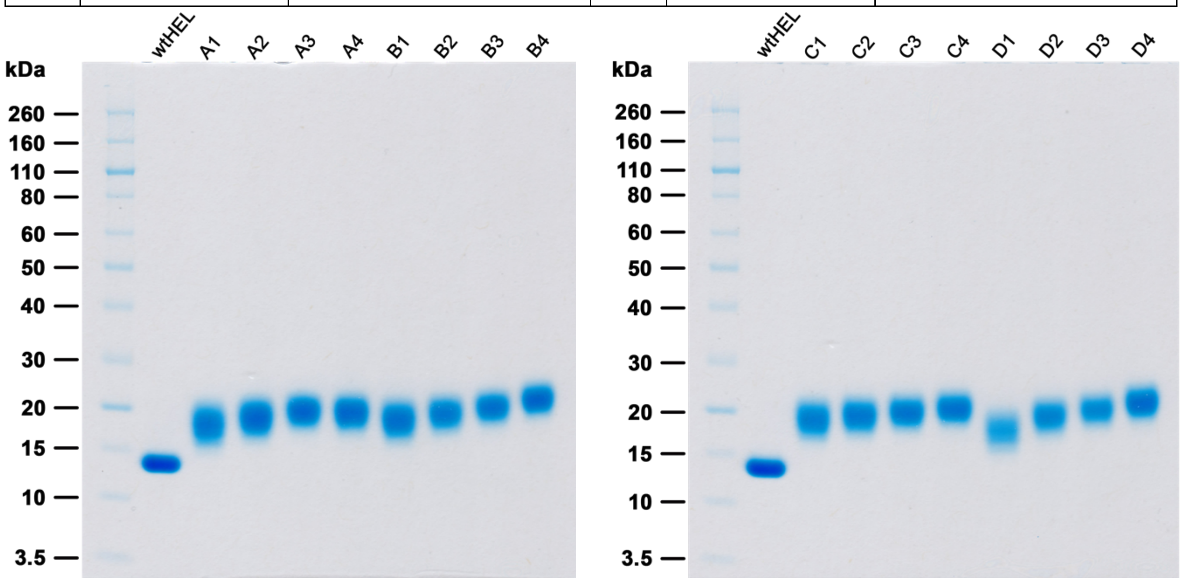


**Figure III-1.** SDS-PAGE of the crude reaction after incubation for 12h.

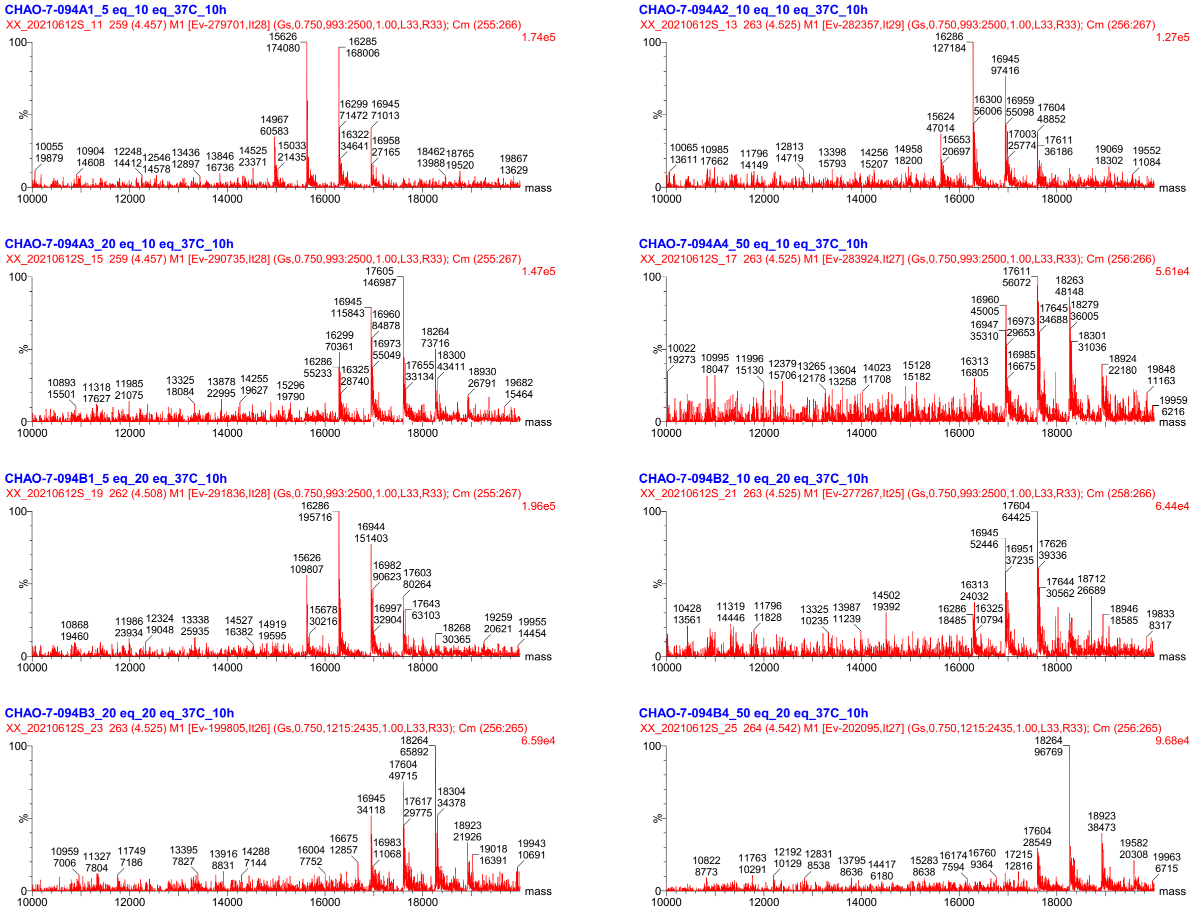

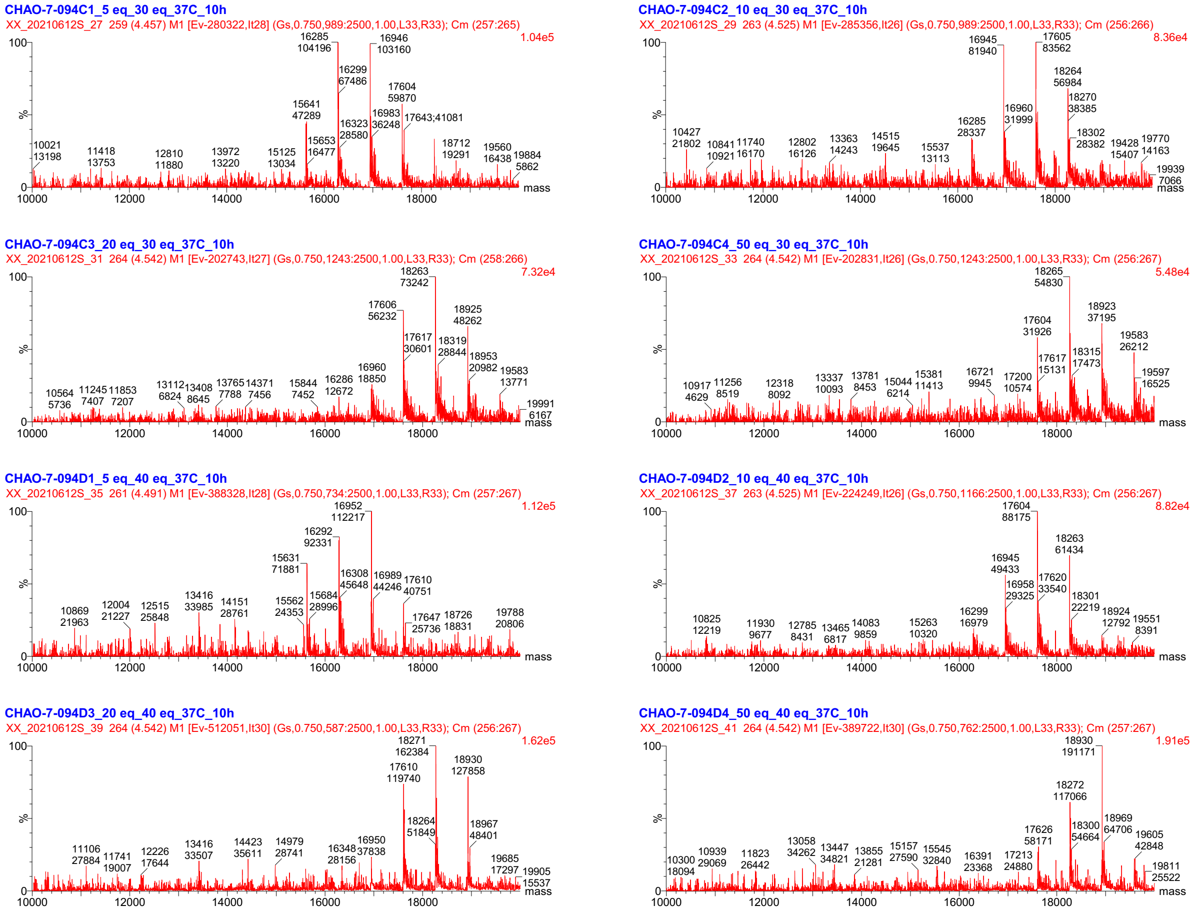


**Figure III-2.** LC-MS of the crude reaction after incubation for 12h. Theoretical MWs: unmodified HEL: 14305; HEL-SiaLac$_1$: 14965; HEL-SiaLac$_2$: 15624; HEL-SiaLac$_3$: 16284; HEL-SiaLac$_4$: 16943; HEL-SiaLac$_5$: 17603; HEL-SiaLac$_6$: 18263; HEL-SiaLac$_7$ (Lys & *N*-terminus): 18922.

From the mass spectra analysis, the modification worked under reductive amination. These data suggest that over-modification occurred, likely reflecting modifications at the *N*-terminus or arginine residues; reductive methylation affected the mass signals; the formaldehyde formed in ozonolysis, and the poor signal/noise ratio indicated a significant loss of protein due to degradation in the presence of sodium cyanoborohydride. Modification condition required to be further optimized.

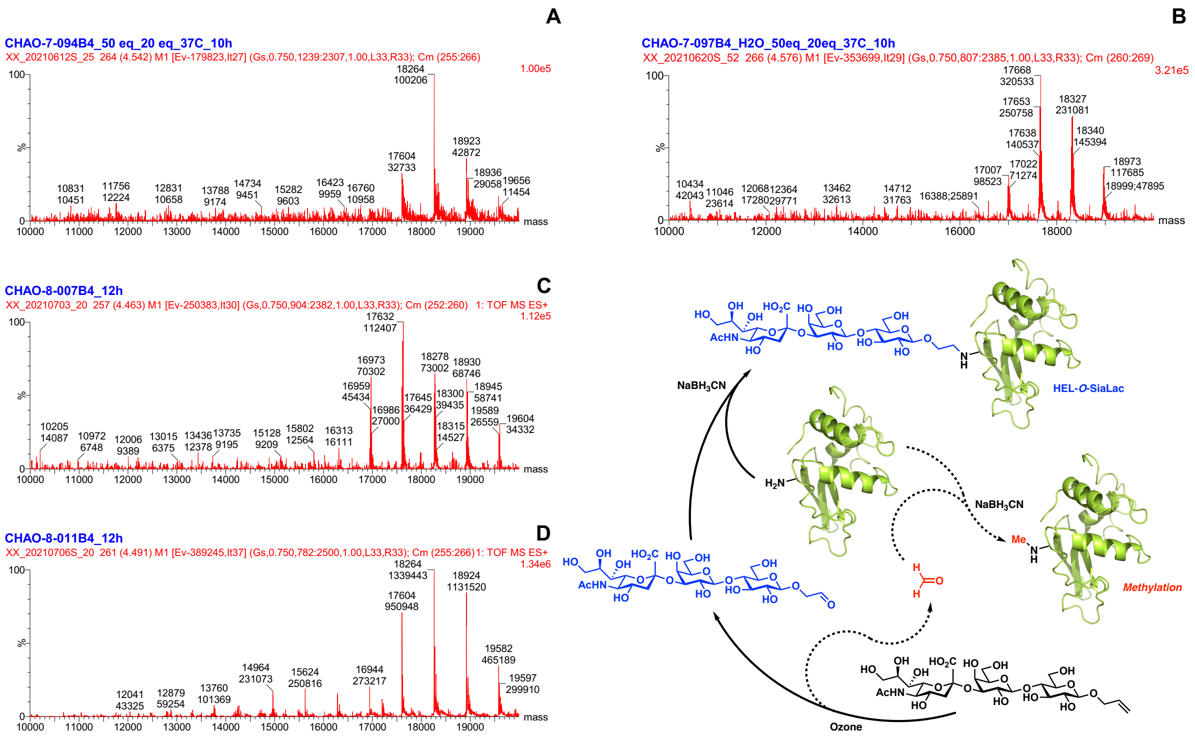

**Figure III-3.** Correlation between sugar purity and modification efficiency.

From the initial data, we know that a relatively ideal condition was a combination of 50 eq. of sugar and 20 eq. of $NaBH_3CN$, as shown in **Figure III-3A**. The second test gave a different spectrum, with a multi-peak (plus 14 Da) for each signal, indicating protein methylation (likely derived from the formaldehyde during ozonolysis) (**Figure III-3B**). Formaldehyde was not completely removed from the crude sugar in vacuum, resulting a competitive methylation reaction of lysine (or arginine) on protein. To get rid of formaldehyde, the crude sugar from ozone cleavage was dissolved in water followed by lyophilization overnight. Mass data (**Figure III-3C**) showed that methylation occurred, indicating formaldehyde was present, and that lyophilization alone was insufficient.

Next, we performed size-exclusion purification. The crude residue (even though appeared pure on TLC) was submitted to LH20. The pooled fractions were combined and concentrated and lyophilized in water to yield a white powder. When the reductive amination modification was repeated with this batch of sugar, no methylation happened. As shown in **Figure III-3D**. Complete removal of formaldehyde is crucial for reductive amination of protein. It's notable that: 1) methylation does happen at lysine (or arginine) residue, which blocks the expected sugar modification, and 2) since methylation occurs randomly at the lysines, arginines, and *N*-terminus, methylated sites are much more stable for proteinases. Stabilizing immunogens may have inadvertent effects on antigen presentation and immune responses. Therefore, we suggest a rethinking of the immunological data from previous papers in that they always employed crude material to conjugate glycans to carrier proteins for animal experiments.

A

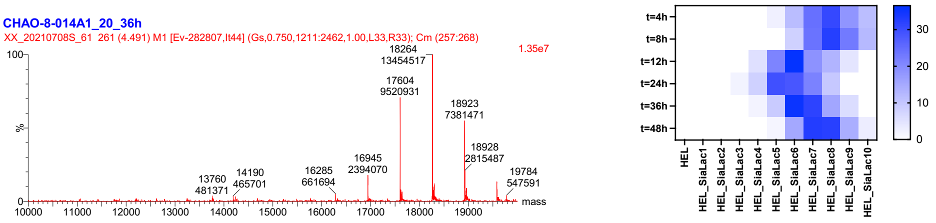
CHAO-8-014A1_20_36h
XX_20210708S_61 261 (4.491) M1 [Ev-282807,It44] (Gs,0.750,1211:2462,1.00,L33,R33); Cm (257:268)
1.35e7
18264 13454517
17604 9520931
18923 7381471
18928 2815487
16945 2394070
16285 661694
14190 465701
13760 481371
19784 547591
%
mass
t=4h
t=8h
t=12h
t=24h
t=36h
t=48h
HEL
HEL_SiaLac1
HEL_SiaLac2
HEL_SiaLac3
HEL_SiaLac4
HEL_SiaLac5
HEL_SiaLac6
HEL_SiaLac7
HEL_SiaLac8
HEL_SiaLac9
HEL_SiaLac10

B

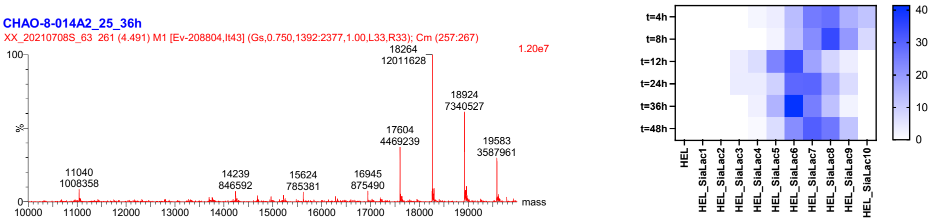
CHAO-8-014A2_25_36h
XX_20210708S_63 261 (4.491) M1 [Ev-208804,It43] (Gs,0.750,1392:2377,1.00,L33,R33); Cm (257:267)
1.20e7
18264 12011628
18924 7340527
17604 4469239
19583 3587961
11040 1008358
14239 846592
15624 785381
16945 875490
%
mass
t=4h
t=8h
t=12h
t=24h
t=36h
t=48h
HEL
HEL_SiaLac1
HEL_SiaLac2
HEL_SiaLac3
HEL_SiaLac4
HEL_SiaLac5
HEL_SiaLac6
HEL_SiaLac7
HEL_SiaLac8
HEL_SiaLac9
HEL_SiaLac10

C

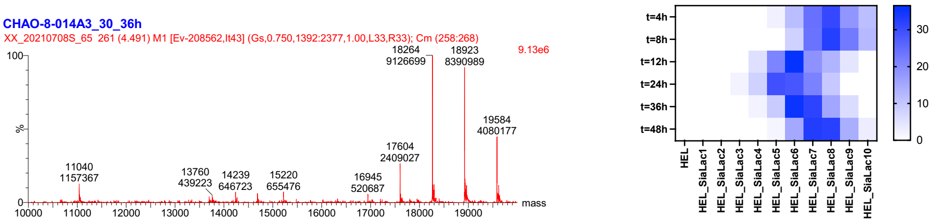
CHAO-8-014A3_30_36h
XX_20210708S_65 261 (4.491) M1 [Ev-208562,It43] (Gs,0.750,1392:2377,1.00,L33,R33); Cm (258:268)
9.13e6
18264 9126699
18923 8390989
19584 4080177
17604 2409027
11040 1157367
13760 439223
14239 646723
15220 655476
16945 520687
%
mass
t=4h
t=8h
t=12h
t=24h
t=36h
t=48h
HEL
HEL_SiaLac1
HEL_SiaLac2
HEL_SiaLac3
HEL_SiaLac4
HEL_SiaLac5
HEL_SiaLac6
HEL_SiaLac7
HEL_SiaLac8
HEL_SiaLac9
HEL_SiaLac10

D

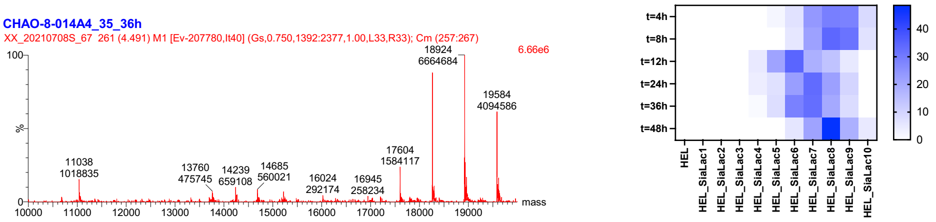
CHAO-8-014A4_35_36h
XX_20210708S_67 261 (4.491) M1 [Ev-207780,It40] (Gs,0.750,1392:2377,1.00,L33,R33); Cm (257:267)
6.66e6
18924 6664684
19584 4094586
17604 1584117
11038 1018835
13760 475745
14239 659108
14685 560021
16024 292174
16945 258234
%
mass
t=4h
t=8h
t=12h
t=24h
t=36h
t=48h
HEL
HEL_SiaLac1
HEL_SiaLac2
HEL_SiaLac3
HEL_SiaLac4
HEL_SiaLac5
HEL_SiaLac6
HEL_SiaLac7
HEL_SiaLac8
HEL_SiaLac9
HEL_SiaLac10

E

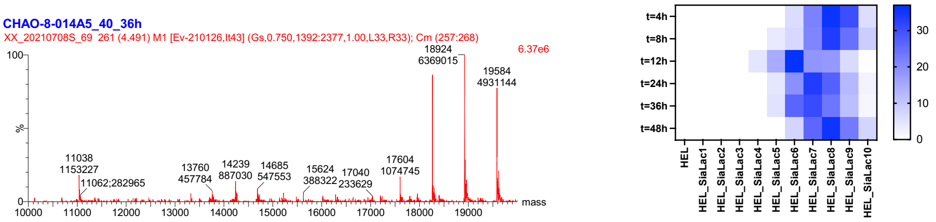
CHAO-8-014A5_40_36h
XX_20210708S_69 261 (4.491) M1 [Ev-210126,It43] (Gs,0.750,1392:2377,1.00,L33,R33); Cm (257:268)
6.37e6
18924 6369015
19584 4931144
11038 1153227
11062;282965
13760 457784
14239 887030
14685 547553
15624 388322
17040 233629
17604 1074745
%
mass
t=4h
t=8h
t=12h
t=24h
t=36h
t=48h
HEL
HEL_SiaLac1
HEL_SiaLac2
HEL_SiaLac3
HEL_SiaLac4
HEL_SiaLac5
HEL_SiaLac6
HEL_SiaLac7
HEL_SiaLac8
HEL_SiaLac9
HEL_SiaLac10

F

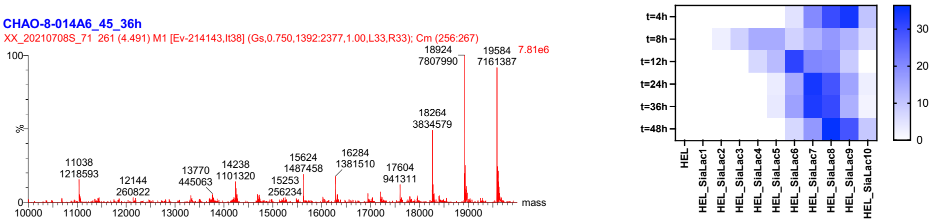
CHAO-8-014A6_45_36h
XX_20210708S_71 261 (4.491) M1 [Ev-214143,It38] (Gs,0.750,1392:2377,1.00,L33,R33); Cm (256:267)
7.81e6
18924 7807990
19584 7161387
18264 3834579
11038 1218593
12144 260822
13770 445063
14238 1101320
15624 1487458
15253 256234
16284 1381510
17604 941311
%
mass
t=4h
t=8h
t=12h
t=24h
t=36h
t=48h
HEL
HEL_SiaLac1
HEL_SiaLac2
HEL_SiaLac3
HEL_SiaLac4
HEL_SiaLac5
HEL_SiaLac6
HEL_SiaLac7
HEL_SiaLac8
HEL_SiaLac9
HEL_SiaLac10

G

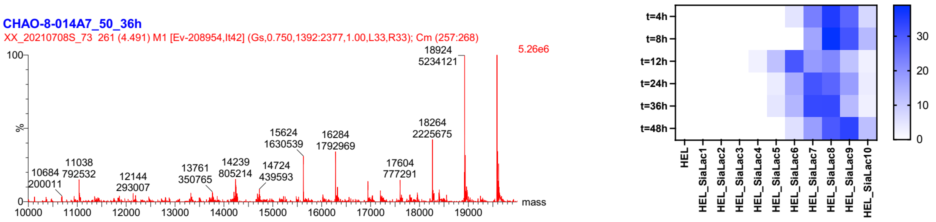
CHAO-8-014A7_50_36h
XX_20210708S_73 261 (4.491) M1 [Ev-208954,It42] (Gs,0.750,1392:2377,1.00,L33,R33); Cm (257:268)
5.26e6
18924 5234121
18264 2225675
15624 1630539
16284 1792969
17604 777291
10684 200011
11038 792532
12144 293007
13761 350765
14239 805214
14724 439593
%
mass
t=4h
t=8h
t=12h
t=24h
t=36h
t=48h
HEL
HEL_SiaLac1
HEL_SiaLac2
HEL_SiaLac3
HEL_SiaLac4
HEL_SiaLac5
HEL_SiaLac6
HEL_SiaLac7
HEL_SiaLac8
HEL_SiaLac9
HEL_SiaLac10

H

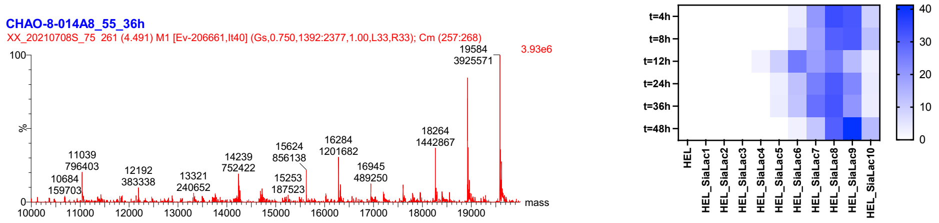


**Figure III-4.** Kinetics of reduction amination modification.

Having purified the sugar, protein modification was tested again. A time course mass analysis and distribution of sugar valency was plotted. For all the pilots here, 20 equivalents of $NaBH_3CN$ was used. As shown in **Figure III-4**, the equivalence used determined the sugar quanities loaded. Notably, over-alkylation occurred (up to ten SiaLac were installed), suggesting some lysine/arginine alkylation or double alkylation, since HEL sequence only covers seven free amine groups (six lysine residues and the *N*-terminus).

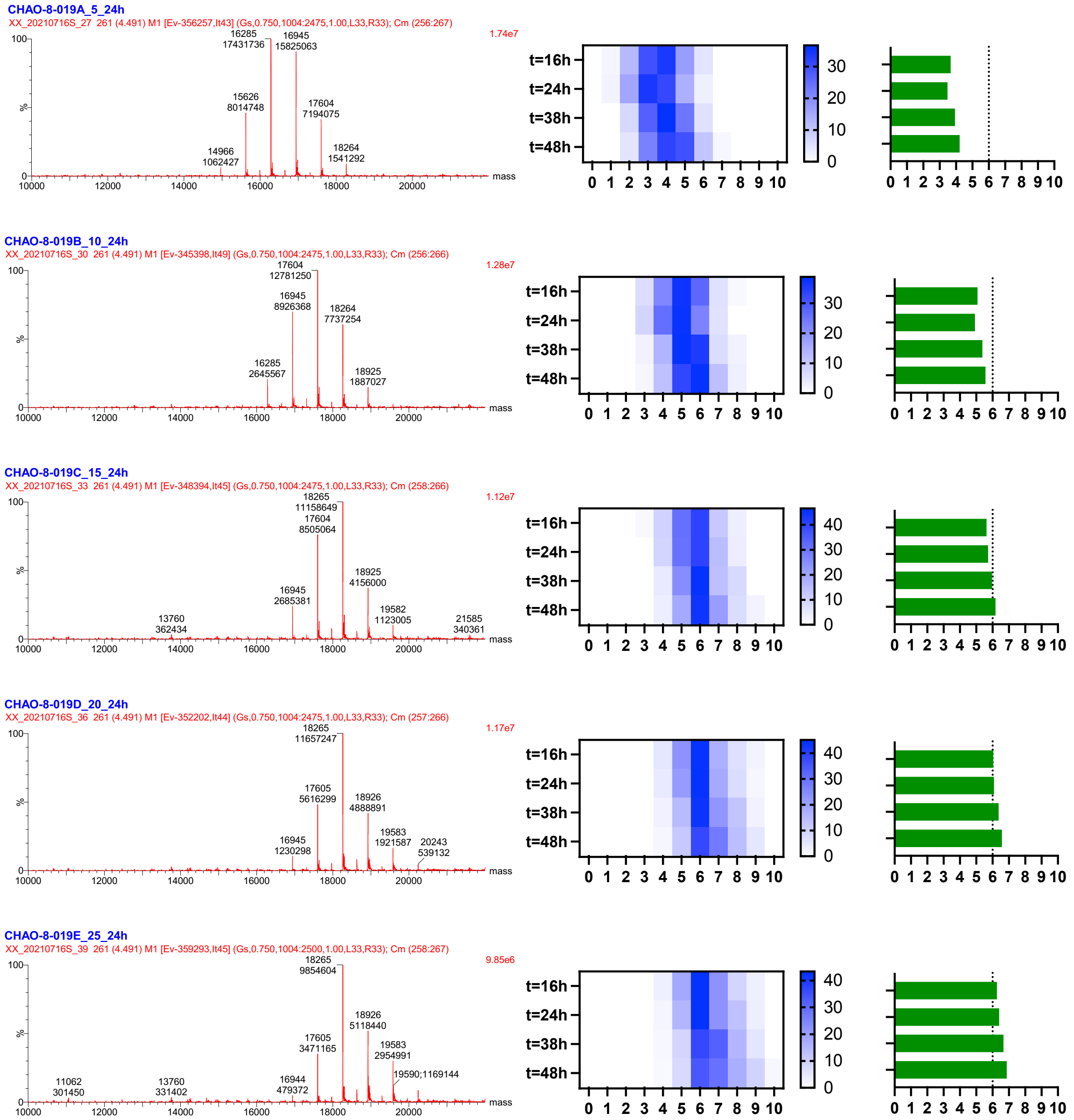

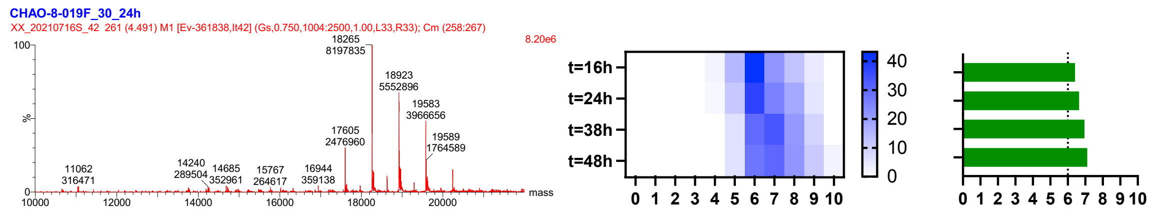


**Figure III-5.** Sugar equivalent and valency loading.

To ensure the O-link product was comparable to that of HEL-[–amidine-GM3g]$_6$, we aimed to get HEL modified in reductive amination conditions with exactly same sugar valency (that is, HEL-[–aminoalkyl-GM3g]$_6$. A second optimization was completed (**Figure III-5**), showing optimal conditions as follows: 1) HEL-[–aminoalkyl-GM3g]$_6$ preparation: 20 eq. Sugar, 20 eq. $NaCH_3CN$, 37 °C 24 h, and 2) HEL-[–aminoalkyl-GM3g]$_{3.7}$ preparation: 5 eq. Sugar, 20 eq. $NaCH_3CN$, 37 °C 24 h.

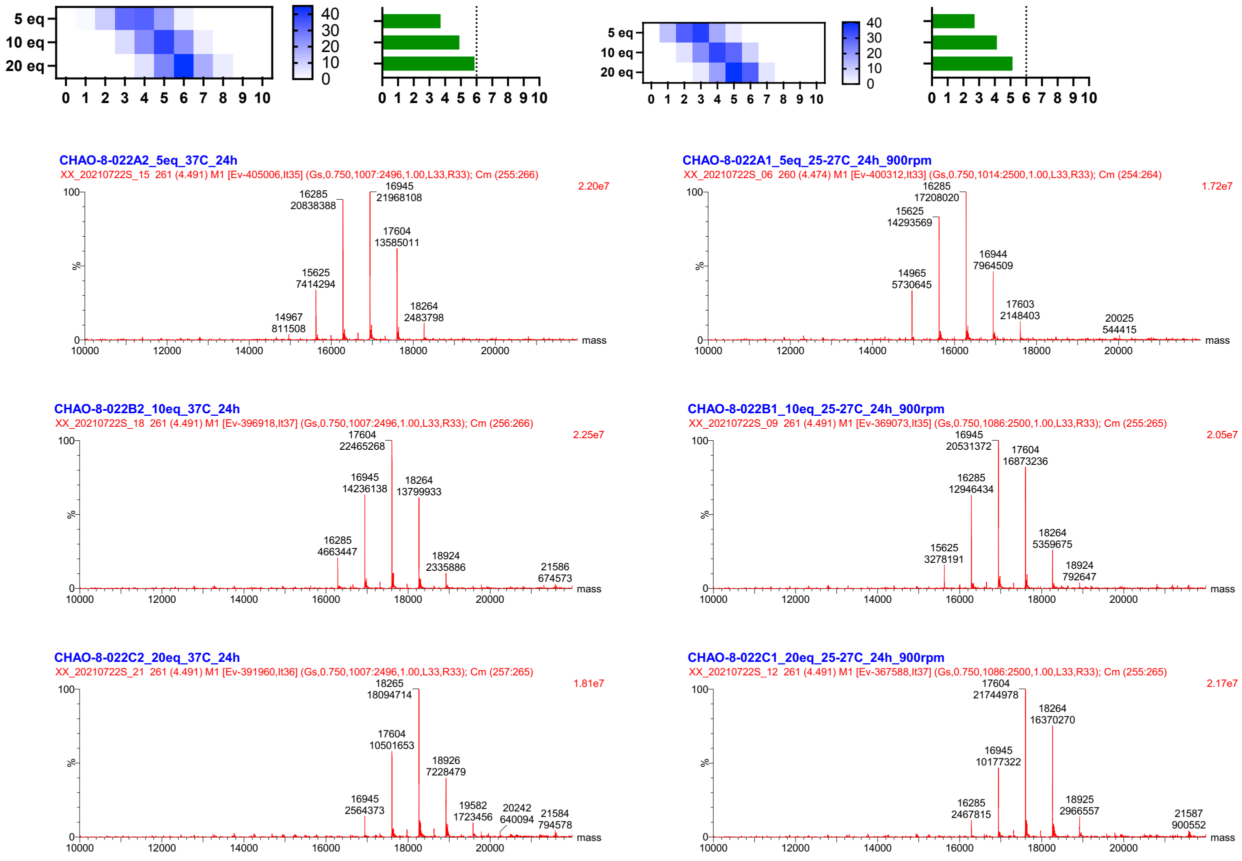


**Figure III-6.** Temperature-dependent valency of loaded SiaLac.

The pilots above were performed in PCR tubes that limited the modification scale. A pilot was done at 37°C, where the modification can be scaled up in the future if necessary. As shown in **Figure III-6**, modification does have a good reproducibility at 37°C. As a comparison, less valency was obtained when carried out in a thermoshaker at ambient temperature. A following scaleup was conducted (**Figure III-7**). After dialysis to PBS, concentration, sterilization, protein quantification and endotoxin assay, modified HEL immunogens were ready for vaccination (**Figures III-8 and III-9**).

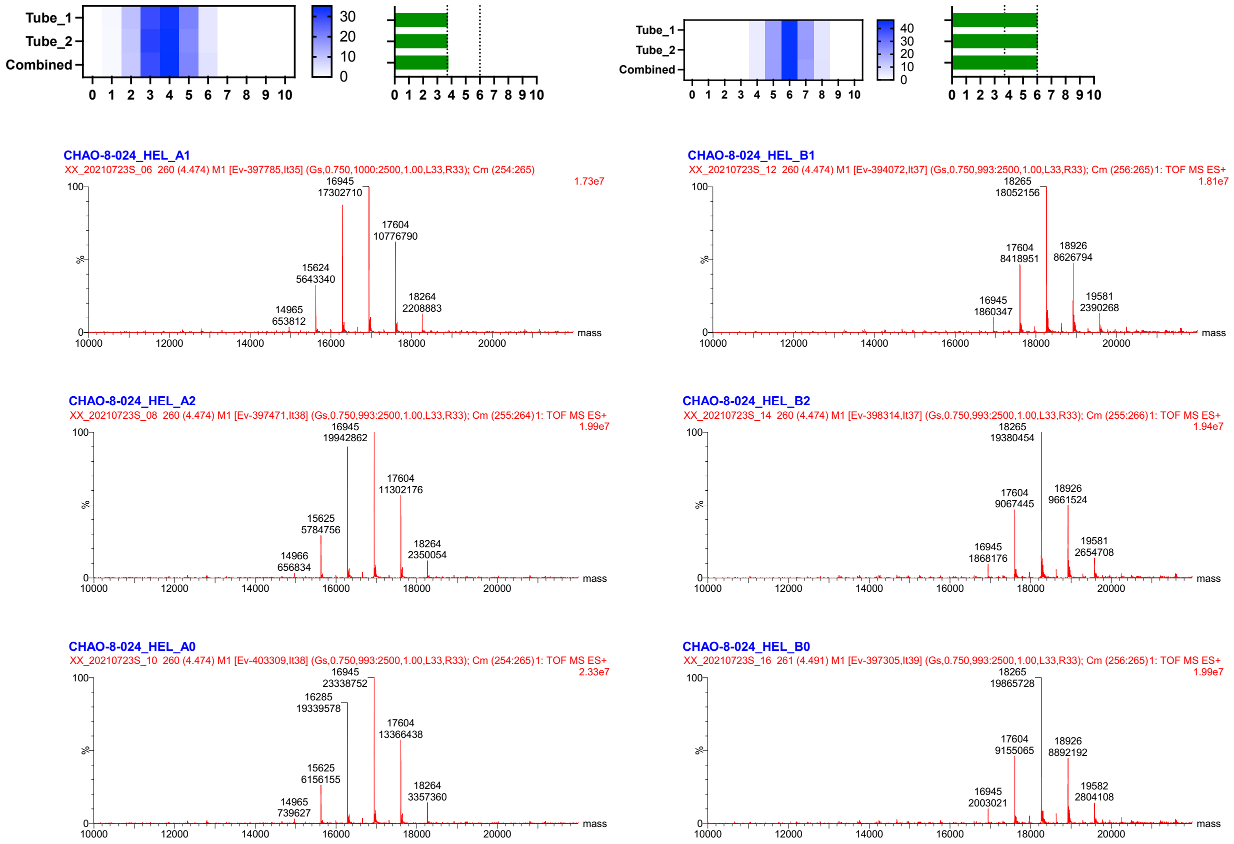


**Figure III-7.** Scale-up of HEL-(*O*-link)-SiaLac.

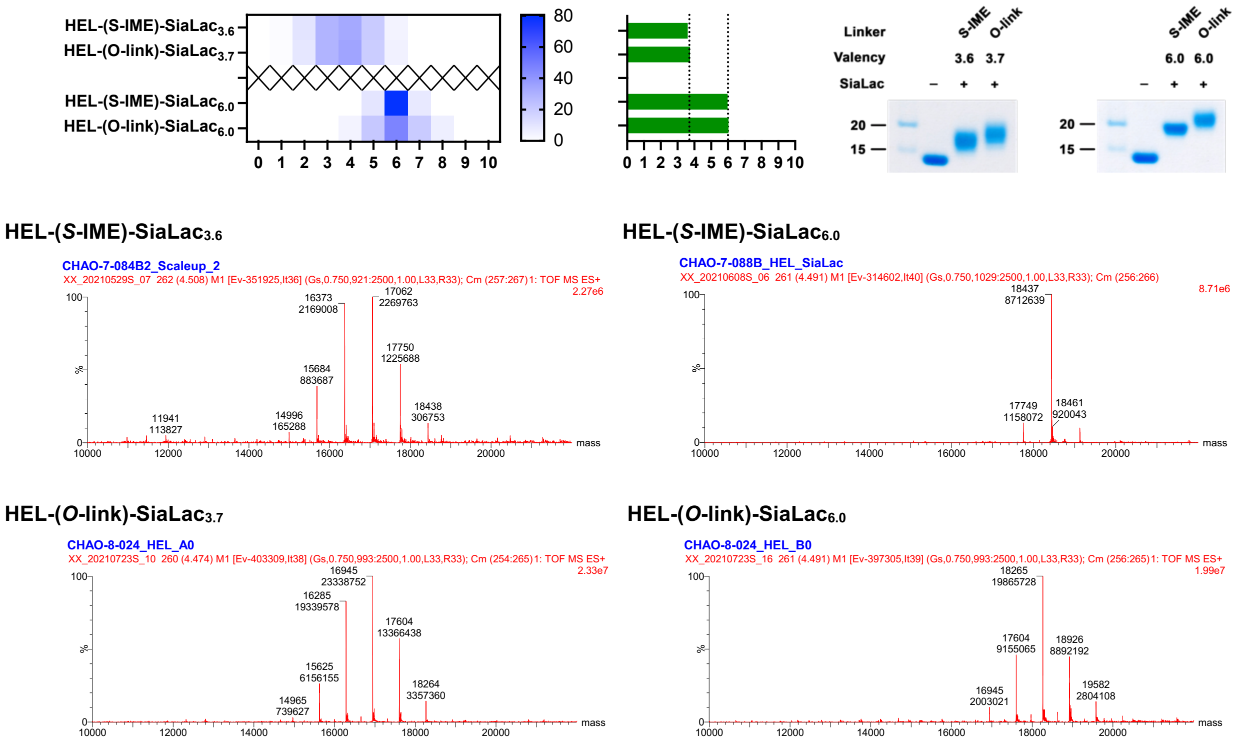


**Figure III-8.** Comparison of HEL-(*S*-IME)-SiaLac and HEL-(*O*-link)-SiaLac.

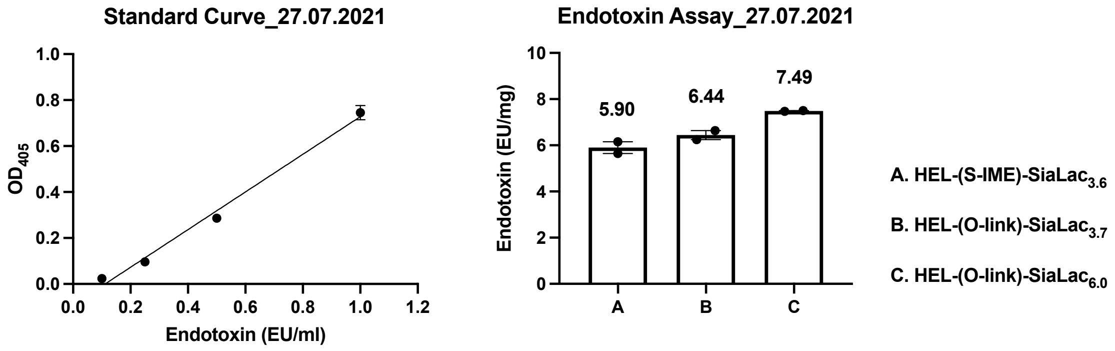


**Figure III-9.** Endotoxin quantitation of immunogens.

Following the same condition, reductive amination of BSA was done as well, yielding the BSA-(*O*-link)-SiaLac as the coating protein of ELISA (**Figure III-10**).

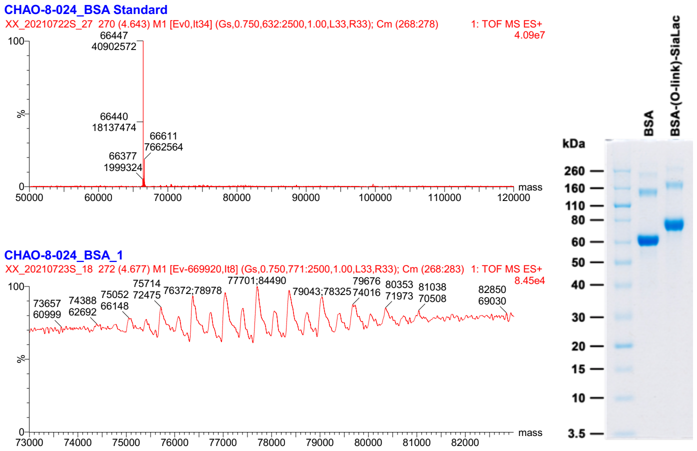


**Figure III-10.** Preparation of BSA-(*O*-link)-SiaLac.

#### IV. HEL-[–amide-GM3g]

The amidine linker is positively charged under physiological conditions. To probe the immunogenicity of amidine, here, we aimed to couple SiaLac to HEL via amide, a bioisostere of amidine.

A known 'two-step' strategy to synthesise this amide linker is possible (*16*). As shown in **Figure IV-1**, an iodo molecule could be introduced via the reaction between a primary amine and the heterobifunctional linker bearing iodo and NHS ester heads. The iodo then could be replaced with free thiol sugar.

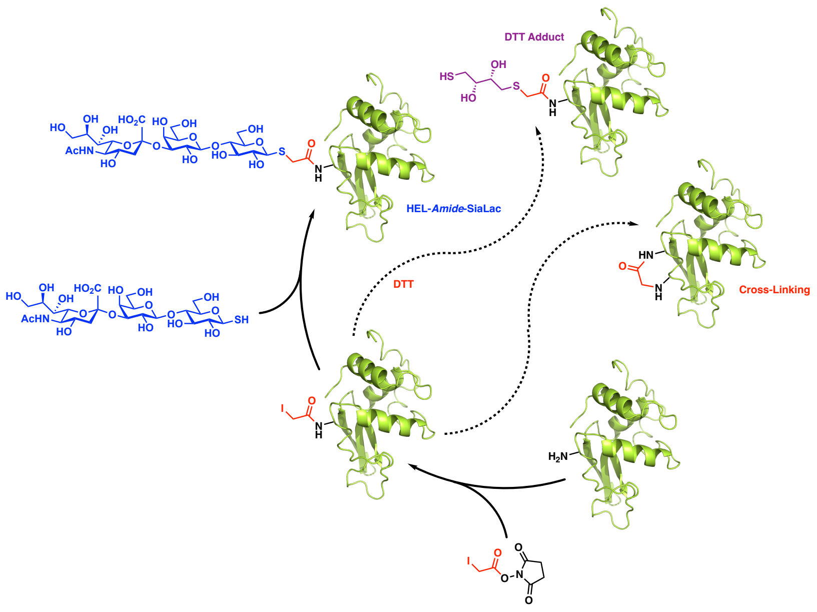


**Figure IV-1.** The diagram of the 'two-step' amidation.

We attempted to synthesise SiaLac-SH (**24**). As shown in **Scheme IV-1**, thiol acetate **4** was hydrolyzed with hydrazine acetate, and thiosugar intermediate was reprotected with Xanthyl group, a stable protecting group of anomeric thiol (*16*). Acetates were removed by the treatment of sodium methoxide in methanol, yielding substrate **26** for enzymatic reaction (54% yield over three steps). Sialic acid was then introduced in the standard conditions, giving trisaccharide **27** (75% yield). Deprotection of xanthyl group in iodine methanol solution generated dimeric SiaLac **28** but monomer SiaLac-SH (**24**), which has been reported previously (*17*). NMR spectra indicated dimer **28** is a symmetric β-isomer. Even though we did not generate monomer **24** directly, it was generated by TCEP reduction *in situ* for protein modification.

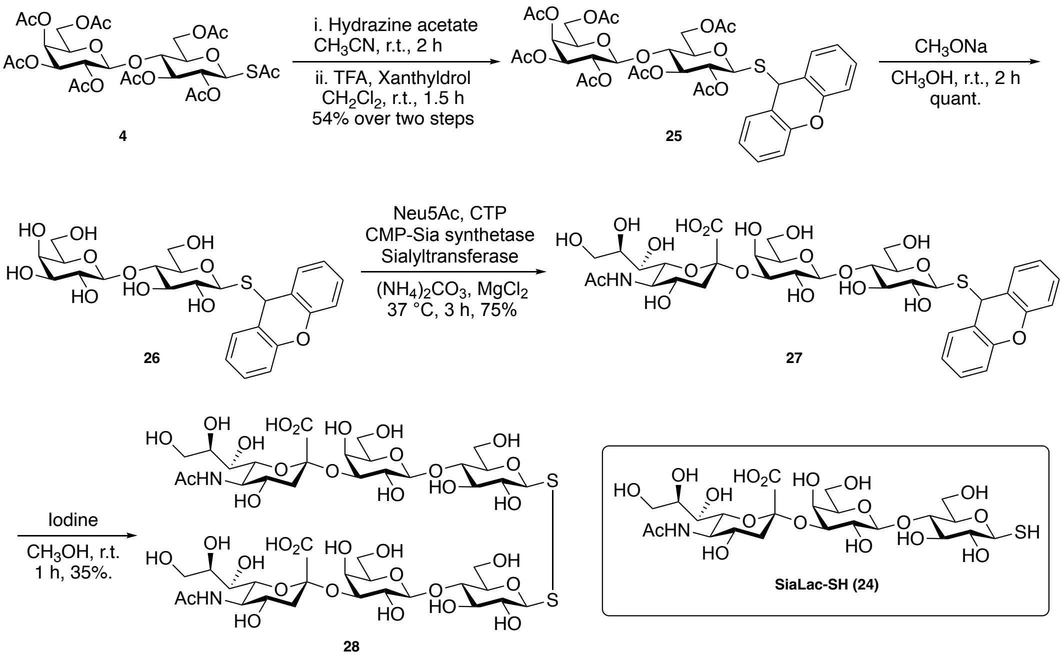


**Scheme IV-1. Synthesis of SiaLac-SH (24)**

Once sugar was ready, incubation of HEL with iodoacetic acid *N*-hydroxysuccinimide ester (I-Short-NHS), 8 eq. per lysine residue gave a good loading with six as the mean valency. For sugar introduction, dimer **28** was reduced with 1.0 eq. of TCEP, the freshly formed SiaLac-SH (**24**) was added into HEL bearing iodo-linker, the desired HEL-(*S*-amide)-SiaLac was obtained after incubation for 3 hours.

Before immunizing mice, a concern of the structure is whether SiaLac-SH coupled to protein in a complete β-isomer. Hence, we repeated the procedure in an NMR tube and assessed the coupling. This was conducted as follows:

1. Collect $^1$H NMR of dimer **28** in $D_2O$, as shown in the bottom.

2. Collect $^1$H NMR of TCEP in $D_2O$ for comparison.

3. Add 1 eq. of TCEP to the dimer solution in step 1, collect $^1$H NMR again after 30 min at room temperature. as shown in the figure, β-isomer formed only ($J_{1,2}$ = 7.8 Hz), and no TCEP left in comparison with TCEP spectrum, which is important, HEL will not be reduced (there're four disulfide bonds).

4. Transfer solution from 3 into coupling buffer (sodium borate in $D_2O$) and check the dynamics up to 12 hours: No α-isomer observed but a slight dimerization occurred. Dimerization is not a problem for protein modification.

5. Add iodoacetamide into the solution above ( $t$ = 12 hours) and leave it for 3 hours at room temperature, collect $^1$H NMR.

6. All the operations above are done in deuterated solvents, for double-checking, repeat 3-5 in water (exactly same as protein modification condition), lyophilize and collect $^1$H NMR in $D_2O$.

7. Collect $^1$H NMR of the β-isomer standard, SiaLac-S-Amide (**31**) that prepared from SiaLac-CN (**1**) (shown later).

8. For $^1$H NMR in 5, 6, and 7, comparison with each other shows SiaLac-S-Amide formed in 5 and 6 are β-isomers.

NOTE: all conditions here (temperature/mass/concentration/eq. of reagents/reaction time) are same as what have been used in the preparation HEL-Amide-SiaLac, but iodoacetamide used as a model.

LIMITATION: *a*. We don't have the α-isomer for comparison, purity of spectra is good, but we have no idea if tiny α-isomer is there (only impurity > 5% can be detected in proton NMR). *b*. It's impossible to reduce protein with TCEP since no TCEP left after sugar reduction. A question is whether the excess SiaLac-SH (2.68 mM in protein solution) can reduce protein. *c*. Protein LC_MS is not that clean if comparing with IME modification, whether the impurity (probably is cross coupling?). Endotoxin assay showed it's clean enough for immunization.

#### V. Free GM3g-amidine, GM3g-alkylamine, GM3g-amide ligands

Initially, we started from SiaLac-CN **1**. As shown in **Scheme V-I**, cyano was converted to imidate by $CH_3ONa$ solution, giving an inseparable mixture of cyano **1** and imidate **7**. Treatment of the mixture with $CH_3NH_2$ afforded another mixture of cyano **1**, imidate **7**, and amidine **8**. The barrier in this route was that SiaLac-amidine **8** was too polar to be purified by normal chromatography.

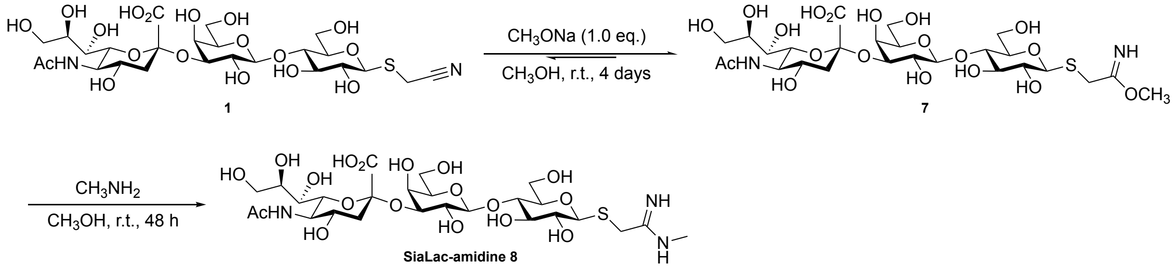


**Scheme V-I.** Assembly of SiaLac-amidine **8** from SiaLac-CN **1**.

Thus, we focused on another strategy in which Lac-CN **6** was used as the material (**Scheme V-2**). Following the previous activation condition, cyano **6** was treated with $CH_3ONa$, yielding the imidate **9** as a white precipitate which was separated from the unreacted **6** by filtration. Imidate **9** then reacted with excessive $CH_3NH_2$. The white solid was collected and characterized as the corresponding Lac-amidine **10**. Sialic acid was then assembled enzymatically, the crude SiaLac-amidine **8** was directly purified by size exclusion chromatography.

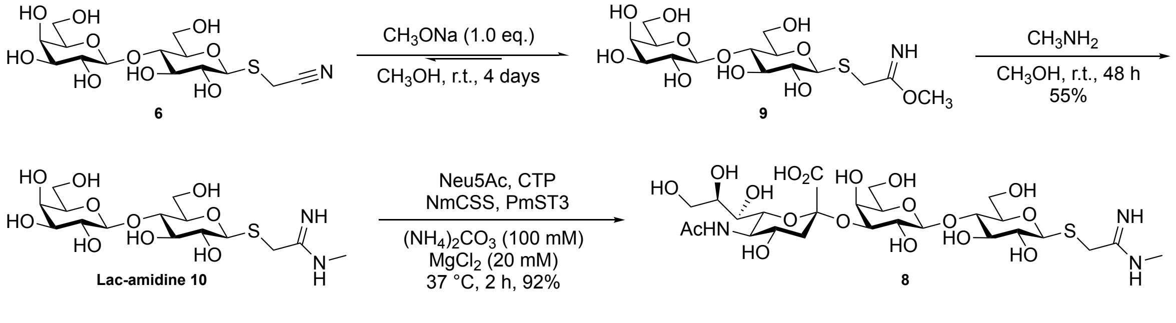


**Scheme V-2.** Assembly of SiaLac-amidine **8** from Lac-CN **6**.

Unfortunately, NMR characterization of amidines **10** and **8** showed both were mixture. Later, LC-MS analysis of **8** gave a mixture of amidine with methyl group, amidine(Me), and amidine without methyl group, amidine(H). This was reminiscent of the previously reported stability of amidine in basic conditions. Based on the literature, the amidine bond formed is stable in acidic buffer; however, it is possible to hydrolysis and cleave at high pH (*18*). A possible explanation: for Lac-amidine **10**, once it's been dissolved in $D_2O$ for NMR experiments, the residue $CH_3ONa$ from the previous activation reacts with $D_2O$, forming NaOD that can hydrolyze amidine. SiaLac-amidine **8** was enzymatically sialylated from **10**, in the enzymatical reaction. When amidine **10** was dissolved in ammonium carbonate (100 mM, pH = 8.5), NaOH was formed from the reaction between $CH_3ONa$ in **10** and water in the buffer, which raised the pH so ammonia in the solution hydrolyzed amidine **8**, yielding amidine(Me) **8** and amidine(H).

Bearing this in mind, a new batch of Lac-amidine **10** was made. Before NMR data collection and the following reaction, solid **10** was redissolved in 1 M HCl solution in methanol followed by adding excess ether to neutralize **10**, giving amidine hydrochloride (salt). For comparison, the basic Lac-amidine **10** (without neutralization) and the amidine HCl salt were submitted for NMR (in $D_2O$) again. From proton NMR, amidine decomposed completely after one day at room temperature; however, salt was stable. From the amidine salt, SiaLac-amidine **8** was obtained in a good purity after multiple flash column chromatography separation and LH-20.

Initially, we speculated that the impurify in the mixture was from contaminants in methylamine solution, but this turned out to not be the case: methylamine solution in MeOH and methylamine solution in THF freshly ordered from supplier were employed for Lac-amidine **10** preparation.

We also synthesized free aminoalkyl- and amide-based SiaLac ligands for competitive ELISA. As shown in **Scheme V-3**, reduction of SiaLacO-$N_3$ (**29**) (*19*) in Staudinger conditions yielded free amine **30**. SiaLacCN (**1**) was treated with a combination of $H_2O_2$ and potassium carbonate in DMSO, giving SiaLac-Amide (**31**) in 97% yield. Starting from Lactose derivative **12**, Lac-*S*-Amide(Me) (**32**) was prepared in EDC-mediated peptide coupling, enzymatic sialylation of **32** afforded SiaLac-*S*-Amide(Me) (**33**) in 92% yield.

**A**

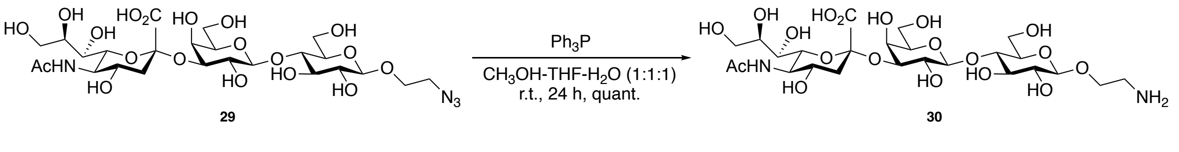


**B**

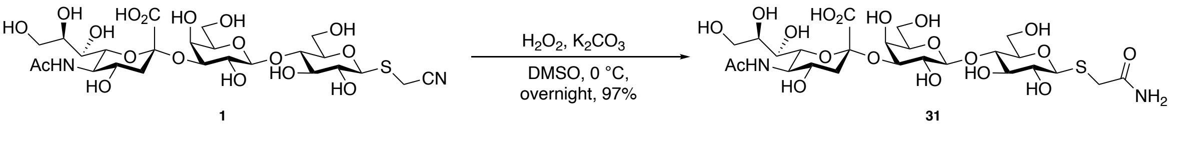

**C**

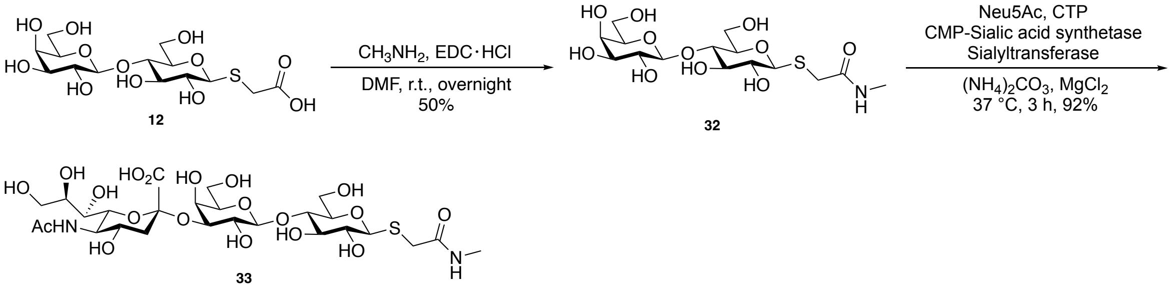


**Scheme V-3. Synthesis of free ligands.**

#### 2. Synthetic Methods

##### General experimental materials for chemistry

All reagents were purchased from commercial sources and were used without further purification unless noted. Molecular sieve (4Å, powder) used in reactions was activated at 350 °C for more than 12 hours. Dry solvents for reactions were purchased from *Sigma-Aldrich*, following abbreviations are used: PE = petroleum ether (*b.p.* 40 – 60 °C), EtOAc = ethyl acetate, THF = tetrahydrofuran. Thin Layer Chromatography (TLC) was carried out using Merck aluminium-backed sheets coated with Kieselgel 60-$F_{254}$ silica gel. Visualization of the reaction components was achieved using UV fluorescence (254 nm) and/or by charring with an acidified *p*-anisaldehyde solution in ethanol or a acidified cerium ammonium molybdate (CAM) solution in water. Organic solvents were evaporated under reduced pressure, and the products were purified by flash column chromatography on silica gel (230–400 mesh) and/or size exclusive chromatography on LH20. Proton Nuclear Magnetic Resonance ($^1$H NMR) spectra were recorded on a Bruker AVB400 (400 MHz), or AV700 (700 MHz) spectrometers, and the chemical shifts are referenced to residual $CHCl_3$ (7.26 ppm, $CDCl_3$), $CHD_2OD$ (3.30 ppm, $CD_3OD$), HDO (4.79 ppm, $D_2O$). Carbon Nuclear Magnetic Resonance ($^{13}$C NMR) spectra were recorded on a Bruker AVB400 (100 MHz), or AV700 (175 MHz) spectrometers and are proton decoupled, and the chemical shifts are referenced to $CDCl_3$ (77.16 ppm) or $CD_3OD$ (49.0 ppm). Assignments of NMR spectra were based on two-dimensional experiments (1H-1H COSY, DEPT-135, HSQC, and HMBC) if required. Reported splitting patterns are abbreviated as s = singlet, d = doublet, t = triplet, q = quartet, p = pentet, hept = heptet, m = multiplet, br = broad, app = apparent. Low Resolution Mass Spectra (LRMS) were recorded on a Micromass Platform 1 spectrometer using electrospray ionization (ESI), or on a Bruker Daltronic MicroTOF spectrometer. High Resolution Mass Spectra (HRMS) were recorded on a Bruker Daltronic MicroTOF spectrometer using electrospray ionization (ESI), *m*/*z* values are reported in Daltons. Optical rotations were measured on a Perkin-Elmer 241 polarimeter at 589 nm (Na D-line) with a path length of 1.0 dm at ambient temperature and are in units of degree $mL \cdot g^{-1} \cdot dm^{-1}$. Infrared (IR) spectra were recorded on a Bruker Tensor 27 Fourier Transform spectrophotometer using attenuated total reflectance (ATR) and Absorption maxima ($\nu$max) are reported in wavenumbers ($cm^{-1}$).

#### General experimental materials for biology

Plasmids pET23a-NmCSS and pET23a-PmST3 were kindly donated from Aziz; BL21(DE3) Competent E. coli (C2527), Q5 High-Fidelity DNA Polymerase, T4 Ligase, SOC Outgrowth medium, Quick-Load® Purple 1kb DNA Ladder (N0552S), and Gel Loading Dye Purple (6×, B7024S) were purchased from *NEW ENGLAND BioLabs* (*NEB*) Inc.; QuikChange II XL Site-Directed Mutagenesis Kit and XL10-gold competent cells were purchased from *Agilent* and stored at – 20 °C and – 80 °C, separately; TAE Buffer (50×) was ordered from *PanReac AppliChem* (ITW Reagents); general chemicals, lysozyme from chicken egg white (HEL, L6876), monoclonal anti-polyhistidine-alkaline phosphatase antibody (A5588-5ML), and BCIP®/NBT liquid substrate system (B1911-100ML) were purchased from *Sigma–Aldrich*; all primers (excluded **T7F** and **T7R**) for sequencing, PCR amplification and mutation were ordered from *Sigma–Aldrich*'s Oligo/Prime Store; DNA sequencing was performed by *Source BioScience*'s sequencing service using free stock primers (**T7F** or **T7R**) or designed primers; sequencing data were viewed and analysed by *SnapGene® Viewer* 4.2.6 together with online tools (ExPASy-translate, BLAST, reverse complement, and multiple/pairwise sequence alignment); PCR programs were set up by *Applied Biosystems* 2720 Thermal Cycler; DNA gel analyses and purification were developed using *Mupid®-One* Submarine Electrophoresis System; performed DNA gels were imaged in *Gel Doc™ XR+ System* (*Bio-Rad* gel documentation systems); SyBR™ Safe DNA Gel Stain (S33102), SDS-PAGE (NuPAGE Novex 4-12% Bis-Tris Protein Gels, 1.0 mm), Novex™ Sharp Unstained Protein Standard (LC5801, 3.5 to 260 kDa), Novex™ Sharp Pre-stained Protein Standard (LC5800, 3.5 to 260 kDa), NuPAGE™ MES SDS Running Buffer (20×) (NP0002), NisPur™ Ni-NTA Resin, and HisTrap™ HP Column (1 mL) were purchased from ThermoFisher SCIENTIFIC; InstantBlue™ Protein Stain (Coomassie Protein Stain for SDS-PAGE) was purchased from *Expedeon*; SDS-PAGEs were developed in Novex X-Cell SureLock™ Mini-Cell using Bio-RAD Powerpac™ Basic as the power; Western Blotting was performed in *i*Blot™ Gel Transfer Device (*Invitrogen*, IB1001); *Vivaspin* concentrators, PD MiniTrap G-25, and PD SpinTrap G-25 were ordered from *GE Healthcare*; concentrations of DNA (fragment or plasmid) were checked using *NanoDrop* 1000 3.8.0 (Nucleic Acid → Sample Type: DNA-50); concentrations of proteins were checked using *NanoDrop* 1000 3.8.0 ($Protein_{280nm}$ → Sample Type: Others) with corresponding molecular weight and extinction coefficient or BCA Assay; absorbance of 96-well plate was monitored on BMG LABTECH's *SPECTROstar Nano*.

#### General methods and operations

*Preparation of grown mediums*

i. *Luria-Bertani (LB) Broth*

LB Borth (25 g, Granulated, formula provides: Tryptone: 10g/L; Sodium Chloride: 10 g/L; Yeast Extract: 5g/L) and distilled water (1 L) were mixed and were autoclaved at 121 °C for 20 min. After cooling to room temperature, the stock solution of kanamycin (1 mL, 50 mg/mL) was added, mixed, and the resulting mixture was ready for use.

ii. *Luria-Bertani (LB) Broth agar plate*

Bacteriological agar (15 g/L, granulated) was added to LB medium prior to autoclaving. Cooled to around 55 °C, the stock solution (100 mg/mL) of ampicillin (1 mL/L) was added, mixed and poured into to plates (~20 mL/plate) which were left overnight at room temperature for cooling, stored at 4 °C for use.

*Buffers preparation*

i. *Buffers for protein purification*

**Binding buffer** (1 L, pH = 7.4): Tris·HCl (25 mM, 3.94 g), NaCl (500 mM, 29.22 g), imidazole (25 mM, 1.70 g), and distilled water (to 1 L). Mixed, adjusted the pH with HCl solution or NaOH solution, filtered with 0.2 μm filter, degassed by sonication, stored on ice prior to purification.

**Elution buffer** (1 L, pH = 7.4): Tris·HCl (25 mM, 3.94 g), NaCl (500 mM, 29.22 g), imidazole (250 mM, 17.02 g), and distilled water (to 1 L). Mixed, adjusted the pH with HCl solution or NaOH solution, filtered with 0.2 μm filter, degassed by sonication, stored on ice prior to purification.

**Phosphate-buffered Saline/PBS buffer** (1 L, pH = 7.4): NaCl (137 mM, 8.0 g), KCl (2.7 mM, 0.2 g), $Na_2HPO_4$ (10 mM, 1.44 g), $KH_2PO_4$ (2 mM, 0.24 g), and distilled water (to 1 L). Mixed, adjusted the pH with HCl solution, filtered with 0.2 μm filter, degassed by sonication, stored on ice prior to purification.

ii. *Buffer for protein LC_Mass analysis*

**Ammonia acetate** (100 mM, pH = 8.0): $CH_3COONH_4$ (770 mg), and distilled water (to 100 mL). Equilibrated, adjusted pH by using either acetic acid or ammonia hydroxide solution, filtered with 0.2 μm filter, degassed by sonication, stored at room temperature for use.

**Formic acid** (1% v/v) in MQ water

iii. <u>*Buffers for Western Blotting*</u>

**Phosphate-buffered Saline/PBS buffer** (pH = 7.4).

**Block buffer** (100 mL): Bovine serum albumin (BSA, 5.00 g, 5%, w/v) was dissolved in PBS buffer (100 mL), stored at 4 °C.

**PBST buffer** (500 mL): Tween-20 (250 µL, 0.05%, v/v) was dissolved in PBS buffer (500 mL), stored at 4 °C.

**Antibody buffer** (20 mL): BSA (0.20 g, 1%, v/v) was dissolved in PBS buffer (20 mL), the resulting solution was mixed with monoclonal anti-polyhistidine-alkaline phosphatase antibody (10 µL, 0.05%, v/v), the freshly prepared solution was used for incubation.

iv. <u>*Buffers for HEL modification*</u>

**Phosphate-buffered Saline/PBS buffer** (pH = 7.4).

**Sodium Borate (SB) buffer** (1 L, pH = 8.5): $H_3BO_3$ (200 mM, 12.37 g), NaOH (200 mM, 8.00 g), and distilled water (to 1 L). Mixed, adjusted the pH with $H_3BO_3$ solution, filtered with 0.2 µm filter, stored at room temperature for use.

#### 5.4. Amplification, expression, and purification of *NmCSS* and *PmST3*

##### *5.4.1. Amplification of plasmids*

i. <u>*Transformation of XL10-gold competent cells*</u>

XL10-gold competent cells were transferred with pET24a-*NmCSS* or pET24a-*PmST3* according to the manufacturers' protocol (Instruction Manual, *QuikChange II XL Site-Directed Mutagenesis Kit*)*, transformed cells were plated onto LB agar plates containing ampicillin (100 µg/mL), plates were therefore incubated at 37 °C for 16 h.

*NZY$^+$ broth was replaced with SOC Outgrowth medium in transformation.

ii. <u>*Plasmid amplification and purification*</u>

Single colonies from corresponding plates were inoculated in LB Broth medium (10 mL × 3) and incubated at 37 °C for 16 h. Overnight cultures were centrifuged, desired plasmids were extracted and purified using standard purification procedure (Quick-StartProtocol, *QIAprep® Spin Miniprep Kit*, QIAGEN®).

iii. <u>*Sequencing*</u>

Extracted plasmids were then submitted for DNA sequencing, 500 ng (100 ng/μL, 5 μL) of plasmid was required for single sequencing operation. Designed primers (3.2 μM, 5 μL) were submitted as well if necessary.

*Expression of NmCSS and PmST3*

i. <u>*Transformation into BL21(DE3)*</u>

*E. coli* BL21(DE3) cells was transferred with pET23a-*NmCSS* or pET23a-*PmST3* according to the manufacturers' protocol (*Transformation Protocol for BL21(DE3) Competent Cells*, *C2527*), transformed cells were plated onto LB agar plates containing ampicillin (100 μg/mL) and incubated at 37 °C for 16 h.

ii. <u>*Culture and expression*</u>

Resuspended a single colony in a small LB medium (10 mL, with ampicillin 100 μg/mL), incubated overnight at 37 °C (200 rpm), Inoculated LB medium (1 L, with ampicillin 100 μg/mL) with the fresh overnight culture, incubated at 37 °C (180~200 rpm) until $OD_{600}$ reached 0.6 − 0.8, it took 3.5 h typically. Protein expression was induced by the addition of the stock solution of IPTG (1 mL/L, 0.5 M) and grown at 37 °C for a further 4 h. Cells were harvested by centrifugation at 9,000 rpm in the JLA-9.1000 Beckman rotor for 20 min at 4 °C. Pellets were stored at − 78 °C for purification.

*Lysis of cells*

The frozen cell pellets were resuspended in binding buffer, cell suspensions were sonicated on ice, this consisted of 5 × 15 amplitude micron bursts of 30 s separated by 59 s intervals (40% *Ampl.*). The lysed cells were centrifuged at 25,000 rpm in a JA-30.50 Beckman rotor for 30 min at 4 °C. The supernatant was filtered through a 0.45 μm filter and stored on ice prior to purification.

*Nickel affinity chromatography by NisPur$^{TM}$ Ni-NTA resin (manual operatioon)*

Ni-resin (typically 1 mL for 1 L of the culture) was washed with M.Q. water (×3), the dried resin was added into supernatants followed an incubation at 4 °C overnight. The suspension was loaded into a small column tube, washed with the chilled binding buffer (20 column volumes), eluted with the chilled elution buffer (20 column volumes), all the fractions were collected on ice and analysed by SDS-PAGE (**Figure I-S1**). Fractions containing the desired band were pooled, concentrated using Vivaspin (20 mL) concentrator (10,000 MWCO PES), the protein was further desalted into PBS buffer (pH = 7.4) by PD-10 column by following the manufacturers' protocol.

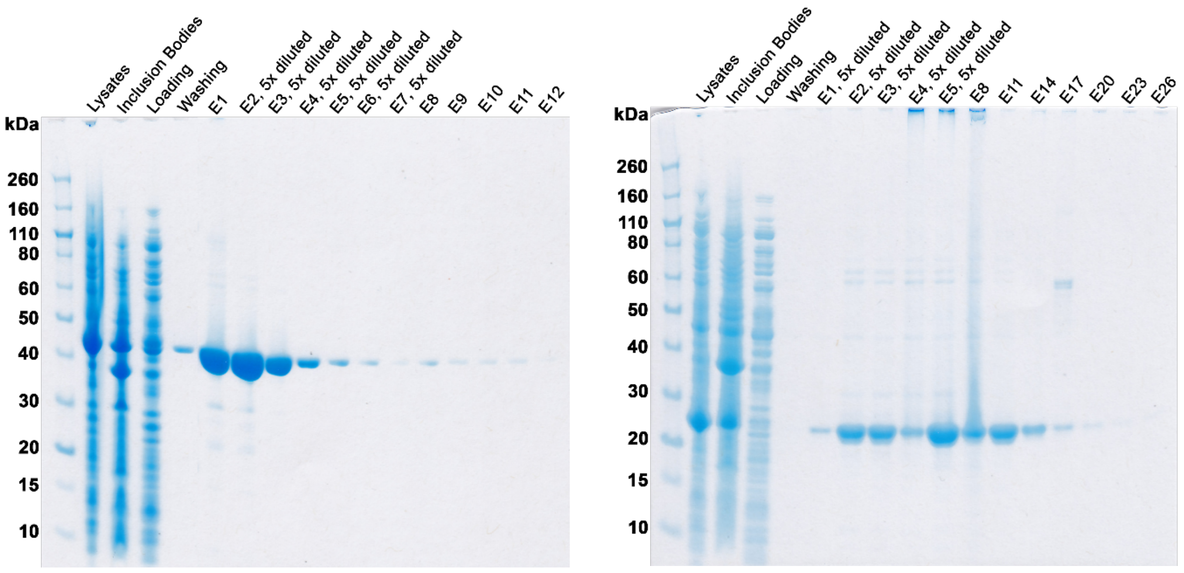


**Figure I-S1.** SDS-PAGE of fractions from manual purification. *NmCSS* (R) and *PmST3* (L).

*Determination of protein concentration using BCA assay*

Protein concentration was determined using BCA Assay by following the protocol listed in *Pierce™ BCA Protein Assay Kit*:

i. *Preparation of diluted albumin (BSA) standards*[a]

**Table I-S1.** Concentrations of BSA solutions for standard curve.

| | **BSA Concentration (mg/ml)** | **Volume of Stock (µl)**[b] | **Volume of Buffer (µl)**[c] | **Final Volume (µl)** |
|---|---|---|---|---|
| **Blank** | 0 | 0 | 1000 | 1000 |
| **1** | 0.025 | 12.5 | 987.5 | 1000 |
| **2** | 0.050 | 25 | 975 | 1000 |
| **3** | 0.100 | 50 | 950 | 1000 |
| **4** | 0.150 | 75 | 925 | 1000 |
| **5** | 0.200 | 100 | 900 | 1000 |
| **6** | 0.300 | 150 | 850 | 1000 |
| **7** | 0.400 | 200 | 800 | 1000 |
| **8** | 0.500 | 250 | 750 | 1000 |
| **9** | 0.600 | 300 | 700 | 1000 |
| **10** | 0.800 | 400 | 600 | 1000 |

[a] BSA standards and stock solution were stored at − 20 °C for use; [b] the concentration of BSA in stock solution is 2 mg/mL; [c] PBS buffer (pH = 7.4) was used for solution preparation.

ii. *Preparation of the BCA working reagent (WR)*

WR was prepared by mixing 50 parts of BCA reagent A with 1 part of BCA reagent B (ratio of A to B is 50:1, v/v).

iii. *Microplate procedure* (sample to WR ratio = 1:8)*

Pipetted 25 μL of each standard and protein solution into a microplate well (96-Well Plate, F-Bottom); added 200 μL of the WR to each well and mixed plate thoroughly; covered plate followed an incubation at 37 °C for 30 min; cooled plate to room temperature; measured the absorbance at 562nm on a plate reader.

*If the protein is concentrated, dilution was required prior to operation; fresh BSA standard curve is needed every time; the working range is 20 μg/mL to 2000 μg/mL.

*Protein characterization*

i. *SDS-PAGE analysis*

Protein fractions were mixed with SDS-PAGE sample loading buffer, the mixture was heated for 10 min at 95 °C, centrifuged, and resulting supernatants were ready for loading. SDS-PAGE was developed in MES buffer (~500 mL, 1×) for 100 min under 110 V at room temperature. The developed gel was stained in InstantBlue™ protein stain (~20 mL) overnight and destained in distilled water for 2 days.

ii. *Western Blotting analysis*

Freshly developed SDS-PAGE was prewashed with M.Q. water. Following the manufacturers' protocol, bands on Gel were transferred onto Membrane (Nitrocellulose, PVDF) using *i*Blot™ Gel Transfer Device ($t$ = 10 min). The transferred membrane was blocked with block buffer (20 mL) for 1 h at room temperature; the blocked membrane was then washed with PBST buffer (20 mL × 3, 5 min × 3); the washed membrane was incubated with antibody buffer (20 mL) overnight at 4 °C followed by washing with PBST buffer (20 mL × 3, 5 min × 3) and PBS buffer (20 mL, 5 min), successively. Finally, washed membrane was stained with BCIP®/NBT liquid substrate system (3 mL) and stopped by distilled water.

iii. *LC_Mass analysis on Waters Xevo G2-S (ESI-qTOF-MS) with UPLC system*

Purified protein (in PBS buffer) was diluted with ammonia acetate buffer, a freshly prepared DTT solution (100 mM in M.Q. water) was added, the final concentration of DTT was 10 mM. The optimal concentration for analysis was 0.03 mg/mL, 5 μL was injected.

Proswift$^{TM}$ RP-2H column (4.6×50 mm SS, ThermoFisher SCIENTIFIC) was run at 0.3 mL/min with eluent A (0.1% formic acid in M.Q. water, v/v) and eluent B (0.1% formic acid in acetonitrile, v/v). Method was programmed as follows (10 min): 5% B (1 min), 5% B to 95% B (6 min), 95% B (1 min), 95% B to 5% B (0.1 min), 5% B (1.9 min). The following MS parameters were used: capillary voltage, 3000 V; sample cone, 20 V; desolvation temperature, 200 °C; source temperature, 80 °C; nitrogen desolvation flow, 700 L/h; no cone flow; pusher cycle time, 94; and ion energy, 34 V; *m*/*z* scan range 200 to 2100; scan time, 1 s; interscan time, 0.1 s; The intact protein LC_MS data were analysed using MassLynx (Waters, version 4.1).

#### HEL Mutants expression in FreeStyle$^{TM}$ 293F cells

1 Tube of 293F cells (P10,1ml) is requested from Pathology CellBank, thaw them into 25ml freestyle medium in the small 125ml shaker flask, wait 3-4 days before doing a count and expanding up.

Expand 293F cells up to 1L volume (seeding at 0.5x10^6 cells/ml each each passage), the day before transfection, diluted healthy cells to 0.7x10^6 cells/ml, on the day of transfection, count cells, make sure viability is over 90% and diluted to 1.0x10^6 cells/ml.

Transfection: cDNA 312 ug for 1L

PEI Mix in tube A: OptiMem 25ml + PEI(1mg/ml) 938 ul

cDNA Mix in tube B: OptiMem 25ml + cDNA solution (sterilized by 0.2uM filter!)

Pour cDNA Mix into PEI Mix (Not the other way round) and mix it gently, incubate at RT for 30 min, then add to cells.

Incubate for 7 days (37C, 8% $CO_2$, 120rpm).

1000 g x 10min at RT pellet cells, supernatant is sterilized by 0.2uM filter unit, stored in cold room.

#### Affinity chromatography

*Protein G column*

4ml Protein G beads are loaded into clean Ecomo-Column chromatography Column (1.5 x 10 cm), pre-equilibrated with 30ml PBS, supernatant is loaded in cold room.

Prewashing: 30ml PBS

Elution: 0.1M glycine, pH=3.0 (~25ml), the eluted solution is immediately neutralized with 1M Tris (pH = 9.0, 150μl per 5ml elute)

Postwashing1: 0.1M glycine, pH=3.0 (10ml)

Postwashing2: 30ml PBS

Postwashing3: PBS containing 0.05% $NaN_3$.

Column is sealed (top and bottom), stored at fridge for reuse.

Protein desalted/dialysized against PBS, concentrated, sterilized, quantified by BCA assay.

*HyHEL-9 column*

HyHEL-9 column is pre-equilibrated with 30ml PBS, HEL supernatant is loaded in cold room.

Prewashing: 30ml PBS

Elution: 3M $MgCl_2$ (10mM Tris, pH=8.0, ~25ml)

Postwashing1: 3M $MgCl_2$ (10mM Tris, pH=8.0, ~10ml)

Postwashing2: 30ml PBS

Postwashing3: PBS containing 0.05% $NaN_3$.

Column is sealed (top and bottom), stored at fridge for reuse.

Protein desalted/dialysized against PBS, concentrated at 4 °C, sterilized, quantified by BCA assay.

*D1.3 column*

D1.3 column is pre-equilibrated with 30ml PBS, HEL supernatant is loaded in cold room.

Prewashing: 30ml PBS

Elution: 3M $MgCl_2$ (10mM Tris, pH=8.0, ~25ml)

Postwashing1: 3M $MgCl_2$ (10mM Tris, pH=8.0, ~10ml)

Postwashing2: 30ml PBS

Postwashing3: PBS containing 0.05% $NaN_3$.

Column is sealed (top and bottom), stored at fridge for reuse.

Protein desalted/dialysized against PBS, concentrated at 4 °C, sterilized, quantified by BCA assay.

#### Reagent synthesis and characterization data

**Cyanomethyl (5-acetamido-3,5-dideoxy-D-*glycero*-α-D-*galacto*-non-2-ulopyranosylonic acid)-(2→3)-β-D-galactopyranosyl-(1→4)-1-thio-β-D-glucopyranoside (1):**

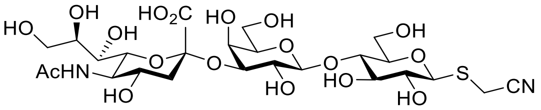


Substrate **6** (200 mg, 503.3 μmol, the final concentration is 10 mM) was dissolved in fresh ammonium carbonate (50.33 mL, 100 mM, containing 20 mM $MgCl_2$, pH = 8.5) buffer in conical flask, *N*-acetylneuraminic acid (163.4 mg, 528.4 μmol), cytidine-5'-triphosphate disodium salt (663.1 mg, 1.258 mmol), CMP-sialic acid synthetase (42.34 μL, 2.5 μg per mg substrate, 11.81 mg/mL in PBS buffer, *NmCSS*), and 2,3-sialyltransferase (41.48 μL, 3.0 μg per mg substrate, 14.465 mg/mL in PBS buffer, *PmST3*) were added, the resulting mixture was incubated at 37 C/200rpm. After 2 to 3 h, the reaction was quenched by adding equal volume of cold ethanol (200 mL). The proteins and insoluble precipitates were removed by centrifugation, the supernatant was concentrated in vacuum, the crude residue was purified by flash column chromatography ($H_2O$–*i*PrOH–EtOAc, 1:2:3 to 1:2:2) followed by size exclusion chromatography (LH20, $CH_3OH$–$H_2O$, 1:1), the combined fractions were concentrated, lyophilized in water, yielding sialyllactose **1** (331 mg, 96%) as a white powder: $R_f$ = 0.61 ($H_2O$–*i*PrOH–EtOAc, 1:2:2); mp 169 – 170 °C; $[\alpha]_D^{25}$ − 19.1 (*c* 1.00, $H_2O$); FT-IR (film): $\nu_{max}$ = 3017, 2349, 1612, 1108, 1030, 618 $cm^{-1}$; $^1H$ NMR (700 MHz, $D_2O$): $\delta$ 4.77 (d, $J_{1,2}$ = 10.0 Hz, 1H, H-1), 4.55 (d, $J_{1',2'}$ = 7.8 Hz, 1H, H-1′), 4.12 (dd, $J_{2',3'}$ = 9.9 Hz, $J_{3',4'}$ = 3.2 Hz, 1H, H-3′), 4.01 (dd, $J_{5',6'a}$ = 2.2 Hz, $J_{6'a,6'b}$ = 12.5 Hz, 1H, H-6′a), 3.97 (app d, *J* = 3.1 Hz, 1H, H-4′), 3.90 (ddd, $J_{8'',7''}$ = 8.9 Hz, $J_{8'',9''a}$ = 6.2 Hz, $J_{8'',9''b}$ = 2.5 Hz, 1H, H-8″), 3.89−3.84 (m, 3H, H-9″a, H-5″, H-6b), 3.82 (d, $^2J$ = 17.5 Hz, 1H, $SCH_2CN$), 3.79−3.65 (m, 9H, H-6′a, H-4, H-6′b, H-5′, $SCH_2CN$, H-4″, H-3, H-5, H-9″b), 3.64 (dd, $J_{6'',5''}$ = 9.0 Hz, $J_{6'',7''}$ = 2.0 Hz, 1H, H-6″), 3.60 (dd, $J_{7'',6''}$ = 1.8 Hz, $J_{7'',8''}$ = 8.9 Hz, 1H, H-7″), 3.59 (dd, $J_{2',1'}$ = 7.9 Hz, $J_{2',3'}$ = 9.9 Hz, 1H, H-2′), 3.49 (dd, $J_{2,1}$ = 9.8 Hz, $J_{2,3}$ = 9.1 Hz, 1H, H-2), 2.77 (dd, $J_{3''eq,3''ax}$ = 12.5 Hz, $J_{3''eq,4''}$ = 4.7 Hz, 1H, H-3″eq), 2.04 (s, 3H, $CH_3CONH$), 1.81 (t, $J_{3''ax,3''eq}$ = 12.3 Hz, $J_{3''ax,4''}$ = 12.3 Hz, 1H, H-3″ax) *ppm*; $^{13}C$ NMR (175 MHz, $D_2O$) $\delta$ 175.0 ($CH_3CONH$), 173.9 ($J_{C1'',H3''ax}$ = 4.7 Hz, *C*OOH, *C*-1″), 118.6 ($SCH_2CN$), 102.6 (*C*-1′), 99.8 (*C*-2″), 84.4 (*C*-1), 78.9 (*C*-5), 77.7 (*C*-4), 75.6 (*C*-3), 75.5 (*C*-3′), 75.2 (*C*-5′), 72.9 (*C*-6″), 71.8 (*C*-8″), 71.7 (*C*-2), 69.4 (*C*-2′), 68.3 (*C*-4″), 68.1 (*C*-7″), 67.5 (*C*-4′), 62.6 (*C*-9″), 61.0 (*C*-6′), 60.0 (*C*-6), 51.7 (*C*-5″), 39.6 (*C*-3″),

22.0 ($CH_3$CONH), 14.5 (S$CH_2$CN) *ppm*; HRMS (ESI): *m*/*z* calcd for $C_{25}H_{40}N_2NaO_{18}S$ $[M+Na]^+$ 711.1889. Found: 711.1887.

**2,3,4,6-Tetra-*O*-acetyl-β-D-galactopyranosyl-(1→4)-1,2,3,6-tetra-*O*-acetyl-D-glucopyranose (2):**

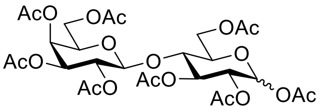


A suspension of sodium acetate (7.5 g) in acetic anhydride (270 mL) was heated to reflux, D-lactose (30 g) was then added into the mixture portion-wise without heating. After complete addition, the mixture was stirred under reflux for another 3 h, giving a clear solution. The hot solution was then poured into a mixture of ice and water (500 mL) under vigorous stirring. The resulted suspension was then extracted with $CH_2Cl_2$ (200 mL × 1, 50 mL × 3), the combined organic layers were washed with saturated $NaHCO_3$ (400 mL, aq.) solution and brine (500 mL) successively, dried over $Na_2SO_4$, filtered and concentrated. The crude solid was recrystallized from hot ethanol, yielding the desired lactose octaacetate **2** (54 g, 96%, α:β=1:9) as an amorphous white solid: $R_f$ = 0.31 (PE–EtOAc, 1:1); mp 90 – 92 °C; $[\alpha]_D^{25}$ + 7.5 (*c* 1.00, $CH_2Cl_2$); FT-IR (film): $\nu_{max}$ = 2361, 1748, 1434, 1369, 1217, 1172, 1049, 954, 900, 669 $cm^{-1}$; $^1H$ NMR (700 MHz, $CDCl_3$): $\delta$ 6.24 (d, $J_{1,2}$ = 3.7 Hz, 1H, $^{\alpha}$H-1), 5.66 (d, $J_{1,2}$ = 8.3 Hz, 1H, $^{\beta}$H-1), 5.45 (t, $J_{2,3}$ = 9.7 Hz, $J_{3,4}$ = 9.7 Hz, 1H, $^{\alpha}$H-3), 5.35 (app d, $J_{3',4'}$ = 3.3 Hz, 1H, $^{\alpha}$H-4′), 5.34 (app d, $J_{3',4'}$ = 3.3 Hz, 1H, $^{\beta}$H-4′), 5.23 (t, $J_{2,3}$ = 9.2 Hz, $J_{3,4}$ = 9.2 Hz, 1H, $^{\beta}$H-3), 5.12–5.08 (m, $^{\alpha}$H-2′, $^{\beta}$H-2′), 5.03 (t, $J_{1,2}$ = 8.7 Hz, $J_{2,3}$ = 8.7 Hz, 1H, $^{\beta}$H-2), 5.00 (dd, $J_{1,2}$ = 3.7 Hz, $J_{2,3}$ = 10.3 Hz, 1H, $^{\alpha}$H-2), 4.96–4.93 (m, $^{\alpha}$H-3′, $^{\beta}$H-3′), 4.48–4.43 (m, $^{\alpha}$H-1′, $^{\beta}$H-1′, $^{\beta}$H-6a, $^{\alpha}$H-6a), 4.15–4.05 (m, $^{\alpha}$H-6b, $^{\beta}$H-6b, $^{\beta}$H-6′a, $^{\alpha}$H-6′a, $^{\alpha}$H-6′b, $^{\beta}$H-6′b), 3.99 (ddd, $J_{4,5}$ = 10.0 Hz, $J_{5,6a}$ = 1.7 Hz, $J_{5,6b}$ = 3.8 Hz, 1H, $^{\alpha}$H-5), 3.88–3.79 (m, $^{\alpha}$H-4, $^{\beta}$H-4, $^{\beta}$H-5′, $^{\alpha}$H-5′), 3.75 (ddd, $J_{4,5}$ = 9.8 Hz, $J_{5,6a}$ = 1.8 Hz, $J_{5,6b}$ = 4.8 Hz, 1H, $^{\beta}$H-5), 2.17 (s, 3H, $^{\alpha}$C*H*$_3$CO), 2.15 (s, 3H, $^{\alpha}$C*H*$_3$CO), 2.14 (s, 3H, $^{\beta}$C*H*$_3$CO), 2.12 (s, 3H, $^{\alpha}$C*H*$_3$CO), 2.11 (s, 3H, $^{\beta}$C*H*$_3$CO), 2.09 (s, 3H, $^{\beta}$C*H*$_3$CO), 2.06 (s, $^{\beta}$C*H*$_3$CO, $^{\alpha}$C*H*$_3$CO), 2.05 (s, 3H, $^{\alpha}$C*H*$_3$CO), 2.044 (s, 3H, $^{\alpha}$C*H*$_3$CO), 2.040 (s, 3H, $^{\beta}$C*H*$_3$CO), 2.03 (s, 3H, $^{\beta}$C*H*$_3$CO), 2.02 (s, 3H, $^{\beta}$C*H*$_3$CO), 2.00 (s, 3H, $^{\alpha}$C*H*$_3$CO), 1.96 (s, 3H, $^{\alpha}$C*H*$_3$CO), 1.95 (s, 3H, $^{\beta}$C*H*$_3$CO) *ppm*. Identical to the previous report (*20*).

**2,3,4,6-Tetra-*O*-acetyl-β-D-galactopyranosyl-(1→4)-2,3,6-tri-*O*-acetyl-α-D-glucopyranosyl bromide (3):**

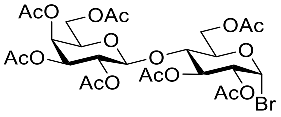

To a stirred solution of lactose octaacetate **2** (3.6 g, 5.3 mmol) in dry $CH_2Cl_2$ (25 mL) was added hydrogen bromide (7 mL, 33 wt.% in acetic acid) solution dropwise at 0 °C, the reaction solution was allowed to warm to room temperature and was stirred for 1 h. The brown solution was diluted with $CH_2Cl_2$ (100 mL), washed with cold saturated $NaHCO_3$ (200 mL) solution, the organic layer was separated, the aqueous layer was then extracted with $CH_2Cl_2$ (30 mL × 2), the combined organic layer was washed with brine (200 mL), dried over $Na_2SO_4$ and filtered, the solution was concentrated to almost dry followed by an immediate addition of cold ether (100 mL, 0 °C), giving white precipitate. The solid was collected by filtration and dried in vacuum to yield the bromosugar **3** (3.11 g, 84%) as an amorphous white solid: $R_f$ = 0.44 (PE–EtOAc, 1:1); $[\alpha]_D^{25}$ + 69.7 (*c* 0.60, $CH_2Cl_2$); FT-IR (film): $\nu_{max}$ = 2924, 2360, 1746, 1434, 1369, 1215, 1174, 1111, 1050, 955, 903, 736, 648 $cm^{-1}$; $^1H$ NMR (700 MHz, $CDCl_3$): $\delta$ 6.53 (d, $J_{1,2}$ = 4.1 Hz, 1H, H-1), 5.56 (t, $J_{2,3}$ = 9.6 Hz, $J_{3,4}$ = 9.6 Hz, 1H, H-3), 5.36 (app d, $J_{3',4'}$ = 3.3 Hz, 1H, H-4′), 5.13 (dd, $J_{1',2'}$ = 8.0 Hz, $J_{2',3'}$ = 10.4 Hz, 1H, H-2′), 4.96 (dd, $J_{2',3'}$ = 10.4 Hz, $J_{3',4'}$ = 3.4 Hz, 1H, H-3′), 4.76 (dd, $J_{1,2}$ = 4.1 Hz, $J_{2,3}$ = 9.6 Hz, 1H, H-2), 4.52 (d, $J_{1',2'}$ = 7.6 Hz, 1H, H-1′), 4.50 (app d, $J_{6a,6b}$ = 9.8 Hz, 1H, H-6a), 4.22−4.18 (m, 2H, H-5, H-6b), 4.15 (dd, $J_{5',6a'}$ = 6.4 Hz, $J_{6a',6b'}$ = 11.2 Hz, 1H, H-6′a), 4.08 (dd, $J_{5',6b'}$ = 7.3 Hz, $J_{6a',6b'}$ = 11.2 Hz, 1H, H-6′b), 3.90 (app t, $J_{5',6a'}$ = 6.9 Hz, $J_{5',6b'}$ = 6.9 Hz, 1H, H-5′), 3.86 (t, $J_{3,4}$ = 9.6 Hz, $J_{4,5}$ = 9.6 Hz, 1H, H-4), 2.16 (s, 3H, C$H_3$CO), 2.14 (s, 3H, C$H_3$CO), 2.10 (s, 3H, C$H_3$CO), 2.07 (s, 3H, C$H_3$CO), 2.065 (s, 3H, C$H_3$CO), 2.057 (s, 3H, C$H_3$CO), 1.97 (s, 3H, C$H_3$CO) *ppm*. Identical to the previous report (*21*).

**2,3,4,6-Tetra-*O*-acetyl-β-D-galactopyranosyl-(1→4)-2,3,6-tri-*O*-acetyl-1-*S*-acetyl-1-thio-β-D-glucopyranose (4):**

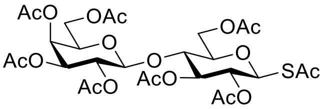

To a stirred solution of bromosugar **3** (1.39 g, 1.99 mmol) in dry $CH_3CN$ (10 mL) was added potassium thioacetate (430 mg, 2.98 mmol) at room temperature. After stirring for 3 h, the reaction mixture was diluted with $CH_2Cl_2$ (100 mL), washed with saturated $NaHCO_3$ (100 mL) solution, the organic layer was separated, the aqueous layer was then extracted with $CH_2Cl_2$ (20 mL × 2), the combined organic layer was washed with brine (150 mL), dried over $Na_2SO_4$, filtered and concentrated, the crude residue was purified by flash column chromatography (PE–EtOAc, 1:1) to give the desired thioester **4** (1.27 g, 92%) as a white foam: $R_f$ = 0.35 (PE–EtOAc, 1:1); mp 79 – 80 °C; $[\alpha]_D^{25}$ + 4.4 (*c* 0.70, $CH_2Cl_2$); FT-IR (film): $\nu_{max}$ = 2981, 2360, 2341, 1750, 1713, 1434, 1370, 1222, 1170, 1130, 1055, 954, 913, 623 $cm^{-1}$; $^1H$ NMR (400 MHz, $CDCl_3$): $\delta$ 5.35 (dd, $J_{3',4'}$ = 3.4 Hz, $J_{4',5'}$ = 0.9 Hz, 1H, H-4′), 4.26 (t, $J_{2,3}$ = 9.0 Hz, $J_{3,4}$ = 9.0 Hz, 1H, H-3), 5.21 (d, $J_{1,2}$ = 10.4 Hz, 1H, H-1), 5.11 (dd, $J_{1',2'}$ = 7.9 Hz, $J_{2',3'}$ = 10.4 Hz, 1H, H-2′), 5.04 (dd, $J_{1,2}$ = 10.4 Hz, $J_{2,3}$ = 9.2 Hz, 1H, H-2), 4.94 (dd, $J_{2',3'}$ = 10.4 Hz, $J_{3',4'}$ = 3.2 Hz, 1H, H-3′), 4.46 (d, $J_{1',2'}$ = 7.9 Hz, 1H, H-1′), 4.45 (dd, $J_{5',6a'}$ = 1.8 Hz, $J_{6'a,6b'}$ = 12.1 Hz, 1H, H-6′a), 4.15−4.05 (m, 3H, H-6a, H-6b, H-6′b), 3.86 (app dt, $J_{4',5'}$ = 0.9 Hz, $J_{5',6b'}$ = 7.0 Hz, 1H, H-5′), 3.82 (dd, $J_{3,4}$ = 8.8 Hz, $J_{4,5}$ = 10.0 Hz, 1H, H-4), 3.75 (ddd, $J_{4,5}$ = 10.0 Hz, $J_{5,6a}$ = 4.6 Hz, $J_{5,6b}$ = 1.8 Hz, 1H, H-5), 2.37 (s, 3H, C*H₃*COS), 2.15 (s, 3H, C$H_3$CO), 2.11 (s, 3H, C$H_3$CO), 2.07 (s, 3H, C$H_3$CO), 2.05 (s, 3H, C$H_3$CO), 2.04 (s, 3H, C$H_3$CO), 2.02 (s, 3H, C$H_3$CO), 1.96 (s, 3H, C$H_3$CO) *ppm*. Identical to the previous report (*22*).

**Cyanomethyl 2,3,4,6-tetra-*O*-acetyl-β-D-galactopyranosyl-(1→4)-2,3,6-tri-*O*-acetyl-1-thio-β-D-glucopyranoside (5):**

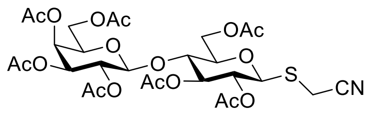


To a stirred solution of thioester **4** (456 mg, 656 μmol) in dry $CH_3CN$ (5 mL) were added hydrazine acetate (72.6 mg, 788 μmol), triethylamine (183 μL, 1.31 mmol), and chloroacetonitrile (831 μL, 13.13 mmol) at room temperature. After stirring for 3 h, the reaction mixture was diluted with $CH_2Cl_2$ (100 mL), washed with saturated $NaHCO_3$ (100 mL, aq.) solution, the organic layer was separated, the aqueous layer was then extracted with $CH_2Cl_2$ (30 mL × 2), the combined organic layer was washed with brine (200 mL), dried over $Na_2SO_4$, filtered and concentrated, the crude residue was purified by flash column chromatography (PE–EtOAc, 1:1) to give the desired **5** (429 mg, 94%) as a white foam: $R_f$ = 0.30 (PE–EtOAc, 1:1); mp 70 – 71 °C; $[\alpha]_D^{25}$ − 33.2 (*c* 0.65, $CH_2Cl_2$); FT-IR (film): $\nu_{max}$ = 1746, 1434, 1370, 1220, 1170, 1137, 1048, 913, 736 $cm^{-1}$; $^1H$ NMR (700 MHz, $CDCl_3$): $\delta$ 5.35 (app d, $J_{3',4'}$ = 3.4 Hz, 1H, H-4′), 5.25 (t, $J_{2,3}$ = 9.2 Hz, $J_{3,4}$ = 9.2 Hz, 1H, H-3), 5.10 (dd, $J_{1',2'}$ = 9.0 Hz, $J_{2',3'}$ = 10.3 Hz, 1H, H-2′), 4.98 (t, $J_{1,2}$ = 9.7 Hz, $J_{2,3}$ = 9.7 Hz, 1H, H-2), 4.96 (dd, $J_{2',3'}$ = 10.6 Hz, $J_{3',4'}$ = 3.4 Hz, 1H, H-3′), 4.69 (d, $J_{1,2}$ = 10.0 Hz, 1H, H-1), 4.55 (dd, $J_{5,6a}$ = 1.5 Hz, $J_{6a,6b}$ = 12.1 Hz, 1H, H-6a), 4.49 (d, $J_{1',2'}$ = 8.0 Hz, 1H, H-1′), 4.14−4.07 (m, 3H, H-6′a, H-6b, H-6′b), 3.87 (app t, $J_{5',6'a}$ = 6.7 Hz, $J_{5',6'b}$ = 6.7 Hz, 1H, H-5′), 3.82 (t, $J_{3,4}$ = 9.7 Hz, $J_{4,5}$ = 9.7 Hz, 1H, H-4), 3.68 (ddd, $J_{4,5}$ = 9.7 Hz, $J_{5,6a}$ = 1.5 Hz, $J_{5,6b}$ = 4.8 Hz, 1H, H-5), 3.59 (d, *J* = 17.0 Hz, 1H, SC*H*HCN), 3.29 (d, *J* = 17.0 Hz, 1H, SCH*H*CN), 2.15 (s, 3H, C$H_3$CO),

2.13 (s, 3H, C$H_3$CO), 2.06 (s, 6H, C$H_3$CO × 2), 2.05 (s, 3H, C$H_3$CO), 2.04 (s, 3H, C$H_3$CO), 1.96 (s, 3H, C$H_3$CO) *ppm*; $^{13}$C NMR (175 MHz, $CDCl_3$) $\delta$ 170.48 ($CH_3$*C*O), 170.47 ($CH_3$*C*O), 170.25 ($CH_3$*C*O), 170.19 ($CH_3$*C*O), 169.9 ($CH_3$*C*O), 169.7 ($CH_3$*C*O), 169.1 ($CH_3$*C*O), 115.8 ($SCH_2$*C*N), 101.2 (*C*-1′), 81.6 (*C*-1), 77.4 (*C*-5), 75.9 (*C*-4), 73.4 (*C*-3), 71.1 (*C*-3′), 70.9 (*C*-5′), 70.0 (*C*-2), 69.2 (*C*-2′), 66.7 (*C*-4′), 61.8 (*C*-6), 61.0 (*C*-6′), 21.0 (*C*$H_3$CO), 20.9 (*C*$H_3$CO), 20.78 (*C*$H_3$CO), 20.77 (*C*$H_3$CO), 20.76 (*C*$H_3$CO), 20.7 (*C*$H_3$CO), 20.6 (*C*$H_3$CO), 14.6 (S*C*$H_2$CN) *ppm*; HRMS (ESI): *m*/*z* calcd for $C_{28}H_{37}NNaO_{17}S$ $[M+Na]^+$ 714.1674. Found: 714.1668.

**Cyanomethyl β-D-galactopyranosyl-(1→4)-1-thio-β-D-glucopyranoside (6):**

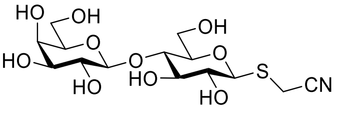


To a stirred solution of **5** (1.99 g, 2.88 mmol) in dry $CH_3OH$ (57.6 mL) was added triethylamine (4.01 mL, 28.77 mmol), the mixture was heated to 40 °C for 24 h. The resulting solution was neutralized by DOWEX 50WX8 (100−200 mesh, hydrogen form) resin, the resin was then removed by filtration, filtrate was concentrated and the crude residue was purified by flash column chromatography ($H_2O$–*i*PrOH–EtOAc, 1:2:4) followed by lyophilisation in water, giving the desired **6** (872 mg, 72%) as a white powder: $R_f$ = 0.40 ($H_2O$–*i*PrOH–EtOAc, 1:2:4); mp 105 – 106 °C; $[\alpha]_D^{25}$ – 65.7 (*c* 0.30, $CH_3OH$); FT-IR (film): $\nu_{max}$ = 3366, 2920, 2361, 2342, 1653, 1398, 1074, 889, 669 cm$^{-1}$; $^1$H NMR (400 MHz, $D_2O$): $\delta$ 4.77 (d, $J_{1,2}$ = 9.0 Hz, 1H, H-1), 4.49 (d, $J_{1',2'}$ = 7.8 Hz, 1H, H-1′), 4.01 (dd, $J_{5',6'a}$ = 2.2 Hz, $J_{6'a,6'b}$ = 12.5 Hz, 1H, H-6′a), 3.95 (app d, $J_{3',4'}$ = 3.4 Hz, 1H, H-4′), 3.86−3.68 (m, 10H, H-6′b, SC*H*HCN, H-6a, H-6b, H-4, H-5′, SCH*H*CN, H-3, H-5, H-3′), 3.57 (dd, $J_{1',2'}$ = 7.8 Hz, $J_{2',3'}$ = 9.9 Hz, 1H, H-2′), 3.50 (dd, $J_{1,2}$ = 9.1 Hz, $J_{2,3}$ = 9.8 Hz, 1H, H-2) *ppm*; $^{13}$C NMR (100 MHz, $D_2O$) $\delta$ 118.6 ($SCH_2$*C*N), 102.8 (*C*-1′), 84.3 (*C*-1), 78.8 (*C*-3′), 77.8 (*C*-5′), 75.6 (*C*-3), 75.3 (*C*-4), 72.5 (*C*-5), 71.7 (*C*-2), 70.9 (*C*-2′), 68.5 (*C*-4′), 61.04 (*C*-6), 60.00 (*C*-6′) *ppm*; HRMS (ESI): *m*/*z* calcd for $C_{14}H_{23}NNaO_{10}S$ $[M+Na]^+$ 420.0935. Found: 420.0934. Identical to the previous report (*23*).

**2-Imino-2-methoxyethyl (5-acetamido-3,5-dideoxy-D-*glycero*-α-D-*galacto*-non-2-ulopyranosylonic acid)-(2→3)-β-D-galactopyranosyl-(1→4)-1-thio-β-D-glucopyranoside (7):**

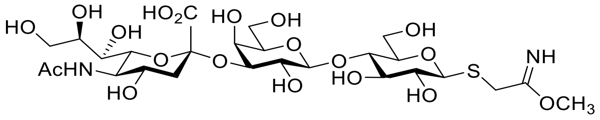

To a stirred solution of trisaccharide **1** (6.89 mg, 10 μmol) in dry $CH_3OH$ (480 μL) was added $CH_3ONa$ solution (20 μL, 0.5 M in dry $CH_3OH$), the resulting mixture was stirred at room temperature for 4 days. The cloudy solution was then concentrated and dried in high vacuum, giving a mixture of substrate **1** (50%) and imidate **7** (50%): $R_f$ = 0.47 ($H_2O$–*i*PrOH–EtOAc, 1:2:2); $^1$H NMR (400 MHz, $CD_3OD$): $\delta$ 4.59 (d, $J_{1,2}$ = 9.7 Hz, $H^1$-1), 4.43 (d, $J_{1',2'}$ = 7.8 Hz, $H^1$-1′), 4.42 (d, $J_{1',2'}$ = 9.7 Hz, $H^7$-1′), 4.35 (d, $J_{1,2}$ = 7.8 Hz, $H^7$-1), 4.06−3.24 (m, 23H), 2.85 (dd, dd, $J_{3''eq,4''}$ = 2.4 Hz, $J_{3''ax,3''eq}$ = 12.2 Hz, 1H, $H^1$-3″eq & $H^7$-3″eq), 2.00 (s, 3H, $CH_3$CONH for **1** and **7**), 1.74−1.69 (m, 1H, $H^1$-3″ax & $H^7$-3″ax) *ppm*; $^{13}$C NMR (100 MHz, $CD_3OD$): $\delta$ 175.5 ($CH_3$*C*ONH), 174.9 (*C*OOH, *C*-1″), 173.0 (*C*(NH)$OCH_3$), 118.6 ($SCH_2$*C*N), 105.0 ($C^1$-1′ & $C^7$-1′), 101.1 ($C^1$-2″ & $C^7$-2″), 86.6 ($C^7$-1), 85.2 ($C^1$-1), 80.8, 80.7, 80.62, 80.58, 77.79, 77.76, 77.6, 77.1, 74.9, 74.1, 73.0, 70.8, 70.1, 69.4, 69.0, 64.6, 62.7, 62.1, 62.0, 53.9, 42.1 ($C^1$-3″ & $C^7$-3″), 22.6 (*C*$H_3$CONH) *ppm*; HRMS (ESI): *m*/*z* calcd for $C_{26}H_{45}O_{19}N_2S$ $[M+H]^+$ 721.2327. Found: 721.2332.

**$N^1$-Methyl-2-[(5-acetamido-3,5-dideoxy-D-*glycero*-α-D-*galacto*-non-2-ulopyranosylonic acid)-(2→3)-β-D-galactopyranosyl-(1→4)-1-β-D-glucopyranosyl)]sulfonylethanimidamide hydrochloride (8):**

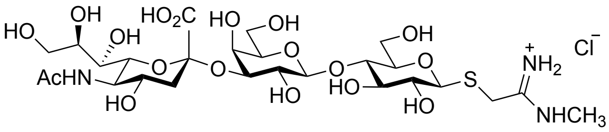

Substrate **10** (51.9 mg, 121 μmol, the final concentration is 10 mM) was dissolved in fresh ammonium carbonate (12.1 mL, 100 mM, containing 20 mM $MgCl_2$, pH = 8.5) buffer in 50mL-Falcon tube, *N*-acetylneuraminic acid (39.3 mg, 127 μmol), cytidine-5’-triphosphate disodium salt (191.3 mg, 363 μmol), CMP-sialic acid synthetase (11 μL, 2.5 μg per mg substrate, 11.81 mg/mL in PBS buffer, *NmCSS*), and 2,3-sialyltransferase (10.8 μL, 3.0 μg per mg substrate, 14.465 mg/mL in PBS buffer, *PmST3*) were added, the resulting mixture was incubated at 37°C/200rpm. After 2 h, the reaction was quenched by adding equal volume of cold ethanol (12.1 mL). The proteins and insoluble precipitates were removed by centrifugation, the supernatant was concentrated in vacuum, the crude residue was purified by flash column chromatography (*i*PrOH–EtOAc–Ammonia(35%)–$H_2O$, 40:40:3:50)

followed by size exclusion chromatography (LH20, $CH_3OH$–$H_2O$, 1:1), the combined fractions were concentrated, lyophilized in water, yielding SiaLac-amidine **8** (35.7 mg, 41%) as a white powder: $R_f$ = 0.32 (*i*PrOH–EtOAc–Ammonia(35%)–$H_2O$, 40:40:3:50); mp 197 – 198 °C; $[\alpha]_D^{25}$ – 10.7 (*c* 0.50, $H_2O$); FT-IR (film): $\nu_{max}$ = 3269, 2360, 1697, 1607, 1260, 1071, 1029, 797 $cm^{-1}$; $^1H$ NMR (400 MHz, $D_2O$) $\delta$ 4.61 (d, $J_{1,2}$ = 9.9 Hz, 1H, H-1), 4.52 (d, $J_{1',2'}$ = 7.8 Hz, 1H, H-1′), 4.10 (dd, $J_{3',2'}$ = 9.8 Hz, $J_{3',4'}$ = 3.1 Hz, 1H, H-3′), 3.95–3.80 (m, 7H), 3.76–3.54 (m, 12H), 3.40 (dd, $J_{2,1}$ = 9,9 Hz, $J_{2,3}$ = 8.8 Hz, 1H, H-2), 2.94 (s, 3H, $SCH_2C(NH)NHC\mathit{H_3}$), 2.75 (dd, $J_{3''eq,3''ax}$ = 12.5 Hz, $J_{3''eq,4}$ = 4.7 Hz, 1H, H-3″eq), 2.02 (s, 3H, $C\mathit{H_3}CONH$), 1.78 (app t, $J_{3''ax,3''eq}$ = 12.1 Hz, $J_{3''ax,4}$ = 12.1 Hz, 1H, H-3″ax) *ppm*; $^{13}C$ NMR (100 MHz, $D_2O$) $\delta$ 175.0 ($CH_3\mathit{C}ONH$), 173.8 (C-1″), 166.1 ($SCH_2\mathit{C}(NH)NHCH_3$), 102.6 (*C*-1′), 99.8 (*C*-2″), 85.0 (*C*-1), 78.7, 77.6, 75.5 (2**C*), 75.2, 72.9, 71.8, 71.7, 69.3, 68.3, 68.1, 67.4, 62.6, 61.0, 60.0, 51.7 (*C*-5″), 39.6 (*C*-3″), 30.6 ($S\mathit{C}H_2C(NH)NHCH_3$), 28.6 ($SCH_2C(NH)NH\mathit{C}H_3$), 22.0 ($\mathit{C}H_3CONH$) *ppm*. HRMS (ESI): *m*/*z* calcd for $C_{26}H_{46}O_{18}N_3S$ $[M+H]^+$ 720.2486. Found: 720.2492.

***N*¹-Methyl-2-[β-D-galactopyranosyl-(1→4)-1-β-D-glucopyranosyl)]sulfonylethanimidamide hydrochloride (10)**:

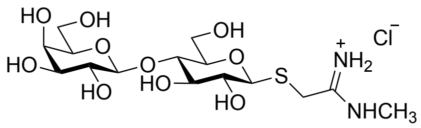


To a stirred solution of disaccharide **6** (100 mg, 252 μmol) in dry $CH_3OH$ (12.1 mL) was added $CH_3ONa$ solution (503 μL, 0.5 M in dry $CH_3OH$), the resulting mixture was stirred at room temperature for 4 days. The white precipitate was collected by filtration, washed with dry $CH_3OH$, dried in high vacuum, affording the imidate intermediate. Imidate **9** was then suspended in dry $CH_3OH$ (5 mL) again, $CH_3NH_2$ solution (503 μL, 2 M in THF) solution was add, after stirring at room temperature for 2 days, the cloudy solution was filtered, the solid was washed with dry $CH_3OH$. Redissolved in HCl solution (1 mL, 1 M in $CH_3OH$) , excess ether was added to precipitate amidine salt. After centrifuging, sugar pellet was harvested, dried in vacuum, giving the corresponding Lac-amidine **10** (59 mg, 55%) as a white powder: $R_f$ = 0.38 (EtOAc–$CH_3OH$–AcOH–$H_2O$, 3:3:3:2); mp 194 – 195 °C; $[\alpha]_D^{25}$ – 24.4 (*c* 0.83, $H_2O$); FT-IR (neat): $\nu_{max}$ = 3386, 1610, 1415, 1371, 1170, 1119, 1078, 1036, 992, 881, 795, 697 $cm^{-1}$; $^1H$ NMR (400 MHz, $D_2O$): $\delta$ 4.60 (d, $J_{1,2}$ = 9.9 Hz, 1H, H-1), 4.44 (d, $J_{1',2'}$ = 7.8 Hz, 1H, H-1′), 3.94−3.90 (m, 2H, H-6a, H-4′), 3.84−3.74 (m, 4H, $SC\mathit{H}HC(NH)NHCH_3$, H-6b, H-6a′,

H-6b′), 3.72−3.68 (m, 2H, H-5′, SCH*H*C(NH)NHCH$_3$), 3.66−3.57 (m, 4H, H-4, H-3′, H-3, H-5), 3.52 (dd, $J_{2',1'}$ = 7.8 Hz, $J_{2',3'}$ = 10.0 Hz, 1H, H-2′), 3.40 (dd, $J_{2,1}$ = 9.9 Hz, $J_{2,3}$ = 8.8 Hz, 1H, H-2), 2.94 (s, 3H, SCHHC(NH)NHC*H$_3$*) ppm; $^{13}$C NMR (100 MHz, D$_2$O): $\delta$ 166.1 (SCH$_2$*C*(NH)NHCH$_3$), 102.8 (*C*-1′), 85.0 (*C*-1), 78.7 (*C*-5), 77.7 (*C*-4), 75.5 (*C*-3′), 75.4 (*C*-5′), 72.5 (*C*-3), 71.7 (*C*-2), 70.9 (*C*-2′), 68.5 (*C*-4′), 61.0 (*C*-6′), 60.0 (*C*-6), 30.6 (S*C*H$_2$C(NH)NHCH$_3$), 28.6 (SCH$_2$C(NH)NH*C*H$_3$) ppm. HRMS (ESI): *m*/*z* calcd for $C_{15}H_{29}O_{10}N_2S$ [M+H]$^+$ 429.1533. Found: 429.1537.

**Carboxymethyl 2,3,4,6-tetra-*O*-acetyl-β-D-galactopyranosyl-(1→4)-2,3,6-tri-*O*-acetyl-1-thio-β-D-glucopyranoside (11):**

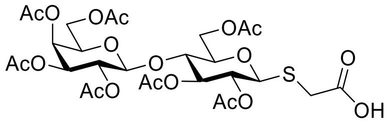


To a stirred solution of thioester **4** (4.574 g, 6.58 mmol) in dry CH$_3$CN (66 mL) were added hydrazine acetate (727.6 mg, 7.90 mmol), triethylamine (3.67 mL, 26.34 mmol), and bromoacetic acid (1.83 g, 13.17 mmol) at room temperature. After stirring for 2 h, the reaction was concentrated, the residue was suspended in CH$_2$Cl$_2$ (150 mL), washed with HCl (200 mL, 1 M, aq.) solution, the organic layer was separated and the aqueous layer was then extracted with CH$_2$Cl$_2$ (50 mL × 2), the combined organic layer was dried over Na$_2$SO$_4$, filtered and concentrated, the residue was purified by flash column chromatography (CH$_2$Cl$_2$–CH$_3$OH, 10:1 to 4:1) to give the desired **11** (4.164 g, 89%) as a white foam: $R_f$ = 0.32 (CH$_2$Cl$_2$–CH$_3$OH, 10:1); mp 96 – 97 °C; $[\alpha]_D^{25}$ − 20.6 (*c* 1.00, CH$_2$Cl$_2$); FT-IR (film): $\nu_{max}$ = 2981, 1745, 1371, 1220, 1140, 1049, 955, 913 cm$^{-1}$; $^1$H NMR (400 MHz, CDCl$_3$): $\delta$ 5.34 (dd, $J_{3',4'}$ = 3.3 Hz, $J_{4',5'}$ = 0.7 Hz, 1H, H-4′), 5.22 (t, $J_{2,3}$ = 9.2 Hz, $J_{3,4}$ = 9.2 Hz, 1H, H-3), 5.09 (dd, $J_{1',2'}$ = 7.8 Hz, $J_{2',3'}$ = 10.4 Hz, 1H, H-2′), 4.97 (t, $J_{1,2}$ = 8.8 Hz, $J_{2,3}$ = 8.8 Hz, 1H, H-2), 4.96 (dd, $J_{2',3'}$ = 10.6 Hz, $J_{3',4'}$ = 3.3 Hz, 1H, H-3′), 4.62 (d, $J_{1,2}$ = 10.4 Hz, 1H, H-1), 4.55 (dd, $J_{5,6a}$ = 1.6 Hz, $J_{6a,6b}$ = 12.1 Hz, 1H, H-6a), 4.49 (d, $J_{1',2'}$ = 7.8 Hz, 1H, H-1′), 4.15−4.04 (m, 3H, H-6′a, H-6′b, H-6b), 3.88 (dt, $J_{4',5'}$ = 0.6 Hz, $J_{5',6'a}$ = 7.0 Hz, $J_{5',6'b}$ = 7.0 Hz, 1H, H-5′), 3.80 (t, $J_{3,4}$ = 9.6 Hz, $J_{4,5}$ = 9.6 Hz, 1H, H-4), 3.63 (ddd, $J_{4,5}$ = 9.9 Hz, $J_{5,6a}$ = 1.7 Hz, $J_{5,6b}$ = 4.7 Hz, 1H, H-5), 3.50 (d, *J* = 15.3 Hz, 1H, SC*H*HCOOH), 3.26 (d, *J* = 15.3 Hz, 1H, SCH*H*COOH), 2.14 (s, 3H, C*H$_3$*CO), 2.13 (s, 3H, C*H$_3$*CO), 2.06 (s, 6H, C*H$_3$*CO), 2.045 (s, 3H, C*H$_3$*CO), 2.038 (s, 3H, C*H$_3$*CO), 2.03 (s, 3H, C*H$_3$*CO), 1.95 (s, 3H, C*H$_3$*CO) *ppm*; $^{13}$C NMR (100 MHz, CDCl$_3$): $\delta$ 173.7 (SCH$_2$*C*OOH), 171.1 (CH$_3$*C*O), 170.5 (CH$_3$*C*O), 170.3 (CH$_3$*C*O), 170.2

($CH_3$*C*O), 169.9 ($CH_3$*C*O), 169.8 ($CH_3$*C*O), 169.2 ($CH_3$*C*O), 101.1 (*C*-1′), 82.3 (*C*-1), 77.4 (*C*-5), 75.9 (*C*-4), 73.6 (*C*-3), 71.1 (*C*-3′), 70.9 (*C*-5′), 69.9 (*C*-2), 69.2 (*C*-2′), 66.8 (*C*-4′), 61.8 (*C*-6), 61.0 (*C*-6′), 31.3 (S*C*$H_2$COOH), 21.0 (*C*$H_3$CO), 21.9 (*C*$H_3$CO), 20.8 (*C*$H_3$CO × 4), 20.6 (*C*$H_3$CO) *ppm*; HRMS (ESI): *m*/*z* calcd for $C_{28}H_{38}NaO_{19}S$ $[M+Na]^+$ 733.1620. Found: 733.1612.

**Carboxymethyl β-D-galactopyranosyl-(1→4)-1-thio-β-D-glucopyranoside (12):**

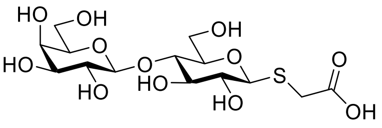

To a stirred solution of **11** (3.94 g, 5.54 mmol) in a mixture of $CH_3OH$ (20 mL) and water (20 mL) was added sodium methoxide (30 mL, 0.5 M in methanol) solution. After stirring overnight at room temperature, the reaction was neutralized by DOWEX 50WX8 (100−200 mesh, hydrogen form) resin, the resin was then removed by filtration, filtrate was concentrated, lyophilized in water, giving **12** (1.81 g, 78%) as a white powder: $R_f$ = 0.32 ($H_2O$–*i*PrOH–EtOAc, 1:2:2); mp 127 – 128 °C; $[\alpha]_D^{25}$ – 25.1 (*c* 1.00, $H_2O$); FT-IR (film): $\nu_{max}$ = 3344, 1708, 1582, 1376, 1021, 891, 783, 702, 611 $cm^{-1}$; $^1$H NMR (700 MHz, $D_2O$): $\delta$ 4.61 (d, $J_{1,2}$ = 9.9 Hz, 1H, H-1), 4.47 (d, $J_{1',2'}$ = 7.8 Hz, 1H, H-1′), 3.96 (dd, $J_{5,6a}$ = 2.2 Hz, $J_{6a,6b}$ = 12.5 Hz, 1H, H-6a), 3.94 (app d, $J_{3',4'}$ = 3.3 Hz, 1H, H-4′), 3.83−3.76 (m, 3H, H-6b, H-6′a, H-6′b), 3.75−3.70 (m, 2H, H-4, H-5′), 3.68 (dd, $J_{2',3'}$ = 10.0 Hz, $J_{3',4'}$ = 3.4 Hz, 1H, H-3′), 3.67 (t, $J_{2,3}$ = 8.9 Hz, $J_{3,4}$ = 8.9 Hz, 1H, H-3), 3.60 (ddd, $J_{4,5}$ = 9.7 Hz, $J_{5,6a}$ = 2.2 Hz, $J_{5,6b}$ = 4.8 Hz, 1H, H-5), 3.56 (dd, $J_{1',2'}$ = 7.8 Hz, $J_{2',3'}$ = 9.9 Hz, 1H, H-2′), 3.55 (d, *J* = 15.4 Hz, 1H, SC*H*HCOOH), 3.45 (d, *J* = 15.4 Hz, 1H, SCH*H*COOH), 3.43 (dd, $J_{1,2}$ = 9.7 Hz, $J_{2,3}$ = 9.0 Hz, 1H, H-2) *ppm*; $^{13}$C NMR (100 MHz, $D_2O$): $\delta$ 176.2 (SCH$_2$*C*OOH), 102.8 (*C*-1′), 84.6 (*C*-1), 78.7 (*C*-5), 77.9 (*C*-5′), 75.6 (*C*-3), 75.3 (*C*-4), 72.5 (*C*-3′), 71.8 (*C*-2), 70.9 (*C*-2′), 68.5 (*C*-4′), 61.0 (*C*-6′), 60.1 (*C*-6), 32.8 (S*C*$H_2$COOH) *ppm*; HRMS (ESI): *m*/*z* calcd for $C_{14}H_{24}NaO_{12}S$ $[M+Na]^+$ 439.0881. Found: 439.0881.

**2-(β-D-galactopyranosyl-(1→4)-β-D-glucopyranosylthio)aceto-1,2′-lactone (14):**

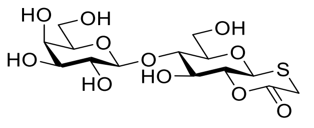

To a stirred solution of **12** (12.3 mg, 29.54 μmol) in dry DMSO (1.0 mL) were added *N*-hydroxysuccinimide (3.57 mg, 31.02 μmol), and *N*-(3-dimethylaminopropyl)-*N′*-ethylcarbodiimide hydrochloride (5.95 mg, 31.02 μmol) at room temperature. After stirring overnight, the solvent was removed by lyophilisation, the crude residue was purified by flash column chromatography ($H_2O$–*i*PrOH–EtOAc, 1:2:4) to give the lactone **14** (7.2 mg, 61%) as a colorless syrup: $R_f$ = 0.51 ($H_2O$–*i*PrOH–EtOAc, 1:2:4); $[\alpha]_D^{25}$ + 68.3 (*c* 1.00, $H_2O$); FT-IR (film): $\nu_{max}$ = 3367, 1704, 1219, 1066, 1016, 951, 709, 656 $cm^{-1}$; $^1H$ NMR (400 MHz, $D_2O$): $\delta$ 5.12 (d, $J_{1,2}$ = 9.8 Hz, 1H, H-1), 4.48 (d, $J_{1',2'}$ = 7.8 Hz, 1H, H-1′), 4.37 (t, $J_{1,2}$ = 9.6 Hz, $J_{2,3}$ = 9.6 Hz, 1H, H-2), 4.18 (d, $J$ = 14.7 Hz, 1H, SC*H*HCO), 4.06 (dd, $J_{2,3}$ = 9.3 Hz, $J_{3,4}$ = 8.5 Hz, 1H, H-3), 3.99 (dd, $J_{5,6a}$ = 2.1 Hz, $J_{6a,6b}$ = 12.5 Hz, 1H, H-6a), 3.94 (app d, $J_{3',4'}$ = 3.0 Hz, 1H, H-4′), 3.87−3.71 (m, 6H, H-4, H-6b, H-6′a, H-6′b, H-5′, H-5), 3.68 (dd, $J_{2',3'}$ = 10.0 Hz, $J_{3',4'}$ = 3.4 Hz, 1H, H-3′), 3.57 (dd, $J_{1',2'}$ = 7.7 Hz, $J_{2',3'}$ = 10.0 Hz, 1H, H-2′), 3.28 (d, $J$ = 14.7 Hz, 1H, SCH*H*CO) *ppm*; $^{13}C$ NMR (100 MHz, $D_2O$): $\delta$ 171.6 ($SCH_2$*C*O), 103.0 (*C*-1′), 79.7 (*C*-5), 78.4 (*C*-2), 78.3 (*C*-4), 76.6 (*C*-1), 75.4 (*C*-5′), 72.51 (*C*-3), 72.46 (*C*-3′), 70.9 (*C*-2′), 68.5 (*C*-4′), 60.9 (*C*-6′), 59.9 (*C*-6), 26.0 (S*C*$H_2$CO) *ppm*; HRMS (ESI): *m*/*z* calcd for $C_{14}H_{22}NaO_{11}S$ $[M+Na]^+$ 421.0775. Found: 421.0775.

**(2-Carboxamido)methyl (5-acetamido-3,5-dideoxy-D-*glycero*-α-D-*galacto*-non-2-ulopyranosylonic acid)-(2→3)-β-D-galactopyranosyl-(1→4)-1-thio-β-D-glucopyranoside:**

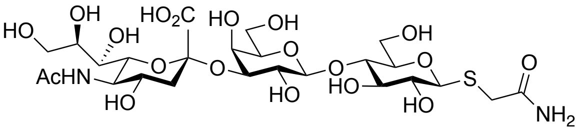


To a stirred solution of **1** (10 mg, 29 μmol) in dry DMSO (0.5 mL) were successively added potassium carbonate (8 mg, 116 μmol) and hydrogen peroxide solution (118.5 μl, 232 μmol, 30% w/w in water) at 0 °C. After stirring overnight at room temperature, the mixture was concentrated in vacuum, the crude residue was directly purified with size exclusion chromatography (LH20, $CH_3OH$–$H_2O$, 1:1, v/v), yielding the amide (10 mg, 97%) as a white powder after lyophilisation in water: $R_f$ = 0.29 ($H_2O$–*i*PrOH–EtOAc, 1:2:2); mp 193 – 194 °C; $[\alpha]_D^{25}$ – 6.5 (*c* 0.60, $H_2O$); FT-IR (neat): $\nu_{max}$ = 3657, 3314, 2981, 2889, 2489, 1610, 1473, 1462, 1383, 1251, 1139, 1072, 1030, 955, 895, 816, 676, 620 $cm^{-1}$; $^1H$ NMR (600 MHz, $D_2O$): $\delta$ 4.60 (d, $J_{1,2}$ = 9.9 Hz, 1H, H-1), 4.54 (d, $J_{1',2'}$ = 7.9 Hz, 1H, H-1′), 4.12 (dd, $J_{2',3'}$ = 9.9 Hz, $J_{3',4'}$ = 3.1 Hz, 1H, H-3′), 3.98–3.95 (m, 2H), 3.91–3.56 (m, 15H), 3.53 (d, $J$ = 15.4 Hz, 1H, SC*H*HCON$H_2$), 3.42 (d, $J$ = 15.4 Hz, 1H, SCH*H*CON$H_2$), 3.41 (t, $J_{2,1}$ = 9.4 Hz, $J_{2,3}$ = 9.4 Hz, 1H, H-2), 2.76 (dd, $J_{3''eq,3''ax}$ = 12.5 Hz, $J_{3''eq,4''}$ = 4.7 Hz, 1H, H-3″eq), 2.04 (s, 3H, C*H*$_3$CONH), 1.80 (t, $J_{3''ax,3''eq}$ = 12.5 Hz, $J_{3''ax,4''}$ = 12.5 Hz, 1H, H-3″ax) *ppm*; $^{13}C$ NMR (150 MHz, $D_2O$): $\delta$ 175.5 ($SCH_2$*C*ON$H_2$), 175.0 ($CH_3$*C*ONH), 173.9 (*C*-1″), 102.6 (*C*-1′), 99.8 (*C*-

2″), 85.0 (*C*-1), 78.7, 77.8, 75.6, 75.5, 75.2, 72.9, 71.9, 71.8, 69.3, 68.3, 68.1, 67.5, 62.6, 61.0, 60.0, 51.7, 39.6, 32.8, 22.0 (S*C*$H_2$CON$H_2$) *ppm*; HRMS (ESI): *m*/*z* calcd for $C_{25}H_{41}O_{19}N_2S$ $[M–H]^-$ 705.2030. Found: 705.2022.

***N*^1-Methyl-2-[β-D-galactopyranosyl-(1→4)-1-β-D-glucopyranosyl)]sulfonylacetamide**:

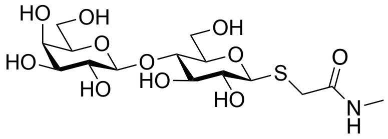

To a stirred solution of **12** (150 mg, 360 µmol) in dry DMF (5.0 mL) were added methylamine (270 µl, 540 µmol, 2 M in THF), and *N*-(3-dimethylaminopropyl)-*N′*-ethylcarbodiimide hydrochloride (82.8 mg, 432 µmol) at room temperature. After stirring overnight, the solvent was removed by concentration, the crude residue was purified by flash column chromatography ($H_2O$–*i*PrOH–EtOAc, 1:2:2) followed by size exclusion chromatography (LH20, $CH_3OH$–$H_2O$, 1:1, v/v) to give amide (77.3 mg, 50%) as a white powder after lyophilisation in water: $R_f$ = 0.42 ($H_2O$–*i*PrOH–EtOAc, 1:2:2); mp 97 – 98 °C; $[\alpha]_D^{25}$ – 22.2 (*c* 0.60, $H_2O$); FT-IR (neat): $\nu_{max}$ = 3326, 2884, 2483, 1635, 1558, 1460, 1408, 1036, 886, 822, 781, 700, 624 $cm^{-1}$; $^1H$ NMR (600 MHz, $D_2O$): $\delta$ 4.56 (d, $J_{1,2}$ = 9.9 Hz, 1H, H-1), 4.45 (d, $J_{1',2'}$ = 7.9 Hz, 1H, H-1′), 3.93 (dd, $J_{6'a,5}$ = 2.0 Hz, $J_{6'a,6'b}$ = 11.8 Hz, 1H, H-6′a), 3.92 (d, $J_{4',3'}$ = 3.6 Hz, 1H, H-4′), 3.81–3.71 (m. 4H, H-6′b, H-6a, H-6b, H-5), 3.70–3.62 (m, 3H, H-4, H-3′, H-3), 3.57 (ddd, $J_{5,4}$ = 2.0 Hz, $J_{5,6a}$ = 4.7 Hz, $J_{5,6b}$ = 9.4 Hz, 1H, H-5), 3.54 (dd, $J_{2',1'}$ = 8.0 Hz, $J_{2',3'}$ = 10.0 Hz, 1H, H-2′), 3.51 (d, $^2J$ = 15.4 Hz, 1H, SC*H*HCONHC$H_3$), 3.40 (t, $J_{2,1}$ = 8.9 Hz, $J_{2,3}$ = 8.9 Hz, 1H, H-2), 3.89 (d, $^2J$ = 15.4 Hz, 1H, SCH*H*CONHC$H_3$), 2.77 (s, 3H, SC$H_2$CONHC*$H_3$*) *ppm*; $^{13}C$ NMR (150 MHz, $D_2O$): $\delta$ 173.0 (SC$H_2$*C*ONHC$H_3$), 102.8 (*C*-1′), 85.0 (*C*-1), 78.6 (*C*-5), 77.8 (*C*-4), 75.6 (*C*-3), 75.3 (*C*-5′), 72.5 (*C*-3′), 71.8 (*C*-2), 70.9 (*C*-2′), 68.5 (*C*-4′), 61.0 (*C*-6), 60.1 (*C*-6′), 33.2 (S*C*$H_2$CONHC$H_3$), 26.2 (SC$H_2$CONH*C*$H_3$) *ppm*; HRMS (ESI): *m*/*z* calcd for $C_{15}H_{27}O_{11}NNaS$ $[M+Na]^+$ 452.1197. Found: 452.1197.

***N*^1-Methyl-2-[(5-acetamido-3,5-dideoxy-D-*glycero*-α-D-*galacto*-non-2-ulopyranosylonic acid)-(2→3)-β-D-galactopyranosyl-(1→4)-1-β-D-glucopyranosyl)]sulfonylacetamide:**

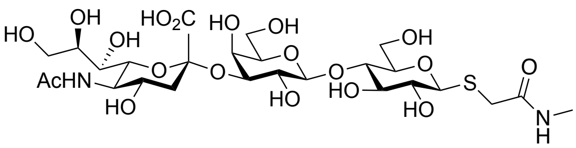

Substrate (5 mg, 11.64 μmol, the final concentration is 10 mM) was dissolved in fresh ammonium carbonate (1.164 mL, 100 mM, containing 20 mM $MgCl_2$, pH = 8.5) buffer, *N*-acetylneuraminic acid (3.78 mg, 12.23 μmol), cytidine-5'-triphosphate disodium salt (15.34 mg, 29.11 μmol), CMP-sialic acid synthetase (1.06 μL, 2.5 μg per mg substrate, 11.81 mg/mL in PBS buffer, *NmCSS*), and 2,3-sialyltransferase (1.04 μL, 3.0 μg per mg substrate, 14.465 mg/mL in PBS buffer, *PmST3*) were added, the resulting mixture was incubated at 37 °C/200rpm. After shaking for 3 h, the reaction was quenched by adding equal volume of cold ethanol (1.164 mL). The proteins and insoluble precipitates were removed by centrifugation, the supernatant was concentrated in vacuum, the crude residue was purified by flash column chromatography ($H_2O$–*i*PrOH–EtOAc, 1:2:2) followed by size exclusion chromatography (LH20, $CH_3OH$–$H_2O$, 1:1, v/v), the combined fractions were concentrated, lyophilized in water, yielding product (7.7 mg, 92%) as a white powder: $R_f$ = 0.36 ($H_2O$–*i*PrOH–EtOAc, 1:2:2); mp 181 – 182 °C; $[\alpha]_D^{25}$ – 14.1 (*c* 0.60, $H_2O$); FT-IR (neat): $\nu_{max}$ = 3657, 3276, 2981, 2888, 1631, 1473, 1462, 1382, 1251, 1149, 1072, 954, 895, 816, 679, 616 $cm^{-1}$; $^1H$ NMR (600 MHz, $D_2O$): $\delta$ 4.58 (d, $J_{1,2}$ = 9.9 Hz, 1H, H-1), 4.55 (d, $J_{1',2'}$ = 7.9 Hz, 1H, H-1′), 4.13 (dd, $J_{3',2'}$ = 9.8 Hz, $J_{3',4'}$ = 3.1 Hz, 1H, H-3′), 3.98–3.95 (m, 2H), 3.92–3.82 (m, 4H), 3.79–3.58 (m. 11H), 3.53 (d, $^2J$ = 15.3 Hz, 1H, SC*H*HCONHCH$_3$), 3.42 (t, $J_{2,1}$ = 9.1 Hz, $J_{2,3}$ = 9.1 Hz, 1H, H-2), 3.41 (d, $^2J$ = 15.3 Hz, 1H, SCH*H*CONHCH$_3$), 2.79 (s, 3H, SCH$_2$CONHC*H$_3$*), 2.78 (dd, $J_{3''eq,3''ax}$ = 12.5 Hz, $J_{3''eq,4''}$ = 4.6 Hz, 1H, H-3″eq), 2.05 (s, 3H, C*H$_3$*CONH), 1.82 (t, $J_{3''ax,3''eq}$ = 12.2 Hz, $J_{3''ax,4''}$ = 12.2 Hz, 1H, H-3″ax) *ppm*; $^{13}C$ NMR (150 MHz, $D_2O$): $\delta$ 175.0 (CH$_3$*C*ONH), 173.8 (*C*-1″), 173.0 (SCH$_2$*C*ONHCH$_3$), 102.6 (*C*-1′), 99.8 (*C*-2″), 85.1 (*C*-1), 78.7, 77.8, 75.6, 75.5, 75.2, 72.9, 71.85, 71.76, 69.4, 68.3, 68.1, 67.5, 62.6, 61.0, 60.1, 51.7, 39.6 (*C*-3″), 33.3 (S*C*H$_2$CONHCH$_3$), 26.2 (SCH$_2$CONH*C*H$_3$), 22.0 (*C*H$_3$CONH) *ppm*; HRMS (ESI): *m*/*z* calcd for $C_{26}H_{43}O_{19}N_2S$ $[M–H]^-$ 719.2186. Found: 719.2177.

**2-Aminoethyl (5-acetamido-3,5-dideoxy-D-*glycero*-α-D-*galacto*-non-2-ulopyranosylonic acid)-(2→3)-β-D-galactopyranosyl-(1→4)-β-D-glucopyranoside:**

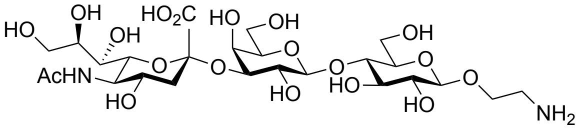


To a stirred solution of **SiaLacOCH$_2$CH$_2$N$_3$** (10 mg, 14.23 μmol) in a mixture of methanol (1 mL) and water (1 mL) was added the solution of triphenylphosphine (37.3 mg, 142.3 μmol) in THF (1 mL). After stirring at room temperature for one day, concentrated, the residues were suspended in water (5 mL), washed with $CH_2Cl_2$ (5 mL × 5) to remove excessive triphenylphosphine and its oxide, the aqueous layer was separated, concentrated, lyophilisation in water gave the desired amine (9.7 mg, quant.) as a white powder: $R_f$ = 0.43 (EtOAc–$CH_3OH$–AcOH–$H_2O$, 3:3:3:2); mp 221 – 223 °C; $[\alpha]_D^{25}$ + 0.3 (*c* 0.64, $H_2O$); FT-IR (neat): $\nu_{max}$ = 3273, 2929, 1609, 1437, 1379, 1319, 1113, 1069, 1030, 897, 838, 816, 783, 721, 696, 617 $cm^{-1}$; $^1H$ NMR (600 MHz, $D_2O$): $\delta$ 4.57 (d, $J_{1,2}$ = 7.8 Hz, 1H, H-1), 4.55 (d, $J_{1',2'}$ = 7.5 Hz, 1H, H-1′), 3.79–3.58 (m, 2H, H-3′, OC*H*HC$H_2NH_2$), 4.03–3.85 (m, 7H), 3.80–3.59 (m, 17H), 3.40 (t, $J_{2,1}$ = 8.4 Hz, $J_{2,3}$ = 8.4 Hz, 1H, H-2), 3.27 (t, *J* = 5.0 Hz, 2H, $OCH_2CH_2NH_2$), 2.79 (dd, $J_{3''eq,3''ax}$ = 12.4 Hz, $J_{3''eq,4''}$ = 4.4 Hz, 1H, H-3″eq), 2.06 (s, 3H, $CH_3$CONH), 1.82 (t, $J_{3''ax,3''eq}$ = 12.2 Hz, $J_{3''ax,4''}$ = 12.2 Hz, 1H, H-3″ax) *ppm*; $^{13}C$ NMR (150 MHz, $D_2O$): $\delta$ 175.1 ($CH_3$*C*ONH), 173.9 (*C*-1″), 102.7 (*C*-1′), 102.0 (*C*-1), 99.8 (*C*-2″), 78.1, 75.5 (*C*-3′), 75.2, 74.8, 74.2, 72.9, 72.7, 71.8, 69.4 (*C*-2′), 68.3, 68.1, 67.5, 66.2 ($OCH_2CH_2NH_2$), 62.6, 61.0, 59.9, 51.7, 39.7 (*C*-3″), 39.5 ($OCH_2CH_2NH_2$), 22.0 ($CH_3$CONH) *ppm*; HRMS (ESI): *m*/*z* calcd for $C_{25}H_{43}O_{19}N_2$ $[M–H]^-$ 675.2466. Found: 675.2459.

**Allyl 2,3,4,6-tetra-*O*-acetyl-β-D-galactopyranosyl-(1→4)-2,3,6-tri-*O*-acetyl-β-D-glucopyranoside (:**

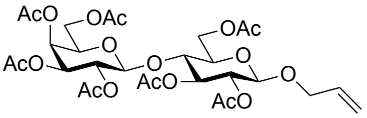


To a stirred solution of **lactose octaacetate** (10 g, 14.74 mmol) and allyl alcohol (1.20 ml, 17.68 mmol) in dry $CH_2Cl_2$ (100 mL) was added boron trifluoride etherate (2.73 ml, 22.10 mmol) dropwise at 0 °C, the reaction mixture was allowed to worm to room temperature and stirred for one day. The resulting yellow solution was quenched with triethylamine, washed with saturated $NaHCO_3$ solution, the organic layer was separated, the aqueous layer was extracted with $CH_2Cl_2$ (30 mL × 3), organic layers were combined, dried over $Na_2SO_4$, filtered and concentrated, the residue was purified with flash column chromatography (PE–EtOAc, 1:1), giving glycoside (4.47 g, 45%) as a white foam: $R_f$ = 0.49 (PE–EtOAc, 1:1); mp 57 – 58 °C; $[\alpha]_D^{25}$ – 7.7 (*c* 1.71, $CH_2Cl_2$); FT-IR (film): $\nu_{max}$ = 1745, 1432, 1369, 1217, 1271, 1133, 1048, 954, 902, 737 $cm^{-1}$; $^1H$ NMR (400 MHz, $CDCl_3$): $\delta$ 5.83 (dddd, $^3J_{trans}$ = 17.3 Hz, $^3J_{cis}$ = 10.6 Hz, $^3J_A$ = 4.9 Hz, $^3J_B$ = 6.1 Hz, 1H, $OCH_2CH$=$CH_2$), 5.34 (dd, $J_{4',3'}$ = 3.4 Hz, $J_{4',5'}$ = 1.0 Hz, 1H, H-4′), 5.25 (dddd, $^3J_{trans}$ = 17.2 Hz, $^2J$ = 1.6 Hz, $^4J_A$ = 1.6 Hz, $^4J_B$ = 1.6 Hz, 1H, $OCH_2CH$=C$H^{trans}$H), 5.193 (t, $J_{3,2}$ = 9.2 Hz, $J_{3,4}$ = 9.2 Hz, 1H, H-3), 5.192 (dddd, $^3J_{cis}$ = 10.4 Hz, $^2J$ = 1.6 Hz, $^4J_A$ = 1.3 Hz, $^4J_B$ = 1.3 Hz, 1H, $OCH_2CH$=CH$H^{cis}$), 5.10 (dd, $J_{2',1'}$ = 7.9 Hz, $J_{2',3'}$ = 10.4 Hz, 1H, H-2′), 4.95 (dd, $J_{3',2'}$ = 10.4 Hz, $J_{3',4'}$ = 3.4 Hz, 1H, H-3′), 4.92 (dd, $J_{2,1}$ = 7.9 Hz, $J_{2,3}$ = 9.5 Hz, 1H, H-2), 4.52 (d, $J_{1,2}$ = 7.9 Hz, 1H, H-1), 4.51–4.47 (m, 1H, H-6a),

4.78 (d, $J_{1',2'}$ = 7.8 Hz, 1H, H-1′), 4.30 (dddd, $^2J$ = 13.2 Hz, $^3J$ = 4.9 Hz, $^4J_{trans}$ = 1.5 Hz, $^4J_{cis}$ = 1.5 Hz, 1H, OC$H^A$HCH=CH$_2$), 4.15-4.04 (m, 4H, H-6′a, H-6′b, H-6b, OCH$H^B$CH=CH$_2$), 3.87 (ddd, $J_{5',4'}$ = 0.9 Hz, $J_{5',6'a}$ = 6.8 Hz, $J_{5',6'b}$ = 6.8 Hz, 1H, H-5′), 3.80 (t, $J_{4,3}$ = 9.6 Hz, $J_{4,5}$ = 9.6 Hz, 1H, H-4), 3.59 (ddd, $J_{5,4}$ = 9.8 Hz, $J_{5,6a}$ = 2.1 Hz, $J_{5,6b}$ = 5.0 Hz, 1H, H-5), 2.15 (s, 3H, C$H_3$CO), 2.12 (s, 3H, C$H_3$CO), 2.06 (s, 3H, C$H_3$CO), 2.04 (s, 9H, C$H_3$CO × 3), 1.96 (s, 3H, C$H_3$CO) *ppm*. Identical to previous report (*24*).

**Allyl β-D-galactopyranosyl-(1→4)-β-D-glucopyranoside:**

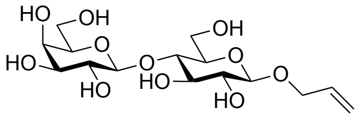

To a stirred solution of **peracetate** (4.47 g, 6.60 mmol) in methanol (40 mL) was added sodium methoxide solution (10 mL, 0.5 M in methanol) at room temperature, after stirring for 2 hours, reaction mixture was neutralised with DOWEX 50WX8 (100−200 mesh, hydrogen form) resin, the resin was then removed by filtration, the filtrate was concentrated to give the desired product (2.53 g, quant.) as a white powder after lyophilisation in water: $R_f$ = 0.32 ($H_2O$–*i*PrOH–EtOAc, 1:2:4); mp 160 – 161 °C; $[\alpha]_D^{25}$ + 1.0 (*c* 1.04, $H_2O$); FT-IR (neat): $\nu_{max}$ = 3267, 1428, 1327, 1218, 1166, 1045, 1020, 936, 892, 784, 700 cm$^{-1}$; $^1$H NMR (400 MHz, $D_2O$): $\delta$ 5.98 (dddd, $^3J_{trans}$ = 17.3 Hz, $^3J_{cis}$ = 10.4 Hz, $^3J_A$ = 6.0 Hz, $^3J_B$ = 6.0 Hz, 1H, OCH$_2$C$H$=CH$_2$), 5.38 (dddd, $^3J_{trans}$ = 17.3 Hz, $^2J$ = 1.5 Hz, $^4J_A$ = 1.5 Hz, $^4J_B$ = 1.5 Hz, 1H, OCH$_2$CH=C$H^{trans}$H), 5.29 (dddd, $^3J_{cis}$ = 10.4 Hz, $^2J$ = 1.1 Hz, $^4J_A$ = 1.1 Hz, $^4J_B$ = 1.1 Hz, 1H, OCH$_2$CH=CH$H^{cis}$), 4.53 (d, $J_{1,2}$ = 8.0 Hz, 1H, H-1), 4.45 (d, $J_{1',2'}$ = 7.8 Hz, 1H, H-1′), 4.39 (dddd, $^2J$ = 12.7 Hz, $^3J$ = 6.4 Hz, $^4J_{trans}$ = 1.1 Hz, $^4J_{cis}$ = 1.1 Hz, 1H, OC$H^A$HCH=CH$_2$), 4.23 (dddd, $^2J$ = 12.7 Hz, $^3J$ = 5.6 Hz, $^4J_{trans}$ = 1.2 Hz, $^4J_{cis}$ = 1.2 Hz, OCH$H^B$CH=CH$_2$), 3.98 (dd, $J_{6a,5}$ = 2.1 Hz, $J_{6a,6b}$ = 12.4 Hz, 1H, H-6a), 3.92 (d, $J_{4',5'}$ = 3.3 Hz, 1H, H-4′), 3.82–3.70 (m, 5H, H-6b, H-4, H-6′a, H-6′b, H-5′), 3.68–3.57 (m, 3H, H-3′, H-3, H-5), 3.54 (dd, $J_{2',1'}$ = 7.8 Hz, $J_{2',3'}$ = 9.9 Hz, 1H, H-2′), 3.37–3.31 (m, 1H, H-2) *ppm*. Identical to previous report (*24*).

**Allyl (5-acetamido-3,5-dideoxy-D-*glycero*-α-D-*galacto*-non-2-ulopyranosylonic acid)-(2→3)-β-D-galactopyranosyl-(1→4)-β-D-glucopyranoside:**

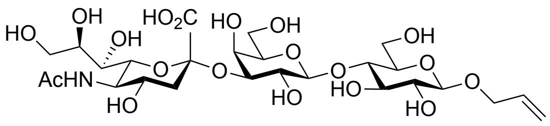

Substrate lactoside (700 mg, 1.83 mmol, the final concentration is 10 mM) was dissolved in fresh ammonium carbonate (183 mL, 100 mM, containing 20 mM $MgCl_2$, pH = 8.5) buffer,

*N*-acetylneuraminic acid (594.5 mg, 1.92 mmol), cytidine-5'-triphosphate disodium salt (2.41 g, 4.58 mmol), CMP-sialic acid synthetase (148.2 μL, 2.5 μg per mg substrate, 11.81 mg/mL in PBS buffer, *NmCSS*), and 2,3-sialyltransferase (145.2 μL, 3.0 μg per mg substrate, 14.465 mg/mL in PBS buffer, *PmST3*) were added, the resulting mixture was incubated at 37 °C/200rpm. After shaking for 3 h, the reaction was quenched by adding equal volume of cold ethanol (183 mL). The proteins and insoluble precipitates were removed by centrifugation, the supernatant was concentrated in vacuum, the crude residue was purified by flash column chromatography ($H_2O$–*i*PrOH–EtOAc, 1:2:2) followed by size exclusion chromatography (LH20, $CH_3OH$–$H_2O$, 1:1, v/v), the combined fractions were concentrated, lyophilized in water, yielding product (1.13 g, 92%) as a white powder: $R_f$ = 0.53 ($H_2O$–*i*PrOH–EtOAc, 1:2:2); mp 167 – 168 °C; $[\alpha]_D^{25}$ – 2.7 (*c* 1.00, $H_2O$); FT-IR (neat): $\nu_{max}$ = 3035, 1802, 1606, 1444, 1343, 1292, 1239, 1195, 1177, 1129, 1106, 1074, 1033, 997, 962, 903, 881, 838, 814, 752, 681, 638, 623 cm$^{-1}$; $^1$H NMR (600 MHz, $D_2O$): $\delta$ 6.00 (dddd, $^3J_{trans}$ = 17.2 Hz, $^3J_{cis}$ = 10.4 Hz, $^3J_A$ = 5.5 Hz, $^3J_B$ = 5.5 Hz, 1H, $OCH_2CH{=}CH_2$), 5.40 (dddd, $^3J_{trans}$ = 17.3 Hz, $^2J$ = 1.5 Hz, $^4J_A$ = 1.5 Hz, $^4J_B$ = 1.5 Hz, 1H, $OCH_2CH{=}CH^{trans}H$), 5.29 (dddd, $^3J_{cis}$ = 10.4 Hz, $^2J$ = 1.4 Hz, $^4J_A$ = 1.4 Hz, $^4J_B$ = 1.4 Hz, 1H, $OCH_2CH{=}CHH^{cis}$), 4.554 (d, $J_{1,2}$ = 8.0 Hz, 1H, H-1), 4.549 (d, $J_{1',2'}$ = 7.9 Hz, 1H, H-1′), 4.41 (dddd, $^2J$ = 12.7 Hz, $^3J$ = 5.6 Hz, $^4J_{trans}$ = 1.3 Hz, $^4J_{cis}$ = 1.3 Hz, 1H, $OCH^AHCH{=}CH_2$), 4.25 (dddd, $^2J$ = 12.7 Hz, $^3J$ = 6.4 Hz, $^4J_{trans}$ = 1.3 Hz, $^4J_{cis}$ = 1.3 Hz, $OCHH^BCH{=}CH_2$), 4.13 (dd, $J_{3',2'}$ = 9.9 Hz, $J_{3',4'}$ = 3.2 Hz, 1H, H-3′), 4.01 (dd, $J_{6a,5}$ = 2.2 Hz, $J_{6a,6b}$ = 12.3 Hz, 1H, H-6a), 3.98 (app d, $J_{4',3'}$ = 3.1 Hz, 1H, H-4′), 3.93–3.83 (m, 4H), 3.80–3.59 (m, 11H), 3.35 (dd, $J_{2,1}$ = 8.2 Hz, $J_{2,3}$ = 9.0 Hz, 1H, H-2), 2.78 (dd, $J_{3''eq,3''ax}$ = 12.5 Hz, $J_{3''eq,4''}$ = 4.7 Hz, 1H, H-3″eq), 2.05 (s, 3H, $CH_3CONH$), 1.82 (t, $J_{3''ax,3''eq}$ = 12.2 Hz, $J_{3''ax,4''}$ = 12.2 Hz, 1H, H-3″ax) *ppm*. Identical to previous report (*25*).

**5-Acetamido-3,5-dideoxy-D-*glycero*-α-D-*galacto*-non-2-ulopyranosylonic acid)-(2→3)-β-D-galactopyranosyl-(1→4)-β-D-glucopyranosyl-2-glycolaldehyde:**

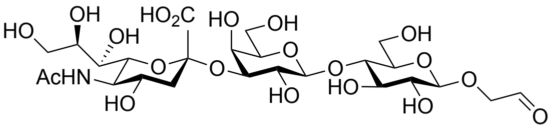


Ozone was bubbled into the solution of allyl glycoside (20mg) in methanol (5 ml) at – 78 °C until a blue colour appeared and remained for 30 min. the remaining ozone was then removed with oxygen bubbling (around 10 min). After dimethyl sulfide (0.2 mL) was added, the resulting solution was allowed to warm to room temperature and was stirred for 2 h prior to concentration in vacuo. The crude product was pelleted with ether (10 × volume of methanol), the solid was then purified by size exclusion chromatography (LH20, $CH_3OH$–

$H_2O$, 1:1, v/v), the pooled fractions were lyophilized in water, giving a white powder, which was used directly in protein modification: $R_f$ = 0.48 ($H_2O$–*i*PrOH–EtOAc, 1:2:2).

**Xanthen-9-yl 2,3,4,6-tetra-*O*-acetyl-β-D-galactopyranosyl-(1→4)-2,3,6-tri-*O*-acetyl-1-thio-β-D-glucopyranoside:**

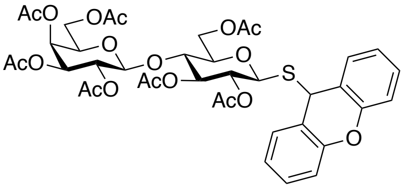


To a stirred solution of thioester **4** (5 g, 7.20 mmol) in dry $CH_3CN$ (20 mL) was added hydrazine acetate (796 mg, 8.64 mmol) at room temperature. After stirring for 2 h, the reaction mixture was diluted with $CH_2Cl_2$ (100 mL), washed with saturated $NaHCO_3$ (100 mL, aq.) solution, the organic layer was separated, the aqueous layer was then extracted with $CH_2Cl_2$ (30 mL × 2), the combined organic layer was washed with brine (200 mL), dried over $Na_2SO_4$, filtered and concentrated, giving the crude **1-thiosugar** as a yellow syrup. The crude residue above was then dissolved in dry $CH_2Cl_2$ (54 mL), trifluoroacetic acid (2.16 mL, 4%, v/v) was added. After stirring for 15 min at room temperature, xanthydrol (2.14 g, 10.8 mmol) was added dropwise. The resulting solution was stirred for 90 min at room temperature. Then, reaction mixture was diluted with $CH_2Cl_2$ (100 mL), washed with saturated $NaHCO_3$ (100 mL, aq.) solution, the organic layer was separated, the aqueous layer was then extracted with $CH_2Cl_2$ (50 mL × 2), the combined organic layer was dried over $Na_2SO_4$, filtered and concentrated, the crude residue was with flash column chromatography (PE–EtOAc, 1:1) to give thioglycoside (3.24 g, 54%) as a white foam: $R_f$ = 0.43 (PE–EtOAc, 1:1); mp 85 – 86 °C; $[\alpha]_D^{25}$ – 37.7 (*c* 1.02, $CH_2Cl_2$); FT-IR (film): $\nu_{max}$ = 1748, 1479, 1458, 1369, 1218, 1047, 901, 757 $cm^{-1}$; $^1H$ NMR (600 MHz, $CDCl_3$): $\delta$ 7.39–7.38 (m, 1H, Ar), 7.33–7.32 (m, 1H, Ar), 7.30–7.28 (m, 2H, Ar), 7.13–7.09 (m, 4H, Ar), 5.52 (s, 1H, SC*H*Ar), 5.32 (dd, $J_{4',3'}$ = 3.4 Hz, $J_{4',5'}$ = 0.7 Hz, 1H, H-4′), 5.07 (dd, $J_{2',1'}$ = 7.9 Hz, $J_{2',3'}$ = 10.4 Hz, 1H, H-2′), 5.03 (t, $J_{3,2}$ = 9.1 Hz, $J_{3,4}$ = 9.1 Hz, 1H, H-3), 4.92 (dd, $J_{3',2'}$ = 10.4 Hz, $J_{3',4'}$ = 3.8 Hz, 1H, H-3′), 4.84 (dd, $J_{2,1}$ = 10.0 Hz, $J_{2,3}$ = 9.3 Hz, 1H, H-2), 4.42 (d, $J_{1',2'}$ = 7.9 Hz, 1H, H-1′), 4.25 (dd, $J_{6a,5}$ = 1.9 Hz, $J_{6a,6b}$ = 12.1 Hz, 1H, H-6a), 4.18 (d, $J_{1,2}$ = 10.1 Hz, 1H, H-1), 4.09 (dd, $J_{6'a,5'}$ = 6.4 Hz, $J_{6'a,6'b}$ = 11.3 Hz, 1H, H-6′a), 4.05 (dd, $J_{6'b,5'}$ = 7.4 Hz, $J_{6'b,6'a}$ = 11.3 Hz, 1H, H-6′b), 4.00 (dd, $J_{6b,5}$ = 5.0 Hz, $J_{6b,6a}$ = 12.1 Hz, 1H, H-6b), 3.82 (app t, $J_{5',6'a}$ = 7.2 Hz, $J_{5',6'b}$ = 7.2 Hz, 1H, H-5′), 3.71 (t, $J_{4,3}$ = 9.4 Hz, $J_{4,5}$ = 9.4 Hz, 1H, H-4), 3.34 (ddd, $J_{5,4}$ = 9.9 Hz, $J_{5,6a}$ = 1.9 Hz, $J_{5,6b}$ = 4.9 Hz, 1H, H-5), 2.13 (s, 3H, C*H₃*CO), 2.10 (s, 3H, C$H_3$CO), 2.06 (s, 3H, C$H_3$CO), 2.03 (s, 3H, C$H_3$CO), 1.98 (s, 3H, C$H_3$CO), 1.95 (s, 3H, C$H_3$CO), 1.84

(s, 3H, C$H_3$CO) *ppm*; $^{13}$C NMR (150 MHz, $CDCl_3$): $\delta$ 170.3 ($CH_3$*C*O), 170.2 ($CH_3$*C*O), 170.1 ($CH_3$*C*O), 170.0 ($CH_3$*C*O), 169.6 ($CH_3$*C*O), 169.5 ($CH_3$*C*O), 169.1 ($CH_3$*C*O), 152.6 (Ar), 152.0 (Ar), 129.4 (Ar), 129.2 (Ar), 129.1 (Ar), 128.9 (Ar), 123.7 (Ar), 123.3 (Ar), 121.3 (Ar), 120.2 (Ar), 116.8 (Ar), 116.7 (Ar), 101.0 (*C*-1′), 81.9 (*C*-1), 76.4 (*C*-5), 76.0 (*C*-4), 73.8 (*C*-3), 71.0 (*C*-3′), 70.6 (*C*-5′), 70.0 (*C*-2), 69.1 (*C*-2′), 66.6 (*C*-4′), 62.2 (*C*-6), 60.7 (*C*-6′), 41.9 (S*C*HAr), 20.9 (*C*$H_3$CO), 20.72 (*C*$H_3$CO), 20.65 (*C*$H_3$CO), 20.59 (*C*$H_3$CO), 20.58 (*C*$H_3$CO), 20.53 (*C*$H_3$CO), 20.46 (*C*$H_3$CO) *ppm*; HRMS (ESI): *m*/*z* calcd for $C_{41}H_{43}O_{18}NaS$ [M+Na]$^+$ 855.2165. Found: 855.2132.

**Xanthen-9-yl β-D-galactopyranosyl-(1→4)-1-thio-β-D-glucopyranoside:**

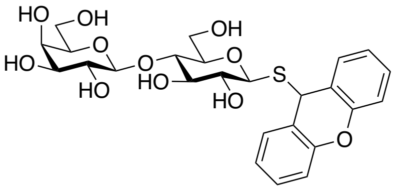

To a stirred solution of protected thioglycoside (3.05 g, 3.66 mmol) in methanol (30 mL) was added sodium methoxide solution (5 mL, 0.5 M in methanol) at room temperature, after stirring for 2 hours, reaction mixture was neutralised with DOWEX 50WX8 (100−200 mesh, hydrogen form) resin, the resin was then removed by filtration, the filtrate was concentrated to give the desired product (1.98 g, quant.) as a white powder after lyophilisation in water: $R_f$ = 0.65 ($H_2O$–*i*PrOH–EtOAc, 1:2:4); mp 201 – 202 °C; $[\alpha]_D^{25}$ – 96.4 (*c* 0.62, DMSO); FT-IR (neat): $\nu_{max}$ = 3392, 2980, 1604, 1574, 1481, 1453, 1377, 1326, 1256, 1214, 1182, 1120, 1080, 1032, 992, 900, 831, 783, 742, 697, 664, 615 cm$^{-1}$; $^1$H NMR (600 MHz, DMSO-$d_6$ with 5% $D_2O$): $\delta$ 7.44–7.43 (m, 1H, Ar), 7.39–7.37 (m, 1H, Ar), 7.35–7.31 (m, 2H, Ar), 7.20–7.14 (m, 4H, Ar), 5.69 (s, 1H, SC*H*Ar), 4.19 (app d, $J_{1',2'}$ = 7.7 Hz, 1H, H-1′), 4.00 (d, $J_{1,2}$ = 9.7 Hz, 1H, H-1), 3.88 (dd, $J_{6a,5}$ = 1.4 Hz, $J_{6a,6b}$ = 12.4 Hz, 1H, H-6a), 3.67–3.61 (m, 2H, H-5, H-6b), 3.52-3.44 (m, 3H, H-6′a, H-6′b, H-5′), 3.31-3.29 (m, 4H, H-2′, H-3′, H-4, H-4′), 3.19 (t, $J_{3,2}$ = 8.4 Hz, $J_{3,4}$ = 8.4 Hz, 1H, H-3), 3.07 (dd, $J_{2,1}$ = 9.7 Hz, $J_{2,3}$ = 8.6 Hz, 1H, H-2) *ppm*; $^{13}$C NMR (150 MHz, DMSO-$d_6$ with 5% $D_2O$): $\delta$ 152.2 (Ar), 151.4 (Ar), 129.2 (Ar), 128.9 (Ar), 128.8 (Ar), 128.7 (Ar), 123.7 (Ar), 123.2 (Ar), 122.0 (Ar), 121.3 (Ar), 116.4 (Ar), 116.1 (Ar), 103.5 (*C*-1′), 83.0 (*C*-1), 80.4 (*C*-4), 79.0 (*C*-4′), 76.2 (*C*-3), 75.2 (*C*-5′), 72.8 (*C*-3′), 72.1 (*C*-2), 70.3 (*C*-2′), 67.9 (*C*-5), 60.5 (*C*-6), 60.2 (*C*-6′), 38.6 (S*C*HAr) *ppm*; HRMS (ESI): *m*/*z* calcd for $C_{25}H_{30}O_{11}NaS$ [M+Na]$^+$ 561.1401. Found: 561.1400.

**Xanthen-9-yl 5-acetamido-3,5-dideoxy-D-*glycero*-α-D-*galacto*-non-2-ulopyranosylonic acid)-(2→3)-β-D-galactopyranosyl-(1→4)-β-D-glucopyranoside:**

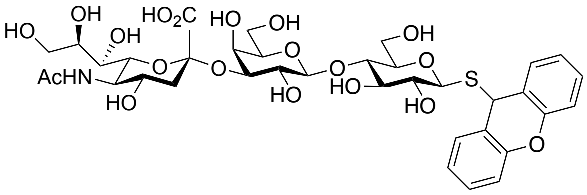

Substrate lactoside (500 mg, 928.4 μmol, the final concentration is 10 mM) was dissolved in fresh ammonium carbonate (93 mL, 100 mM, containing 20 mM $MgCl_2$, pH = 8.5) buffer and DMF (4.64 ml, 5% of buffer, optimised), *N*-acetylneuraminic acid (301.5 mg, 974.8 μmol), cytidine-5'-triphosphate disodium salt (1.22 g, 2.32 mmol), CMP-sialic acid synthetase (105.8 μL, 2.5 μg per mg substrate, 11.81 mg/mL in PBS buffer, *NmCSS*), and 2,3-sialyltransferase (103.7 μL, 3.0 μg per mg substrate, 14.465 mg/mL in PBS buffer, *PmST3*) were added, the resulting mixture was incubated at 37 °C/200rpm. After shaking for 3 h, the reaction was quenched by adding equal volume of cold ethanol (93 mL). The proteins and insoluble precipitates were removed by centrifugation, the supernatant was concentrated in vacuum, the crude residue was purified by flash column chromatography ($H_2O$–*i*PrOH–EtOAc, 1:2:3) followed by size exclusion chromatography (LH20, $CH_3OH$–$H_2O$, 1:1, v/v), the combined fractions were concentrated, lyophilized in water, yielding product (578 mg, 75%) as a white powder: $R_f$ = 0.51 ($H_2O$–*i*PrOH–EtOAc, 1:2:3); mp 78 – 79 °C; $[\alpha]_D^{25}$ – 31.2 (*c* 0.50, $H_2O$); FT-IR (neat): $\nu_{max}$ = 3294, 2161, 2031, 1604, 1478, 1458, 1378, 1324, 1255, 1068, 1026, 899, 817, 755, 681, 615 $cm^{-1}$; $^1H$ NMR (700 MHz, $D_2O$): $\delta$ 7.52–7.50 (m, 1H, Ar), 7.46–7.42 (m, 1H, Ar), 7.41–7.36 (m, 2H, Ar), 7.25–7.16 (m, 4H, Ar), 5.69 (s, 1H, SC*H*Ar), 4.48 (d, $J_{1',2'}$ = 7.8 Hz, 1H, H-1′), 4.12 (d, $J_{1,2}$ = 10.0 Hz, 1H, H-1), 4.09 (dd, $J_{3',2'}$ = 9.9 Hz, $J_{3',4'}$ = 3.2 Hz,1H, H-3′), 3.94 (app d, $J_{4',3'}$ = 3.2 Hz, 1H, H-4′), 3.90–3.87 (m, 2H), 3.86 (t, $J_{5'',4''}$ = 10.2 Hz, $J_{5'',6''}$ = 10.2 Hz,1H, H-5″), 3.74–3.63 (m, 8H), 3.61–3.58 (m, 2H), 3.54 (dd, $J_{2',1'}$ = 7.9 Hz, $J_{2',3'}$ = 9.8 Hz,1H, H-2′), 3.41 (t, $J_{3,2}$ = 8.9 Hz, $J_{3,4}$ = 8.9 Hz, 1H, H-3), 3.26 (ddd, $J_{5,4}$ = 9.9 Hz, $J_{5,6a}$ = 3.0 Hz, $J_{5,6b}$ = 3.0 Hz, 1H, H-5), 3.24 (dd, t, $J_{2,1}$ = 10.0 Hz, $J_{2,3}$ = 9.2 Hz, 1H, H-2), 2.76 (dd, $J_{3''eq,3''ax}$ = 12.4 Hz, $J_{3''eq,4''}$ = 4.7 Hz, 1H, H-3″eq), 2.04 (s, 3H, C$H_3$CONH), 1.80 (t, $J_{3''ax,3''eq}$ = 12.1 Hz, $J_{3''ax,4''}$ = 12.1 Hz, 1H, H-3″ax) *ppm*; $^{13}C$ NMR (150 MHz, $D_2O$): $\delta$ 175.0 ($CH_3$*C*ONH), 173.9 (*C*-1″), 152.3 (Ar), 151.6 (Ar), 129.7 (Ar), 129.4 (Ar), 129.1 (Ar), 124.2 (Ar), 123.9 (Ar), 122.3 (Ar), 120.6 (Ar), 116.44 (Ar), 116.39 (Ar), 102.5 (*C*-1′), 99.8 (*C*-2″), 83.9 (*C*-1), 78.4 (*C*-5), 77.7 (*C*-4), 75.8 (*C*-3), 75.5 (*C*-3′), 75.1, 72.9 (*C*-6″), 71.8, 71.4 (*C*-2), 69.3 (*C*-2′), 68.4, 68.1, 67.5 (*C*-4′), 62.6, 61.0, 59.9 (*C*-6), 51.7 (*C*-5″), 41.5 (S*C*HAr), 39.7 (*C*-3″), 22.0 (*C*$H_3$CONH) *ppm*; HRMS (ESI): *m*/*z* calcd for $C_{36}H_{47}NO_{19}NaS$ $[M+Na]^+$ 852.2355. Found: 852.2351.

**Bis[5-acetamido-3,5-dideoxy-D-*glycero*-α-D-*galacto*-non-2-ulopyranosylonic acid)-(2→3)-β-D-galactopyranosyl-(1→4)-1-thio-β-D-glucopyranosyl]1,1′-disulfide:**

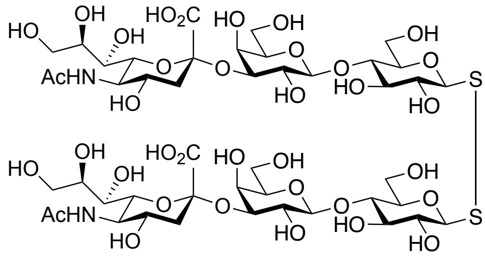

To a stirred solution of thioglycoside (20 mg, 24.1 μmol) in dry methanol (4.0 mL) was added iodine (61.2 mg, 241 μmol), after being stirred for one hour at room temperature, ether (10 × volume of methanol) was added to precipitate sugars. After centrifugation, the pellet was then purified by size exclusion chromatography (LH20, $CH_3OH$–$H_2O$, 1:1, v/v) to give the desired dimer (11 mg, 35%) as a white powder after lyophilization in water: $R_f$ = 0.08 ($H_2O$–*i*PrOH–EtOAc, 1:2:2); mp 207 – 208 °C; $[\alpha]_D^{25}$ – 34.1 (*c* 0.20, $H_2O$); FT-IR (neat): $\nu_{max}$ = 3335, 1614, 1568, 1435, 1396, 1377, 1320, 1295, 1238, 1211, 1071, 1035, 947, 894, 881, 814, 781, 677, 614 $cm^{-1}$; $^1H$ NMR (600 MHz, $D_2O$): $\delta$ 4.64 (d, $J_{1,2}$ = 9.4 Hz, 1H, H-1), 4.56 (d, $J_{1',2'}$ = 7.8 Hz, 1H, H-1′), 4.13 (dd, $J_{3',2'}$ = 9.8 Hz, $J_{3',4'}$ = 2.9 Hz,1H, H-3′), 4.03–3.98 (m, 2H), 3.92–3.85 (m, 4H), 3.80–3.58 (m, 12H), 2.78 (2.76 (dd, $J_{3''eq,3''ax}$ = 12.5 Hz, $J_{3''eq,4''}$ = 4.6 Hz, 1H, H-3″eq), 2.05 (s, 3H, C$H_3$CONH), 1.82 (t, $J_{3''ax,3''eq}$ = 12.1 Hz, $J_{3''ax,4''}$ = 12.1 Hz, 1H, H-3″ax) *ppm*; $^{13}C$ NMR (150 MHz, $D_2O$): $\delta$ 175.0 ($CH_3$*C*ONH), 173.9 (*C*-1″), 102.6 (*C*-1′), 99.8 (*C*-2″), 89.4 (*C*-1), 79.1, 77.5, 75.54, 75.49, 75.2, 72.9, 71.8, 71.0, 69.4, 68.4, 68.1, 67.5, 62.6, 61.1, 60.1, 51.7, 39.7 (*C*-3″), 22.0 (*C*$H_3$CONH) *ppm*; HRMS (ESI): *m*/*z* calcd for $C_{46}H_{75}N_2O_{36}S_2$ $[M–H]^-$ 1295.3535. Found: 1295.3521.

**2-Acetamido-3,4,6-tri-*O*-acetyl-2-deoxy-α-D-glucopyranosyl Chloride (15):**

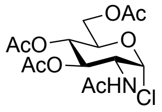

*N*-Acetyl-D-glucosamine (20 g, 90.4 mmol) was added to acetyl chloride (58 ml, 814 mmol) portion-wise in 5 mins under an argon atmosphere. After stirring for 2 days at room temperature, $CH_2Cl_2$ (100 ml) was added to the dark-red solution, poured into a mixture of ice (100 gram) and water (100 ml) followed by neutralization with saturated $NaHCO_3$ solution, the suspension was then extracted with $CH_2Cl_2$ (150 ml × 3), the combined organic layers were dried over $Na_2SO_4$, filtered and concentrated. The crude residue was dissolved in the minimum amount of $CH_2Cl_2$ and recrystallised with chilled ether. The precipitates were collected with filtration, dried in high vacuo, giving the desired chlorosugar **15** (28 g, 85%) as yellow crystals: $R_f$ = 0.65 (pure EtOAc); mp 123 – 124 °C; $[\alpha]_D^{25}$ + 121.7 (*c* 1.00, $CH_2Cl_2$); FT-IR (film): $\nu_{max}$ = 1745, 1667, 1536, 1434, 1368, 1223, 1116, 1077, 1043, 960, 910, 768, 735, 673, 645 cm$^{-1}$; $^1$H NMR (400 MHz, $CDCl_3$): $\delta$ 6.18 (d, $J_{1,2}$ = 3.7 Hz, 1H, H-1), 5.83 (d, $J_{2,NH}$ = 8.8 Hz, 1H, N*H*), 5.32 (dd, $J_{2,3}$ = 9.5 Hz, $J_{3,4}$ = 9.8 Hz, 1H, H-3), 5.21 (t, $J_{3,4}$ = 9.8 Hz, $J_{4,5}$ = 9.8 Hz, 1H, H-4), 4.53 (ddd, $J_{1,2}$ = 3.7 Hz, $J_{2,NH}$ = 8.8 Hz, $J_{2,3}$ = 9.5 Hz, 1H, H-2), 4.30−4.25 (m, 2H, H-6a, H-5), 4.14−4.11 (m, 1H, H-6b), 2.10 (s, 3H, C*H₃*CO), 2.05 (s, 6H, C*H₃*CO × 2), 1.98 (s, 3H, C*H₃*CO) *ppm*. Identical to the previous report (*26*).

**(2-Acetamido-3,4,6-tri-*O*-acetyl-2-deoxy-β-D-glucopyranosyl)-1-isothiouronium Chloride (16):**

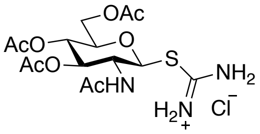


To a stirred solution of **15** (20.5 g, 56 mmol) in dry acetone (100 mL) was added thiourea (8.53 g, 112 mmol), the suspension was then refluxed at 80°C for 2 h under an argon atmosphere. After cooling to room temperature, the precipitate was collected with filtration, washed with chilled ethanol, dried under high vacuo overnight, yielding **16** (23.2 g, 94%) as white crystals: $R_f$ = 0.22 (pure $CH_3OH$); mp 173 – 174 °C; $[\alpha]_D^{25}$ – 26.3 (*c* 1.00, $H_2O$); FT-IR (film): $\nu_{max}$ = 1749, 1648, 1542, 1434, 1367, 1301, 1223, 1207, 1097, 1077, 1055, 1027, 978, 943, 909, 875, 813, 686, 652, 636, 620 cm$^{-1}$; $^1$H NMR (400 MHz, DMSO-$d_6$): $\delta$ 9.40 (br s, 2H, SC(*NH₂*)NH₂), 9.20 (br s, 2H, SC(NH₂)*NH₂*), 8.41(br d, $J_{2,NH}$ = 9.2 Hz, 1H, N*H*), 5.67 (d, $J_{1,2}$ = 10.3 Hz, 1H, H-1), 5.12 (t, $J_{2,3}$ = 9.7 Hz, $J_{3,4}$ = 9.7 Hz, 1H, H-3), 4.93 (t, $J_{3,4}$ = 9.7 Hz, $J_{4,5}$ = 9.7 Hz, 1H, H-4), 4.23−4.15 (m, 2H, H-5, H-6a), 4.07−3.97 (m, 2H, H-6b, H-2), 2.01 (s, 3H, C*H₃*CO), 1.98 (s, 3H, C*H₃*CO), 1.93 (s, 3H, C*H₃*CO), 1.80 (s, 3H, C*H₃*CO) *ppm*. Identical to the previous report (*27*).

**Cyanomethyl 2-acetamido-3,4,6-tri-*O*-acetyl-2-deoxy-1-thio-β-D-glucopyranoside (17):**

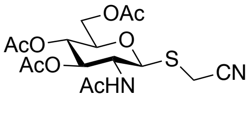


To a stirred solution of **16** (19.32 g, 43.72 mmol) in a mixture of water (120 ml) and acetone (120 ml) were added sodium metabisulfite (18.29 g, 96.19 mmol), potassium carbonate (7.86 g, 56.84 mmol), and chloroacetonitrile (55.30 ml, 874 mmol), successively. After stirring overnight at room temperature, the suspension was poured into a mixture of ice (100

g) and water (100 ml) followed by extraction with $CH_2Cl_2$ (150 ml × 3), the combined organic layers were dried over $Na_2SO_4$, filtered and concentrated in high vacuo, giving **17** (17.5 g, 99%) as an amorphous solid: $R_f$ = 0.52 (pure EtOAc); mp 175 – 176 °C; $[\alpha]_D^{25}$ – 96.3 (*c* 1.00, $CH_2Cl_2$); FT-IR (film): $\nu_{max}$ = 2117, 1743, 1662, 1534, 1434, 1371, 1229, 1046, 963, 916, 820, 736 $cm^{-1}$; $^1H$ NMR (500 MHz, $CDCl_3$): $\delta$ 5.90 (br d, $J_{2,NH}$ = 9.3 Hz, 1H, N*H*), 5.19 (t, $J_{2,3}$ = 9.5 Hz, $J_{3,4}$ = 9.5 Hz, 1H, H-3), 5.12 (t, $J_{3,4}$ = 9.8 Hz, $J_{4,5}$ = 9.8 Hz, 1H, H-4), 4.78 (d, $J_{1,2}$ = 10.4 Hz, 1H, H-1), 4.25−4.16 (m, 3H, H-6a, H-6b, H-2), 3.77 (ddd, $J_{4,5}$ = 9.8 Hz, $J_{5,6a}$ = 2.4 Hz, $J_{5,6b}$ = 4.9 Hz, 1H, H-5), 3.67 (d, $^2J$ = 17.0 Hz, 1H, SC*H*HCN), 3.32 (d, $^2J$ = 17.0 Hz, 1H, SCH*H*CN), 2.09 (s, 3H, C$H_3$CO), 2.05 (s, 3H, C$H_3$CO), 2.04 (s, 3H, C$H_3$CO), 1.97 (s, 3H, C$H_3$CONH) *ppm*; $^{13}C$ NMR (125 MHz, $CDCl_3$) $\delta$ 171.2 ($CH_3$*C*O), 170.7 ($CH_3$*C*O), 170.5 ($CH_3$*C*ONH), 169.2 ($CH_3$*C*O), 116.1 ($SCH_2$*C*N), 83.0 (*C*-1), 76.3 (*C*-5), 73.3 (*C*-3), 68.0 (*C*-4), 61.8 (*C*-6), 52.8 (*C*-2), 23.1 (*C*$H_3$CONH), 20.7 (*C*$H_3$CO), 20.63 (*C*$H_3$CO), 20.56 (*C*$H_3$CO), 14.6 (S*C*$H_2$CN) *ppm*; HRMS (ESI): *m*/*z* calcd for $C_{16}H_{23}N_2O_8S$ $[M+H]^+$ 403.1170. Found: 403.1171.

**Cyanomethyl 2-acetamido-2-deoxy-1-thio-β-D-glucopyranoside (18):**

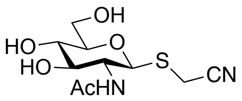


To a stirred solution of **17** (16.8 g, 41.75 mmol) in dry $CH_3OH$ (200 mL) was added triethylamine (58.2 mL, 417.5 mmol), the mixture was heated to 40 °C for 24 h. The resulting precipitates were collected with filtration, washed with chilled methanol, dried under high vacuo and lyophilised in water, giving the desired **18** (9.9 g, 86%) as a white powder: $R_f$ = 0.24 ($CH_2Cl_2$–$CH_3OH$, 5:1); mp 170 – 171 °C; $[\alpha]_D^{25}$ – 76.3 (*c* 1.00, $H_2O$); FT-IR (film): $\nu_{max}$ = 3285, 2245, 1648, 1538, 1373, 1312, 1285, 1060, 1028, 1005, 945, 882, 816, 693, 607 $cm^{-1}$; $^1H$ NMR (500 MHz, $CD_3OD$): $\delta$ 4.66 (d, $J_{1,2}$ = 10.2 Hz, 1H, H-1), 3.91 (dd, $J_{5,6a}$ = 1.7 Hz, $J_{6a,6b}$ = 12.3 Hz, 1H, H-6a), 3.86 (d, $^2J$ = 17.1 Hz, 1H, SC*H*HCN), 3.80 (t, $J_{1,2}$ = 10.2 Hz, $J_{2,3}$ = 10.2 Hz, 1H, H-2), 3.70–3.67 (m, 1H, H-6b), 3.58 (d, $^2J$ = 17.1 Hz, 1H, SCH*H*CN), 3.49–3.46 (m, 1H, H-3), 3.37–3.34 (m, 2H, H-4, H-5), 1.98 (s, 3H, C$H_3$CO) *ppm*; $^{13}C$ NMR (125 MHz, $CD_3OD$) $\delta$ 173.7 ($CH_3$*C*O), 118.6 ($SCH_2$*C*N), 84.7 (*C*-1), 82.5 (*C*-5), 76.9 (*C*-3), 71.9 (*C*-4), 62.9 (*C*-6), 55.7 (*C*-2), 22.8 (*C*$H_3$CO), 14.8 (S*C*$H_2$CN) *ppm*; HRMS (ESI): *m*/*z* calcd for $C_{10}H_{17}N_2O_5S$ $[M+H]^+$ 277.0853. Found: 277.0853.

**2,3,4,6-Tetra-*O*-acetyl-α-D-galactopyranosyl 2,2,2-trichloroacetimidate (L-2):**

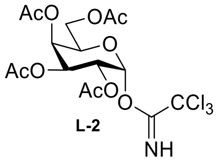


**L-2**

To a solution of the commercially available *β*-D-pentaacetyl galactose (**L-1**) (11 g, 28.3 mmol) in dry DMF (100 mL) was added hydrazine acetate (2.88 g, 31.2 mmol), under nitrogen. The mixture was stirred at 50 °C for 5 h, then recovered with EtOAc (200 mL), washed with satd. solution of $NaHCO_3$ (3 x 125 mL) and brine (125 mL). The organic layer was dried over $MgSO_4$ and evaporated to afford the hemiacetal intermediate as a colourless oil which was used directly without purification. To a solution of hemiacetal intermediate (9.87 g, 25.6 mmol, 1 eq.) in DCM (100 mL) were added trichloro acetonitrile (25.67 mL, 256 mmol, 10 eq.) and DBU (766 µL, 5.12 mmol, 0.2 eq.), under nitrogen. The mixture was stirred at r.t. for 16 h, then evaporated and purified by flash column chromatography over silica (PE–EtOAc, 2:1 to 1:1) to afford the titled imidate **L-2** as a light yellow solid (9.15 g, 18.6 mmol, 73%): $R_f$ = 0.5 (PE–EtOAc, 2:1); $[\alpha]_D^{25}$ +105.8 (*c* = 1.00, $CHCl_3$); IR (ATR) 1745.24 (CO); $^1H$ NMR ($CDCl_3$, 400 MHz) *δ* 8.65 (s, 1 H, NH), 6.51 (d, *J* = 3.3 Hz, 1 H, H-1), 5.47 (d, *J* = 2.0 Hz, 1 H, H-4), 5.30 (ddd, *J* = 26.2, 10.9, 3.2 Hz, 2 H, H-3, H-2), 4.36 (t, *J* = 6.5 Hz, 1 H, H-5), 4.08 (dd, *J* = 11.3, 6.5 Hz, 1 H, H-6a), 4.04–3.88 (m, 1 H, H-6b), 2.09, 1.95, 1.93, 1.93 (4 x s, 4 x 3 H, 4 x OAc) *ppm*; $^{13}C$ NMR ($CDCl_3$, 100 MHz) *δ* 170.18, 170.03, 169.97, 169.87 (4 x CO), 160.74 (CNH), 93.43 (C-1), 90.70 ($CCl_3$), 68.93 (C-5), 67.42, 67.30 (C-3, C-4), 66.82 (C-2), 61.18 (C-6), 20.58, 20.54, 20.53, 20.46 (4 x OAc) *ppm*; MS ($ESI^+$) m/z = 493.70 [M+H]. Identical to the previous report (*28*).

**Ethyl 2,3,4,-tri-*O*-acetyl-1-thio-β-L-fucopyranoside (L-4):**

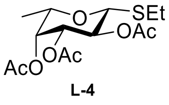


**L-4**

To a solution of 1,2,3,4-tetra-*O*-acetyl-6-deoxy-*β*-L-fucopyranoside (**L-3**)[6] (29.4 g, 79.7 mmol, 1 eq.) in DCM (dry, 40 mL), were added HBr (33% in AcOH, 34.6 mL, 199.3 mmol, 2.5 eq.) and $Ac_2O$ (6.02 mL, 63.8 mmol, 0.8 eq.). The mixture was stirred at r.t. for 1 h, poured into ice/water (250 mL), recovered with EtOAc (200 mL) and separated. The organic layer was washed with $H_2O$ (125 mL), satd. solution of $NaHCO_3$ (3 x 125 mL), dried over $MgSO_4$ and evaporated to afford bromosugar as a yellow oil which was unable and was used directly without further purification. To a solution of bromosugar (31.25 g, 88.5 mmol,

1 eq.) in ACN (180 mL), was added thiourea (7.07 g, 92.9 mmol, 1.05 eq.) and the mixture was refluxed at 80 °C for 30 min. The reaction was filtered, the precipitate washed with ACN and re-suspended in ACN (180 mL). Triethylamine (30.8 mL, 221.25 mmol, 2.5 eq.) and EtBr (9.9 mL, 132.75 mmol, 1.5 eq.) were added, and the reaction stirred at r.t for 1 h. The mixture was filtered, the precipitate dissolved in DCM (150 mL) and washed with $H_2O$ (125 mL), HCl 1M (125 mL) and brine (125 mL), dried over $MgSO_4$ and evaporated to afford the titled thioglycoside **L-4** as a colourless oil (22.20 g, 66.4 mmol, 75%): $R_f$ = 0.9 (EtOAc–PE, 4:1); $[\alpha]_D^{25}$ + 7.4 (*c* = 1.00, $CHCl_3$); IR (ATR) 1745.70 (CO); $^1H$ NMR ($CDCl_3$, 200 MHz) $\delta$ 5.25 (ddd, *J* = 18.7, 11.0, 6.5 Hz, 2 H, H-4, H-2), 5.04 (dd, *J* = 9.9, 3.4 Hz, 1 H, H-3), 4.45 (d, *J* = 9.7 Hz, 1 H, H-1), 3.97–3.61 (m, 1 H, H-5), 2.73 (qd, *J* = 7.4, 3.4 Hz, 2 H, $CH_2$), 2.17, 2.06, 1.98 (3 x s, 3 x 3 H, 3 x OAc), 1.29 (d, *J* = 7.4 Hz, 3 H, H-6), 1.21 (d, *J* = 6.4 Hz, 3 H, $CH_3$) *ppm*; $^{13}C$ NMR ($CDCl_3$, 50 MHz) $\delta$ 170.63, 170.52, 170.33 (3 x CO), 83.66 (C-1), 73.33 (C-5), 72.50 (C-3), 70.61 (C-4), 67.52 (C-2), 24.25 ($CH_2$), 20.03, 20.00, 19.98 (3 x Ac), 16.57 ($CH_3$), 14.87 (C-6) *ppm*; MS ($ESI^-$) m/z = 333.37 [M-H]. Identical to the previous report (*29*).

**Ethyl 2,3,4,-tri-*O*-*p*-methoxybenzyl-1-thio-β-L-fucopyranoside (L-5):**

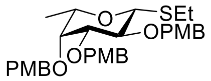


**L-5**

To a solution of **L-4** (22.2 g, 66.4 mmol, 1 eq.) in MeOH (66 mL), was added NaOMe (25% in MeOH, 13.6 mL, 66.4 mmol, 1 eq.) and the mixture was stirred at r.t. for 1 h. The reaction was quenched with Dowex $H^+$, filtered and evaporated to afford the free sugar as a white solid which was used directly without further purification. To a solution of the solid above and TBAI (2.8 g, 7.58 mmol, 0.1 eq.) in DMF (dry, 150 mL), was added, at 0 °C and portion wise NaH (11 g, 455.1 mmol, 6 eq.). The mixture was stirred at 0 °C for 1 h, then PMBCl (41.1 mL, 303.4 mmol, 4 eq.) was added and the reaction stirred at r.t. for 16 h. The reaction was carefully quenched at 0 °C with EtOH, then evaporated, recovered with EtOAc (200 mL) and washed with brine (4 x 250 mL). The organic layer was dried over $MgSO_4$, evaporated and re-crystallised from hot IPA to afford the titled compound **L-5** as a white solid (21 g, 36.9 mmol, 49%): $R_f$ = 0.8 (EtOAc–PE, 4:1); $[\alpha]_D^{25}$ – 2.1 (*c* = 1.00, $CHCl_3$); IR (ATR) 819.66 (arom); $^1H$ NMR ($CDCl_3$, 400 MHz) $\delta$ 7.45–6.66 (m, 12 H, PMB), 4.92–4.64 (m, 6 H, 3 x $OCH_2$), 4.38 (d, J = 9.6 Hz, 1 H, H-1), 3.84–3.80 (m, 9 H, 3 x OMe), 3.77 (d, *J* = 9.5 Hz, 1 H, H-2), 3.57 (d, *J* = 2.6 Hz, 1 H, H-4), 3.53 (dd, *J* = 9.2, 2.8 Hz, 1 H, H-3), 3.45 (q, *J* = 6.3 Hz, 1 H, H-5), 2.85–2.61 (m, 2 H, $SCH_2$), 1.30 (t, *J* = 7.4 Hz, 3 H, $SCH_2CH_3$), 1.17 (d, *J* = 6.3 Hz, 3 H, H-6) *ppm*; $^{13}C$ NMR (CDCl3, 50 MHz) $\delta$ 159.97, 159.46, 159.23 (3 x COMe-Ar), 130.75, 130.64,

130.22 (3 x $CCH_2$-Ar), 129.94, 129.47, 129.12, 113.42, 113.40, 113.28 (12 x CH-Ar), 84.68 (C-1), 74.48, 74.33, 74.29 (3 x $CH_2$), 71.92 (C-5), 73.04 (C-3), 70.58 (C-4), 66.54 (C-2), 55.09, 54.98, 54.67 (3 x OMe), 23.99 ($CH_2$), 16.52 ($CH_3$), 14.47 (C-6) *ppm*; MS ($ESI^-$) m/z = 567.71 [M-H].

**2-Acetamido-3,4,6-tri-*O*-acetyl-2-deoxy-α-D-glucopyranosyl chloride (L-7):**

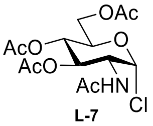

**L-7**

To acetyl chloride (58.1 mL, 817.1 mmol, 9 eq.) was added *N*-acetyl glucosamine (**L-6**) (20 g, 90.4 mmol, 1 eq.) under nitrogen atmosphere and strong stirring. The suspension was stirred for 2 days at r.t., before $CHCl_3$ (200 mL) was added and the mixture was poured into ice (200 g) and $H_2O$ (50 mL). The organic layer was separated and poured into a satd. solution of $NaHCO_3$ (250 mL) and ice (100 g). The mixture was stirred and then shaken into a separatory funnel until the production of gas ceased. The organic phase was then dried over $MgSO_4$ and evaporated to a small volume under vacuum. $Et_2O$ was rapidly added and the product left to crystallise at r.t.. The mother liquor was evaporated and re-crystallised again from $Et_2O$. The product **L-7** was obtained as a white solid (29.07 g, 79.5 mmol, 88%): $R_f$ = 0.9 (EtOAc, 100%); $[\alpha]_D^{25}$ + 118.9 (*c* = 1.04, $CHCl_3$); IR (ATR) 1746.09 (COOR), 1667.69 (CONH); $^1H$ NMR ($CDCl_3$, 400 MHz) *δ* 6.18 (d, *J* = 3.7 Hz, 1 H, H-1), 5.83 (d, *J* = 8.7 Hz, 1 H, NH), 5.32 (t, *J* = 11.9 Hz, 1 H, H-3), 5.21 (t, *J* = 9.8 Hz, 1 H, H-4), 4.53 (ddd, *J* =10.7, 8.8, 3.7 Hz, 1 H, H-2), 4.30–4.24 (m, 2 H, H-5, H-6a), 4.13 (dd, *J* =12.0, 1.6 Hz, 1 H, H-6b), 2.10, 2.05, 1.98 (3 x s, 12 H, 4 x Ac) *ppm*; $^{13}C$ NMR ($CDCl_3$, 101 MHz) *δ* 171.49, 170.58, 170.09, 169.13 (4 x CO), 93.64 (C-1), 70.91 (C-5), 70.14 (C-3), 66.96 (C-4), 61.15 (C-6), 53.51 (C-2), 23.09 (NHAc), 20.71, 20.68, 20.55 (3 x OAc) *ppm*; MS ($ESI^+$) m/z = 366.76 [M+H]. Identical to the previous report (*30*).

**2-Acetamido-3,4,6-tri-*O*-acetyl-2-deoxy-β-D-glucopyranosyl-1-*S*-isothiouronium chloride (L-8):**

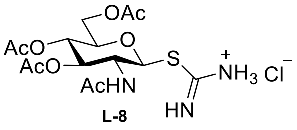

**L-8**

To a solution of **L-7** (29.07 g, 79.5 mmol, 1 eq.) in acetone (200 mL) was added thiourea (12.1 g, 159 mmol, 2 eq.). The mixture was stirred for 1 h at reflux (80 °C). The reaction was cooled to r.t., filtered and the solid washed with cold EtOH, to afford the titled compound **L-8** as a white solid (32.85 g, 74.3 mmol, 93%): $R_f$ = 0.0 (EtOAc, 100%); $[\alpha]_D^{25}$ – 28.0 (*c* = 1.04, $H_2O$); IR (ATR) 3219.6, 3185.60 ($NH_2$),1759.10 (COOR), 1646.90 (CONH); $^1H$ NMR (DMSO-$d_6$, 400 MHz) $\delta$ 9.41, 9.21 (2 x s, 4 H, $NH_2$, $NH_2Cl$), 8.43 (d, *J* = 9.3 Hz, 1 H, NH), 5.68 (d, *J* = 10.5 Hz, 1 H, H-1), 5.13 (t, *J* = 9.8 Hz, 1 H, H-4), 4.93 (t, *J* = 9.7 Hz, 1 H, H-3), 4.24–4.15 (m, 2 H, H-5, H-6a), 4.07–3.97 (m, 2 H, H-2, H-6b), 2.01, 1.98, 1.93, 1.80 (4 x s, 12 H, 4 x Ac) *ppm*; $^{13}C$ NMR (DMSO-$d_6$, 101 MHz) $\delta$ 170.50, 170.36, 170.06, 169.73 (4 x CO), 167.69 ($CNHNH_2$), 81.10 (C-1), 75.18 (C-5), 73.13 (C-3), 68.31 (C-4), 61.93 (C-6), 51.69 (C-2), 22.97 (NHAc), 21.01, 20.85, 20.74 (3 x OAc) *ppm*; MS ($ESI^+$) m/z = 406.42 [M+H]. Identical to the previous report (*31*).

**Cyanomethyl 2-acetamido-3,4,6-tri-*O*-acetyl-2-deoxy-1-thio-β-D-glucopyranoside (L-9):**

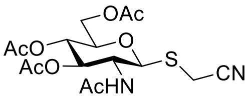


**L-9**

To a solution of **L-8** (20 g, 49.3 mmol, 1 eq.) in $H_2O$:acetone (1:1, 220 mL) were added $Na_2S_2O_5$ (20.6 g, 108.5 mmol, 2.2 eq.), $K_2CO_3$ (8.86 g, 64.1 mmol, 1.3 eq.) and chloroacetonitrile (59.3 mL, 937 mmol, 19 eq.). The mixture was stirred for 16 h at r.t., then it was poured into cold $H_2O$ (400 mL) and stirred for further 1.5 h at r.t.. The reaction was extracted with $CHCl_3$ (3 x 250 mL), the organic layer backwashed with brine (200 mL), dried over $MgSO_4$ and evaporated under vacuum, to afford the titled compound **L-9** as a white solid (17.21 g, 42.8 mmol, 87%): $R_f$ = 0.8 (EtOAc–MeOH, 4:1); $[\alpha]_D^{25}$ – 94.3 (c = 1.0, $CHCl_3$); IR (ATR) 2198.15 (CN), 1738.74 (COOR), 1660.67 (CONH); $^1H$ NMR ($CDCl_3$, 250 MHz) $\delta$ 5.96 (d, *J* = 9.2 Hz, 1 H, NH), 5.20 (t, *J* = 9.4 Hz, 1 H, H-3), 5.09 (t, *J* = 9.4 Hz, 1 H, H-4), 4.79 (d, *J* = 10.4 Hz, 1 H, H-1), 4.16–4.27 (m, 3 H, H-2, H-6), 3.77 (ddd, *J* = 9.2, 4.3, 2.4 Hz, 1 H, H-5), 3.32, 3.67 (2 x d, *J* = 17.0 Hz, 2 H, $CH_2CN$), 2.09, 2.04, 1.96 (3 x s, 12 H, 4 x Ac) *ppm*; $^{13}C$ NMR ($CDCl_3$, 63 MHz) $\delta$ 171.63, 171.17, 171.03 (3 x COO), 169.75 (CONH), 116.67 (CN), 83.44 (C-1), 76.66 (C-5), 73.69 (C-3), 68.49 (C-4), 62.30 (C-6), 53.19 (C-2), 23.58 (NHAc), 21.20, 21.11, 21.05 (3 x OAc), 15.11 ($CH_2CN$) *ppm*; MS ($ESI^+$) m/z = 403.42 [M+H]. Identical to the previous report (*31*).

**Cyanomethyl 2-acetamido-2-deoxy-1-thio-β-D-glucopyranoside (L-10):**

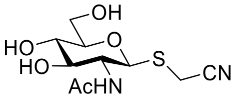

**L-10**

To a solution of **L-9** (23 g, 57 mmol, 1 eq.) in MeOH (445 mL) was added $NEt_3$ (44.5 mL, 319 mmol, 5.6 eq.). The mixture was stirred for 16 h at r.t., then the solvent was removed under vacuum and the slurry obtained co-evaporated twice with toluene and once with $Et_2O$, to afford the titled compound **L-10** as a white solid (19 g, 57 mmol, q.): $R_f$ = 0.2 (EtOAc–MeOH, 4:1); $[\alpha]_D^{25}$ – 64.6 (*c* = 1.00, $H_2O$); IR (ATR) 3428.88 (OH), 2196.43 (CN), 1664.21 (CONH); $^1H$ NMR (DMSO-$d_6$, 400 MHz) $\delta$ 7.89 (d, *J* = 9.2 Hz, 1 H, NH), 5.11 (dd, *J* = 9.0, 5.4 Hz, 2 H, OH-3, OH-4), 4.50 (dd, *J* = 11.3, 4.3 Hz, 2 H, H-1, OH-6), 3.79 (dd, *J* = 41.5, 17.0 Hz, 2 H, $CH_2CN$), 3.74–3.65 (m, 1 H, H-6a), 3.56 (dd, *J* = 19.5, 9.8 Hz, 1 H, H-2), 3.45 (dt, *J* = 11.7, 5.7 Hz, 1 H, H-6b), 3.34–3.29 (m, 1 H, H-3), 3.12 (ddd, *J* = 13.7, 9.0, 3.5 Hz, 2 H, H-4, H-5) *ppm*; $^{13}C$ NMR (DMSO-$d_6$, 125 MHz) $\delta$ 169.31 (CO), 118.10 (CN), 83.11 (C-1), 81.44 (C-5), 74.97 (C-3), 70.45 (C-4), 61.29 (C-6), 54.07 (C-2), 22.90 (Ac), 13.79 ($CH_2CN$) *ppm*; MS ($ESI^+$) m/z = 277.32 [M+H]. Identical to the previous report (*31*).

**Cyanomethyl 2-acetamido-6-*O*-*tert*-butyldimethylsilyl-2-deoxy-1-thio-β-D-glucopyranoside (L-11):**

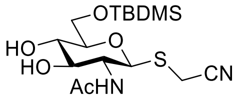

**L-11**

To a solution of **L-10** (8.2 g, 29.7 mmol, 1 eq.) in DMF (60 mL) at 0 °C, were added imidazole (3.03 g, 44.5 mmol, 1.5 eq.) and TBDMSCl (4.92 g, 32.6 mmol, 1.1 eq.). The mixture was stirred for 16 h at r.t., then the solvent was removed under vacuum and the oil obtained precipitated with $H_2O$. The precipitate was filtered, washed with $H_2O$ and co-evaporated with toluene and acetone, to afford the titled compound **L-11** as a white solid (7.72 g, 19.8 mmol, 67%): $R_f$ = 0.85 (EtOAc–MeOH, 9:1); $[\alpha]_D^{25}$ – 89.8 (*c* = 1.00, acetone); IR (ATR) 3291.99 (OH), 2198.37 (CN), 1649.35 (CONH); $^1H$ NMR (Acetone-$d_6$, 400 MHz) $\delta$ 7.32 (d, *J* = 9.0 Hz, 1 H, NH), 4.69 (d, *J* = 10.3 Hz, 1 H, H-1), 4.01 (dd, *J* = 11.3, 2.0 Hz, 1 H, H-6a), 3.89–3.78 (m, 2 H, H-2, H-6b), 3.69 (dd, *J* = 37.0, 17.6 Hz, 2 H, $CH_2CH$), 3.62 (dd, *J* = 10.5, 7.9 Hz, 1 H, H-3), 3.51–3.42 (m, 1 H, H-4), 3.35 (ddd, *J* = 9.7, 5.4, 2.0 Hz, 1 H, H-5), 1.89 (s, 3 H, Ac), 0.91 (s, 9 H, $^t$Bu), 0.10, 0.09 (2 x s, 2 x 3 H, 2 x $CH_3$) *ppm*; $^{13}C$ NMR (Acetone-$d_6$, 100 MHz) $\delta$ 169.74 (CO), 116.87 (CN), 83.29 (C-1), 81.15 (C-5), 76.10 (C-3), 70.65 (C-4),

62.90 (C-6), 54.39 (C-2), 25.41 ($C(CH_3)_3$), 22.25 (Ac), 18.10 ($C(CH_3)_3$), 13.61 ($CH_2CN$), -5.86, -5.96 (2 x $CH_3$) *ppm*; MS ($ESI^+$) m/z = 391.58 [M+H].

**Cyanomethyl 2,3,4,6-tetra-*O*-acetyl-β-D-galactopyranosyl-(1→4)-2-acetamido-6-*O*-*tert*-butyldimethylsilyl-2-deoxy-1-thio-β-D-glucopyranoside (L-12):**

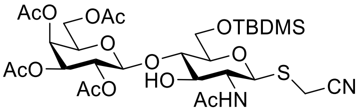

**L-12**

A mixture of **L-11** (1 g, 2.6 mmol, 1.0 eq.) and **L-2** (1.4 g, 2.8 mmol, 1.1 eq.) was co-evaporated with toluene, dried under high vacuum and suspended in anhydrous DCM (7 mL) with MS (4 Å, 2.5 g) and stirred for 1 h. The mixture was cooled to -41 °C and stirred for 15 min prior to addition of boron trifluoride diethyl etherate (370 µL, 3.0 mmol, 1.15 eq.). The solution was stirred at -41 °C for 1 h before triethylamine (1 mL) was added and the crude mixture was directly purified by flash column chromatography over silica (EtOAc–PE, 1:1 to 1:0) to afford the titled disaccharide **L-12** as a white solid (776.7 mg, 1.076 mmol, 42%): $R_f$ = 0.5 (EtOAc, 100%); $[\alpha]_D^{25}$ – 43.3 (*c* = 1.00, $CHCl_3$); IR (ATR) 3347.95 (OH), 2201.76 (CN), 1732.17 (CO),1693.87 (CONH); $^1H$ NMR ($CDCl_3$, 400 MHz) $\delta$ 5.97 (d, *J* = 8.9 Hz, 1 H, NH), 5.42–5.32 (m, 1 H, H-4'), 5.20 (dd, *J* = 10.5, 8.0 Hz, 1 H, H-2'), 4.96 (dd, *J* = 10.5, 3.4 Hz, 1 H, H-3'), 4.67 (d, *J* = 10.3 Hz, 1 H, H-1), 4.61 (d, *J* = 8.1 Hz, 1 H, H-1'), 4.24–4.05 (m, 2 H, H-6'), 3.99 (dd, *J* = 7.4, 6.3 Hz, 1 H, H-5'), 3.86 (dd, *J* = 20.0, 10.1 Hz, 2 H, H-2, H-6a), 3.70 (ddd, *J* = 21.0, 10.0, 5.1 Hz, 3 H, H-6b, H-3, H-4), 3.56 (d, *J* = 16.9 Hz, 1 H, $CH_2$aCN), 3.38 (dd, *J* = 9.1, 1.8 Hz, 1 H, H-5), 3.32 (d, *J* = 16.9 Hz, 1 H, $CH_2$bCN), 2.14, 2.06, 2.04, 2.01, 1.96 (5 x s, 5 x 3 H, 5 x Ac), 0.90 (s, 9 H, $^t$Bu), 0.07, 0.06 (2 x s, 2 x 3 H, 2 x $CH_3$) *ppm*; $^{13}C$ NMR ($CDCl_3$, 100 MHz) $\delta$ 170.90, 170.53, 170.09, 170.00, 169.26 (5 x CO), 116.38 (CN), 101.46 (C-1'), 82.61 (C-1), 80.32 (C-5), 78.84 (C-3), 73.41 (C-5'), 71.32 (C-3'), 70.80 (C-4'), 68.65 (C-4), 66.77 (C-2'), 61.32 (C-6'), 60.42 (C-6), 54.19 (C-2), 25.82 ($C(CH_3)_3$), 23.32 (NHAc), 20.70, 20.60, 20.55, 20.52 (4 x OAc), 18.21 ($C(CH_3)_3$), 14.31 ($CH_2CN$), -4.98, -5.22 (2 x $CH_3$) *ppm*; MS ($ESI^+$) m/z = 721.87 [M+H].

**Cyanomethyl 2,3,4-tri-*O*-*p*-methoxybenzyl-α-L-fucopyranosyl-(1→3)-[2,3,4,6-tetra-*O*-acetyl-β-D-galactopyranosyl-(1→4)]-2-acetamido-6-*O*-*tert*-butyldimethylsilyl-2-deoxy-1-thio-β-D-glucopyranoside (L-13):**

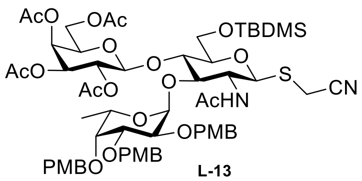

L-13

**L-12** (380 mg, 0.527 mmol, 1.0 eq.) and **L-5** (599.6 mg, 1.054 mmol, 2 eq.), co-evaporated with toluene and dried under high vacuum for 1h, were dissolved in DCM:DMF (dry, 1:1, 1.5 mL). To the mixture were added MS (4 Å, 790 mg) and it was stirred for 1 h at r.t. before copper(II) bromide (235.4 mg, 1.054 mmol, 2.0 eq.) and tetra-butylammonium bromide (356.8 mg, 1.107 mmol, 2.1 eq.) were added too. The mixture was stirred at r.t. for 16 h in the dark, then filtered over Celite and washed with DCM (200 mL). The filtrate was washed with brine (150 mL) and $NaHCO_3$ satd. solution (6 x 120 mL). The aqueous layer was re-extracted with DCM (200 mL) and the combined organic layers where dried over $MgSO_4$, filtered, concentrated in vacuum and directly purified by flash column chromatography over silica (PE–EtOAc, 1:1 to 7:3) to afford the titled trisaccharide **L-13** as a white foam (513.3 mg, 0.418 mmol, 79%): $R_f$ = 0.1 (PE–EtOAc, 1:1); $[\alpha]_D^{25}$ – 60.3 (*c* = 1.00, $CHCl_3$); IR (ATR) 2211.43 (CN), 1716.94 (CO),1696.88 (CONH); $^1$H NMR ($CDCl_3$, 400 MHz) *δ* 7.35-6.77 (m, 12 H, CH-arom), 6.22 (d, *J* = 7.9 Hz, 1 H, NH), 5.36 (dd, *J* = 3.4, 0.8 Hz, 1 H, H-4’), 5.07 (dd, *J* = 10.6, 7.9 Hz, 2 H, H-2’, H-1’’), 4.93 (dd, *J* = 10.5, 3.5 Hz, 1 H, H-3’), 4.88 (d, *J* = 11.4 Hz, 1 H, $CH_2$-PMB), 4.77 (dd, *J* = 9.4, 8.0 Hz, 3 H, $CH_2$-PMB, H-1, H-1’), 4.70 (s, 1 H, $CH_2$-PMB), 4.66 (d, *J* = 11.1 Hz, 1 H, $CH_2$-PMB), 4.61 (d, J = 11.5 Hz, 1 H, $CH_2$-PMB), 4.33 (d, *J* = 6.4 Hz, 1 H, H-5’’), 4.17 (dd, *J* = 11.0, 7.8 Hz, 1 H, H-6’a), 4.11–4.05 (m, 1 H, H-2’’), 4.04–3.97 (m, 3 H, H-6’b, H-3, H-5), 3.89–3.76 (m, 14 H, H-2, H-6, H-5’, H-3’’, 3 x $OCH_3$), 3.65 (d, *J* = 1.6 Hz, 1 H, H-4’’), 3.45 (d, *J* = 17.1 Hz, 1 H, $CH_2$aCN), 3.40–3.32 (m, 1 H, H-4), 3.14 (d, *J* = 17.0 Hz, 1 H, $CH_2$bCN), 2.05, 2.01, 1.96, 1.94 (4 x s, 4 x 3 H, 4 x OAc), 1.77 (s, 3 H, NHAc), 1.15 (d, *J* = 6.5 Hz, 3 H, H-6’’), 0.90 (s, 9 H, $^t$Bu), 0.07, 0.05 (2 x s, 2 x 3 H, 2 x $CH_3$) *ppm*; $^{13}$C NMR ($CDCl_3$, 100 MHz) *δ* 170.50, 170.12, 170.03, 170.00, 169.42 (5 x CO), 159.36, 159.17, 159.02 (3 x $COCH_3$-arom), 130.81, 130.65, 130.52 (3 x C-arom), 129.99, 129.95, 128.77 (3 x CHC-arom), 116.80 (CN), 113.93, 113.76, 113.58 (3 x CHCO-arom), 99.19 (C-1’), 98.34 (C-1’’), 82.02 (C-1), 79.97 (C-4), 79.61 (C-3’’), 76.48, 76.27, 76.04 (C-4’’, C-2’’, C-3), 74.01 , 73.78 (2 x $OCH_2$), 72.97 (C-5), 72.21 ($OCH_2$), 71.29 (C-3’), 70.69 (C-5’), 68.97 (C-2’), 66.79 (C-5’’), 61.18 (C-4’), 60.57 (C-6), 60.39 (C-6’), 55.28, 55.25, 55.22 (3 x $OCH_3$), 55.18 (C-2), 25.84 ($C(CH_3)_3$), 23.09 (NHAc), 21.05, 20.83, 20.61, 20.55 (4 x OAc), 18.17 ($C(CH_3)_3$), 16.78 (C-6’’), 14.22 ($CH_2CN$), -5.07, -5.33 (2 x $SiCH_3$) *ppm*; MS (ESI$^+$) m/z = 1228.45 [M+H].

**Cyanomethyl 2,3,4-tri-*O*-acetyl-α-L-fucopyranosyl-(1→3)-[2,3,4,6-tetra-*O*-acetyl-β-D-galactopyranosyl-(1→4)]-2-acetamido-6-*O*-acetyl-2-deoxy-1-thio-β-D-glucopyranoside (L-14):**

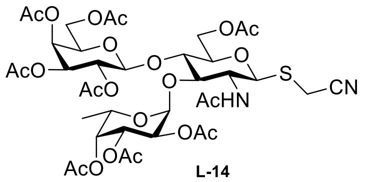

L-14

To **L-13** (531.9 mg, 0.433mmol, 1.0 eq) dissolved in THF (dry, 3.5 mL) was added triethylamine trihydrofluoride (506 µL, 3.10 mmol, 7 eq). The solution was stirred at r.t. for 16 h. The solvent was evaporated and the crude mixture was directly used without purification. To the solution of the crude residue above in ACN:$H_2O$ (9:1, 10 mL) was added CAN (2.87 g, 5.23 mmol, 9 eq.). The solution was stirred at r.t. for 2 h, then $Ac_2O$ (30 mL), pyridine (60 mL) and DMAP (7.1 mg, 0.058 mmol, 0.1 eq.) were added. The reaction was stirred at r.t. for 16 h. The solvents were evaporated, the crude mixture was co-evaporated with toluene and purified by flash column chromatography on silica (EtOAc–PE, 1:1 to 1:0) to afford the title compound **L-14** as a white powder (325 mg, 0.34 mmol, 59%): $R_f$ = 0.5 (PE–EtOAc, 1:1); $[\alpha]_D^{25}$ – 123.8 (*c* = 1.00, $CHCl_3$); IR (ATR) 2206.78 (CN), 1743.13 (CO); $^1$H NMR ($CDCl_3$, 400 MHz) *δ* 5.86 (d, *J* = 9.8 Hz, 1 H, NH), 5.44 (d, *J* = 4.0 Hz, 1 H, H-1''), 5.42 (d, *J* = 3.4 Hz, 1 H, H-4'), 5.39 (d, *J* = 2.7 Hz, 1 H, H-4''), 5.18 (dd, *J* = 11.0, 3.3 Hz, 1 H, H-3''), 5.09 (dd, *J* = 10.3, 8.0 Hz, 1 H, H-2'), 5.00 (dd, *J* = 10.3, 3.7 Hz, 1 H, H-3'), 5.00 (dd, *J* = 10.0, 3.5 Hz, 1 H, H-2''), 4.92 (q, *J* = 6.5 Hz, 1 H, H-5''), 4.70 (dd, *J* = 12.2, 2.1 Hz, 2 H, H-1, H-6a), 4.52 (dd, *J* = 16.5, 7.1 Hz, 2 H, H-6'a, H-1'), 4.32 (dd, *J* = 11.5, 7.9 Hz, 1 H, H-6'b), 4.07 (m, 2 H, H-6b, H-2), 3.94–3.85 (m, 3 H, H-, H-3, H-5'), 3.61 (d, *J* = 17.1 Hz, 1 H, $CH_2$aCN), 3.55 (m,1 H, H-5), 3.28 (d, *J* = 17.1 Hz, 1 H, $CH_2$bCN), 2.20, 2.15, 2.15, 2.10, 2.09, 2.07, 1.99, 1.98, 1.97 (9 x s, 9 x 3 H, 9 x Ac), 1.21 (d, 3 H, *J* = 5.0, H-6'') *ppm*; $^{13}$C NMR (100 MHz, $CDCl_3$) *δ* 171.42, 170.90, 170.78, 170.66, 170.50, 170.31, 170.08, 169.84, 169.16 (9 x CO), 116.42 (CN), 100.59 (C-1'), 95.64 (C-1''), 82.97 (C-1), 77.58 (C-5), 74.90 (C-3), 74.24 (C-4), 71.34 (C-4''), 71.20 (C-2''), 70.97 (C-2), 68.95 (C-2'), 68.06 (C-3'), 67.91 (C-3''), 66.67 (C-4'), 64.49 (C-5'), 64.43 (C-5''), 61.41 (C-6), 60.69 (C-6'), 25.40 (C-6''), 23.38 (NHAc), 20.97, 20.93, 20.90, 20.86, 20.78, 20.70, 20.65, 20.57 (8 x Ac), 14.32 ($CH_2$CN) *ppm*; MS (ESI$^+$) m/z = 921.28 [M+H].

**2-Imino-2-methoxyethyl α-L-fucopyranosyl-(1→3)-[β-D-galactopyranosyl-(1→4)]-2-acetamido-2-deoxy-1-thio-β-D-glucopyranoside ($Le^x$-IME):**

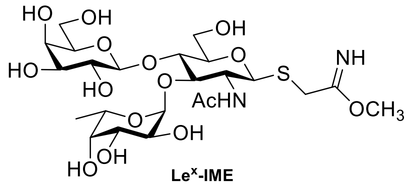

$Le^x$-IME

Cyano 14 (51 mg, 55.4 μmol) was suspended in dry $CH_3OH$ (554 μL) to which was added $CH_3ONa$ (57 L, 25% in $CH_3OH$, 0.025 mM, 0.4 eq.). The reaction was left stirring at r.t. for 16 h, monitored for production formation by mass analysis. The reaction was quenched by solvent evaporation under nitrogen flow followed by 2 h in high vacuum, giving **$Le^x$-IME** together with cyano **L-14** as a 1:1 mixture: $R_f$ = 0.57 (Acetone–$H_2O$, 8:2); $^1$H NMR (500 MHz, $CD_3OD$) $\delta$ 5.08 (d, $J_{H''-1,H''-2}$ = 4.0 Hz, 1H, H″-1), 4.86 (m, 1H, H″-5), 4.46 (d, $J_{H-1,H-2}$ = 10.2 Hz, 1H, H-1 **$Le^x$-IME**), 4.75 (d, $J_{H-1,H-2}$ = 10.0 Hz, 1H, H-1 **L-14**), 4.10 (m, 1H, H-2), 3.98 (m, 3H, H″-3, H-3, H-4), 3.93 (d, *J* = 9.4 Hz, 1H, H-6), 3.90 (m, 3H, H-2, H-5, H′-5), 3.84 (m, 1H, SC*H*HCN), 3.79 (m, 1H, H′-6a), 3.74 (m, 1H, H′-4), 3.70 (m, 3H, H″-4, H-2, H′-6b), 3.64 (d, *J* = 17.0 Hz, 1H, SCH*H*CN), 3.55 (m, 4H, H″-2, H′-2, H-5, H′-3), 1.98 (s, 3H, $CH_3$CONH), 1.21 (d, *J* = 6.6 Hz, 3H, $CH_3$) *ppm*; $^{13}$C NMR (125 MHz, $CD_3OD$) δ 173.7 ($CH_3$*C*ONH), 118.6 ($SCH_2$*C*N), 104.0 (*C*-1′), 100.6 (*C*-1″), 84.6 (*C*-1), 81.9 (*C*-5), 76.8 (*C*-3), 75.0 (*C*-5′), 73.0 (*C*-4), 72.8 (*C*-3′), 71.2 (*C*-3″), 69.7 (*C*-5″), 62.9 (*C*-6′), 61.4 (*C*-6), 55.9 (*C*-2), 22.9 (*C*$H_3$CONH), 16.6 (*C*-6″) *ppm*; HRMS (ESI): *m*/*z* calcd for $C_{23}H_{40}N_2NaO_{15}S$ $[M+Na]^+$ 639.2042. Found: 639.2045.

**Lys–C(NH)NH-GM3g**

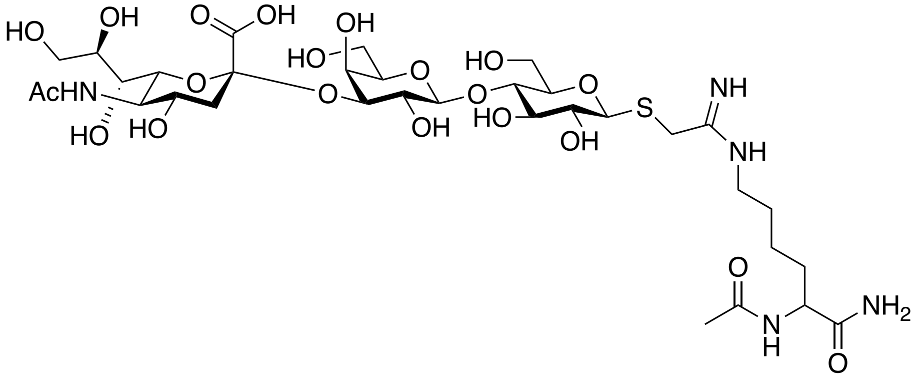


To a solution of SiaLac-IME (90 mg, 0.125 mmol) and Ac-Lys-$NH_2$ (36.2 mg, 0.162 mmol) in dry MeOH (10 mL) was added triethylamine (52 □L, 0.375 mmol) and the mixture stirred at rt for 24 h after which time the solvent was removed under vacuum and resuspend and water and lyophilised. The crude mixture was purified by LH20 twice, eluting with water to give the product as a white solid. Rf: baseline EtOAc:IPA:$H_2O$:AcOH 2:2:1:1; LRMS: m/z (ES+) 876 (100%, $[M+H]^+$), 877 (40%, $[M+H]^+$); HRMS: m/z (ES+) calculated for $C_{33}H_{58}N_5O_{20}S$ $[M+H]^+$ 876.3370; observed 876.3390; IR $\nu_{max}$ 3286 (OH), 2936, 1617 ($COO^-$), 1558, 1432, 1395, 1377, 1317, 1293, 1241, 1212, 1070, 1034, 945, 896, 815, 779, 680, 607.

$^1H$ NMR (950 MHz, $D_2O$) δ 4.66 (d, J = 10.0 Hz, 1H), 4.57 (d, J = 7.8 Hz, 1H), 4.26 (dd, J = 9.1, 5.2 Hz, 1H), 4.14 (dd, J = 9.9, 3.2 Hz, 1H), 4.01 – 3.95 (m, 2H), 3.94 – 3.83 (m, 4H), 3.81 – 3.58 (m, 10H), 3.45 (dd, J = 9.9, 9.0 Hz, 1H), 3.36 (t, J = 7.0 Hz, 2H), 2.79 (dd, J = 12.4, 4.7 Hz, 1H), 1.88 (ddt, J = 15.7, 11.2, 5.9 Hz, 1H), 1.83 (t, J = 12.2 Hz, 1H), 1.78 (dtd, J = 14.0, 9.8, 5.0 Hz, 1H), 1.73 (ddt, J = 17.0, 9.7, 7.0 Hz, 1H), 1.52 (dtt, J = 15.4, 10.5, 5.5 Hz, 1H), 1.50 – 1.42 (m, 1H).

$^{13}C$ NMR (239 MHz, $D_2O$) δ 177.04, 175.03, 174.38, 173.82, 165.31, 102.56, 99.78, 85.03, 78.79, 78.72, 77.62, 75.54, 75.51, 75.22, 72.87, 71.78, 71.70, 69.33, 68.29, 68.07, 67.44, 62.59, 61.04, 59.96, 53.44, 53.37, 51.65, 42.29, 39.65, 39.14, 30.45, 30.36, 26.15, 22.40, 22.00, 21.68, 21.63.

**Neu5Ac-Gal**

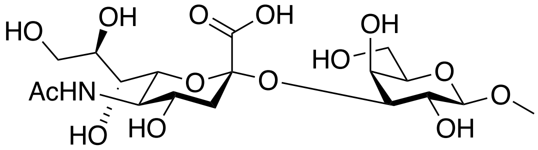


To a solution of methyl beta-D-galactopyranoside (58 mg, 0.299 mmol) in Buffer (20 mL, 100 mM (NH4)2CO3, 20 mM MgCl2, pH 8.5) was added CTP disodium salt ( 395 mg, 0.747 mmol), Neu5Ac (97 mg, 0.314 mmol), Pasteurella multocida Sialyltransferase (2.5 μg per mg of acceptor, 3.52 mg/mL stock) and Neisseria meningitides CMP-sialic acid synthetase (3 μg per mg of acceptor, 2.52 mg/mL stock) and the reaction shaken at 37 °C for 18 h. The solution was concentrated and lyophilised to give the crude solid which was subjected to LH-20 size exclusion and purified as required. This protocol was adapted (*1*).

$^1H$ NMR (950 MHz, $D_2O$) δ 4.32 (1H d, *J* = 8.3 Hz, H-1a), 4.02 (1H, dd, *J* = 10.3, 3.1 Hz, H-3a), 3.88 (1H, d, *J* = 3.3 Hz, H-4a), 3.83 – 3.42 (20H, m), 3 .50 (3H, s, OMe) 2.69 (1H, dd, *J* = 13.0, 4.9 Hz, H-3eq), 1.96 (3H, s, NAc), 1.72 (1H, t, *J* = 12.7 Hz, H-3ax).

### DOCUMENT S2: FURTHER DISCUSSION OF STRUCTURAL ANALYSES OF BAR1 Fab

The crystal contains two complete copies of the complex which are largely identical (rmsd of light chain 0.5 Å). In both the electron density is well ordered for all three sugar rings and the amidine of Lys–C(NH)NH-GM3g, but less well ordered for the lysine (**Fig S12c,d**). The terminal sialic acid makes direct interactions confirming an important role in antibody binding. The sialic acid makes five hydrogen bonds to antibody backbone atoms, the most striking being the bidentate interaction of the sugar carboxylate and with the amide nitrogen atoms of $Ala58_H$ and $Val59_H$. Both the glycerol and N-acetyl moieties of sialic acid are anchored by hydrogen bonds to the protein main-chain. Several highly coordinated water molecules (W) contribute to the binding of the glycan. W1 interacts with $O_9$ of the sialic acid glycerol moiety and residues $Gly38_H$, $His40_H$, $Tyr103_H$. The sialic N-acetyl group is also stabilised by a hydrophobic pi-CH interaction with $Phe37_H$. Notably the indole of $Trp57_H$ stacks against the alpha-face of the Gal sugar of GM3g; the classical pi-CH interaction found in diverse so-called carbohydrate modules (CBMs) (*32, 33*). The conformer of C5-C6-O6 of the Gal residue differs between the two copies. In one copy O6 hydrogen interacts with two highly coordinated water molecules W2 (bridging with $Ser61_H$ and $Asn63_H$) and W3 ($Tyr99_H$ and $Asn63_H$). In the other copy the side chain proximity with the cell symmetric modifies the water networking and the hydroxyl interacts directly with $Asn63_L$ and a water molecule.

The glucose ring of Lys–C(NH)NH-GM3g makes hydrogen bonds to three water molecules, two of which bridge to the protein (including W3 which bridges to $Tyr99_L$, $Asn63_H$ and galactose) but only three direct van der Waal contacts with the protein. These water molecules are found in both copies in the crystallographic asymmetric unit, with additional water molecules in the second copy possibly due to differences in crystal packing. The amidine linkage of Lys–C(NH)NH-GM3g makes hydrogen bonds to the protein ($Tyr97_L$) and to a water molecule (W5) that bridges to the glucose. The amidine also forms a cation-pi interaction with $Tyr37_L$ confirming the contribution of the amidine linker in the binding. The aliphatic side chain of the lysine portion of the ligand makes van der Waal contacts with $Tyr97_L$. The terminal amides of lysine make long (2.9 Å) hydrogen bonds to the side chain of $Thr98_L$.

The interactions of the tip disaccharide of GM3g, and their positions relative to BAR1 Fab (including water molecules) are similar to that described for the stage-specific embryonic antigen-4 (SSEA-4) headgroup (a hexasaccharide) (RCSB 6ug7/6ug8) antibody complex (*34*). Although a different antibody to that described here, the interacting residues in CDRH1 and CDRH2 in both structures are largely conserved (**Fig S13**). However, in the hexasaccharide there is an N-acetylgalactosamine (NGA) in the third position, in contrast to glucose in the ligand studied here. The NGA makes hydrogen bonds and multiple van der Waal interactions with the antibody, as well as hydrogen bonds to bridging water molecules (including one like W3). The glucose and the NGA rings are offset by a 90° rotation ($\varphi$=265) as a result the hexasaccharide forms a U-shape whereas the Lys–C(NH)NH-GM3g adopts a more linear arrangement ($\varphi$=47/57). The hexasaccharide engages with the other loops in the ch28/11 alternative light chain, which would otherwise clash with the BAR-1 light chain.

### DOCUMENT S3: SUPPLEMENTARY GLYCAN MICROARRAY DOCUMENT

#### Supplementary Glycan Microarray Document

#### Based on MIRAGE Guidelines (doi:10.3762/mirage.3)

| Classification | Guidelines |
|---|---|
| **1. Sample: Glycan Binding Sample** | |
| Description of Sample | Sera from mice (diluted 1:200 in 1X PBS + 0.05% Tween-20 |
| Sample modifications | N/A |
| Assay protocol | Please see method section in the main text. |
| **2. Glycan Library** | |
| Glycan description for defined glycans | In-house sialoside array, consisting of 137 defined glycans (Supplementary Table 1). The synthesis of the contained glycans are described in Supplemental Experimental Procedures in (Peng et al., 2017). |
| Glycan description for undefined glycans | No glycans are undefined. |
| Glycan modifications | No modifications after initial synthesis were made. |
| **3. Printing Surface; e.g., Microarray Slide** | |
| Description of surface | NHS-ester functionalized hydro-polymer. |
| Manufacturer | Schott SlideH (Applied Microarrays 1070936). |
| Custom preparation of surface | None |
| Non-covalent Immobilization | All glycans are terminated with primary amine linker (either natural amino acid or chemical linker). |
| **4. Arrayer (Printer)** | |
| Description of Arrayer | MicroGrid II (Digilab) |
| Dispensing mechanism | Contact microarray pins (SMP3, ArrayIt) |
| Glycan deposition | Manufacturer estimation is 0.7nL per spot. However, actual delivery volume of each printed spot is not determined.<br>Each glycan was "pre-spotted" 3 times on Poly-L-Lysine derivatized slides (made in-house) before being spotted on SlideH slides. Each array contains 6 replicate spots of each individual glycan. |

| | |
|---|---|
| Printing conditions | Glycans were diluted to 100uM in 150mM NaPO4 buffer, pH 8.4 + 0.005% Tween-20. 10uL of each glycan was transferred to a 384-well microtiter plate and printed at ambient temperature and relative humidity of 50-65%. |
| **5. Glycan Microarray with "Map"** | |
| Array layout | Each slide contains 3 replicate arrays, consisting of a 4x4 (16) subarray pattern with each subarray containing 12x18 features (not all features contain a printed sample).<br>Array Layout file = "SA.GAL" |
| Glycan identification and quality control | In-house sialoside array, consisting of 137 defined glycans (Supplementary Table 1).<br>Quality control was assessed by incubation with plant lectins, AAL, ECA and SNA, to monitor fucosylations, de-sialylation and NeuAc-α2-6 terminated glycans, respectively. See Supplemental Experimental Procedures 2 in (Peng et al., 2017). |
| **6. Detector and Data Processing** | |
| Scanning hardware | Innoscan 1100AL (Innopsys) |
| Scanner settings | Scanning resolution: 10 µm / pixel<br>Laser channel: 488<br>PMT Voltages: Adjusted for each sample to achieve maximum signal without saturation of any single spot.<br>Scan power: Adjusted for each sample to achieve maximum signal without saturation of any single spot. |
| Image analysis software | Mapix (Innopsys) |
| Data processing | Output .txt files containing calculated data were processed in MS Excel to determine the mean signal value of 6 replicate spots with highest and lowest signals removed (e.g. average of 4 spots). |
| **7. Glycan Microarray Data Presentation** | |
| Data presentation | The microarray binding results are in **Figure X,** and **Supplementary Figure SX**. Binding results are presented as 2D bar graphs with bars representing averaged mean signal of each glycan and error bars representing standard deviation. |
| **8. Interpretation and Conclusion from Microarray Data** | |
| Data interpretation | No software or algorithms were used to interpret processed data. |
| Conclusions | Mice immunized with sialyl-containing antigens are capable to mount an immune response to specific glycan containing epitopes. |
